# Elucidating Molecular Features of White Matter Hyperintensities in Alzheimer’s Disease through Multimodal Imaging and SHAP Analysis

**DOI:** 10.64898/2026.06.18.733283

**Authors:** Claire F. Scott, Cody R. Marshall, Lukasz G. Migas, Léonore E.M. Tideman, Madeline E. Colley, Katerina V. Djambazova, Wilber Romero-Fernandez, Matthew S. Schrag, Raf Van de Plas, Jeffrey M. Spraggins

## Abstract

White matter hyperintensities (WMHs) are a common feature of Alzheimer’s disease and are associated with cognitive decline, yet their molecular composition and spatial heterogeneity remain incompletely defined. Here, we identify distinct lipid signatures in human AD WMHs compared to matched normal-appearing white matter (NAWM) from the same donors. Using an integrated multimodal approach combining magnetic resonance imaging, matrix-assisted laser desorption/ionization imaging mass spectrometry (MALDI IMS), histological staining, and complementary liquid chromatography–tandem mass spectrometry, we resolve spatially localized lipid alterations within tissue sections while preserving spatial context. This approach reveals region-specific heterogeneity in WMH lipid composition across anterior and posterior brain regions that may be obscured by bulk lipidomics alone. Machine learning–based analysis using Shapley additive explanations (SHAP) identified lipid features that contribute to WMH classification, with sulfatide, hexosylceramide, and phosphatidylinositol species emerging as key discriminators. In anterior brain regions, WMHs were associated with differential abundance and depletion of specific sulfatide and hexosylceramide species (SHexCer 44:2;3O, SHexCer 42:2;3O, HexCer 41:1;3O, SHexCer 42:2;2O), whereas posterior WMHs were characterized by reduced phosphatidylinositol species (PI 36:1), demonstrating that AD-associated white matter pathology is governed by region-specific, heterogeneous lipid remodeling rather than uniform global degradation.

## Introduction

As the number of patients living with Alzheimer’s disease (AD) continues to increase, the need to better understand its pathological features becomes more pressing.^1^ To gain insights into neurodegeneration in AD and potential therapeutic targets, numerous studies have characterized the hallmark pathologies associated with the disease. These include well-known neurofibrillary tangles and amyloid-β plaques, as well as other features such as white matter hyperintensities (WMH), dystrophic neurites, vascular pathology, and oxidative damage.^2–6^ AD-associated neuropathologies comprise diverse cellular and molecular microenvironments, some of which remain far less characterized than others, such as WMH. Clinicians identify and diagnose WMHs using T2-weighted Fluid-Attenuated Inversion Recovery (T2-FLAIR) magnetic resonance imaging (MRI), in which they appear as abnormally bright foci within the cerebral white matter. The etiology of WMHs remains a subject of investigation. While historically attributed primarily to cerebral small vessel disease (cSVD) and associated chronic ischemia, emerging evidence suggests that WMHs are not exclusively vascular in origin. Clinically, WMH burden correlates with AD diagnosis and cognitive decline independent of amyloid-β plaque burden and vascular risk factors. Furthermore, histological examination of WMH frequently reveals primary neurodegenerative signatures, including astrogliosis, microglial activation, and demyelination, even in the absence of significant vascular pathology.^7–17^ Together, these findings suggest that in the context of AD, WMHs may represent a dynamic biochemical transition rather than a static consequence of ischemic injury. Strikingly, white matter changes, including WMH, have been shown to appear up to 22 years before the onset of clinical symptoms in patients with familial AD.^18^ Amyloid-β plaques and neurofibrillary tangles have also been detected years before the onset of clinical cognitive AD symptoms and are associated with increased WMH burden and +demyelination.^19^ Proposed mechanisms link increased demyelination to the accumulation of amyloid-β plaques and neurofibrillary tangles through a process in which microglia are metabolically exhausted by processing myelin debris and become dysfunctional at clearing plaques and tangles. Deciphering the exact molecular features driving these early white matter changes is therefore critical to identifying localized therapeutic targets before clinical symptom onset.

Hematoxylin and eosin (H&E) staining can be used in tandem with MRI to define the tissue microenvironment within and surrounding WMH. Central to WMH pathology is demyelination, characterized by damage to the lipid-dense myelin sheaths that insulate neuronal axons and are the defining feature of neural white matter. Oligodendrocytes produce and maintain myelin, which is essential for efficient axon signal propagation. While demyelination is a natural process with age, and human brains have systems for replenishing lost myelin, the extent of demyelination in AD is abnormal.^16,20,21^

Myelin has a much higher lipid-to-protein ratio than most other membranes in the body (70-85% lipid to 15-30% protein, compared to 50:50 in other membranes).^22^ The balance of lipids within oligodendrocytes and myelin is tightly regulated under normal conditions.^15,17,22–24^ However, lipid dysregulation is a known feature of AD.^25,26^ Sulfatide, a sulfated galactosphingolipid largely confined to myelin, together with its precursor galactosylceramide, undergoes a substantial and early depletion reported across post-mortem brain tissue and cerebrospinal fluid studies of AD. However, how these alterations vary across brain regions remains largely unexplored.^27,28^ Liquid chromatography-tandem mass spectrometry (LC-MS/MS) analyses of human AD CSF samples and homogenized whole-brain tissue show global lipid alterations in AD compared to controls. However, despite the broad molecular coverage of LC- MS/MS, these findings lack spatial specificity and therefore cannot be interpreted in the specific context of WMH microenvironments. Classical histopathological and clinical imaging of human brain tissue can be complemented by untargeted lipid imaging, such as matrix-assisted laser desorption ionization imaging mass spectrometry (MALDI IMS), to reveal molecular landscapes associated with WMHs.

MALDI IMS is a highly multiplexed molecular imaging technology that enables the simultaneous mapping of hundreds of lipids with spatial specificity. Rather than requiring tissue homogenization, as in LC-MS/MS, whole tissue sections mounted on glass slides are interrogated pixel by pixel, and the relative intensity of each lipid is displayed as an ion image.^29,30^ One example of the spatial imaging strength of MALDI IMS is demonstrated by Koutarapu et al., who used this technique, guided by immunostaining, to show that amyloid-β plaques in human AD brain tissue are associated with neuronal dystrophy.^31^ They found that these plaques exhibit distinct patterns of amyloid-β truncation and chemical modification compared with other cored plaque deposits. Additionally, Shariatgorji et al. have employed MALDI IMS to visualize changes in the locations and concentrations of neurotransmitters in the brain tissues of animal models following drug treatment.^32^ Given the complexity of AD and the variety of neuropathologies involved, multimodal molecular imaging approaches are essential for characterizing both the molecular and cellular changes associated with AD.

Multimodal MALDI IMS workflows are therefore well-suited to characterize the altered lipid microenvironments within tissue regions affected by WMH. Using a multimodal workflow integrating post-mortem T2-FLAIR MRI, MALDI IMS, LC-MS/MS, and H&E staining, we characterize the lipid microenvironments of WMH and normal-appearing white matter (NAWM) in the AD brain. MRI guided the selection of WMH and NAWM tissue blocks from each donor, providing internal biological controls at every sampling site. Data from all modalities were co-registered to enable integrative cross-modality analysis, and interpretable machine learning was applied to identify lipid signatures most indicative of WMH versus NAWM molecular environments. Despite the known association between WMHs and cognitive decline, the underlying biochemical dysregulation remains elusive; our integrated multimodal approach addresses this by uncovering a previously unrecognized link between localized sulfatide metabolism (likely driven by fatty acid 2-hydroxylase (FA2H) dysregulation) and white matter pathology in the AD brain.

## Results and Discussion

Integrating multiple technologies enabled us to understand AD-associated WMH by revealing the relationships between tissue morphology, molecular distributions, and neuropathologies. **Figure 1** details the multimodal workflow we developed to study WMH in AD, combining clinically relevant MRI data with multimodal molecular imaging data on the same tissue, including autofluorescence (AF) microscopy, MALDI IMS, and histological stains. Specifically, using post-mortem MRI, we differentiated WMH and NAWM regions in human AD brains, which were then dissected into tissue blocks for submerged vibratome sectioning. Each mounted tissue section was analyzed by MALDI IMS for molecular characterization, followed by H&E staining, which revealed white/gray matter boundaries and served as a reference for traditional histopathology. Adjacent tissue sections were collected for LC-MS/MS lipidomics experiments to inform MALDI IMS peak annotations. Following data collection, image co-registration was performed pixel-wise, linking each MALDI IMS pixel to the corresponding AF and H&E-stained microscopy pixels from the same tissue section. This enabled multimodal data integration and analysis, including the application of supervised and interpretable machine learning across modalities.

**Figure 1.**
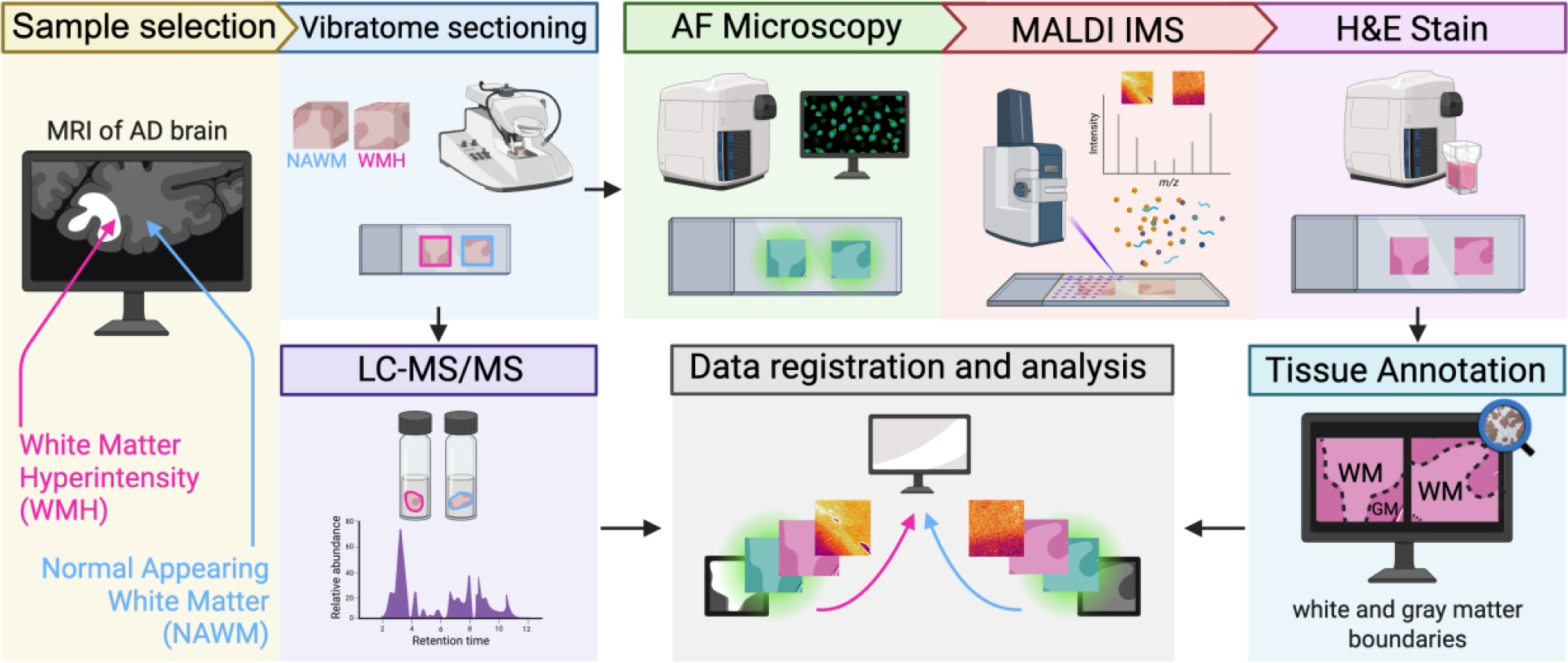
Schematic of the multimodal workflow for integrated lipid mapping and identification in white matter hyperintensities (WMHs). Included in the workflow were T2-weighted Fluid-Attenuated Inversion Recovery magnetic resonance imaging (T2-FLAIR MRI), autofluorescence (AF) microscopy-guided Matrix-assisted laser desorption ionization imaging mass spectrometry (MALDI IMS), histological staining, and liquid chromatography-tandem mass spectrometry (LC-MS/MS). Mounted WMH and normal-appearing white matter (NAWM) tissue sections were used for MALDI IMS, followed by hematoxylin and eosin (H&E) staining and annotation of white matter tissue masks based on H&E stains. Adjacent sections were homogenized for LC-MS/MS. LC-MS/MS data were used to generate a lipid library for annotation of analytes detected by MALDI IMS. Data from all imaging modalities were co-registered to the pixel-plane of the MALDI IMS data, and lipid data were extracted from within white matter masks, separating gray matter data. Supervised and unsupervised analyses were applied to discern differences in the lipid profile of WMH vs. NAWM, consistent across all donors and brain regions.

**Figure 2** highlights the brain tissue sampling method used for all donor brains and H&E-derived tissue section masks. **Figure 2A** is a visual representation of the types of tissue blocks gathered from each of the three AD donor brains (**Table 1**). WMH and NAWM regions were selected from both the anterior and posterior brain regions. While current literature typically evaluates WMH burden as a uniform, global metric, few studies investigate distinct regional differences. Growing evidence suggests that posterior WMH are associated with rapid memory decline, and show greater amyloid-β deposition, entorhinal cortical thinning, and progression to AD.^18,33–35^ Therefore, we characterized WMH in both anterior and posterior brain regions to build on these findings. The four tissue blocks selected from each donor based on MRI scans were sectioned for further analysis. To separate white matter from gray matter in these lipid data, we generated masks based on boundaries visible in H&E-stained images. **Figure 2B** shows the results of this white matter and gray matter annotation for one replicate. Masks of other replicates are shown in **Figure S3**. Additional example data outputs from the multimodal workflow in **Figure 1** are shown in **Figure 3** and **Figure S2-S6**. The entire workflow was applied to human brain tissue from three donors with AD. These tissues were obtained from Vanderbilt University Medical Center in collaboration with the Schrag research group. All three donors showed anterior and posterior WMH on post-mortem MRI scans (**Figure S1, Figure S2)**. WMH do not always appear in both anterior and posterior brain regions, but by selecting these specific tissues, direct, intra-donor comparisons between WMH and NAWM from both regions could be made.

**Figure 2.**
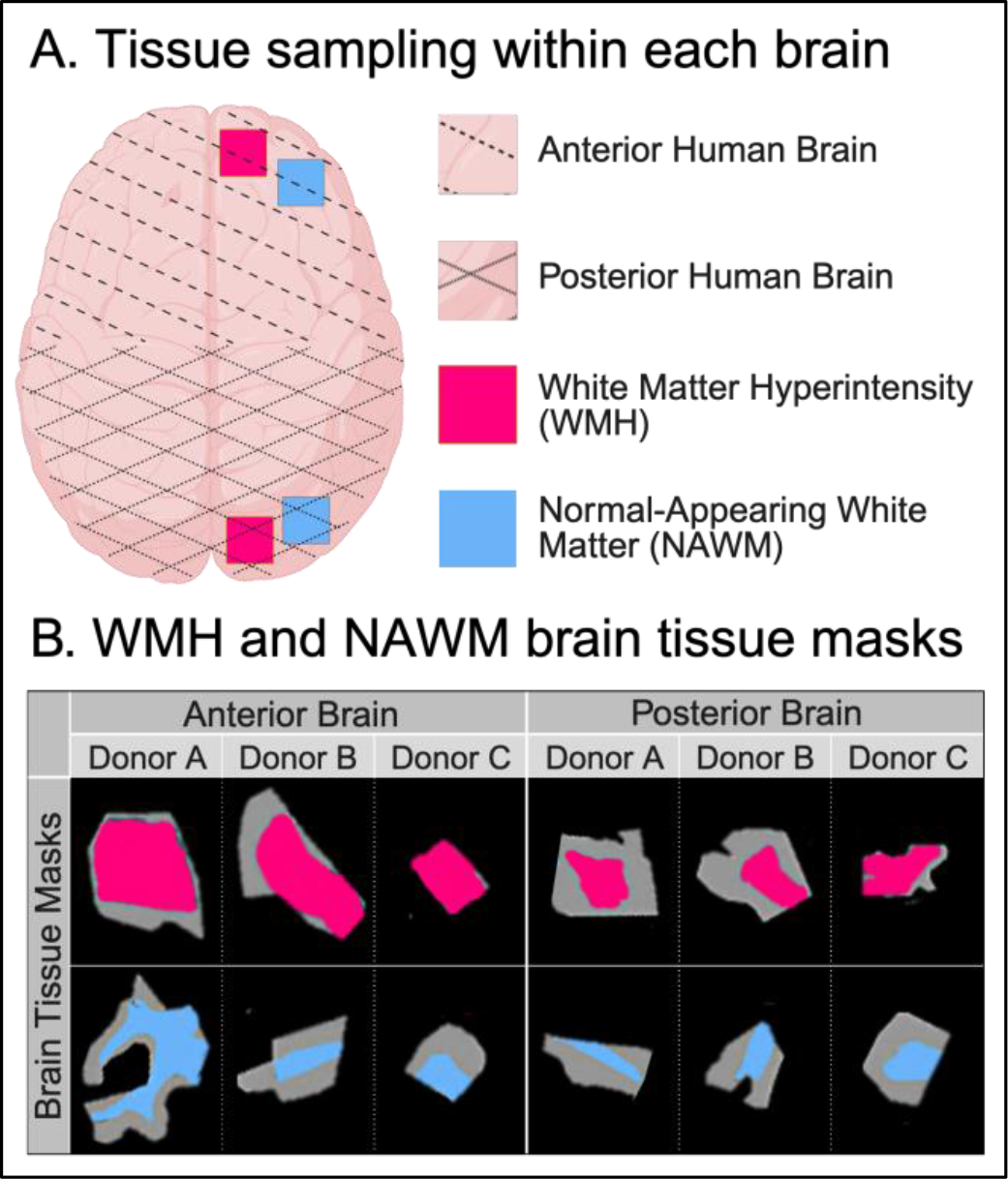
Donor tissue sampling strategy and annotation of white matter masks across modalities. (A) WMH and NAWM tissue samples were collected from both anterior and posterior regions of each human AD donor brain. (B) Representative tissue sections were examined using the multimodal workflow. Masks derived from post-IMS H&E staining delineate gray matter (gray), WMH (pink), and NAWM (blue).

**Figure 3.**
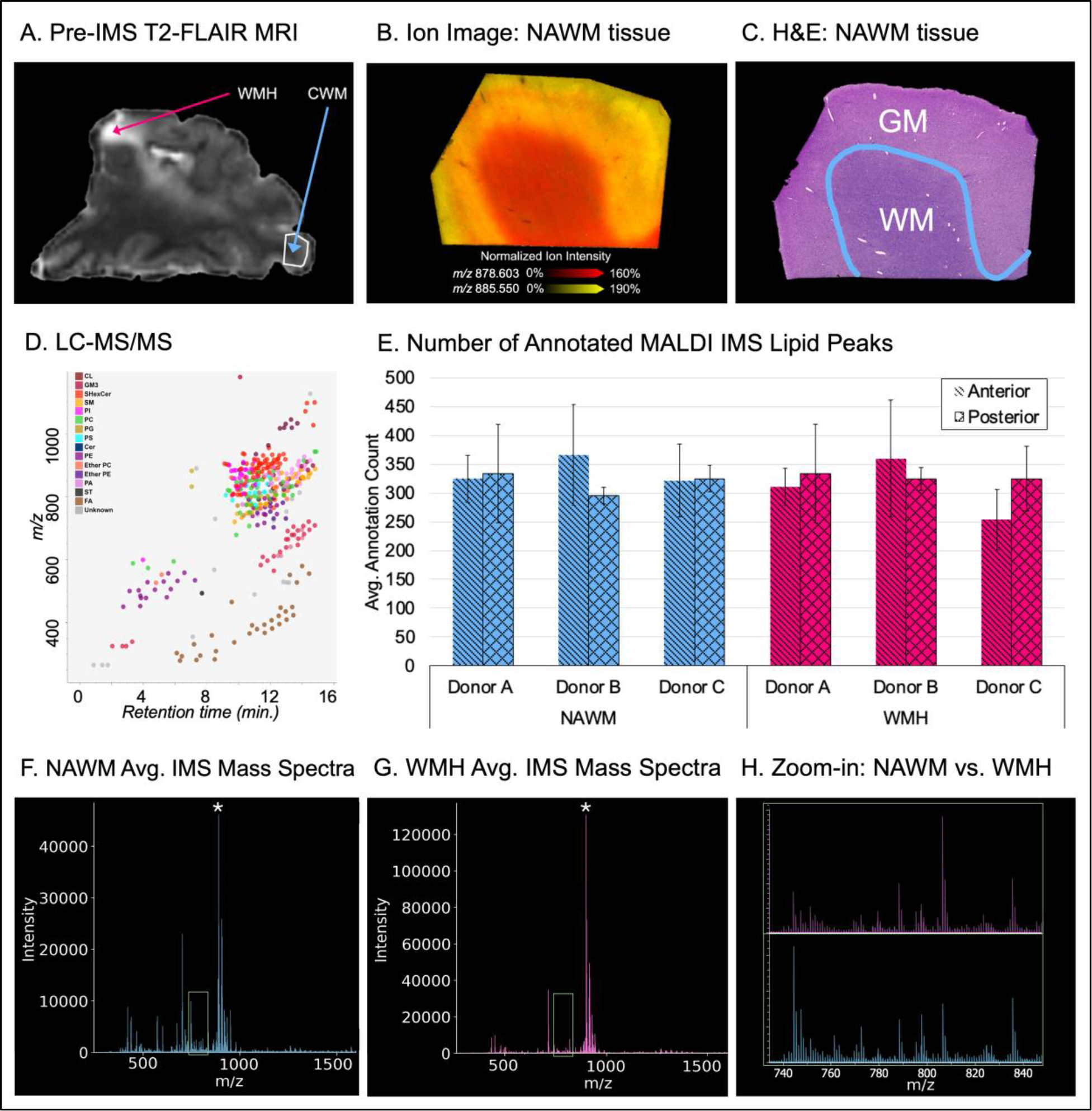
Representative outputs from the multimodal workflow for spatial lipidomic profiling of white matter hyperintensities (WMH) and normal-appearing white matter (NAWM). (A) T2-FLAIR MRI guided the identification of WMH and NAWM regions in each donor brain. (B) MALDI IMS acquired at 20 µm pixel size in negative ionization mode. Representative ion images show distinct lipid distributions across white and gray matter, including m/z 885.550 (PI 38:4), predominantly localized to gray matter, and m/z 878.603 (SHexCer 40:1;3O), predominantly localized to white matter but also present at lower intensity in gray matter. (C) Post-IMS H&E staining and annotation of gray matter and white matter. (D) LC-MS/MS analysis of homogenized tissue, showing detected lipid species plotted by m/z and retention time. (E) Number of MALDI IMS peaks annotated using the LC-MS/MS-derived lipid library across sample types (mean of triplicates, n=3). (F-G) Average MALDI IMS spectra from NAWM (F) and WMH (G) in a representative donor. [SHexCer (42:2;2O)-H]- (m/z 888.624) designated with a white asterisk. (H) Expanded view of the average spectra highlighting differential features.

**Table 1.** Human AD Donor Brain Tissue Block Collection.

|  | **Anterior WMH**<br>**Tissue Block** | **Anterior NAWM**<br>**Tissue Block** | **Posterior WMH**<br>**Tissue Block** | **Posterior NAWM**<br>**Tissue Block** |
| --- | --- | --- | --- | --- |
| **Donor A** | X | X | X | X |
| **Donor B** | X | X | X | X |
| **Donor C** | X | X | X | X |

**Figures 3A-C** show an MRI of the anterior brain tissue, a representative MALDI IMS overlay of two ion images, and an H&E stain of a tissue block containing NAWM from Donor C. A variety of lipid species, including phospholipids, ether phospholipids, sphingolipids, sterol lipids, and fatty acyls, are detected by LC-MS/MS in homogenized adjacent tissue sections (**Figure 3D)**, and the number of MALDI IMS peaks annotated as lipids (based on the LC-MS/MS data) is consistent across brain regions and donors (**Figure 3E).** In the average mass spectrum of NAWM (**Figure 3F**), we observe a rich lipid profile comprising sphingolipids, glycerolipids, glycerophospholipids, oxidized phospholipids, and other species. Fewer lipid species dominate the WMH average mass spectra (**Figure 3E**), and the relative intensities are altered compared to the NAWM. For example, SHexCer (42:2;2O) (*m/z* 888.624) is much higher in intensity in WMH (pink) compared to NAWM (blue) **(Figure 3F-G)**. The selected mass range (*m/z* 740–*m/z* 840) shown in **Figure 3H** highlights the changes in the relative intensities of other lipids between the two regions.

Given the inherent biological variability across human AD donors, we first evaluated whether molecular profiles were consistent across samples before proceeding with region-specific analyses. Uniform Manifold Approximation and Projection (UMAP) was applied to IMS data from all donors simultaneously, with tissue region-driven (rather than donor-driven) component structure indicating reproducible molecular profiles across the cohort. Representative UMAP results are shown in **Figure 4**, with all donor replicates provided in **Figure S3 and Figure S4**. Using H&E-based region annotations (**Figure 2B**) as a guide, Component 0 displays a molecular distribution predominantly localized to gray matter, while Component 1 reflects a distribution localized to the outer white matter and the white/gray matter boundary. Component 2 differs from components 0 and 1, exhibiting more heterogeneous molecular distributions within the white and gray matter, and less so at the border of the white and gray matter. These results demonstrate that the dominant molecular variance in the dataset reflects tissue architecture rather than donor-to-donor variation, establishing a foundation for subsequent region-specific analyses. This structural consistency was preserved despite pronounced variations in global neuropathological staging and mixed co-pathologies across our cohort. For example, tissue profiles from Donor C, which presented with a low AD score and confounding Lewy Body Disease (LBD) co-pathology, clustered together with profiles from the intermediate- and high-stage AD donors. This overlapping distribution underscores that the primary lipidomic features driving the UMAP structure are fundamentally governed by localized tissue anatomy rather than global clinical phenotypes or the specific presence of amyloid-β, tau, or Lewy body pathology. Nevertheless, resolving subtle molecular differences between WMH and NAWM requires greater sensitivity than unsupervised approaches can provide, motivating a more targeted, supervised machine learning approach.

**Figure 4.**
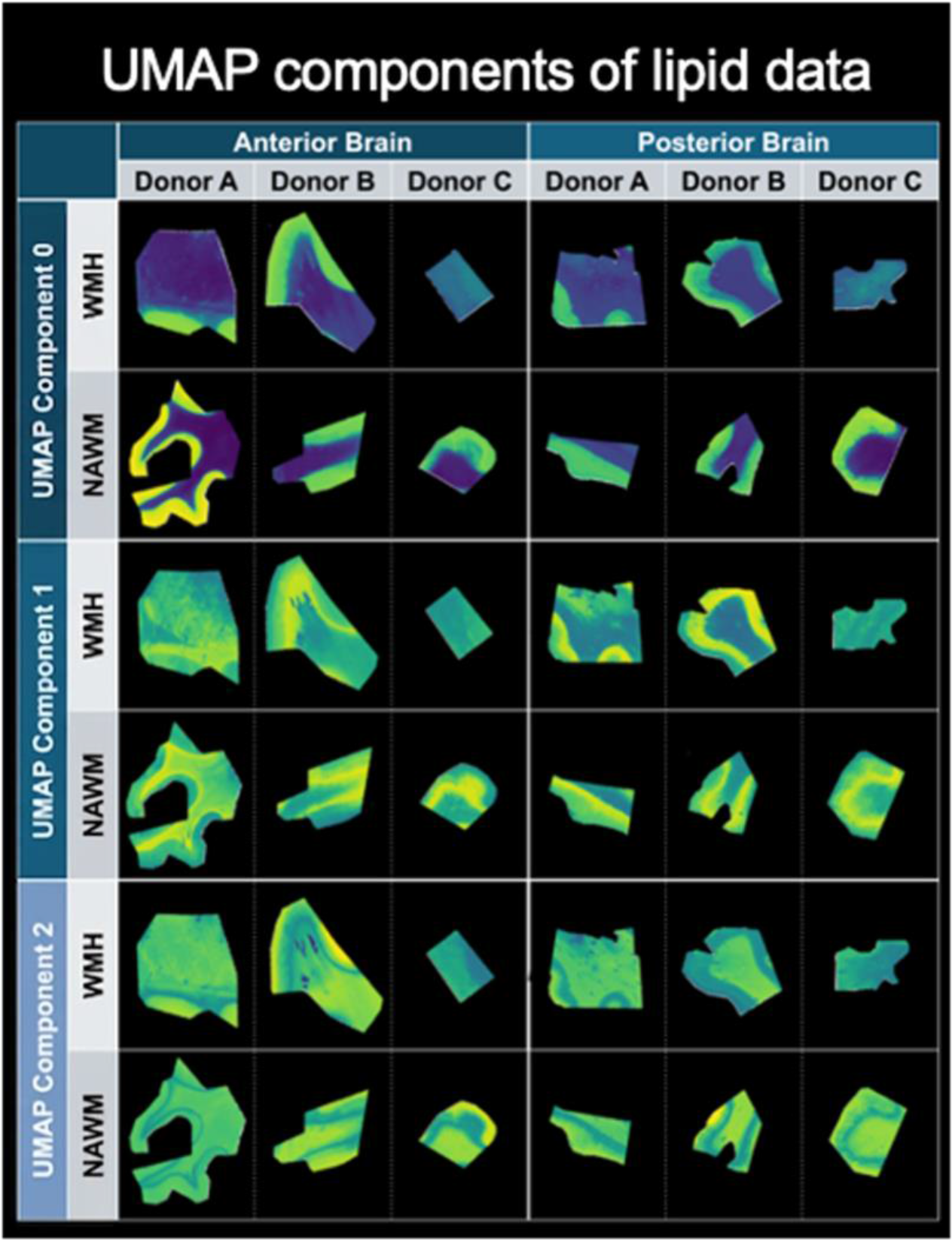
Uniform Manifold Approximation and Projection (UMAP) component images of IMS data from a representative replicate across all samples show three components that distinguish biological features within the human brain, rather than components of human variation. Component 0 captures the molecular variance in the spatial lipidomics dataset, with spatial distributions that separate white matter from gray matter and correspond closely to H&E-based region annotations (Figure 2B). Component 1 is localized to the outer white matter and the white/gray matter boundary. Component 2 shows a molecular distribution within the white and gray matter, lacking localization to the white and gray matter boundary.

We used the interpretable supervised machine learning workflow described by Tideman et al. to detect subtle yet important differences between WMH and NAWM brain tissue. With this workflow, XGBoost models identified relationships between IMS-observed lipid species and H&E-stain-derived white matter masks of WMH and NAWM. Once the XGBoost classification models demonstrated strong predictive performance at distinguishing WMH from NAWM, the models were interpreted using Shapley additive explanations (SHAP) to determine which lipid species drove the models’ predictions and thus held marker potential for distinguishing WMH from NAWM. The predicted labels, performance metrics, and top 20 SHAP features are shown in **Figures S7-S9**.

In **Figure 5**, lipids are ranked by Global SHAP importance score, where a higher score indicates that the lipid is a more significant marker for tissue classification. Spearman rank-order correlation coefficients (ρ) between ion intensity and Shapley values are depicted using a blue-to-red gradient, where red indicates that higher ion intensity is associated with WMHs, whereas blue signifies that lower ion intensity is associated with WMHs.

**Figure 5.**
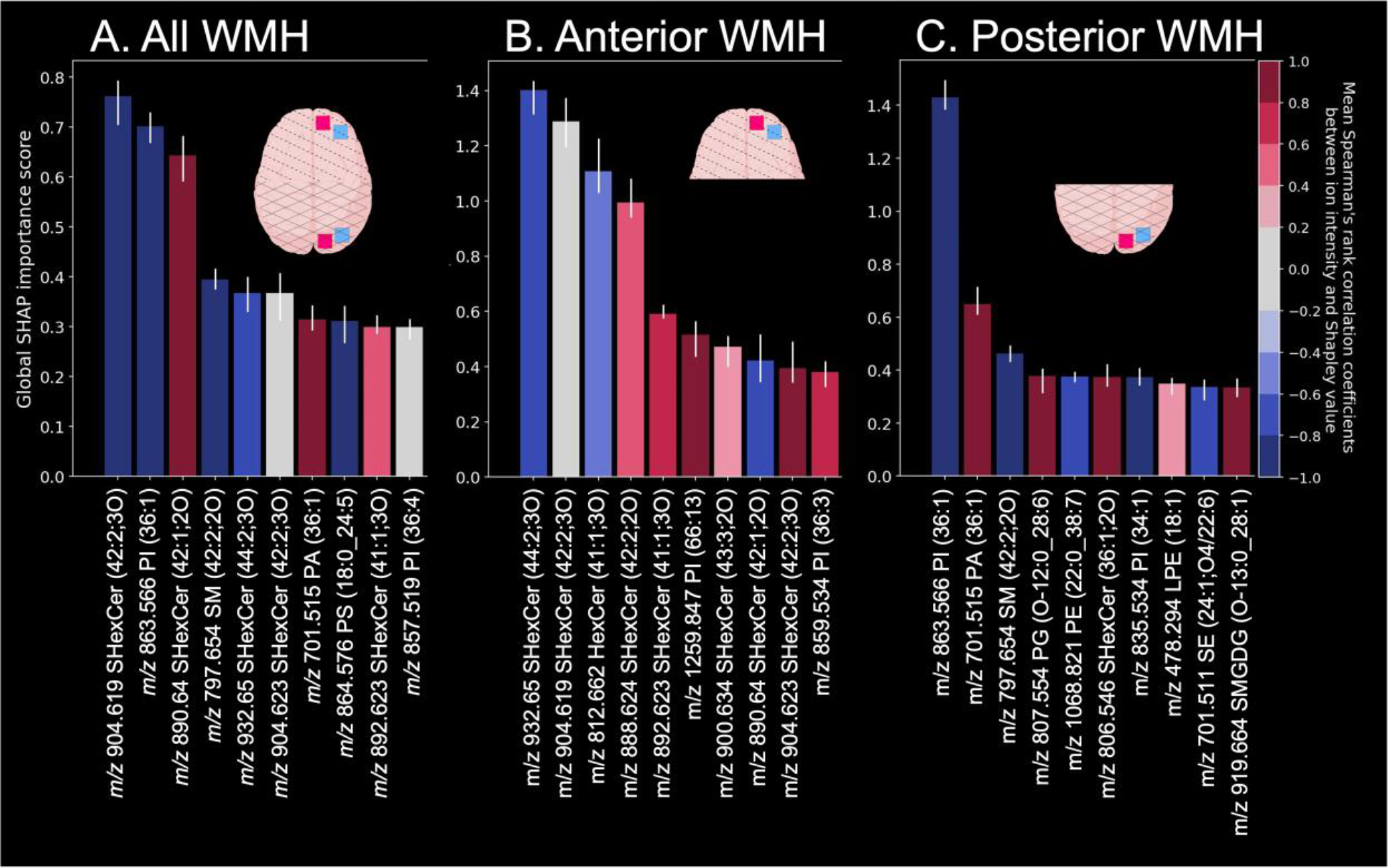
Interpretable machine learning workflows identify predictive lipid signatures distinguishing WMH vs. NAWM. High Global SHAP importance scores indicate a high degree of feature significance in the model’s ability to classify tissue regions. (A) SHAP interpretation of the binary XGBoost models reveals the top ten lipid features across all white matter tissue regions. Spearman rank-order correlation coefficients denote both positive correlations (red, associated with higher ion intensity in WMHs) and negative correlations (blue, associated with lower ion intensity in WMHs). Separate SHAP analyses were conducted for (B) anterior and (C) posterior brain regions.

Ranking by Global SHAP importance score highlights sulfatide SHexCer 42:2;3O (*m/z* 904.619), a structural and signaling lipid in myelin, as the top SHAP-ranked feature whose lower ion intensity is indicative of a WMH rather than NAWM (**Figure 5A)**. Here, the ‘3O’ denotes three additional oxygens built into the core lipid backbone and its acyl chains. These most commonly include two hydroxyl moieties on the sphingoid base and a third on the N-acyl chain. A decrease in the ion intensity of this trihydroxylated sulfatide is associated with WMHs across all tissue sections. In contrast, SHexCer 42:1;2O (*m/z* 890.640, likely with two hydroxyl groups on the ceramide backbone) is positively correlated with WMHs. Bulk lipidomic studies have established that sulfatide is depleted early and markedly in the AD brain, accompanied by a reciprocal increase in ceramide that points to sulfatide degradation rather than reduced biosynthesis.^27,28^ Our spatially resolved measurements localize this depletion to the WMH microenvironments. Further, our findings indicate that the WMH lipidome is remodeled rather than uniformly lost. Although most discriminating sulfatides decrease, species such as SHexCer 42:1;2O increase in WMH, consistent with shifts in acyl-chain length and hydroxylation state. Because FA2H catalyzes the 2-hydroxylation of the fatty acyl chain incorporated into myelin galactosylceramide and sulfatide,^36^ a regional decline in SHexCer 42:2;3O alongside preservation of its non-hydroxylated counterpart SHexCer 42:1;2O would be consistent with reduced FA2H activity, though altered sulfatide catabolism could contribute as well.^27,28,37^ It is noted that FA2H is inferred here from lipid signatures rather than measured directly, so this link remains a hypothesis to be addressed in future targeted studies. Another highly ranked lipid negatively correlated with WMHs is PI 36:1 (*m/z* 863.566), a phosphatidylinositol essential for membrane signaling and cytoskeletal stability, which acts as a structural anchor for proper myelin compaction.^38–40^

To assess regional variability in WMH composition, separate SHAP analyses were additionally performed for anterior and posterior brain regions (**Figure 5B, 5C**). This revealed notable spatial heterogeneity in the top predictive features driving the regional models. Sulfatide species, such as SHexCer 44:2;3O (m/z 932.650) remained primary drivers of classification performance in the anterior regions. While this exact lipid species has not been previously detailed in the Alzheimer’s disease literature, its identification aligns with the understanding that broad sulfatide depletion and remodeling are critical early events in AD-associated white matter disruption.

PI 36:1 (*m/z* 863.566) was the top-ranked posterior marker, suggesting a more prominent role for phosphatidylinositol disruption in posterior WMH regions compared to anterior counterparts. Furthermore, several lipids were highly ranked only in the posterior SHAP models, such as SM 42:2;2O (*m/z* 797.654), potentially reflecting differences in sphingomyelin metabolism across brain regions. These regional differences highlight the spatial heterogeneity inherent to WMH pathology. Bulk profiling of frontal and occipital subcortical WMH in AD has likewise reported region-dependent molecular differences;^8^ our *in situ* analysis extends this by localizing such differences to the lesion itself and naming the specific lipid species that distinguish anterior from posterior WMH. Rather than establishing a direct causal mechanism for white matter degradation, these divergent SHAP profiles reveal distinct, highly localized lipidomic shifts that act as top-contributing molecular signatures of white matter pathology in the Alzheimer’s disease brain.

The underlying mechanisms driving this regional heterogeneity are likely multifactorial. Longitudinal MRI studies demonstrate that AD-associated WMHs initially manifest in the posterior-periventricular region. Subsequently, these lesions expand into the deep parietal and occipital white matter, including the splenium, and later extend into the genu of the corpus callosum, the frontal periventricular region, and the deep frontal areas. This posterior-to-anterior WMH trajectory suggests that anterior WMHs may represent “younger” pathology, whereas posterior WMHs represent “older” pathology characterized by prolonged exposure to lipid dysregulation and resulting lipid dyshomeostasis. Additionally, intrinsic anatomical and functional differences between anterior and posterior brain regions may shape lipidomic responses to damage. Distinct glial or vascular microenvironments could also influence regional vulnerability and repair mechanisms.

## Conclusions

Here, we resolve the lipid landscape of white matter hyperintensities in the human AD brain with both spatial and regional specificity. This study establishes a multimodal workflow combined with interpretable supervised machine learning to identify specific molecular signatures characteristic of the WMH microenvironment in formalin-fixed human AD brain tissue. These findings highlight sulfatide alterations present in anterior WMH and implicate altered FA2H kinetics or downstream degradation pathways as a possible cause. Conversely, PI 36:1 depletion was found to be a key indicator of posterior AD WMHs. These results provide a molecular framework for understanding the role of lipid dysregulation in the AD brain. Together, these findings suggest that WMHs cannot be characterized as homogeneous features; rather, they exhibit spatially distinct molecular signatures that provide context for understanding region-specific disease progression and early therapeutic targeting for AD. Notably, these molecular differentiators emerged from a limited donor cohort, underscoring the sensitivity of the workflow for resolving WMH-specific lipid signatures even at modest sample sizes. Expanding to larger cohorts in future studies would provide the statistical power to resolve more subtle distinctions between NAWM and WMH and to test whether these regional signatures generalize across donors. Additionally, future studies should seek to contextualize AD WMH lipidomic profiles within their protein and cell microenvironments by incorporating multiplexed immunofluorescence microscopy into the multimodal workflow. This multimodal approach can be further extended to characterize the metabolic interplay between WMH and other AD-associated features, including amyloid-β plaques, dystrophic neurites, tau tangles, and vascular pathology.

## Experimental Methods

### Human Brain Tissue

Brain tissues were obtained from the Vanderbilt Brain and Biospecimen Bank at Vanderbilt University Medical Center, Nashville, Tennessee, USA (IRB# 180287). Written informed consent for each anatomical donation was obtained from the patient or their surrogate decision-maker. The Vanderbilt University Medical Center Institutional Review Board provided ethical supervision, and the study was performed in accordance with the ethical principles of the Declaration of Helsinki.

### Post-mortem Magnetic Resonance Imaging

Coronal tissue slabs were sliced coronally and embedded in 1% agarose under degassing conditions, with care to avoid air bubbles. The agarose was at a temperature within the hysteresis range: agarose solution forms a gel at below 30°C and melts at above 90°C. Post-mortem susceptibility weighted imaging (SWI) and T2-weighted FLuid Attenuated Inversion Recovery (FLAIR) MRI were obtained on both 3.0T (Philips Ingenia) and 7.0T (Philips Achieva) human clinical scanners using 32-channel phased array reception. 3.0T SWI (technique = 3D gradient echo, number of echoes = 4, first echo = 7.2 ms, echo spacing = 6.2 ms, repetition time = 31 ms, spatial resolution = 0.6x0.6x2 mm), 3.0T FLAIR (technique =3D turbo-inversion-recovery, echo time = 271 ms, repetition time = 4800 ms, inversion time=1650 ms, spatial resolution=1.0x1.0x1.0 mm) were performed with sequence parameters chosen to parallel in vivo protocols to resemble the clinical conditions.

### Tissue Selection

Post-mortem T2-weighted-Fluid-Attenuated Inversion Recovery (T2-FLAIR) MRI To begin examining the lipidomic profiles within white matter hyperintensities (WMH), three human AD brain donor cases were selected from the Schrag Lab Brain Bank. Brain donor tissue was characterized neuropathologically according to the NIA-AA consensus criteria for the postmortem diagnosis of AD, by an experienced pathologist. ^41^ The severity of AD neuropathological change was determined using the combined ‘ABC’ score. Post-mortem T2-weighted-Fluid-Attenuated Inversion Recovery (T2-FLAIR) magnetic resonance imaging (MRI) was performed on formalin-fixed donor brain tissue. Tissues were stored in 1x TBS /0.02% sodium azide. Donors were selected based on T2-FLAIR MRI, with all three donors exhibiting WMH in both the anterior and posterior regions of the brain. Blocks of tissue encompassing WMH, and separate blocks encompassing NAWM, were excised from each donor, posteriorly and anteriorly (Table 1, Figure S1). White matter was not dissected from gray matter; thus, all blocks also contained some surrounding gray matter. Using a submerged vibratome with chilled 1xTBS, serial tissue sections were cut from each tissue block for MALDI IMS, followed by H&E staining in triplicate. Two 30 μm sections were cut first, followed by eleven 15 μm thick sections (vibratome settings: 30 μm thickness, 0.1 speed, 0.85 amplitude; 15 μm thickness, 0.02 speed, 0.75 amplitude). Tissue blocks were adhered to the chuck with a thin layer of superglue after the tissue mounting surface was dabbed dry. Tissue sections were stored in sequential order, with one tissue section floating in each well of a 24-well plate. Each well contained 2 mL 1x TBS/0.02% sodium azide, and plates were sealed with paraffin film and stored at 4 °C. Three additional serial sections from each block were placed at the bottom of a corresponding glass vial, for separate LC-MS/MS preparation and analysis of each tissue block. Small, clean paintbrushes were used to transfer floating tissue sections between the vibratome chamber and storage containers.

### Tissue Mounting

Four floating sections from each tissue block were mounted onto poly-L-lysine-coated indium-tin-oxide (ITO) glass slides (Delta Technologies, Loveland, CO). After pipetting a 1 mL droplet of chilled 1x TBS onto the slide, a tissue section was carefully transferred into the droplet and, using paintbrushes, unfurled and laid flat, avoiding wrinkles or stretching as much as possible. Depending on the tissue morphology, the droplet was then slowly poured or pipetted off, followed by drying with a low-pressure (1-2 PSI) stream of nitrogen gas.

### Pre-MALDI IMS Autofluorescence Microscopy

Before MALDI IMS, autofluorescence microscopy (Pre-IMS-AF) images of all mounted sections (4 sections/block, 12 blocks, 48 sections imaged in total) were acquired using a Zeiss AxioScan.Z1 slide scanner (Carl Zeiss Microscopy GmbH, Oberkochen, Germany), equipped with a Colibri7 LED light source, capturing channels 350 nm, 488 nm, and 555 nm. A 10× objective (NA ∼0.45) was used, with a resulting pixel sampling of 0.65 µm/pixel. Of the four sections mounted from each block, three advanced to the MALDI IMS stage based on their optimal mounting, as visualized in pre-IMS autofluorescence images.

### MALDI IMS Tissue Preparation

To prepare for MALDI IMS, slides were washed four times on ice with 150 mM ammonium formate for 45 seconds each to remove salts. Slides were then dried with a flow of nitrogen gas (1-2 PSI), followed immediately by sublimation of 4-Dimethylaminocinnamaldehyde (DMACA) solubilized in Tetrahydrofuran at roughly 0.8µg/mm^2^.

### MALDI IMS Data Acquisition

MALDI IMS data from brain tissue were acquired at 20 µm spatial resolution in negative ionization qTOF mode with a scan range of *m/z* 250 to 2500 on a Bruker timsTOF fleX mass spectrometer (Bruker Daltonics). With Beam Scan on, laser power was set to 18% and each pixel consisted of 200 shots from a SmartBeam 3D 10 kHz frequency tripled Nd:YAG laser (355 nm). Red phosphorus was spotted on the target plate and used for mass calibration prior to data acquisition.

### Post-MALDI IMS Autofluorescence Microscopy

Following MALDI IMS data acquisition, autofluorescence microscopy (Post-IMS-AF) images of all slides (3 sections/block, 12 blocks, 36 sections imaged in total) were acquired using a Zeiss AxioScan.Z1 slide scanner (Carl Zeiss Microscopy GmbH, Oberkochen, Germany), equipped with a Colibri7 LED light source, capturing channels 488 nm, 555 nm, and a brightfield image. A 10× objective (NA ∼0.45) was used, with a resulting pixel sampling of 0.65 µm/pixel. The images show the matrix desorption markings created via MALDI IMS. Visualizing these markings is integral to our multimodal image registration.

### Post-MALDI IMS H&E Staining

Following MALDI IMS and post-IMS-autofluorescence microscopy acquisition, the matrix was removed from the slides using ethanol washes (30 second 95% ethanol, 30 second 70% ethanol) followed by three D.I. water rinses. Tissue is then fixed with 10% neutral buffered formalin for 3 minutes, followed by three D.I. water rinses. Slides were then incubated at room temperature in Hematoxylin for 3 minutes, rinsed three times with D.I. water, dipped in ammonium hydroxide, and immediately rinsed three times with D.I. water. Eosin counterstain and dehydration consisted of 20 dips in 70% ethanol, 20 dips in 95% ethanol, 1 minute in Eosin, 20 dips in 95% ethanol, 10 dips in 95% ethanol, 20 dips in 100% ethanol, and a final 20 dips in 100% ethanol. Lastly, slides were washed three times with Xylene 10 seconds then cover-slipped using CytoSeal mounting medium.

### Annotation of white matter masks

Masks of the white matter were drawn on H&E stained images in QuPath v.0.5.1, in collaboration with neurologist Matthew Schrag, MD, PhD. The state of the white matter (NAWM or WMH) and the location (anterior or posterior brain) were predetermined by the tissue block from which the sample was sectioned. Following multimodal image registration, these masks allowed us to select only the MALDI IMS pixels contained within the white matter for analysis and comparison across groups.

### MALDI IMS Data Preprocessing

MALDI IMS data were exported from Bruker timsTOF (.d) files into a custom binary format. Centroided mass spectra from each pixel were recalibrated using Bruker DataAnalysis TF-SDK (v2.21). Spectra were aligned in *m/z* space with the msalign library (v0.2.0) using at least six reference peaks. A minimum of four reference ions were used to perform mass recalibration, resulting in an estimated mass accuracy of ∼1 ppm.

Following alignment, spectra were normalized by total ion current (TIC), and an average spectrum was generated from all tissue pixels. Peak picking performed on the averaged spectrum yielded 550 *m/z* features across human brain tissue samples. Spectra were computationally deisotoped before downstream analysis, although some residual isotopic features may have remained. Detected peaks were subsequently used to extract centroid ion intensities within a ±5 ppm mass tolerance. For WMH and NAWM analyses, H&E-derived annotation masks were transformed into the IMS coordinate space.

### Lipid Extraction and LC-MS/MS data acquisition

Lipid extraction was performed as previously described on protocols.io (https://dx.doi.org/10.17504/protocols.io.ewov1ob5klr2/v1). ^42^ This protocol for bulk untargeted LC-MS/MS lipidomics involves lipid extraction by MTBE and connecting Waters UHPLC to a Bruker timsTOF fleX mass spectrometer via contact closure.

LC-lipidomic data acquired from serial tissue sections in negative ion mode were used to build a reference library for MALDI IMS annotations. MS/MS stepping and a Waters Premier HPLC with a 2.1 x 100 mm CSH-C18 column were used. Sulfatide annotations were manually confirmed in MS-DIAL based on the presence of the characteristic bisulfate ion peak at *m/z* 96.9.

### Image Registration

Image registration was performed in two parts, with the goal of precisely linking microscopy images (Pre-IMS-AF, Post-IMS-AF, H&E) to the MALDI data. Using image2image Elastix, MALDI pixels were linked with the matrix ablation marks present in the post-IMS autofluorescence microscopy image. ^43^ With another software (wsiReg), the pre-IMS-AF, post-IMS-AF, and H&E images were registered together. ^44^ In both image2image and wsiReg, all images were registered and transformed to the “Target Modality” Post-IMS autofluorescence microscopy image.

### Supervised Machine Learning and Feature Importance Analysis

Using white matter masks annotated from H&E-stained sections, three distinct eXtreme Gradient Boosting (XGBoost) binary classification models were developed to differentiate: (1) total WMH vs. NAWM, (2) posterior WMH vs. posterior NAWM, and (3) anterior WMH vs. anterior NAWM. Classification was performed at the pixel level based solely on the corresponding MALDI IMS data, utilizing a total of 3,392,445 spectra for model training and validation. Because each tissue donor contributed paired WMH and NAWM regions, the experimental design inherently controlled for inter-subject biological variability. Each binary classification task was structured to differentiate the positive class (WMH) from the negative class (NAWM) using pixel-wise ion intensity profiles.

#### Ion annotation and dataset preparation

A dataset comprising 550 ions detected by imaging mass spectrometry (IMS) was selected for machine learning classification. To ensure robust characterization, these features were tentatively annotated by matching accurate mass values against a high-confidence lipid database generated via LC-MS/MS analysis of adjacent tissue sections (mass tolerance < 5 ppm; false discovery rate (FDR) ≤ 0.01).

#### Model training and hyperparameter optimization

Discriminative classification models were implemented in Python (v3.9.18) utilizing the scikit-learn (v1.3.0) and XGBoost (v2.1.1) libraries. For each spatial mask, the data were partitioned into a 67% training and 33% testing split. To optimize classification accuracy, hyperparameters were systematically tuned using a grid-search approach on a validation subset of the training data prior to final model construction. *SHAP feature importance and spatial correlation.* To determine which lipid species drove the model predictions, the trained XGBoost models were interpreted using Shapley additive explanations (SHAP), implemented with the SHAP library (v0.46.0). SHAP provides both a global (experiment-wide) and a local (per-pixel) importance score for each lipid species. The global SHAP importance score was obtained by taking the mean of the magnitude of the local SHAP importance scores (Shapley values) across all pixels, and lipid species were ranked in descending order of global SHAP importance score to shortlist potential biomarker candidates. To establish whether a candidate was positively or negatively correlated with WMH, the direction and significance of the relationship between the mean-centered ion intensity and the Shapley values of each lipid species were quantified using the Spearman rank-order correlation coefficient (ρ), where magnitudes greater than 0.2 were considered significant.

## Supporting information

Supplemental Doc 3

Supplemental Doc 1

Supplemental Doc 2

## Acknowledgements

Funding provided by National Institutes on Aging R01AG078803, T32-AG058524, CZI 2021-240339 & 2022-309518. Thanks to our collaborators in the Schrag and Van de Plas labs, the Bruker Daltonics team for supporting this project, and the Mass Spectrometry Research Center at Vanderbilt. Special thanks to the human brain donors and their families. Figures made using BioRender, ImageJ, and SCiLS Lab.

Language and editing assistance were provided by humans and, minimally, by Gemini and ChatGPT strictly to improve the clarity and grammar of the narrative text. The authors carefully verified all technical content, data coordinates, and interpretations, and maintain complete accountability for the final manuscript.

## Abbreviations

T2-FLAIR: T2-weighted Fluid-Attenuated Inversion Recovery
MRI: magnetic resonance imaging
LC-MS/MS: Liquid Chromatography-Tandem Mass Spectrometry
MALDI: Matrix-assisted laser desorption ionization
IMS: imaging mass spectrometry
H&E stain: Hematoxylin and eosin stain
AF: microscopy Autofluorescence microscopy
AD: Alzheimer’s disease
WMH: White matter hyperintensity
NAWM: Normal-appearing white matter
UMAP: Uniform Manifold Approximation and Projection
XGBoost: eXtreme Gradient Boosting
SHAP: Shapley additive explanations
m/z: Mass-to-charge ratio

## Notes

### Competing Interest Statement

The authors have declared no competing interest.

