## Supplemental Doc 1 for "Elucidating Molecular Features of White Matter Hyperintensities in Alzheimer’s Disease through Multimodal Imaging and SHAP Analysis"

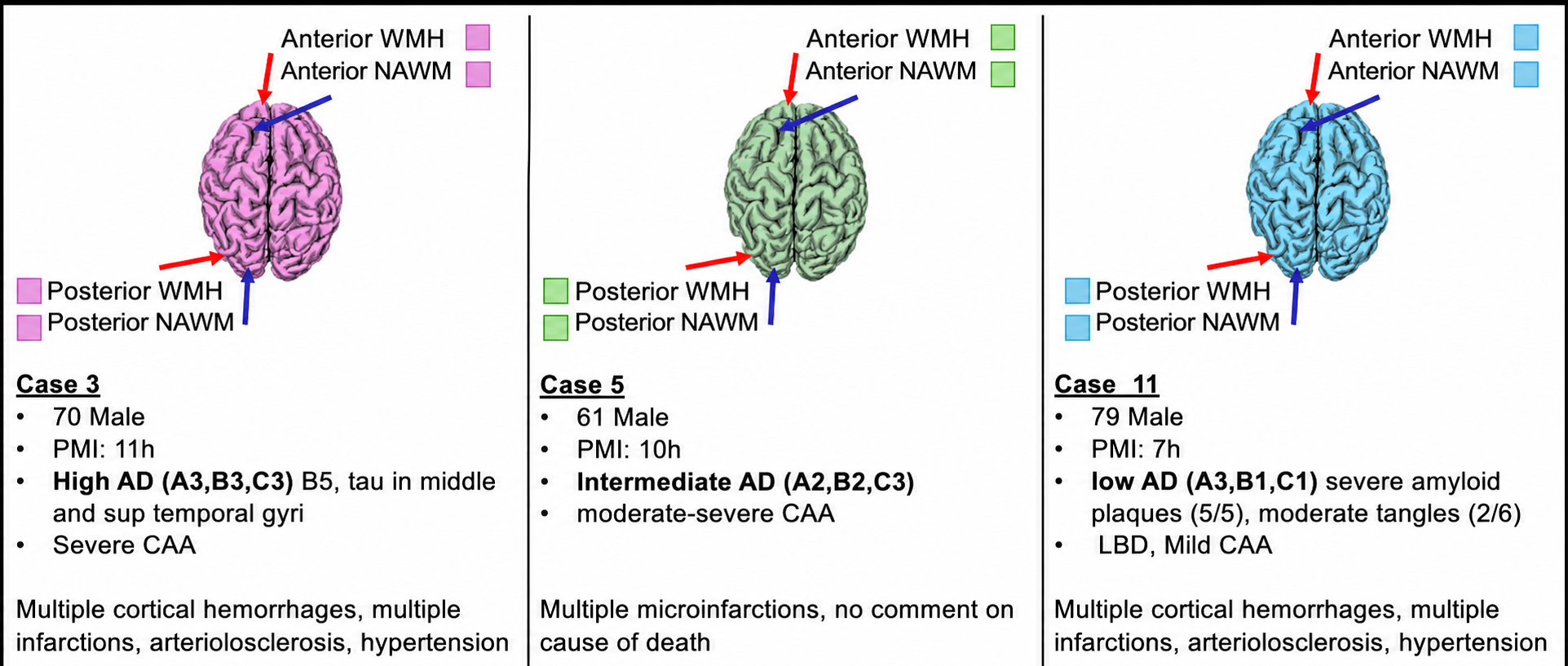

***Figure S1.*** *Sample Collection from three human AD donor cases and corresponding available patient data.* *Note: For data continuity across computational pipelines, Donor A is labeled as Case 3, Donor B as Case 5, and Donor C as Case 11 in subsequent figures (Figures S4, S7, S8, S9).*

| **Acquisition Parameter** | **Value** |
| --- | --- |
| **Transfer** |  |
| Set MALDI Plate Offset | -50.0 V |
| Deflection Delta Sum | -125.0 V |
| Set Funnel 1 RF | 350.0 Vpp |
| IsCID Energy | 0.0 eV |
| Set Funnel 2 RF | 300.0 Vpp |
| Multipole RF | 300.0 Vpp |
| **Collision Cell** |  |
| Set Collision Energy Offset | -10.0 eV |
| Set Collision Cell RF | 2500.0 Vpp |
| **Quadrupole** |  |
| Set Ion Energy Offset | -10.0 eV |
| Set Isolation Mass (MS only) | *m/z* 500.00 |
| **Digitizer** |  |
| Set Transfer Time | 120.0 µs |
| Set PrePulseStorage Time | 12.0 µs |

**Table S1.** Instrument parameters for negative mode qTOF MALDI IMS experiments

| *m/z* | Identity | Mean ppm |
| --- | --- | --- |
| 904.619 | [SHexCer (42:2;3O)-H]^-^ | ± 0.5006 |
| 863.566 | [PI (18:0_18:1)-H]^-^ | ± 1.0748 |
| 890.640 | [SHexCer (42:1;2O)-H]^-^ | ± 4.3609 |
| 932.650 | [SHexCer (44:2;3O)-H]^-^ | ± 1.3623 |
| 812.662 | [HexCer (41:1;3O)-H]^-^ | ± 2.6315 |
| 888.624 | [SHexCer (42:2;2O)-H]^-^ | ± 0.6689 |
| 885.550 | [PI (38:4)-H]^-^ | ± 0.7441 |
| 878.603 | [SHexCer (40:1;3O)-H]^-^ | ± 0.1383 |

**Table S2.** Lipid Identities (an exhaustive list of lipid identities is shown in Supplement 2)

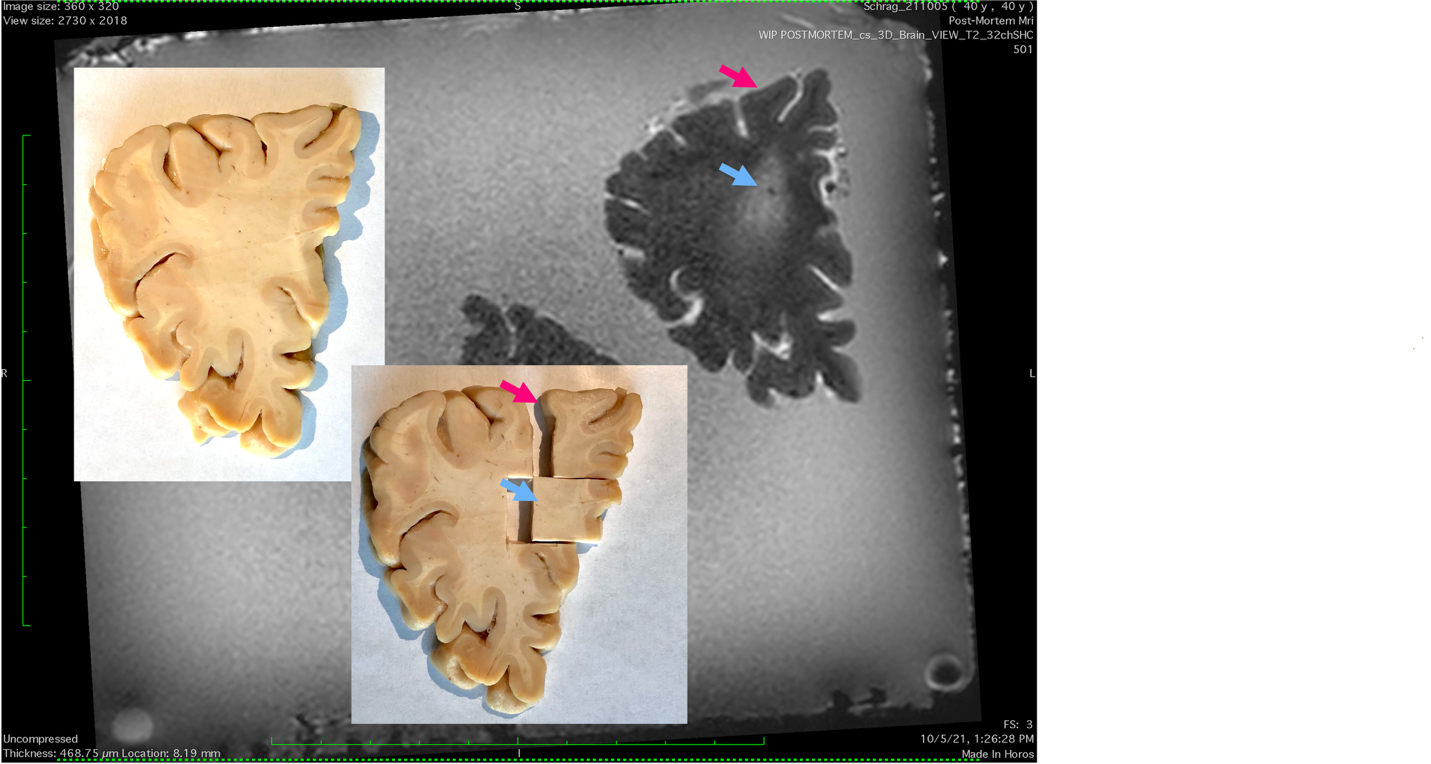

Donor A Anterior WMH and NAWM

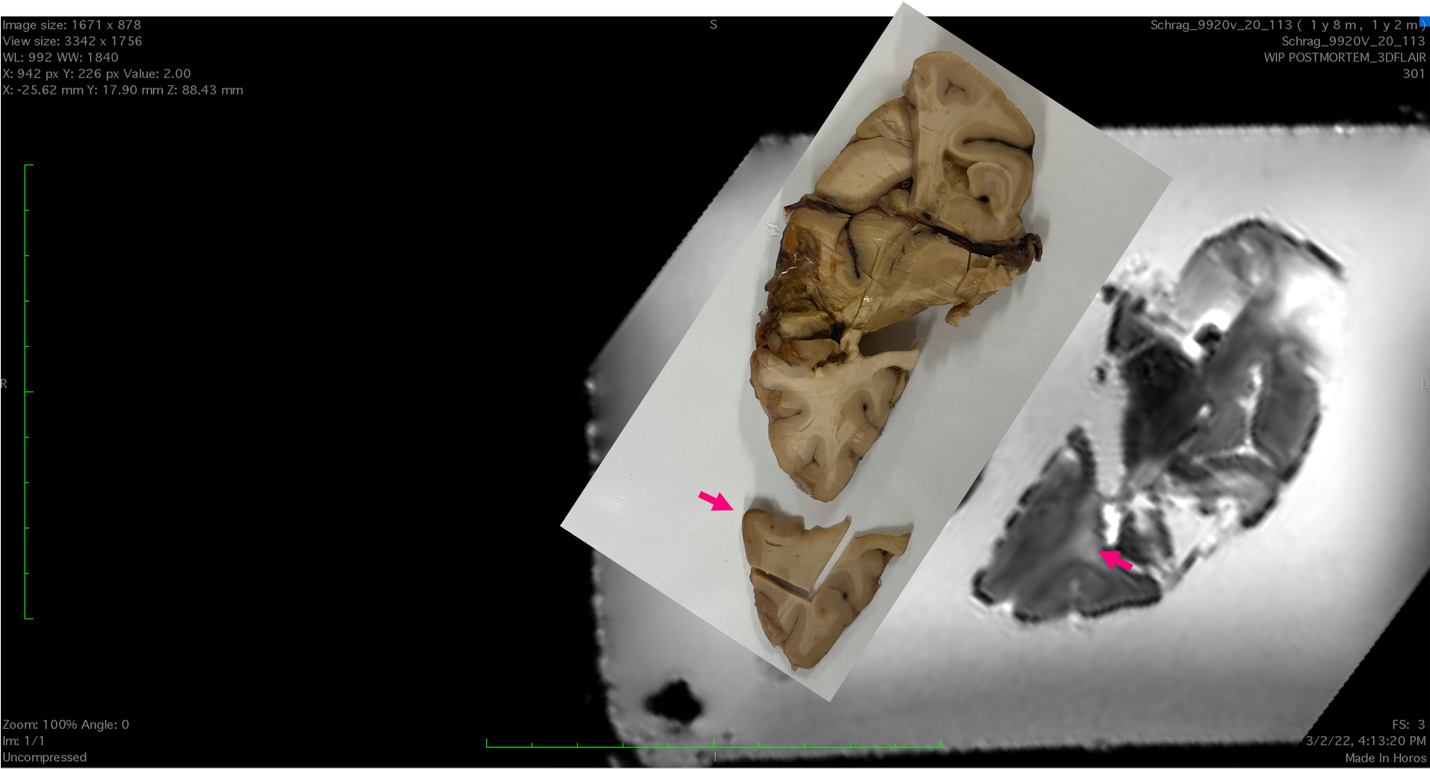

Donor B Anterior WMH and NAWM

*(Figure S2 continued on next page)*

**Figure S2 (continued)**

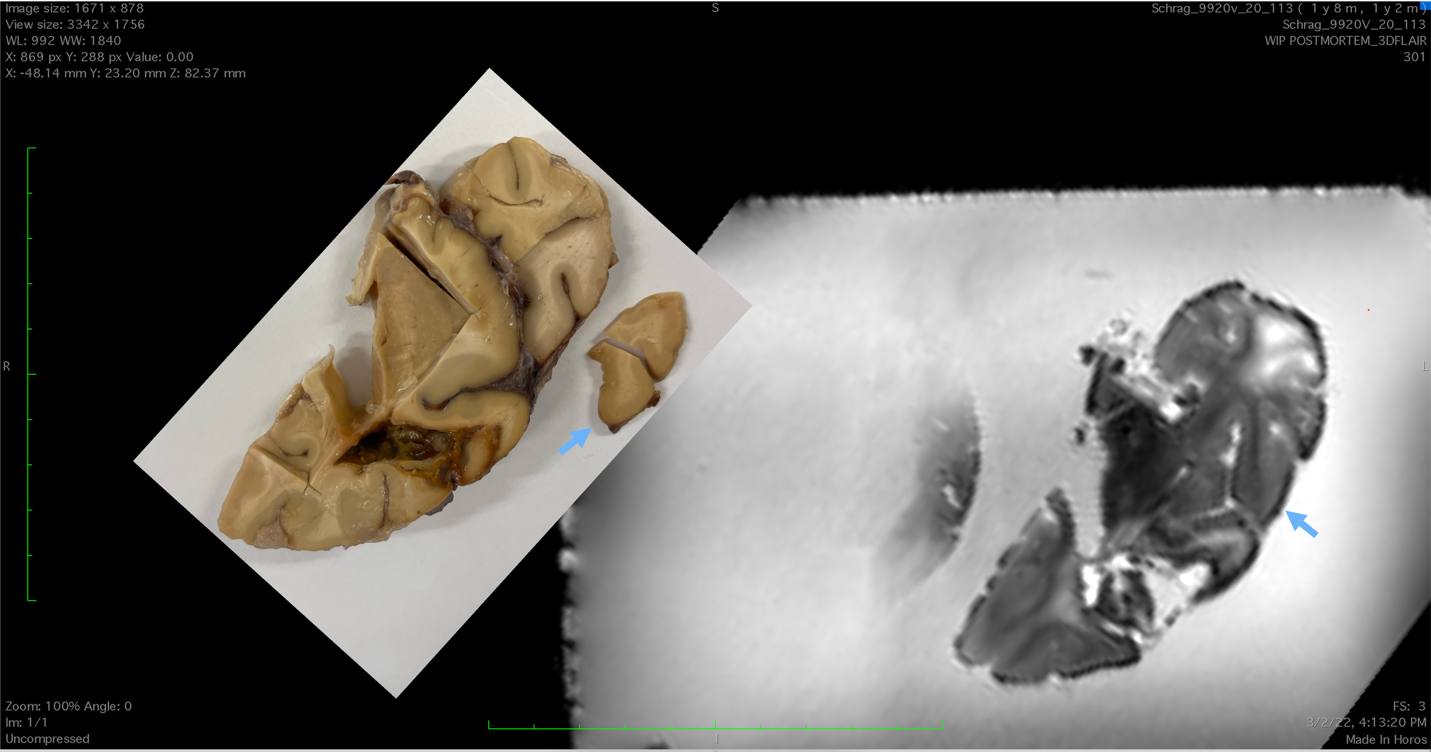

Donor B Anterior NAWM

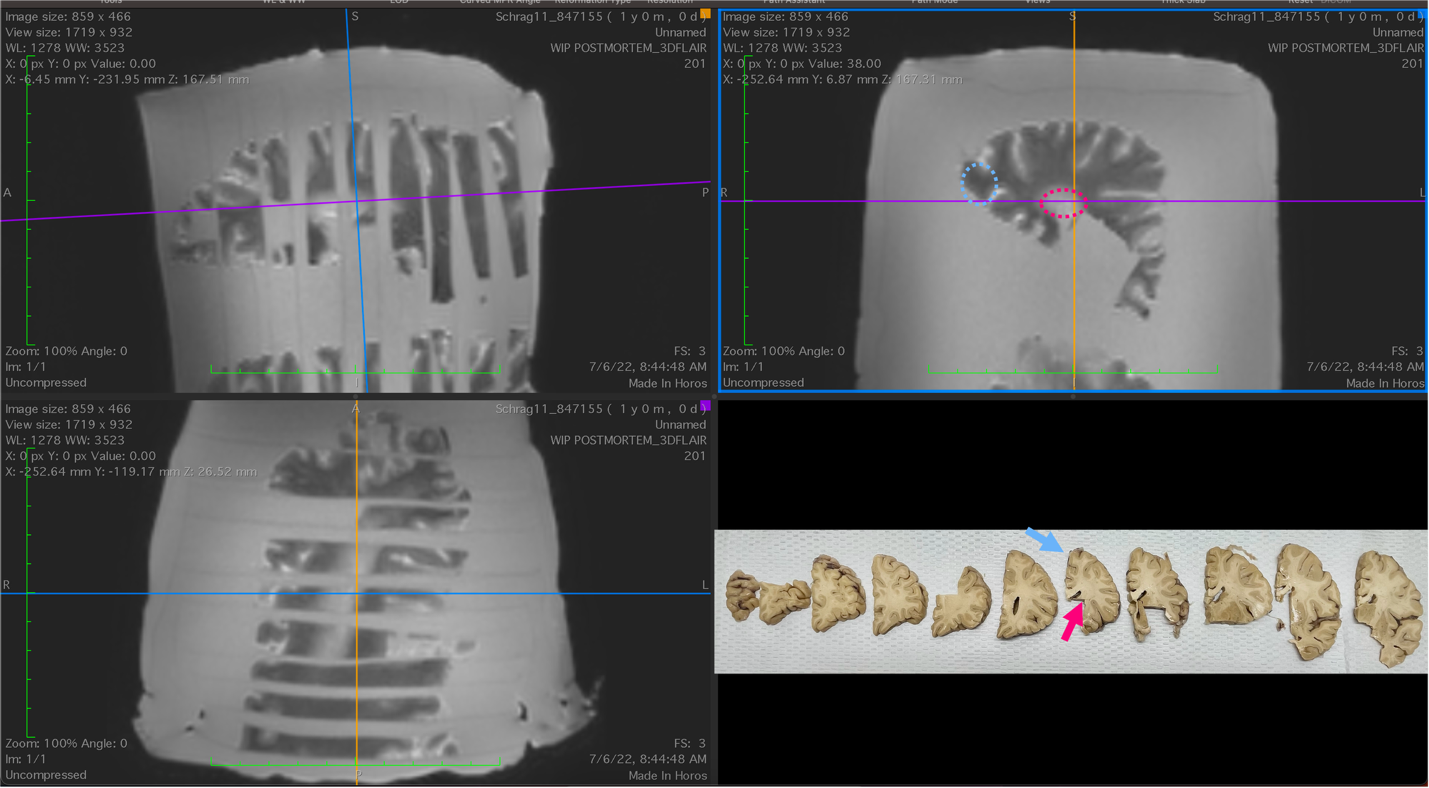

Donor C Anterior WMH and NAWM

*(Figure S2 continued on next page)*

**Figure S2 (continued)**

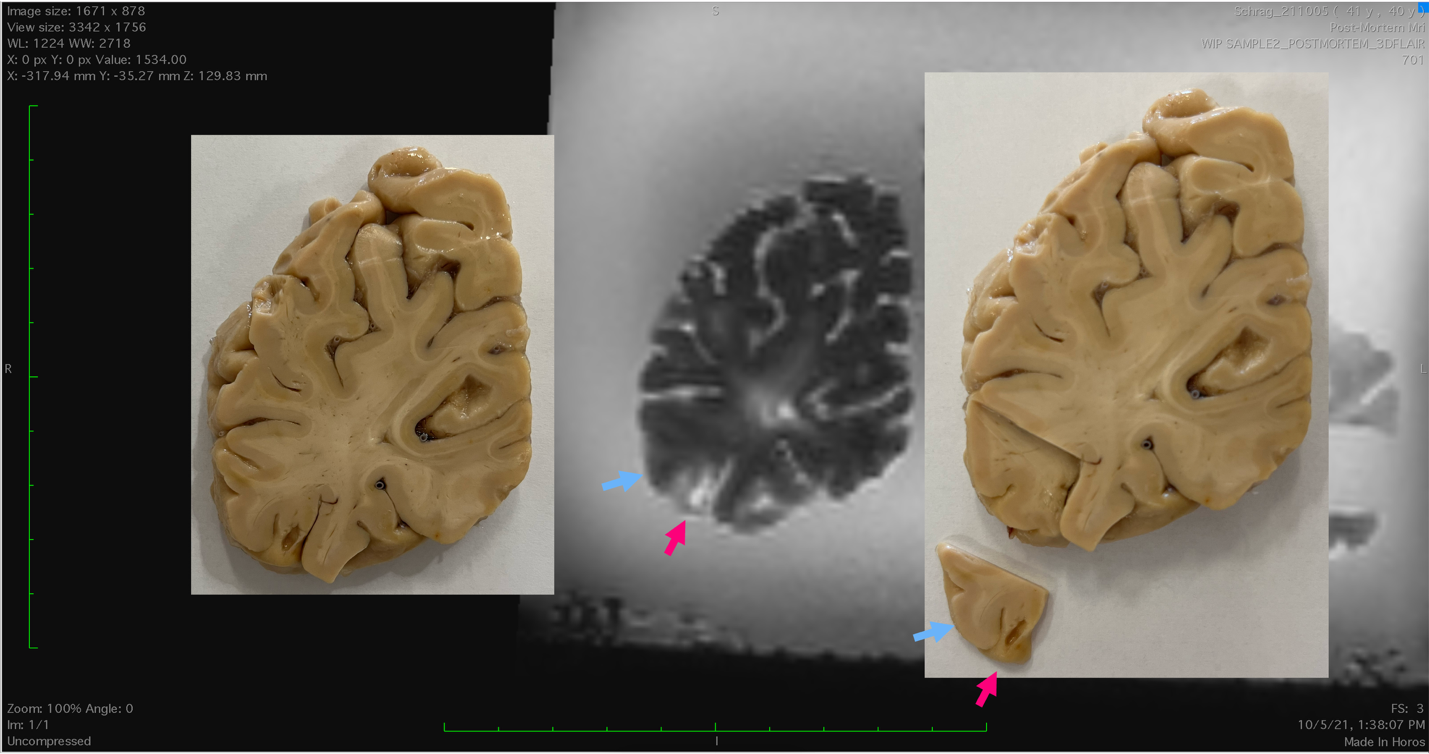

Donor A Posterior WMH and NAWM

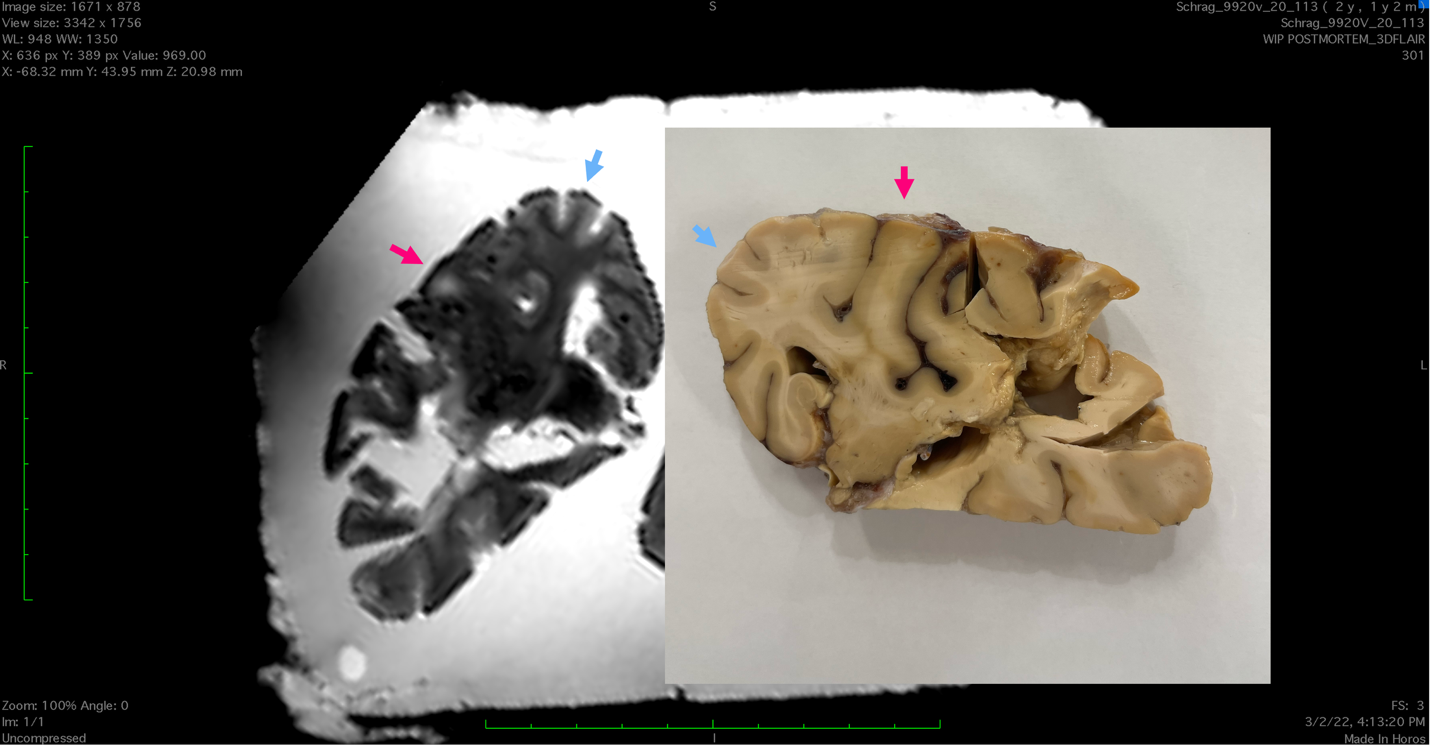

Donor B Posterior WMH and NAWM

*(Figure S2 continued on next page)*

**Figure S2 (continued)**

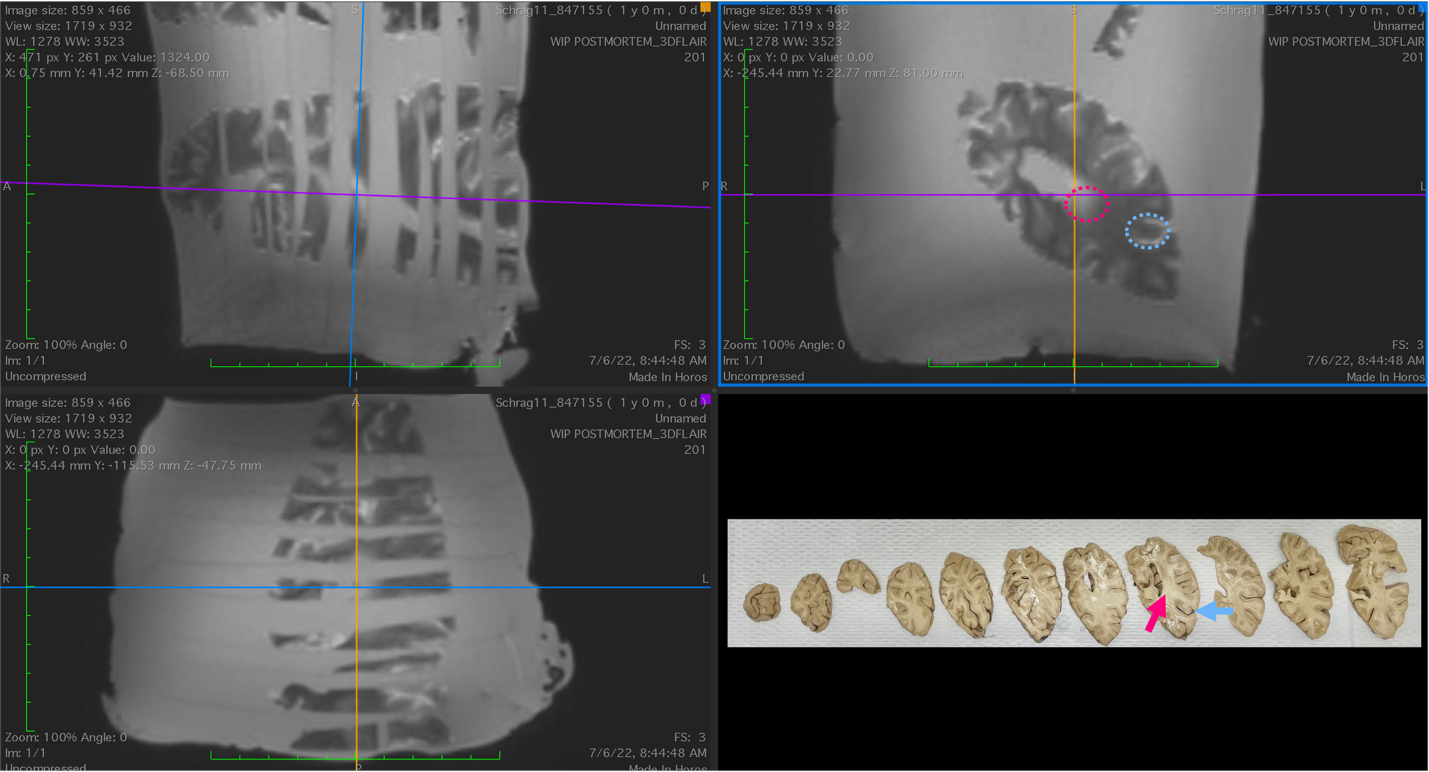

Donor C Posterior WMH and NAWM

***Figure S2.*** *Screen captures of T2-FLAIR MRI images with WMH (pink arrow) and NAWM (blue arrow). Photograph of corresponding brain tissue and dissected WMH and NAWM tissue blocks. Tissue dissections and region identifications were performed by collaborating pathologists and neurologists.*

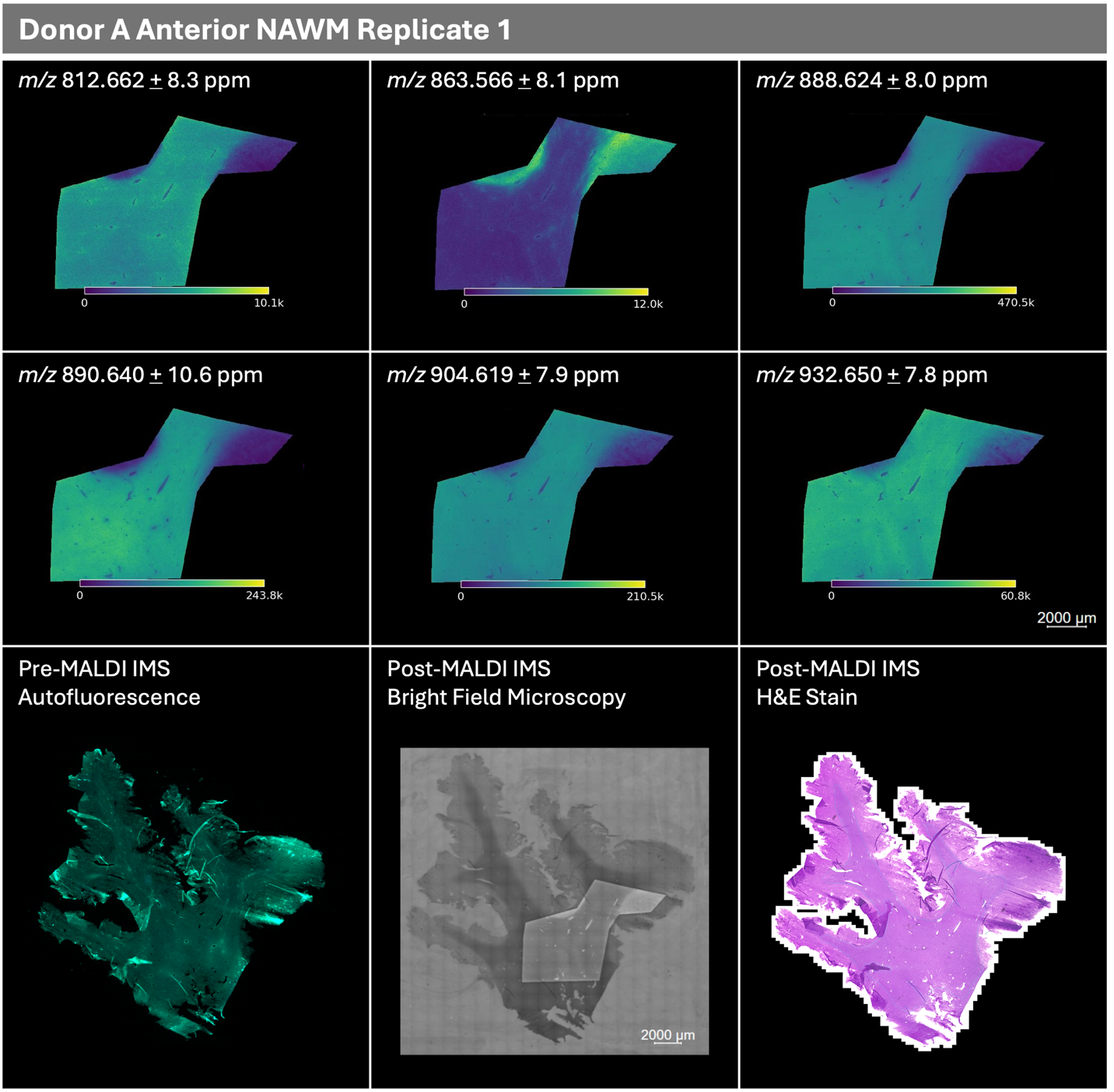

*(Figure S3 continued on next page)*

**Figure S3 (continued)**

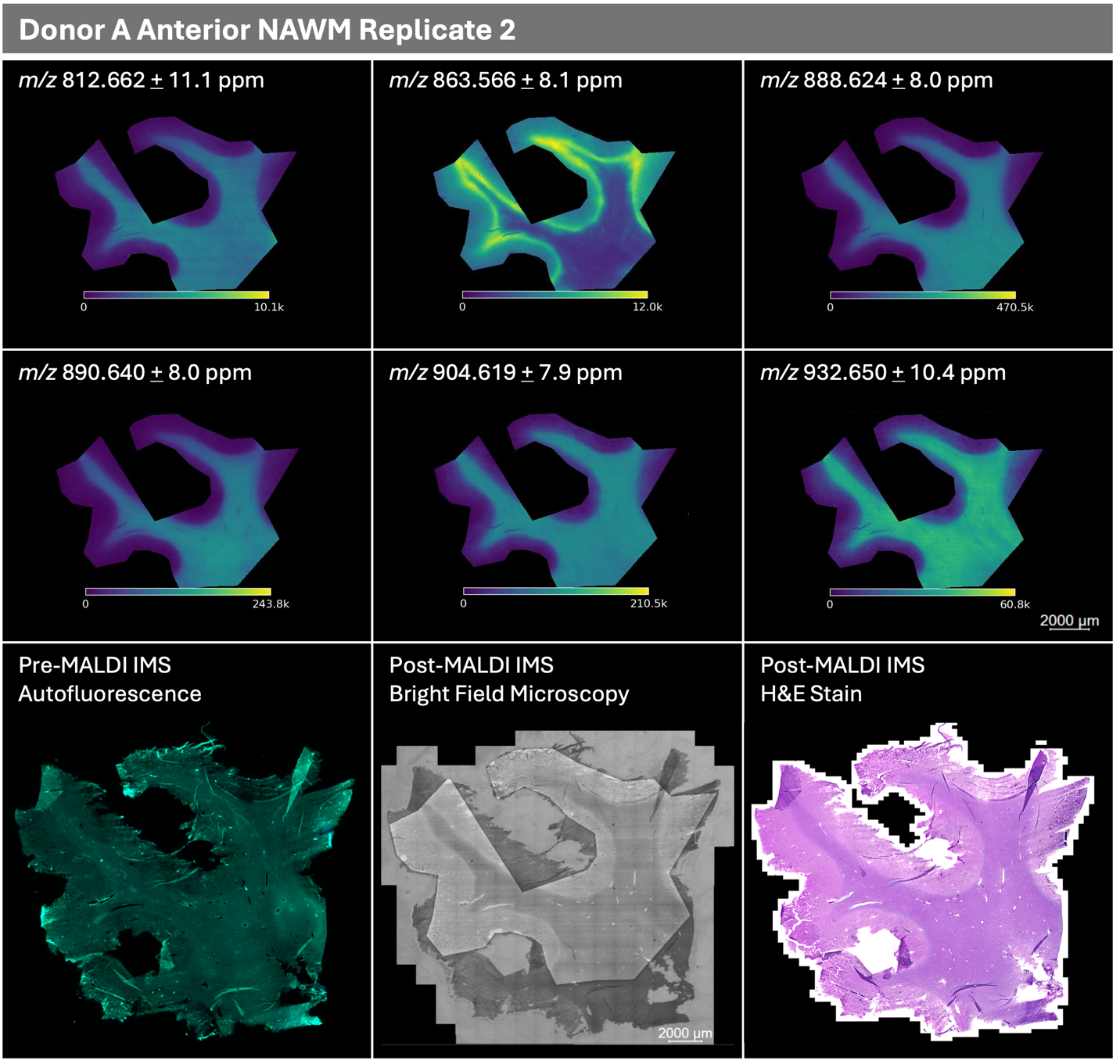

*(Figure S3 continued on next page)*

**Figure S3 (continued)**
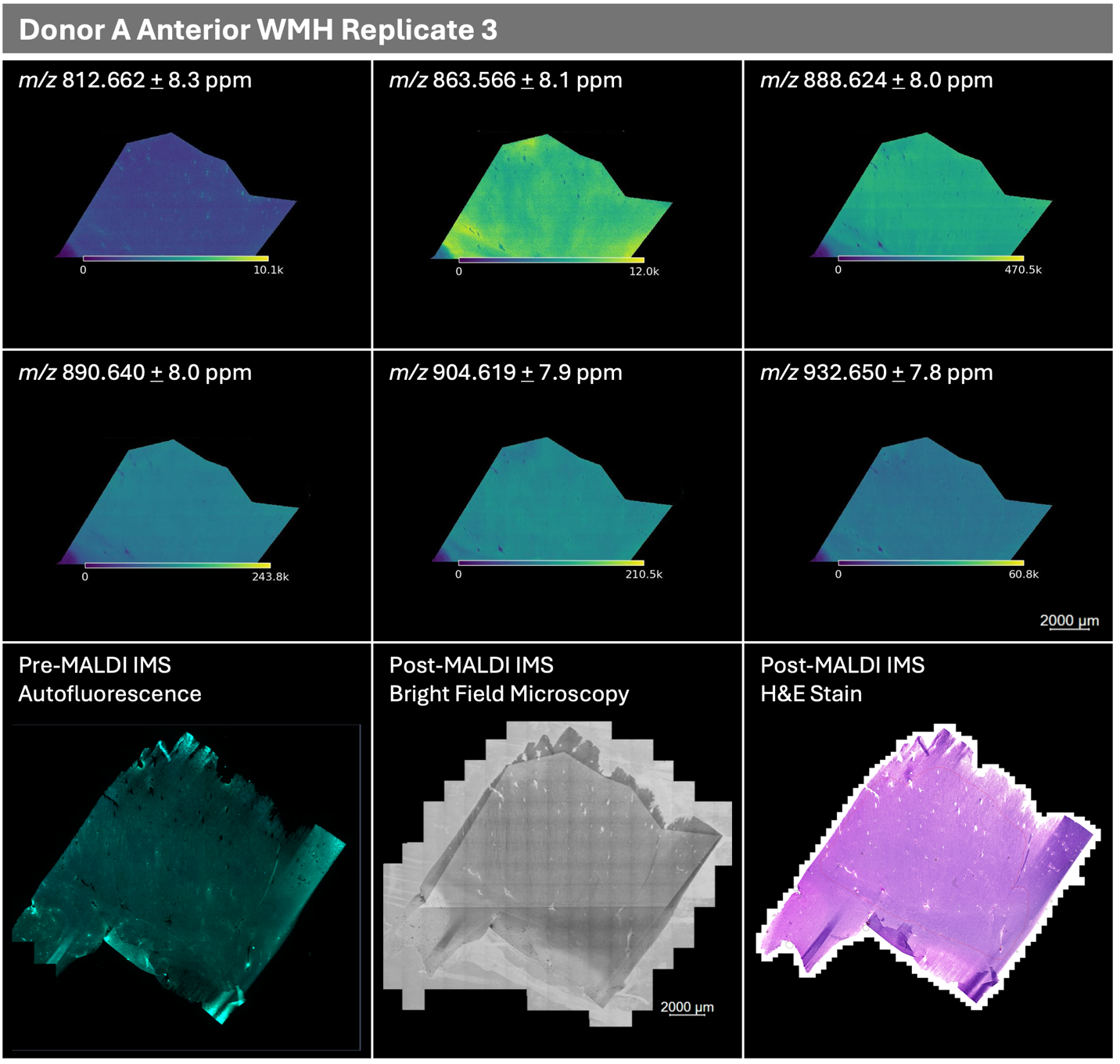

*(Figure S3 continued on next page)*

**Figure S3 (continued)**

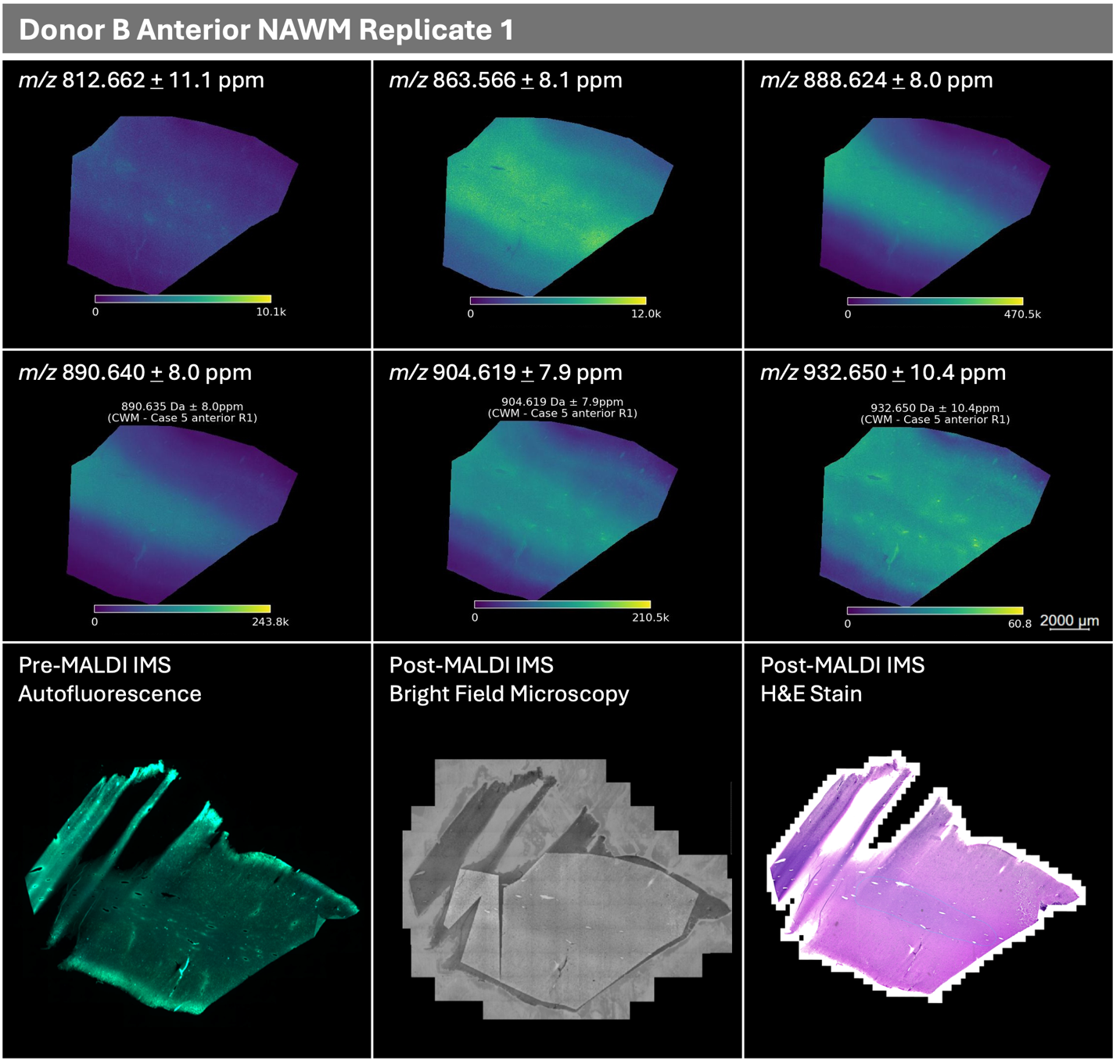

*(Figure S3 continued on next page)*

**Figure S3 (continued)**
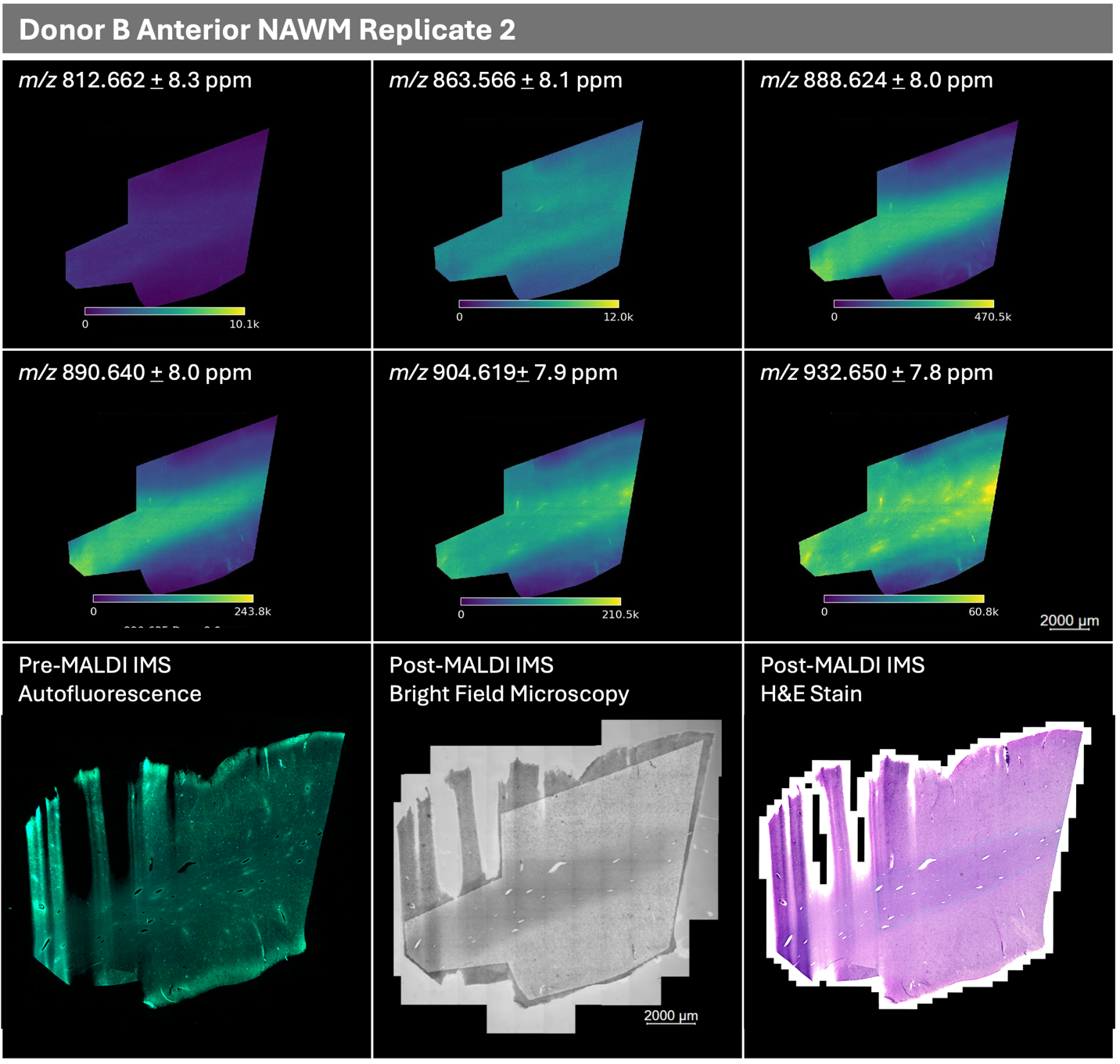

*(Figure S3 continued on next page)*

**Figure S3 (continued)**

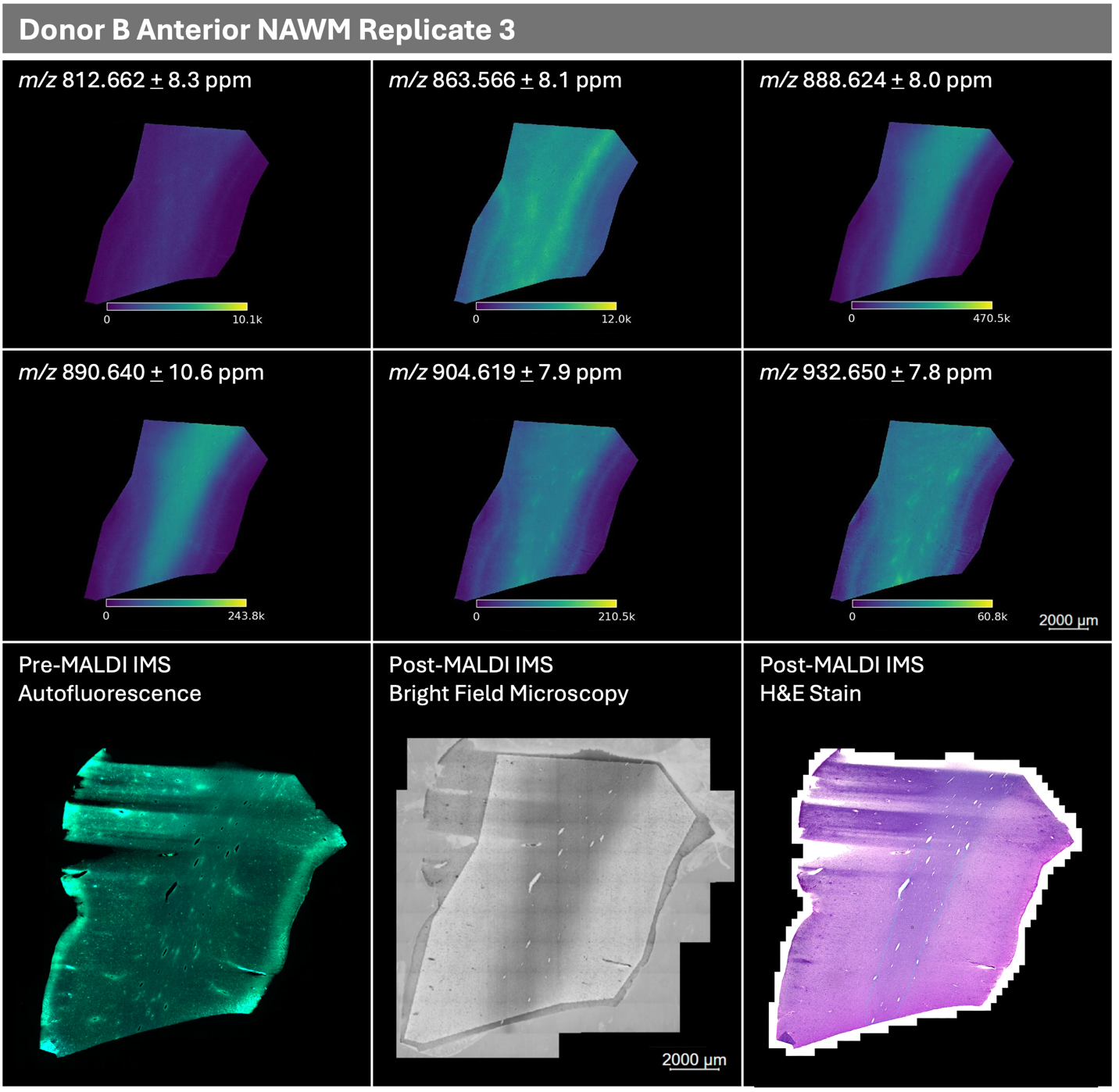

*(Figure S3 continued on next page)*

**Figure S3 (continued)**
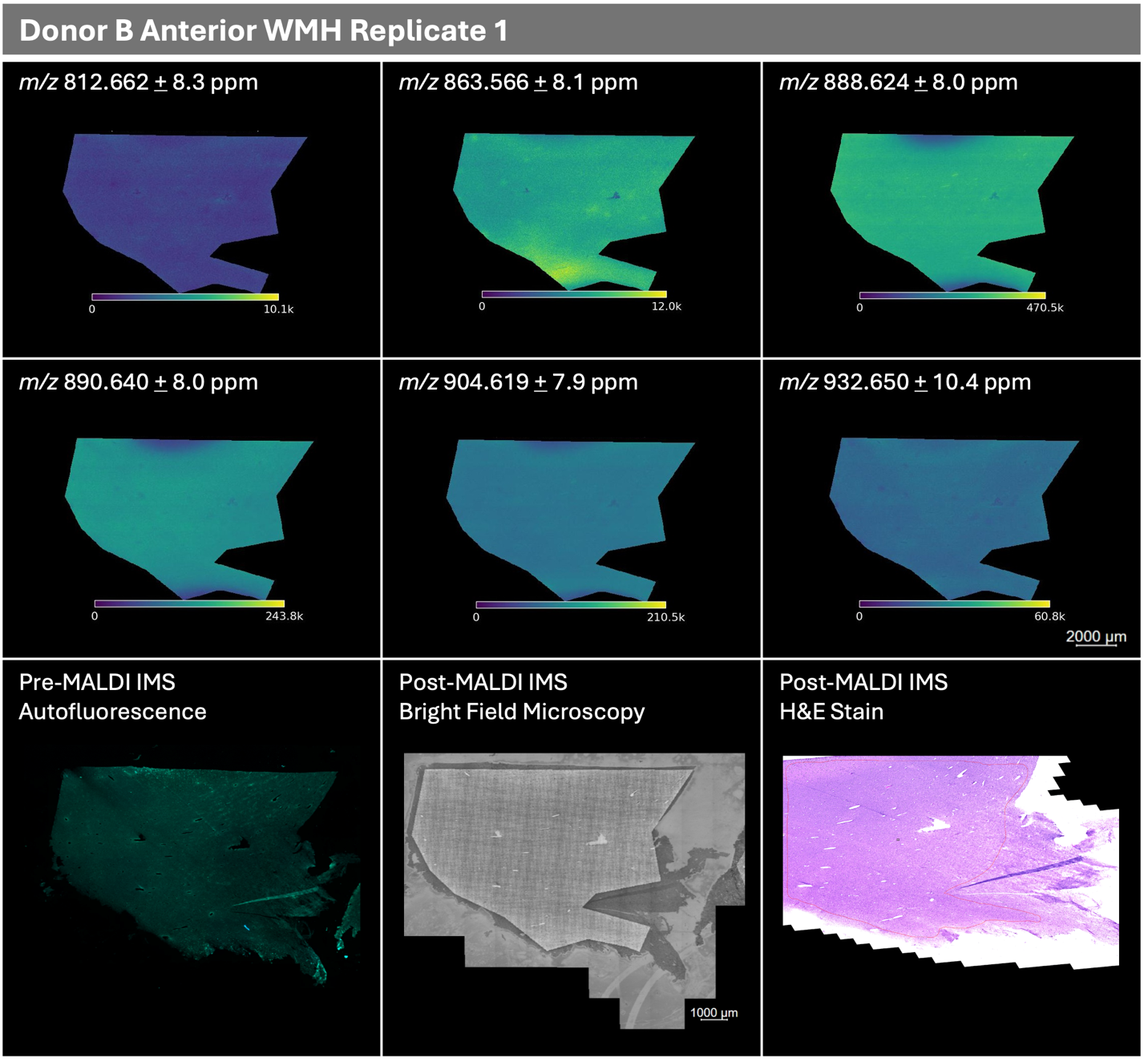

*(Figure S3 continued on next page)*

**Figure S3 (continued)**

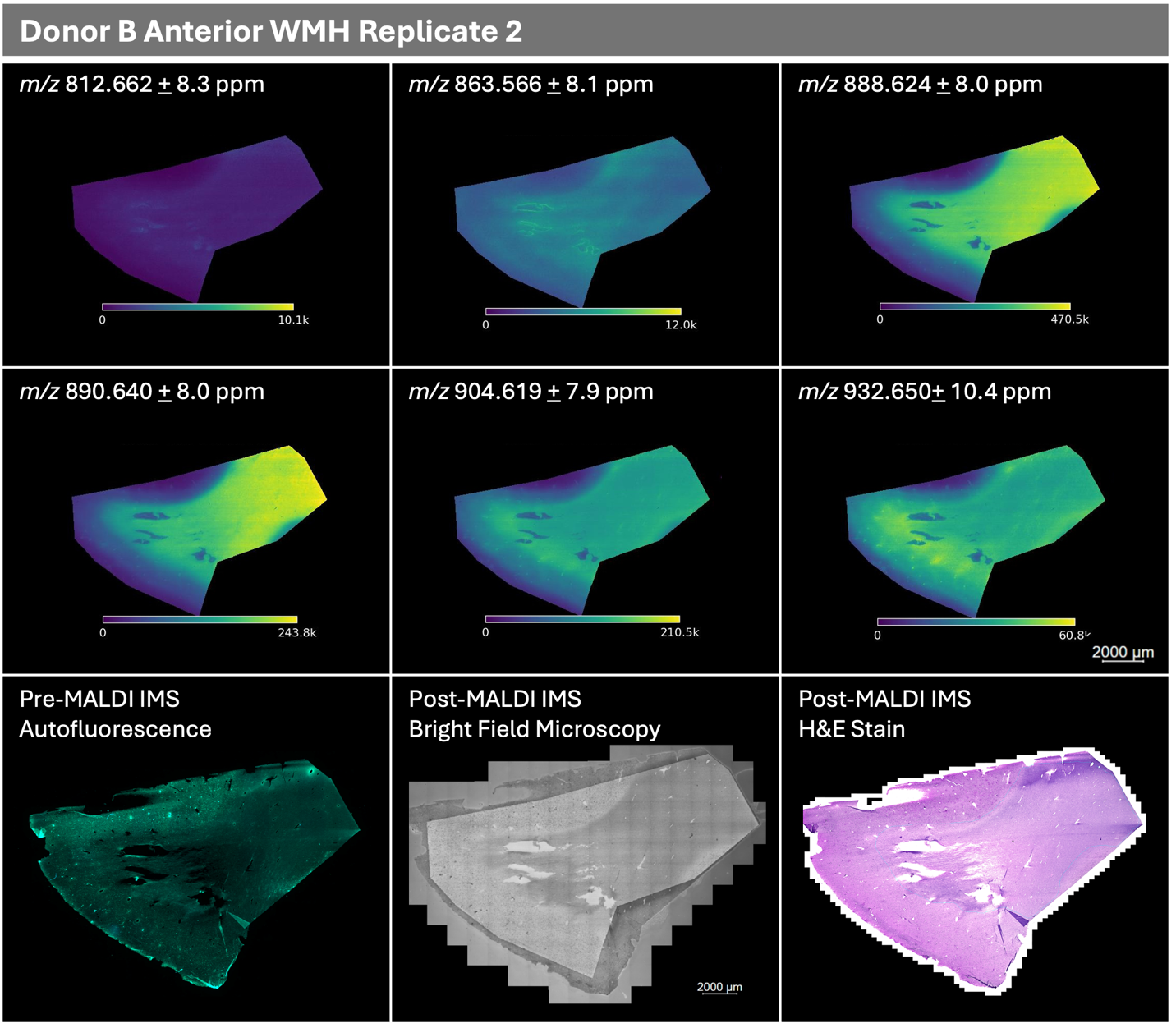

*(Figure S3 continued on next page)*

**Figure S3 (continued)**
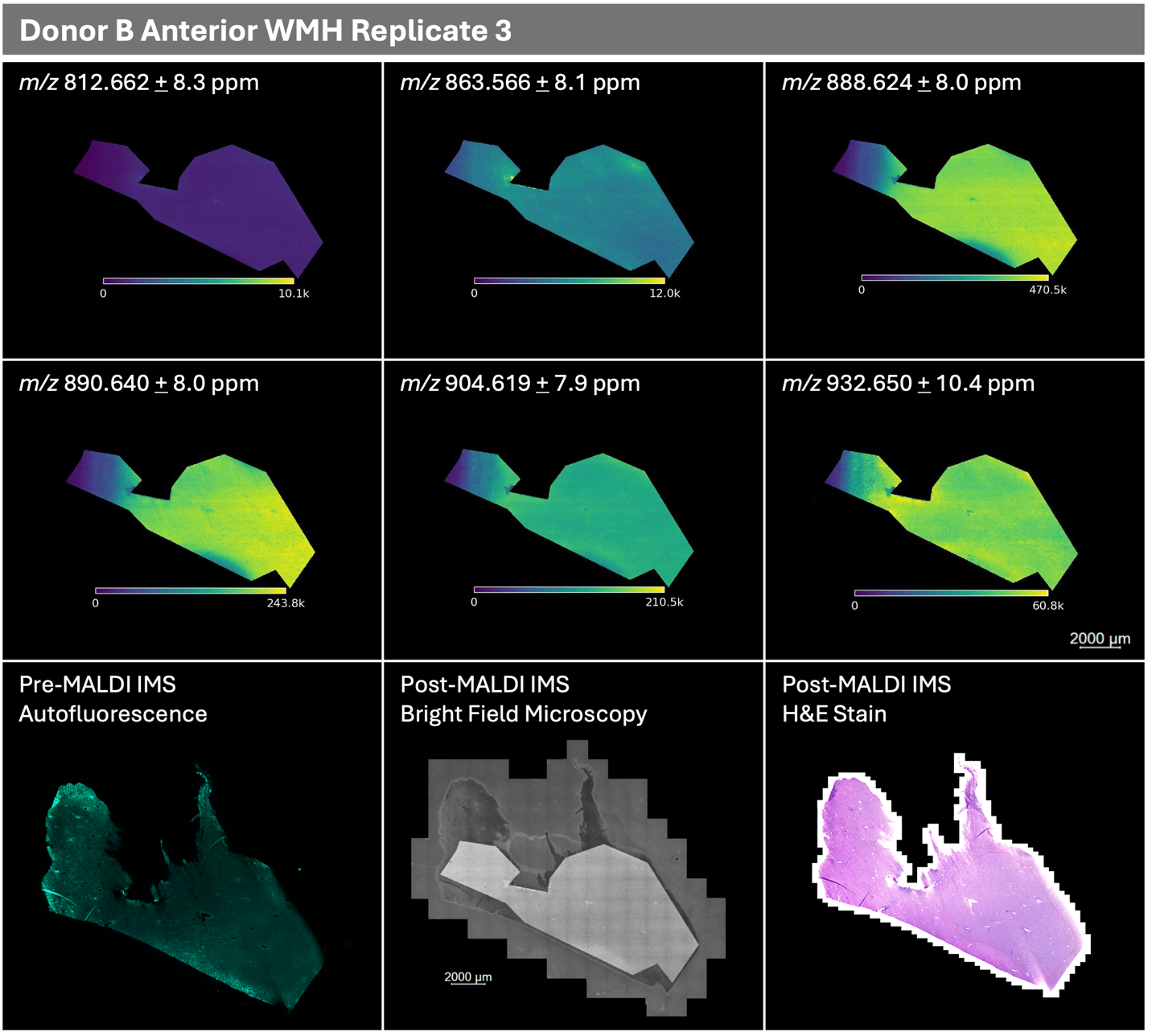

*(Figure S3 continued on next page)*

**Figure S3 (continued)**

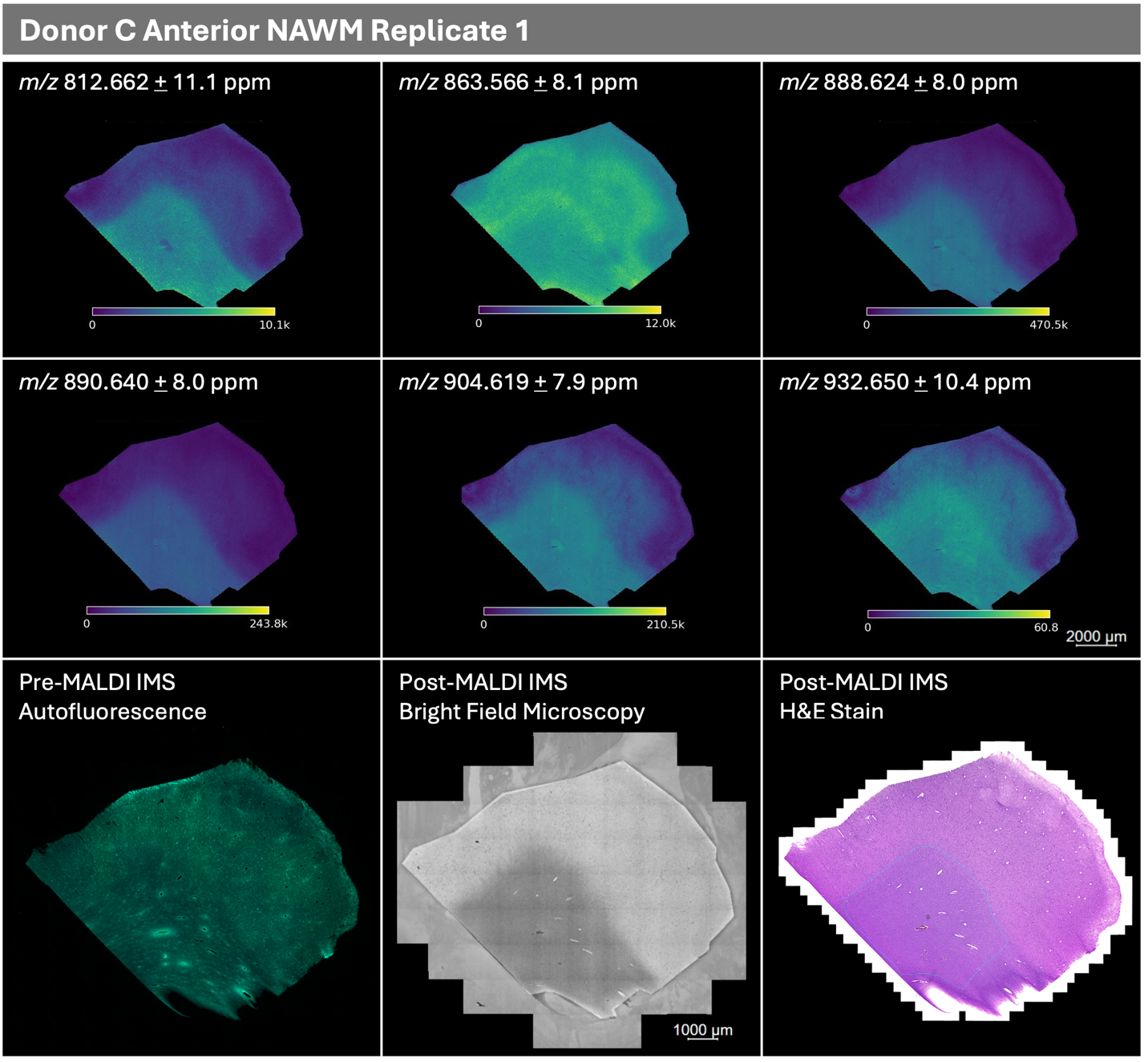

*(Figure S3 continued on next page)*

**Figure S3 (continued)**
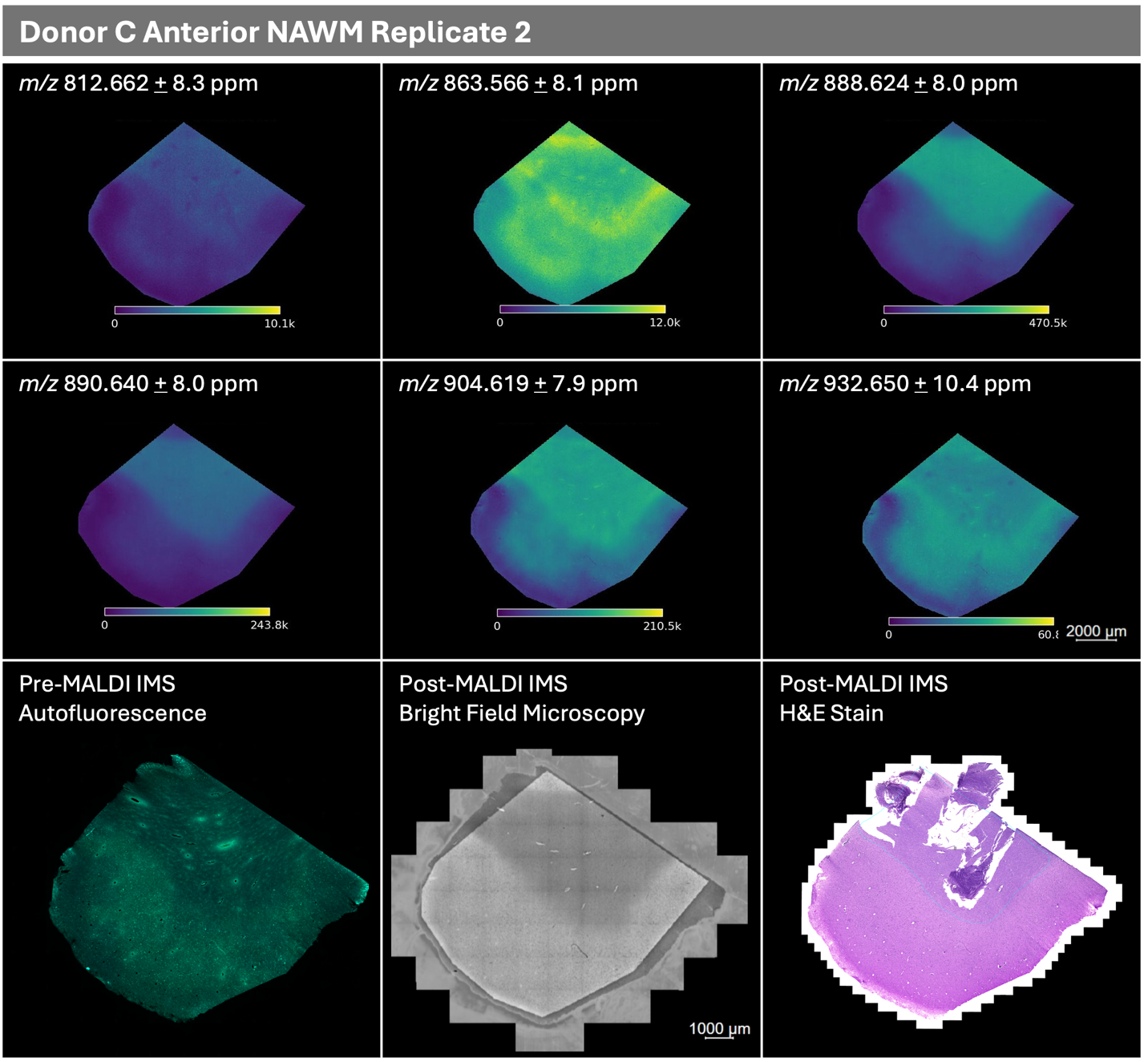

*(Figure S3 continued on next page)*

**Figure S3 (continued)**
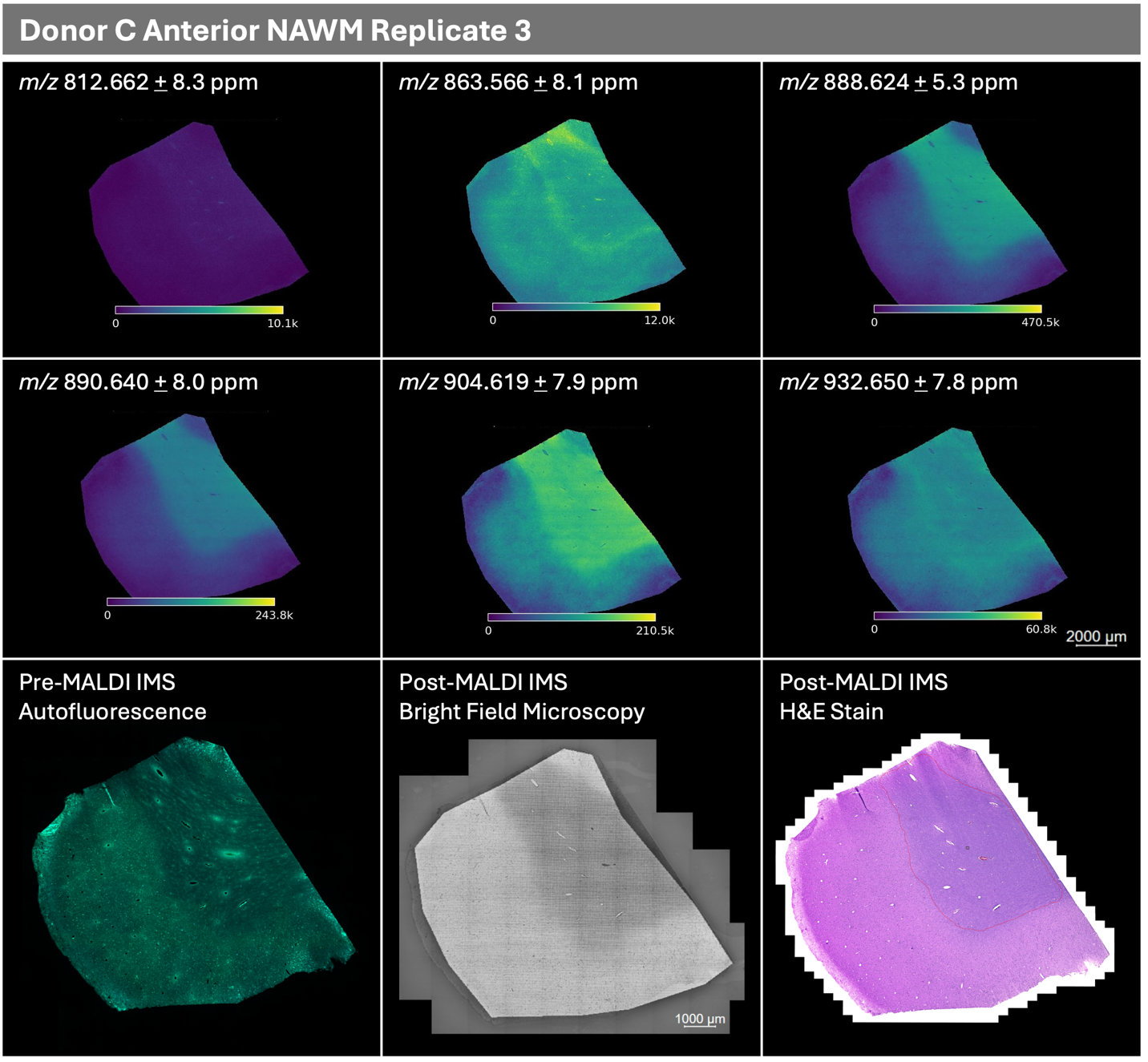

*(Figure S3 continued on next page)*

**Figure S3 (continued)**
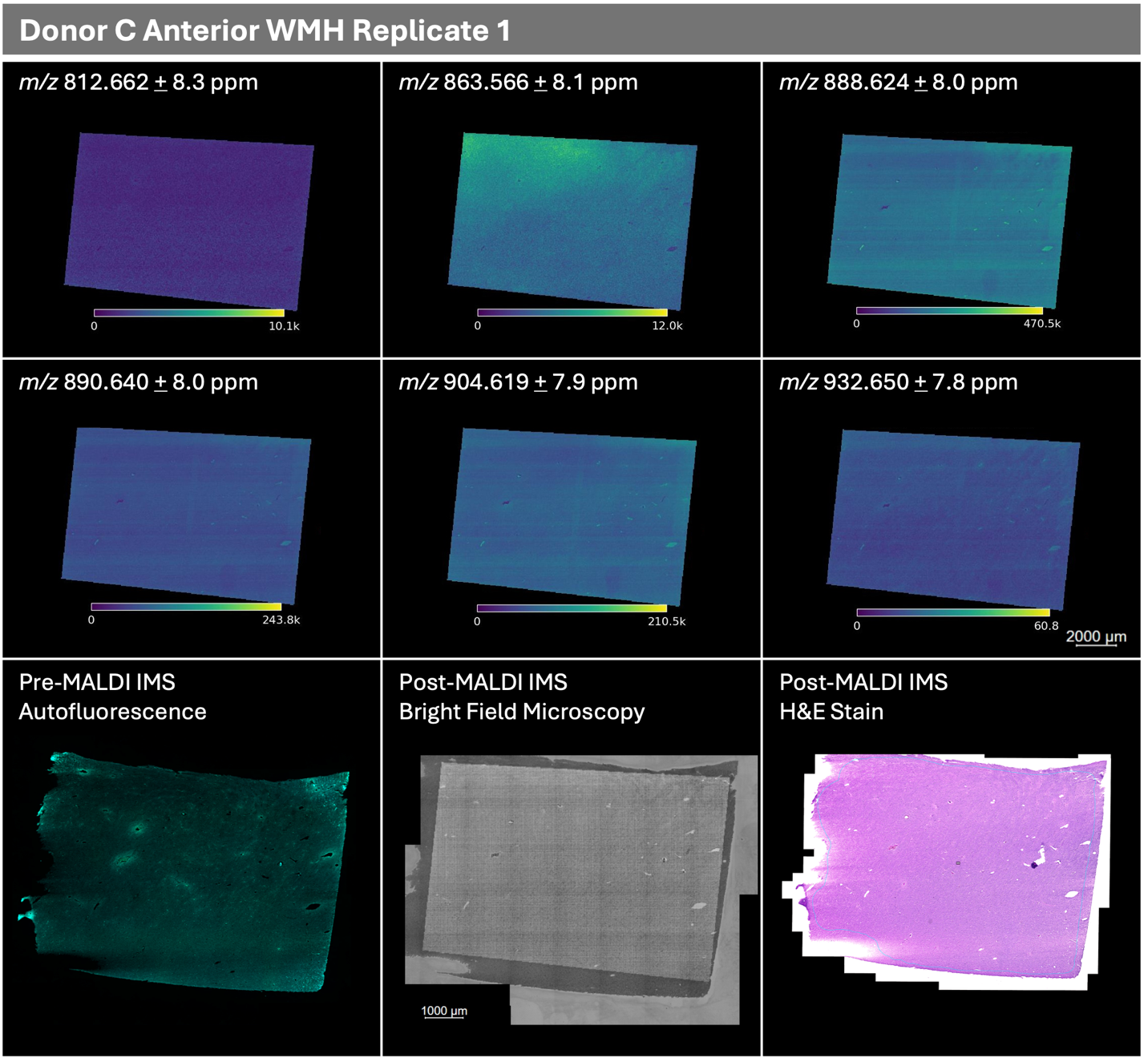

*(Figure S3 continued on next page)*

**Figure S3 (continued)**
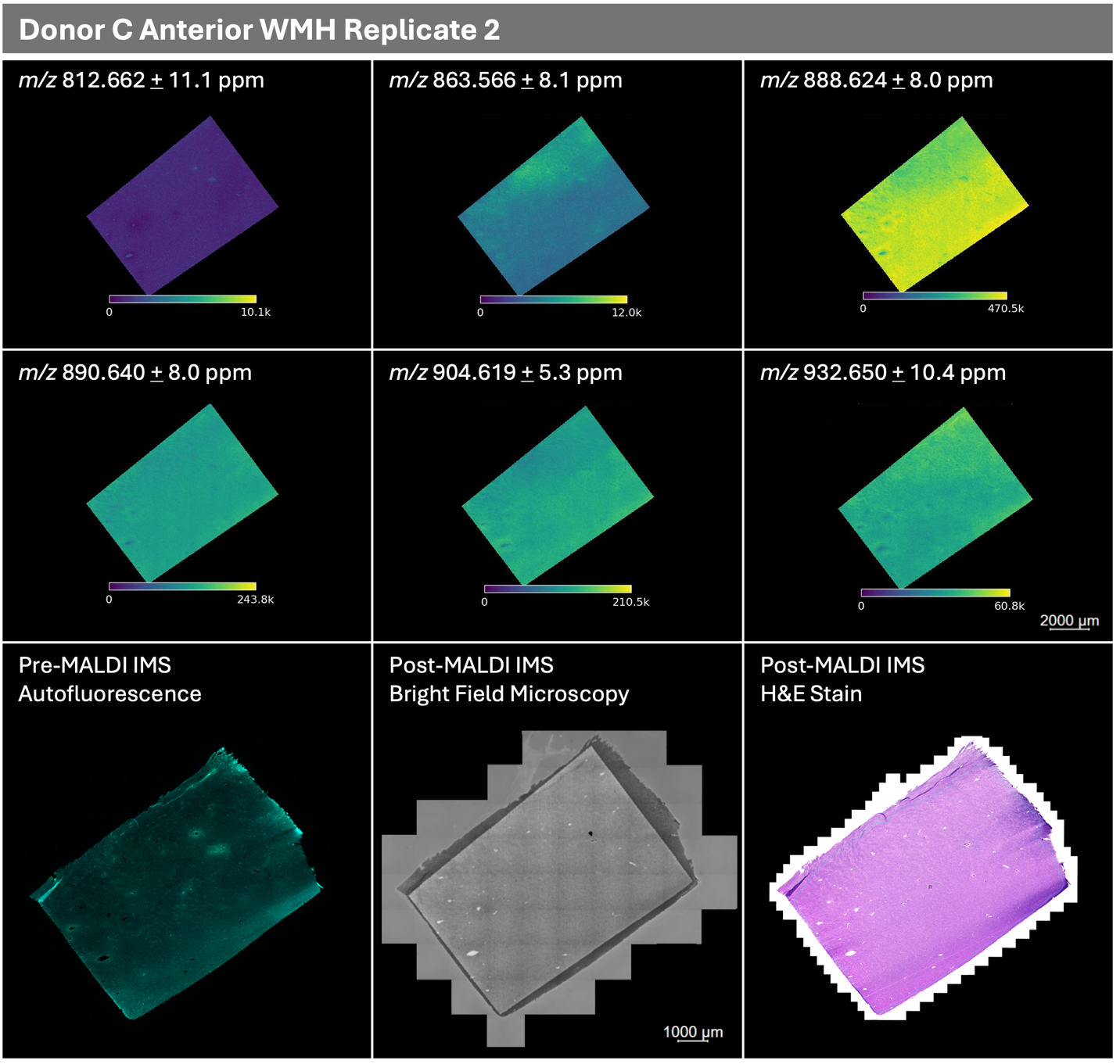

*(Figure S3 continued on next page)*

**Figure S3 (continued)**
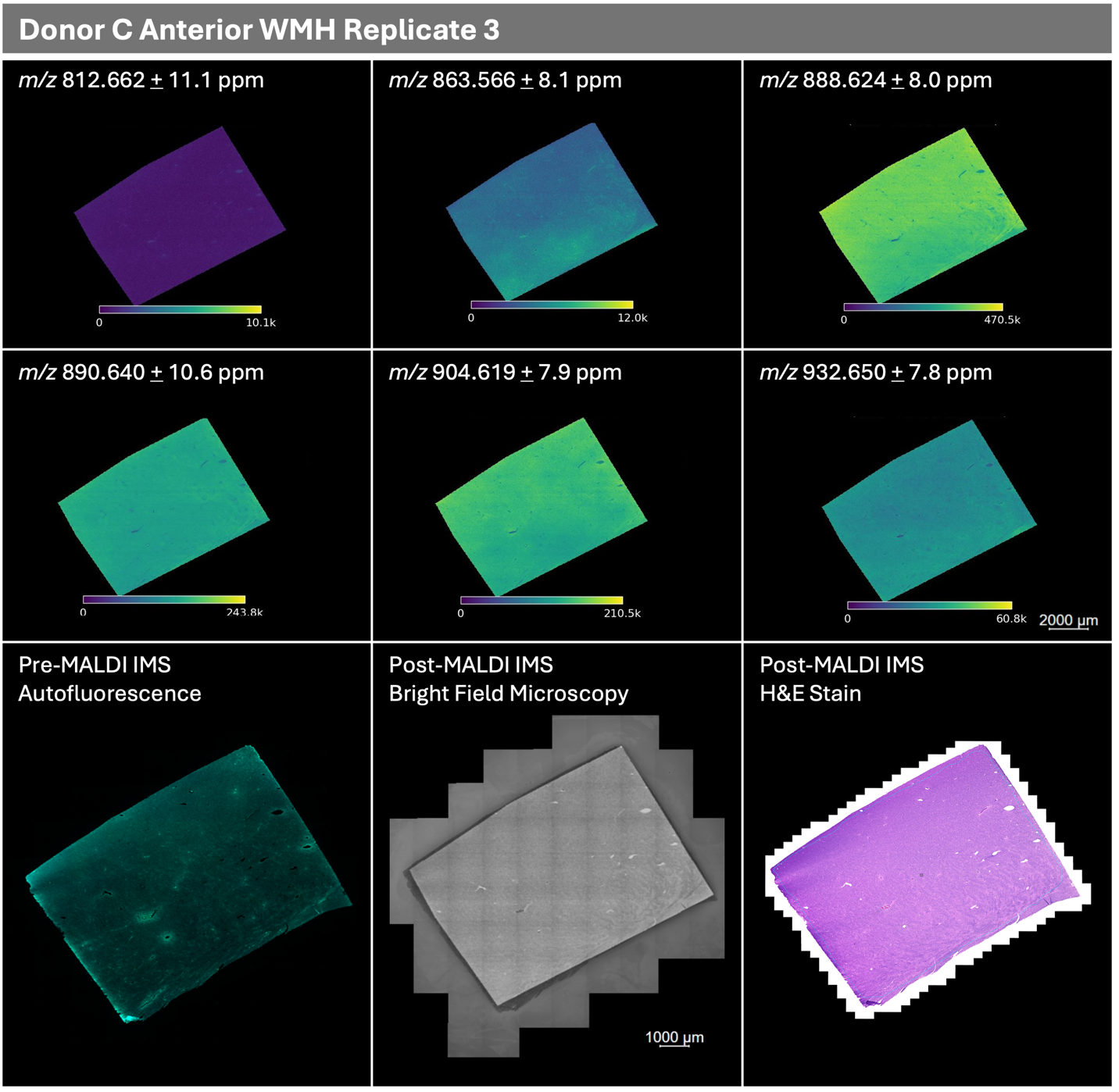

*(Figure S3 continued on next page)*

**Figure S3 (continued)**
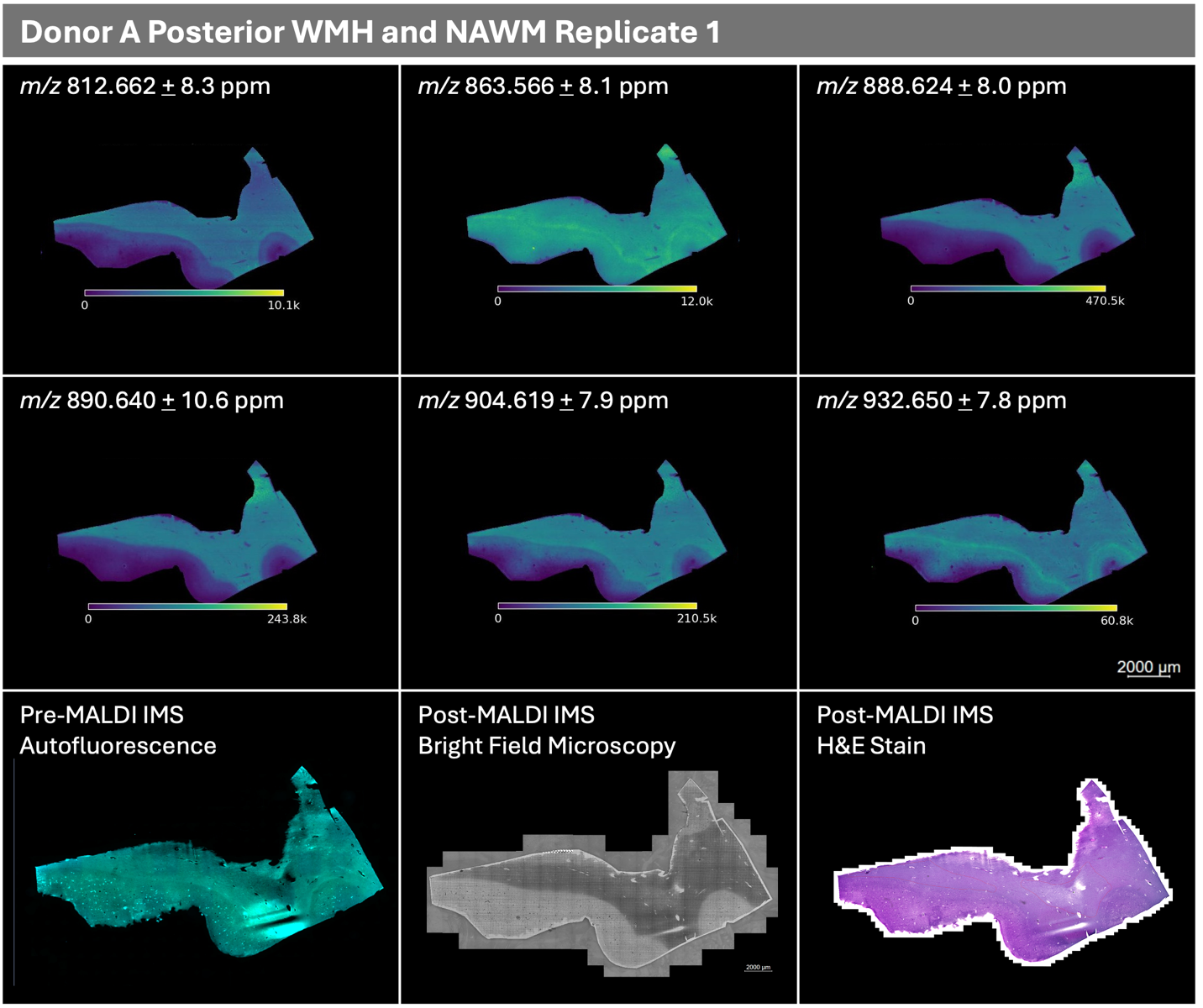

*(Figure S3 continued on next page)*

**Figure S3 (continued)**
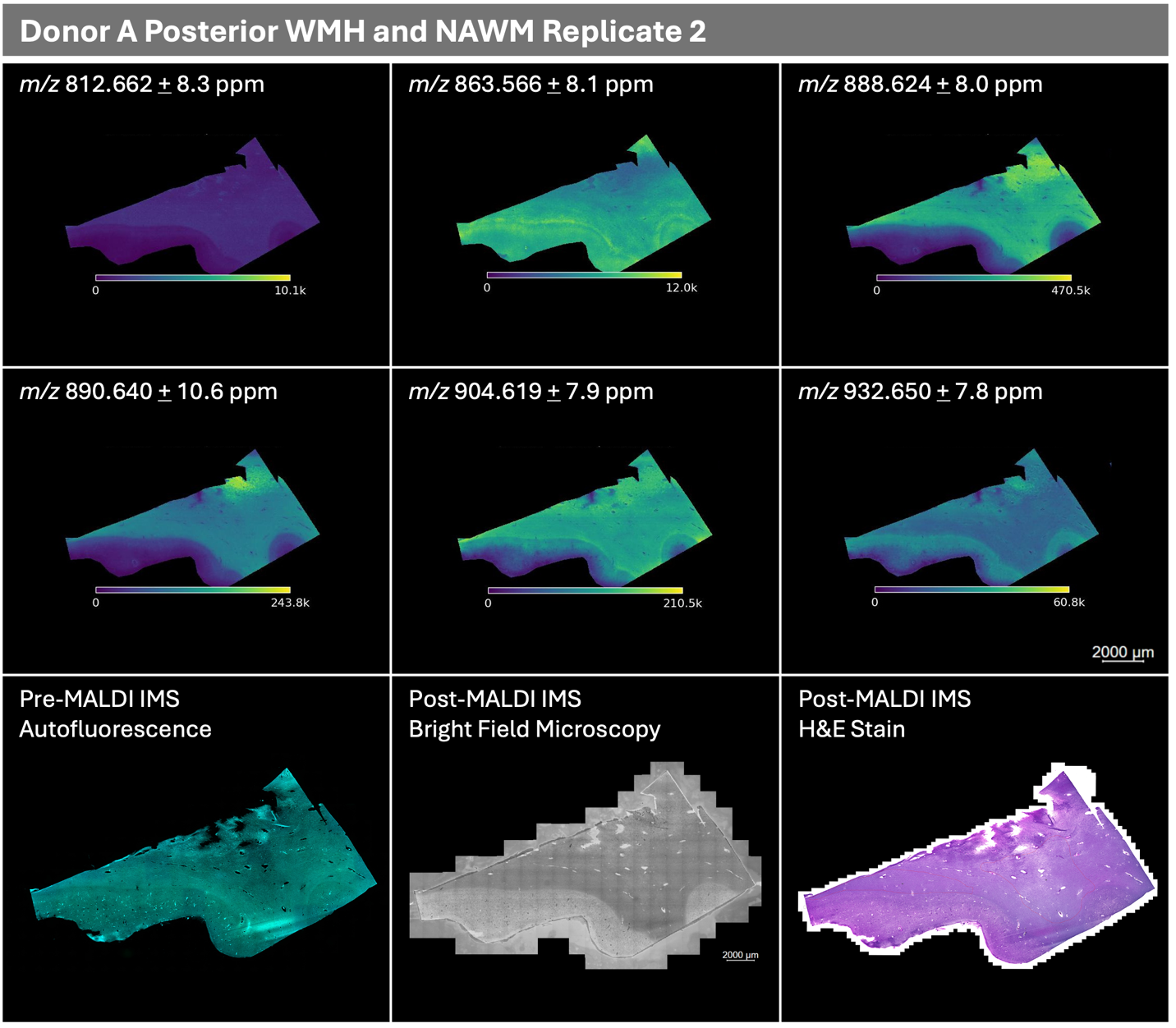

*(Figure S3 continued on next page)*

**Figure S3 (continued)**

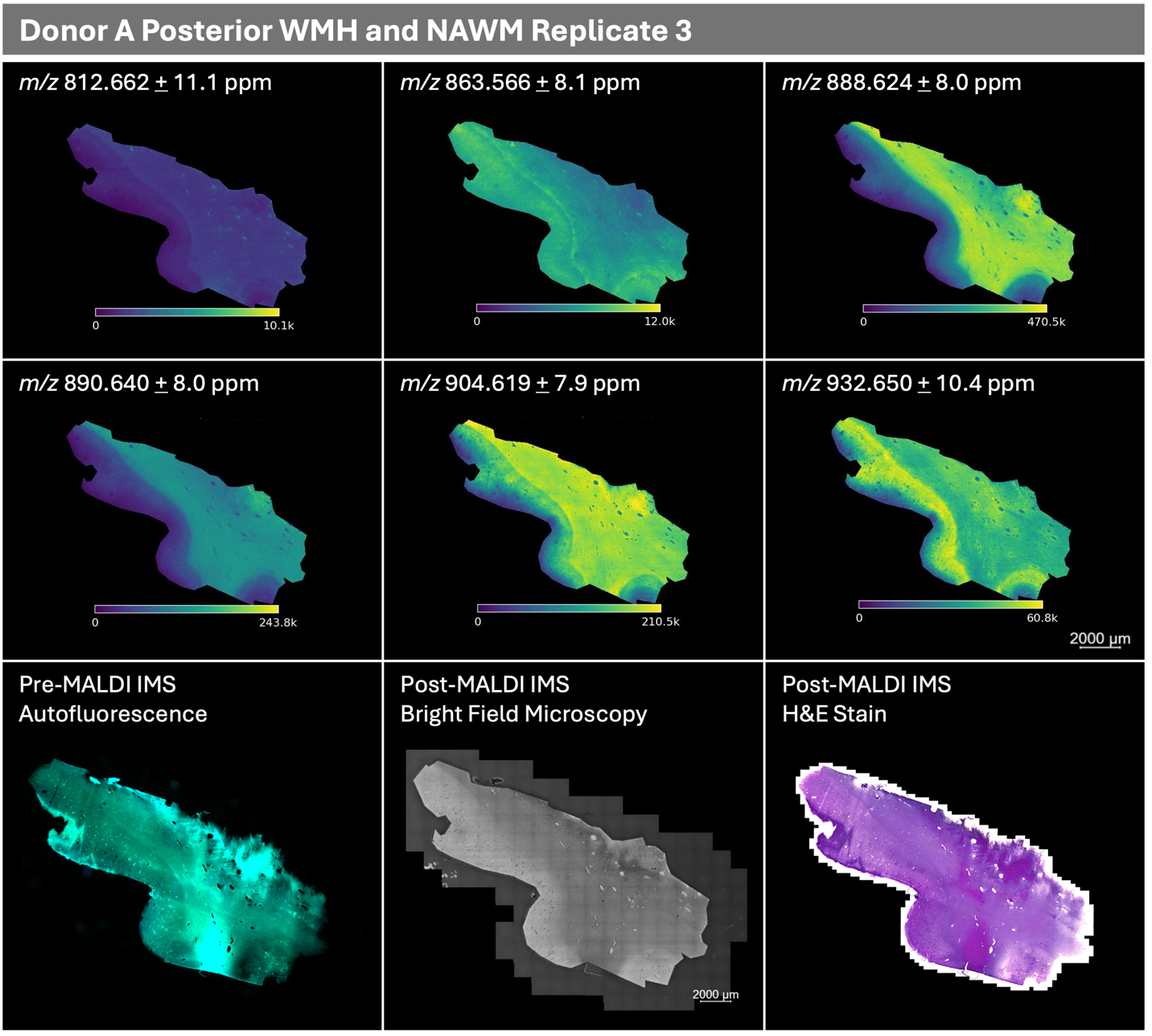

*(Figure S3 continued on next page)*

**Figure S3 (continued)**
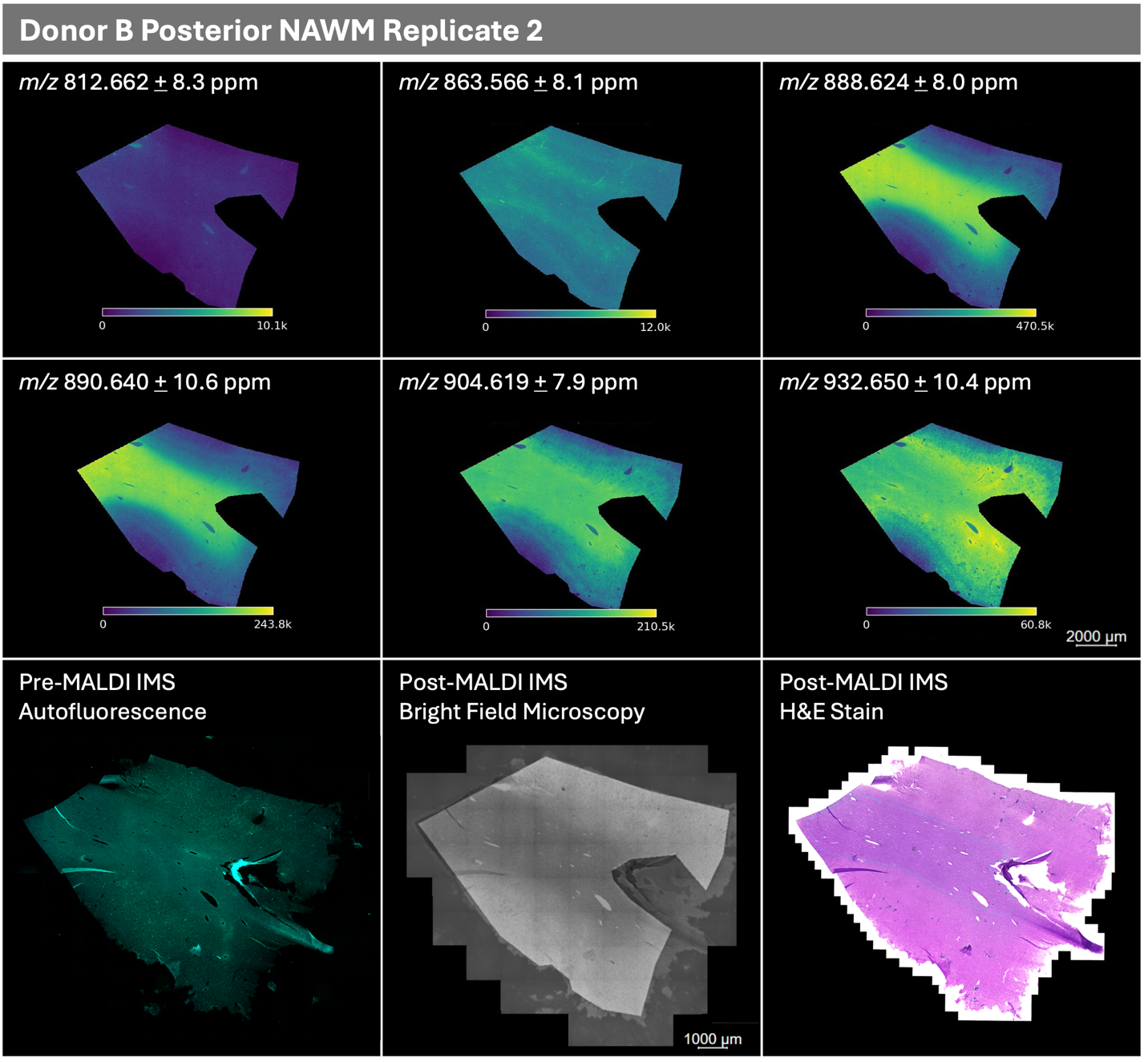

*(Figure S3 continued on next page)*

**Figure S3 (continued)**
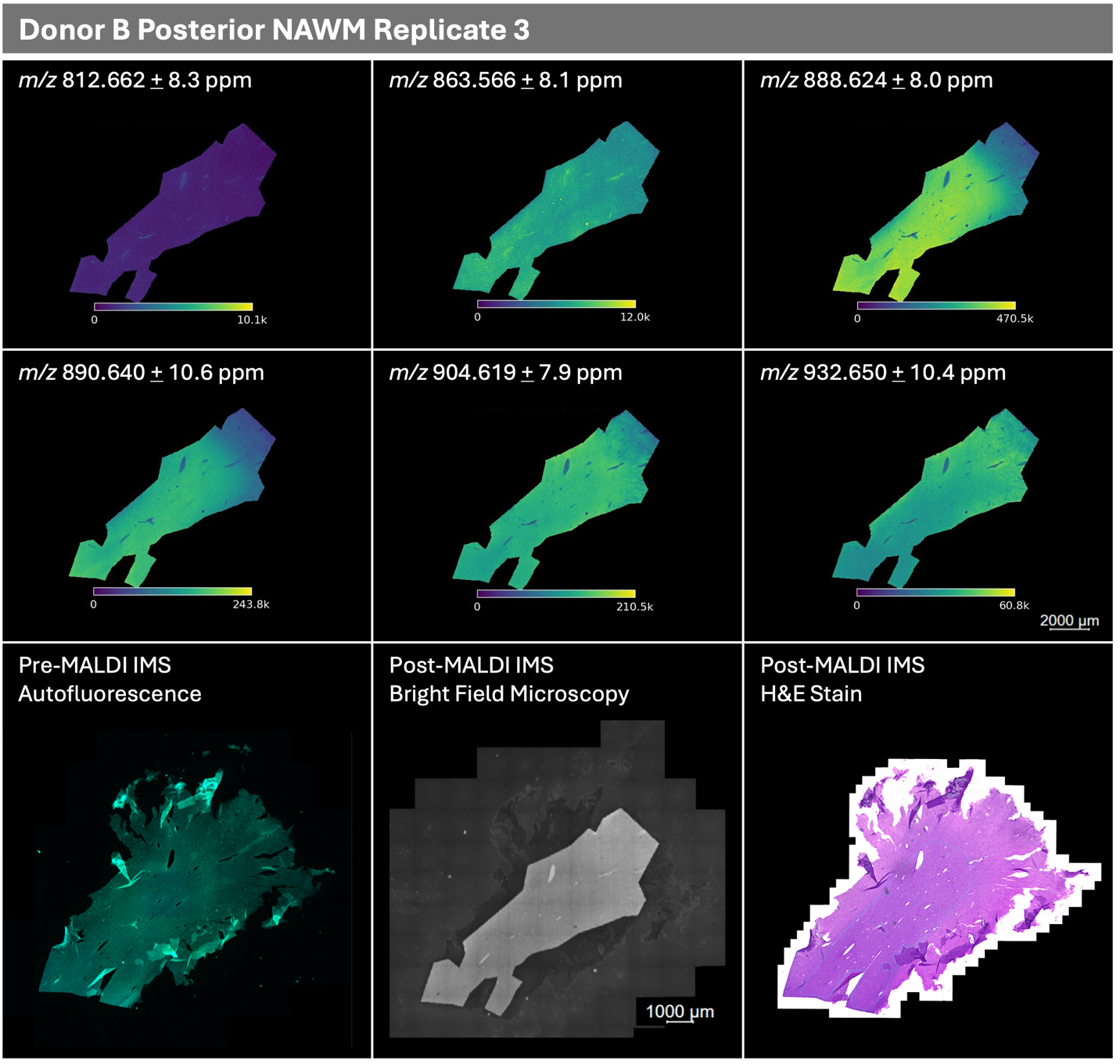

*(Figure S3 continued on next page)*

**Figure S3 (continued)**
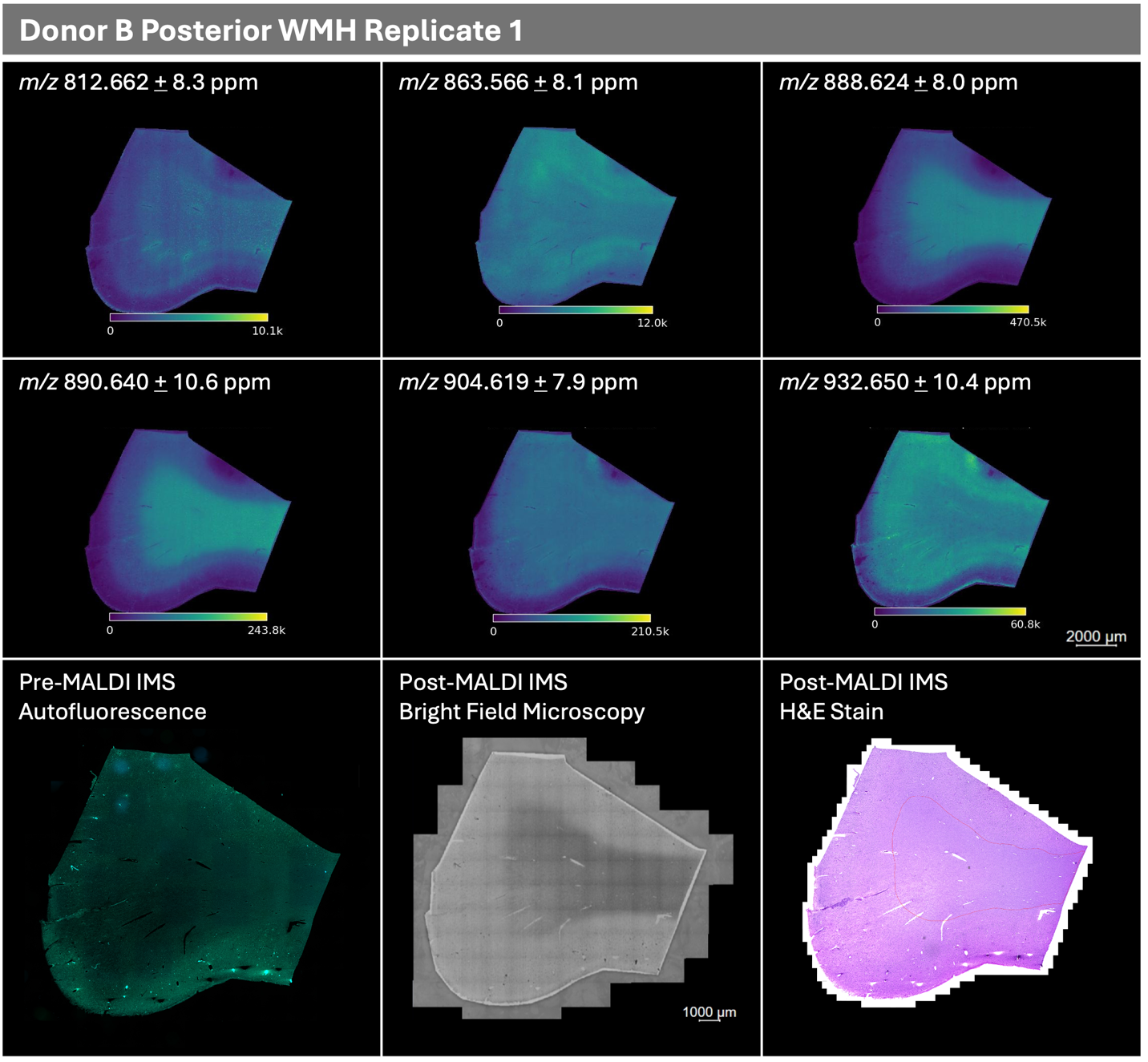

*(Figure S3 continued on next page)*

**Figure S3 (continued)**
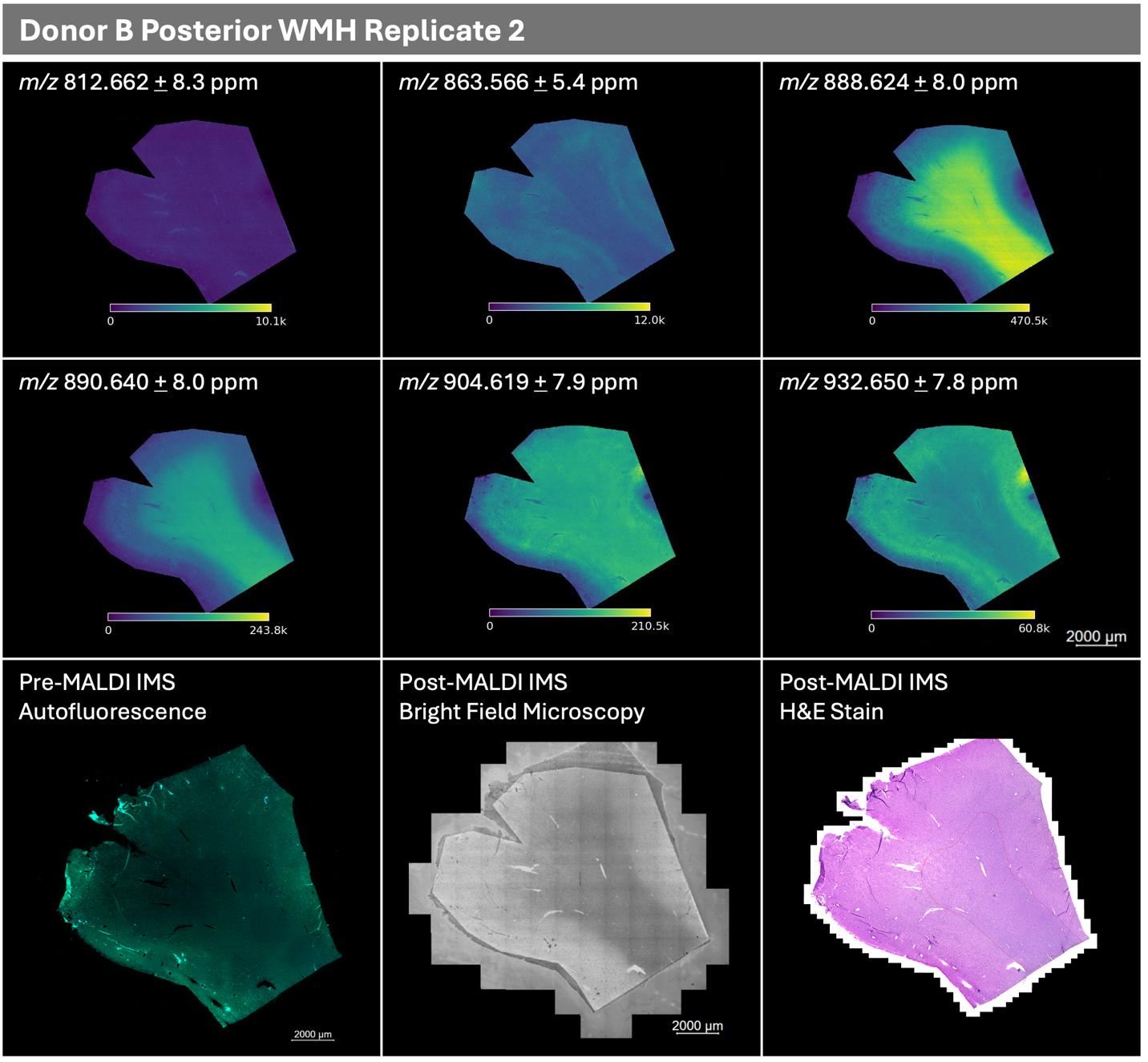

*(Figure S3 continued on next page)*

**Figure S3 (continued)**
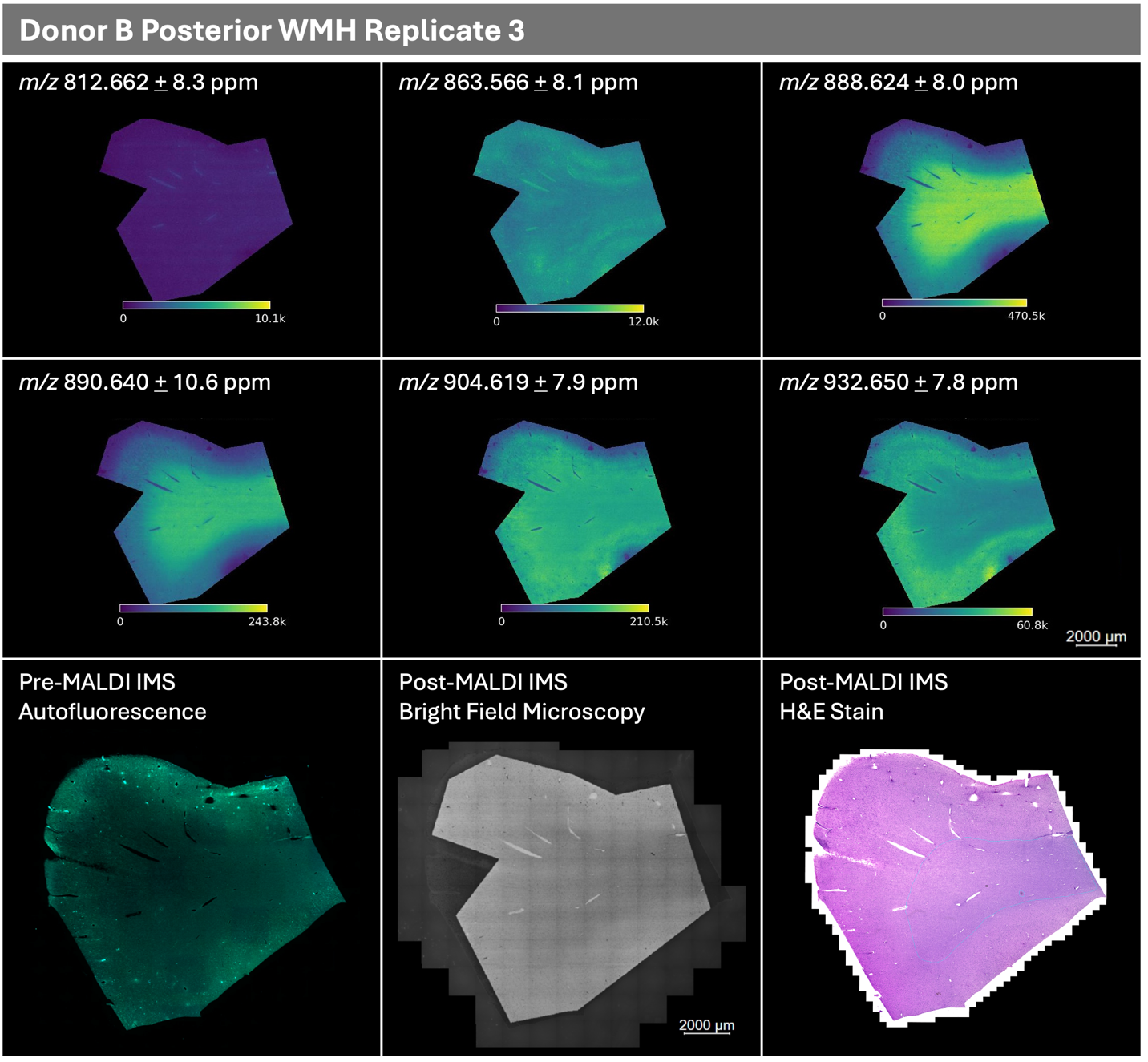

*(Figure S3 continued on next page)*

**Figure S3 (continued)**
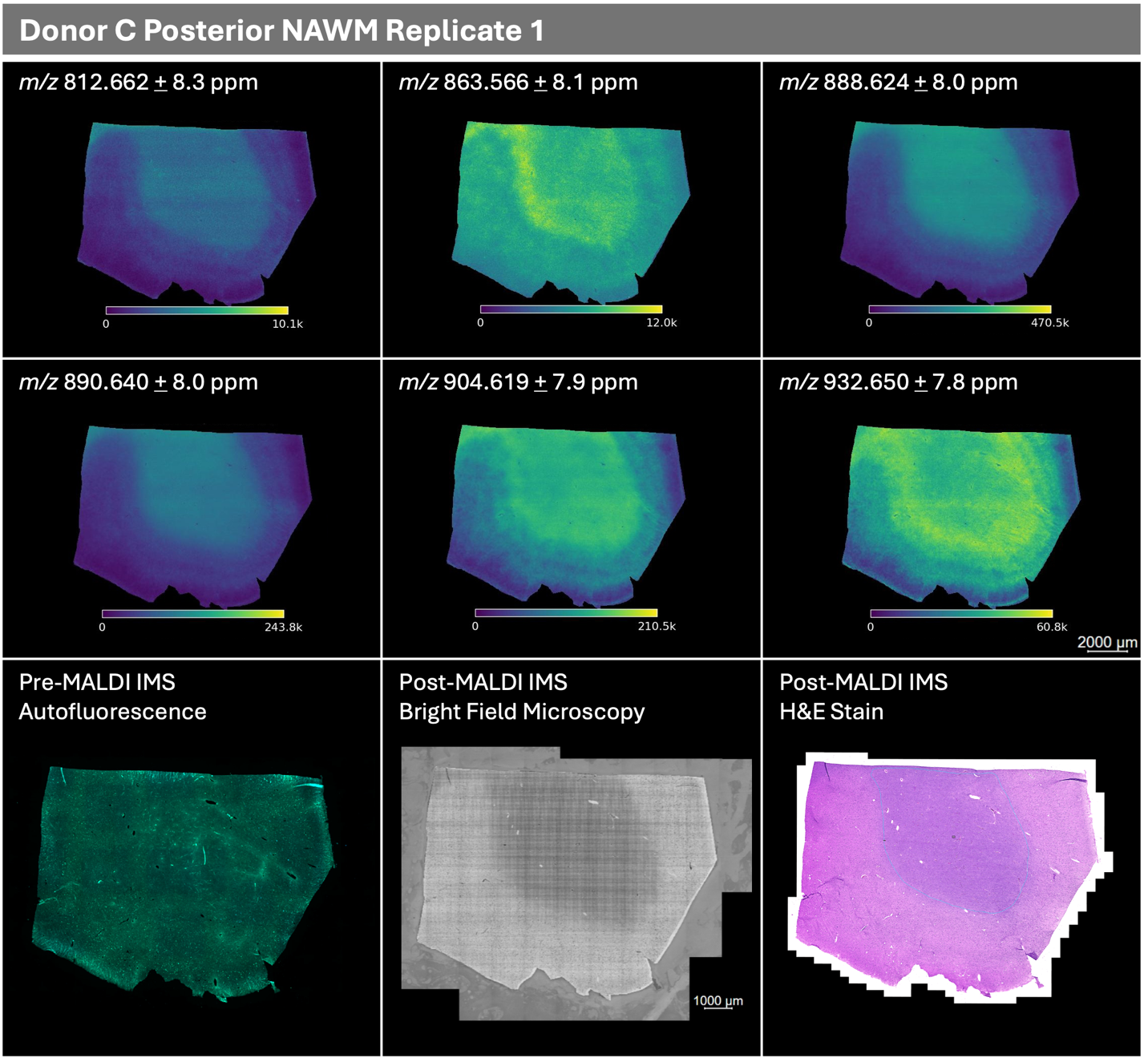

*(Figure S3 continued on next page)*

**Figure S3 (continued)**
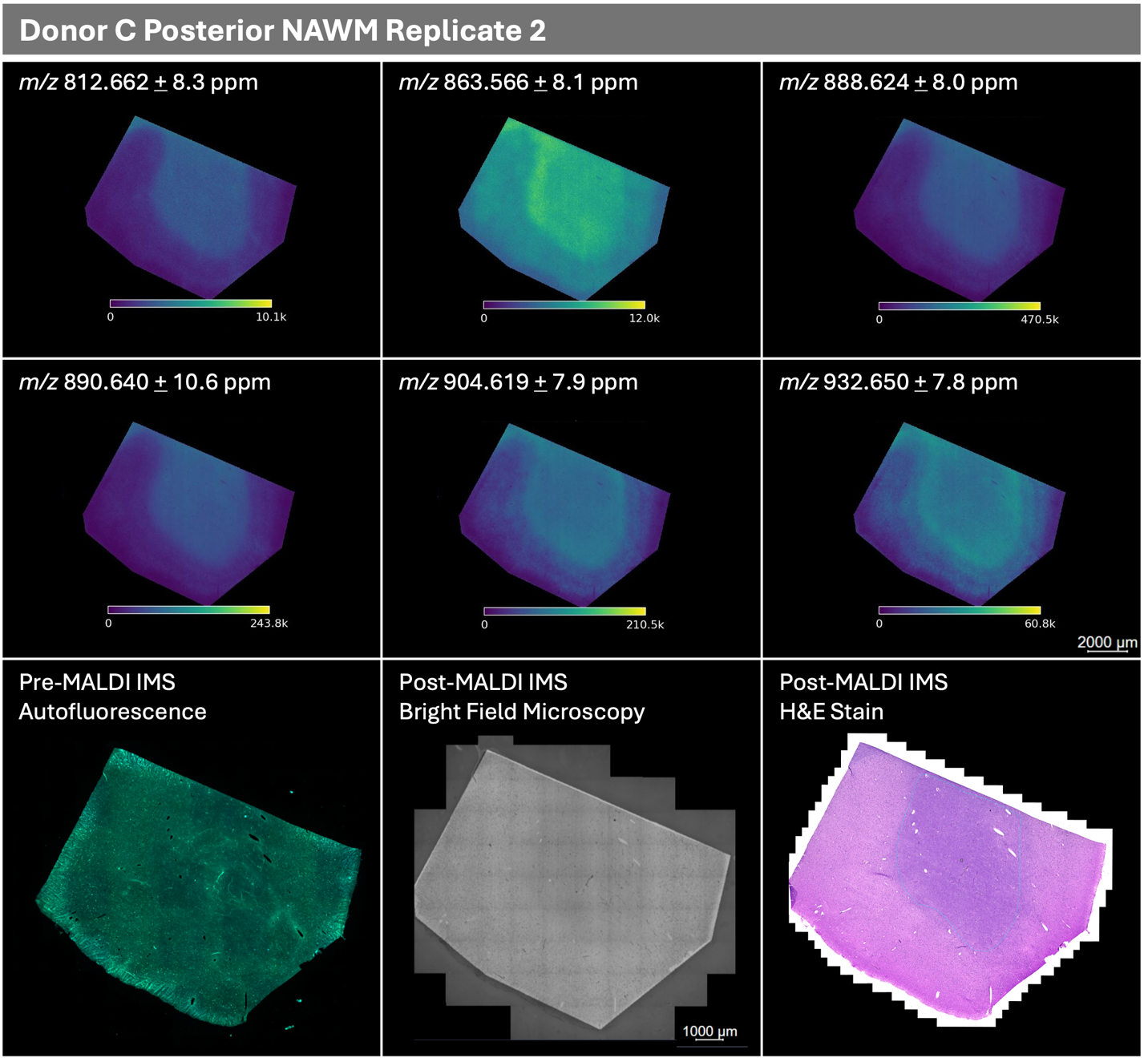

*(Figure S3 continued on next page)*

**Figure S3 (continued)**
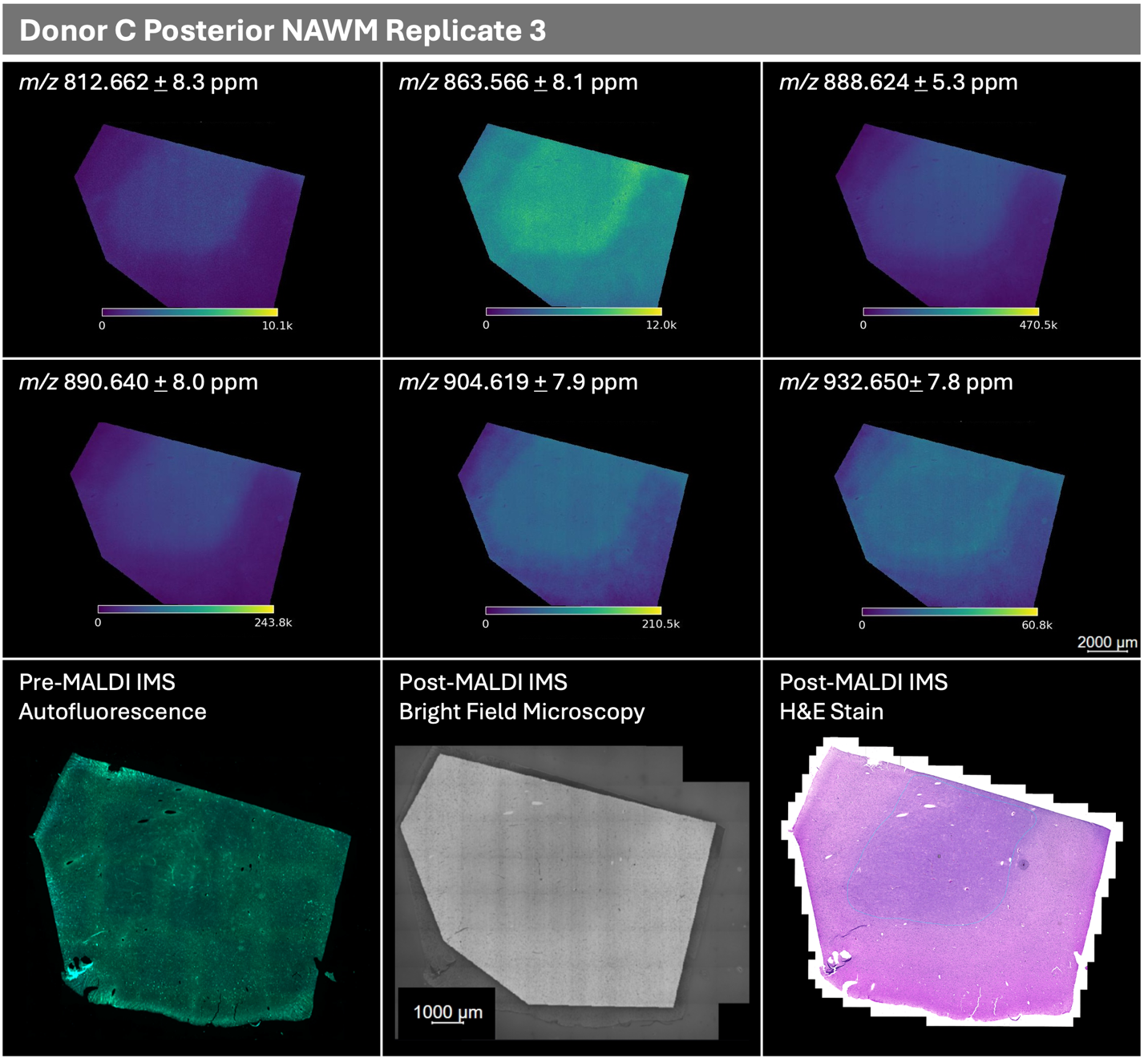

*(Figure S3 continued on next page)*

**Figure S3 (continued)**
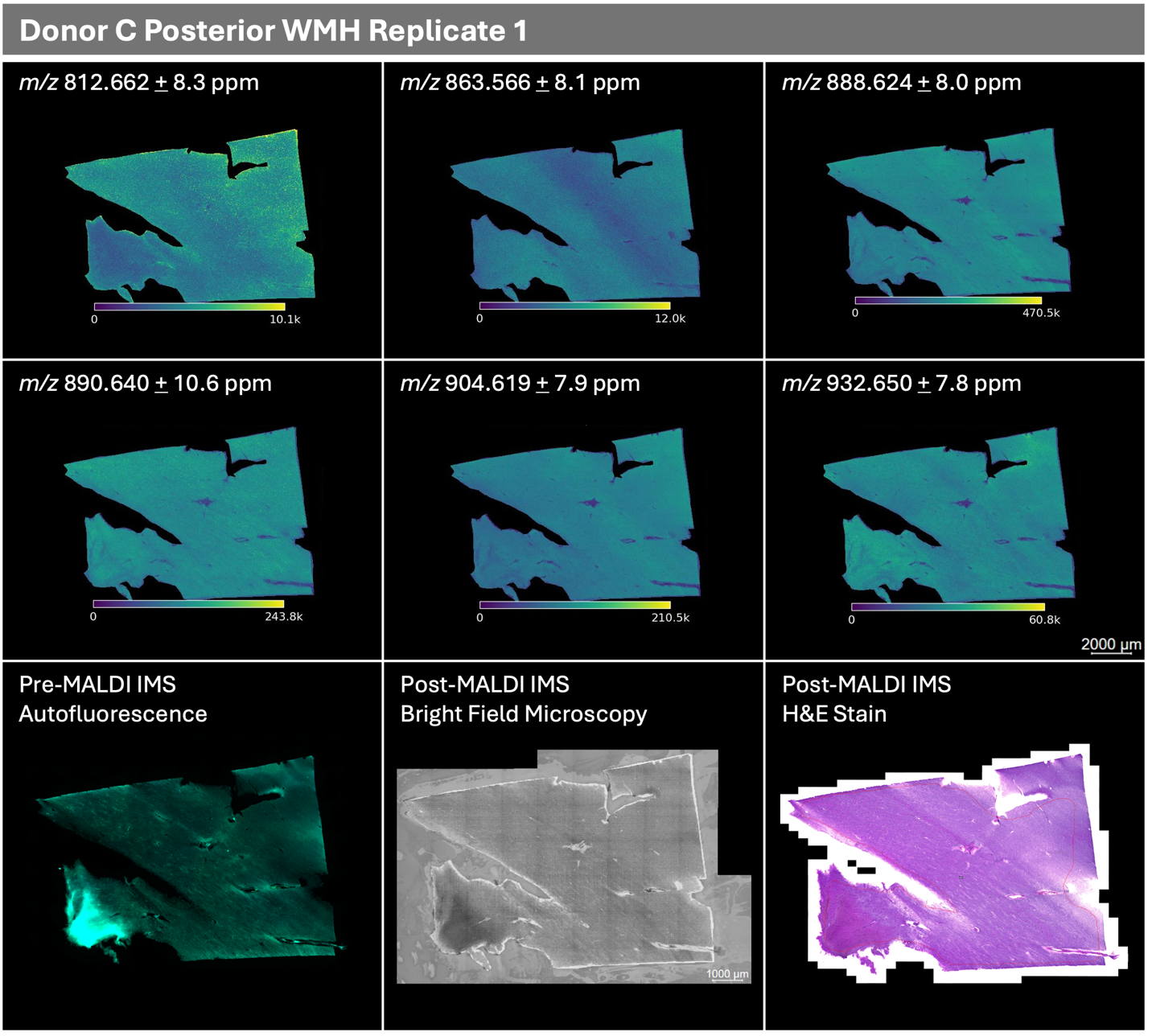

*(Figure S3 continued on next page)*

**Figure S3 (continued)**

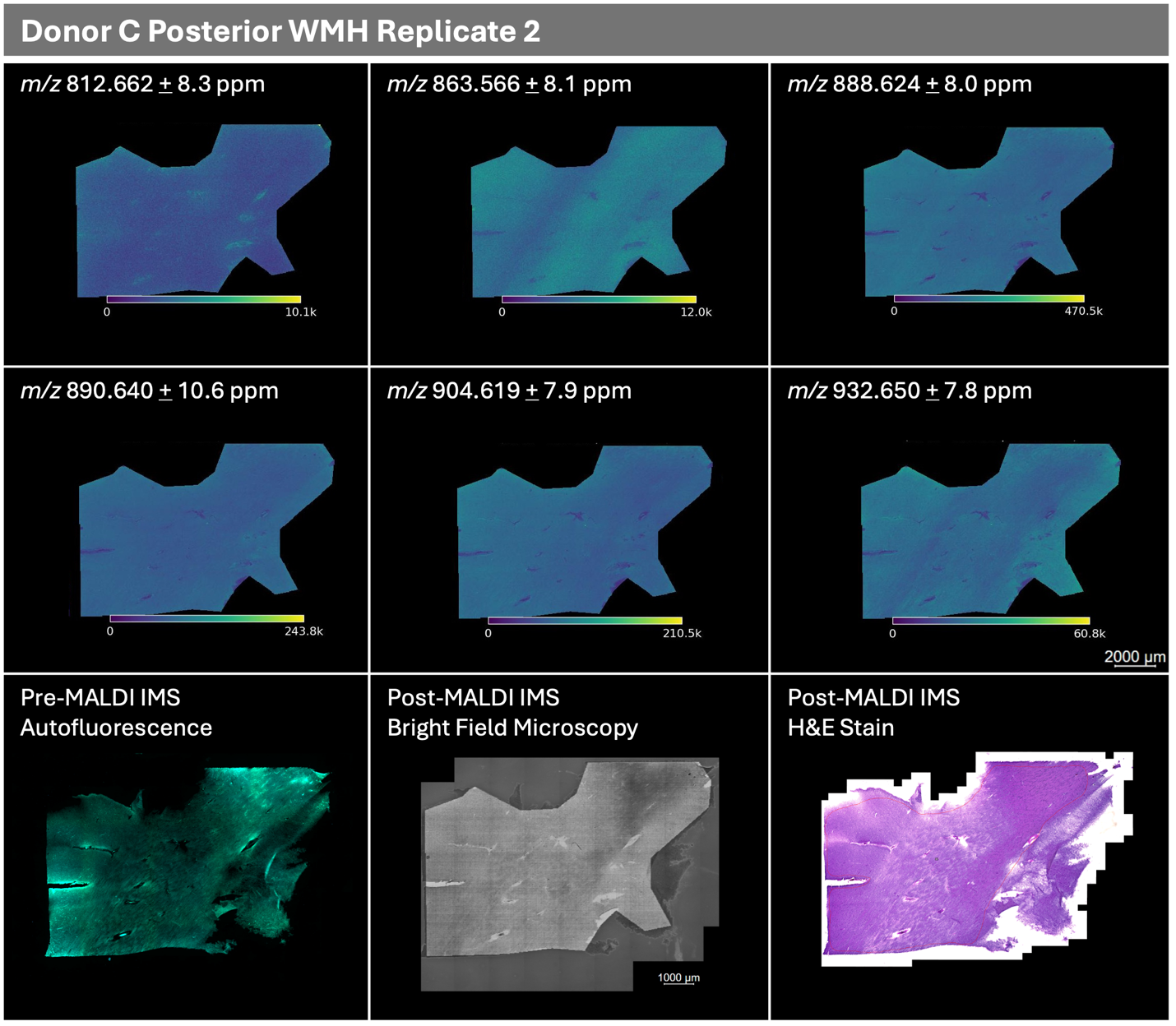

*(Figure S3 continued on next page)*

**Figure S3 (continued)**
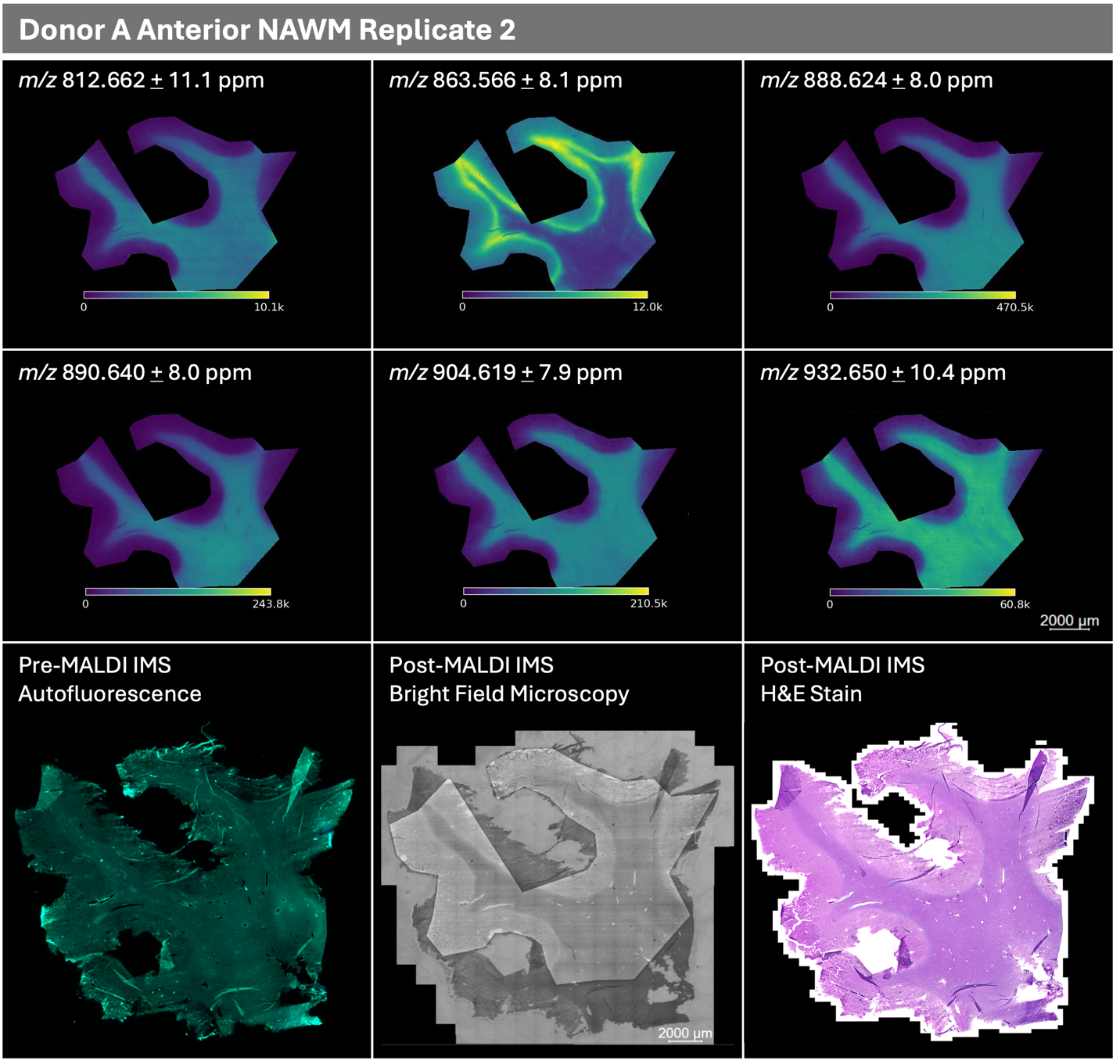

*(Figure S3 continued on next page)*

**Figure S3 (continued)**

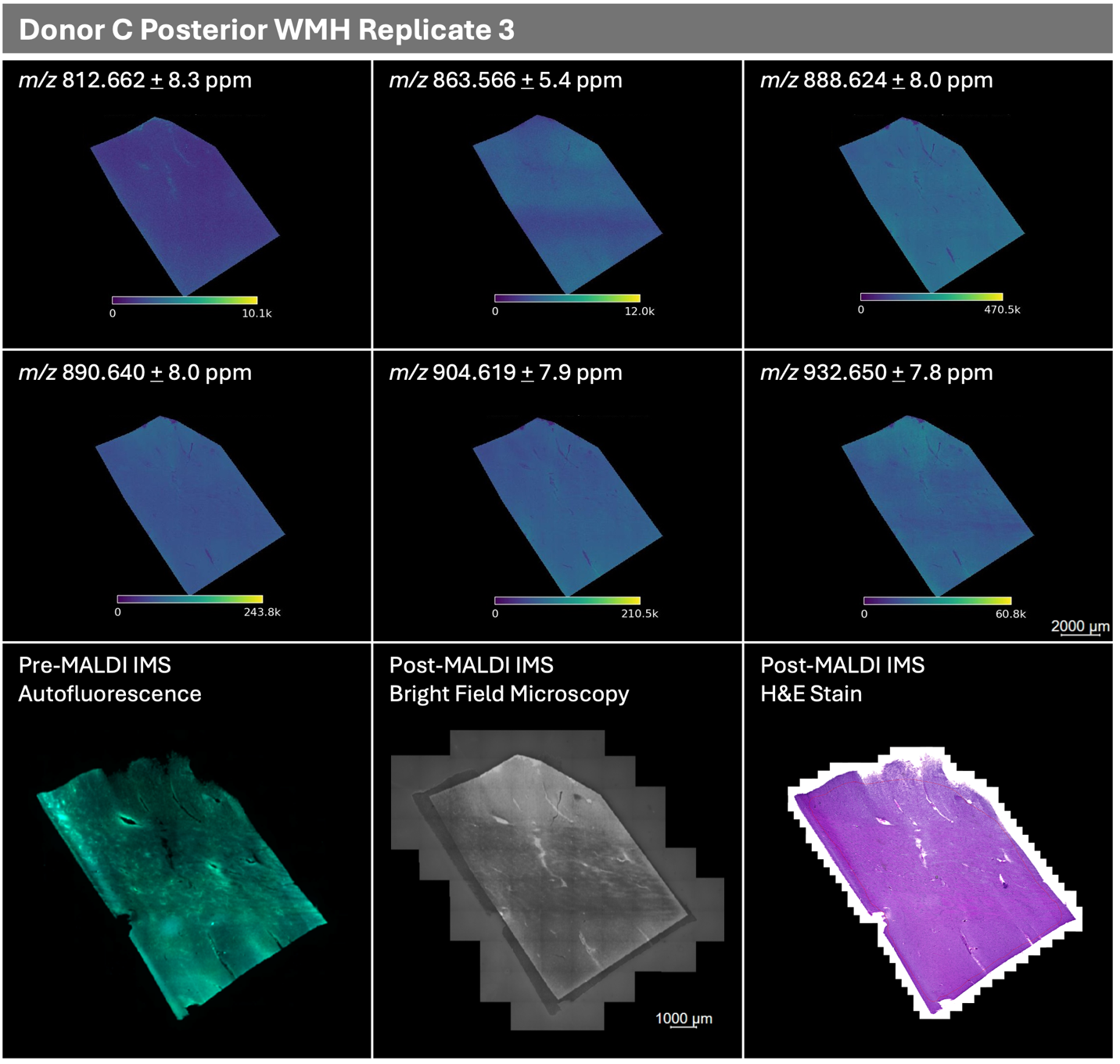

***Figure S3:*** *MALDI IMS ion images, m/z 812.662, m/z 863.566, m/z 888.624, m/z 890.640, m/z 904.619, m/z 932.650. Pre-MALDI IMS autofluorescence microscopy images (overlay of excitation wavelengths 353 nm and 488 nm). Post-MALDI IMS bright field microscopy, which shows MALDI laser ablation pixel marks for image registration. Post-MALDI IMS and post-matrix removal H&E stain, including annotation (thin red line) of the white matter boundaries. All tissues and replicates are shown.*

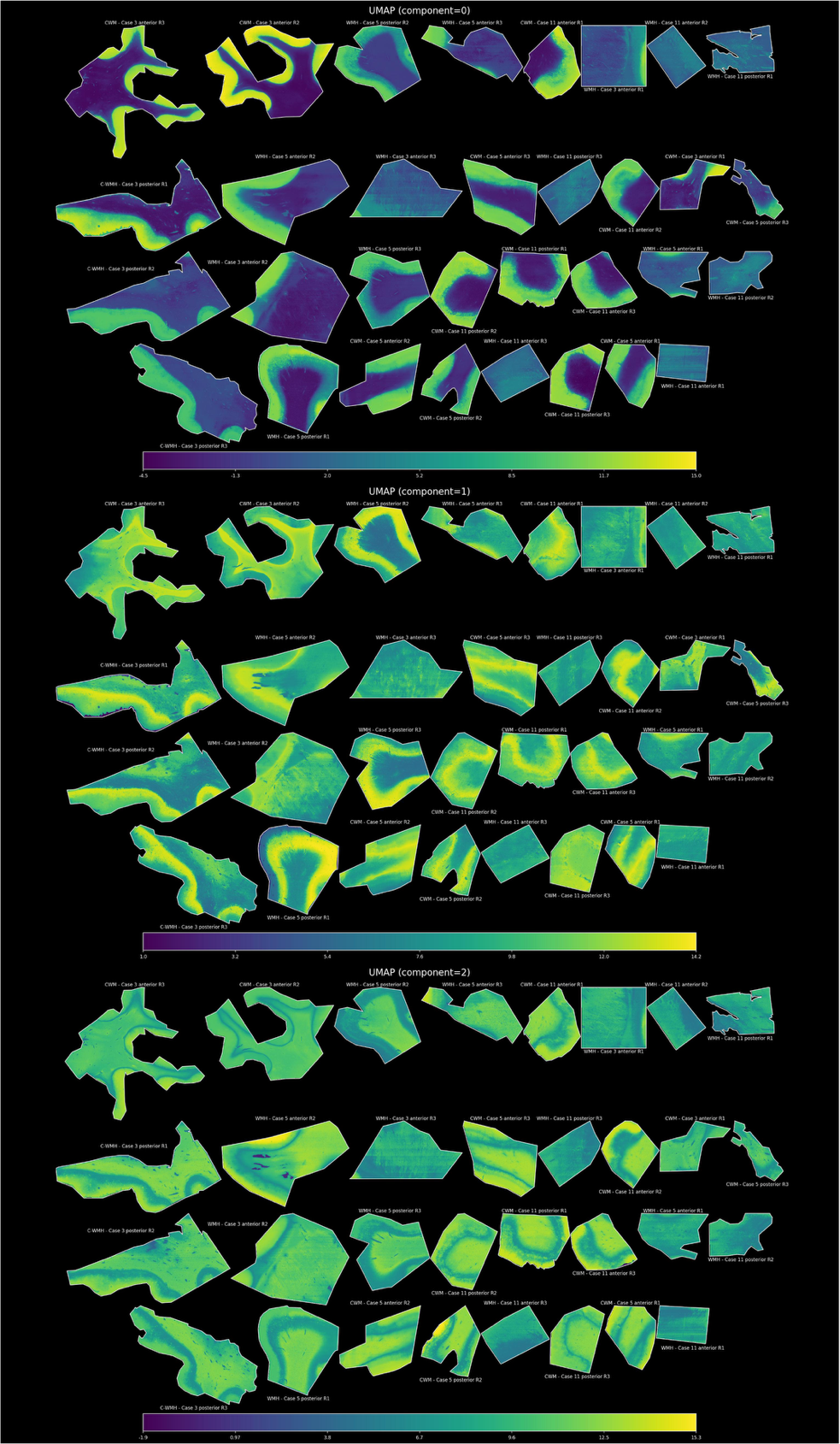

***Figure S4.*** *UMAP Components (0-2, top to bottom), shown across all tissue samples and replicates. *Note:* ***Cases 3, 5, and 11 correspond to Donors A, B, and C, respectively****, matching the clinical profiles in Figure S1. WMH refers to white matter hyperintensity, CWM refers to control white matter, referenced in the main text of this paper as NAWM. Anterior / Posterior refers to the brain sample region, and “R#” refers to the replicate number (1-3).*

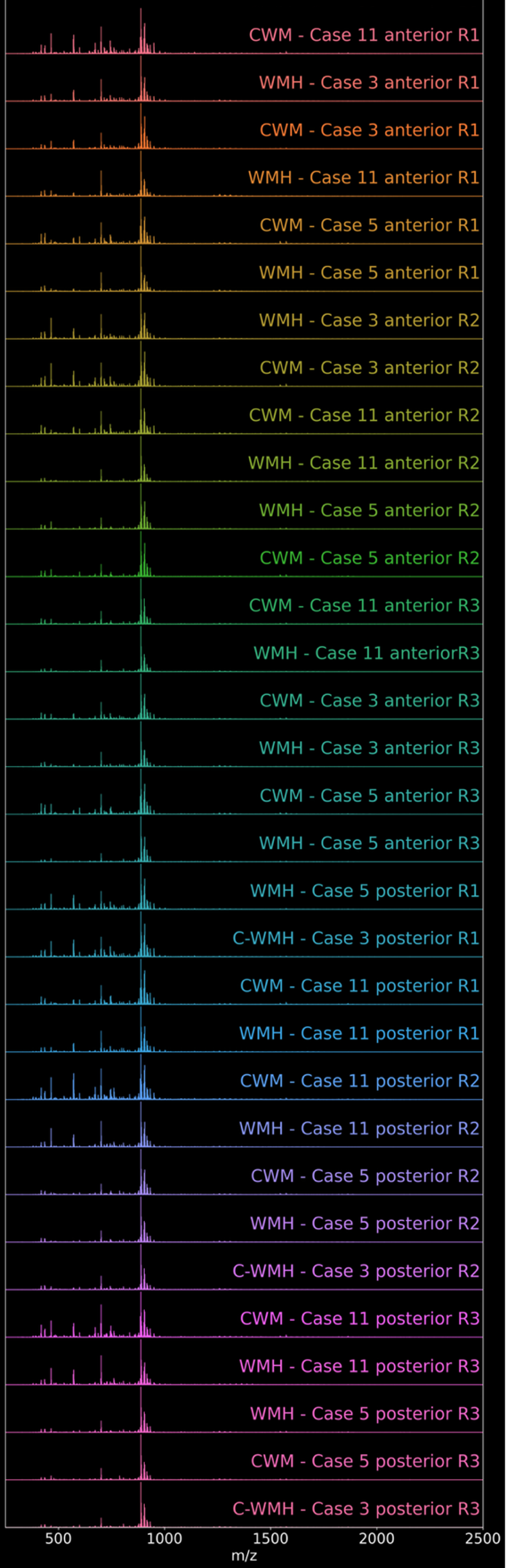

***Figure S5.*** *Waterfall plot of all average mass spectra. WMH refers to white matter hyperintensity, *Note:* ***Cases 3, 5, and 11 correspond to Donors A, B, and C, respectively****, matching the clinical profiles in Figure S1. WMH refers to white matter hyperintensity, CWM refers to control white matter, referenced in the main text of this paper as NAWM. Anterior / Posterior refers to the brain sample region, and “R#” refers to the replicate number (1-3).*

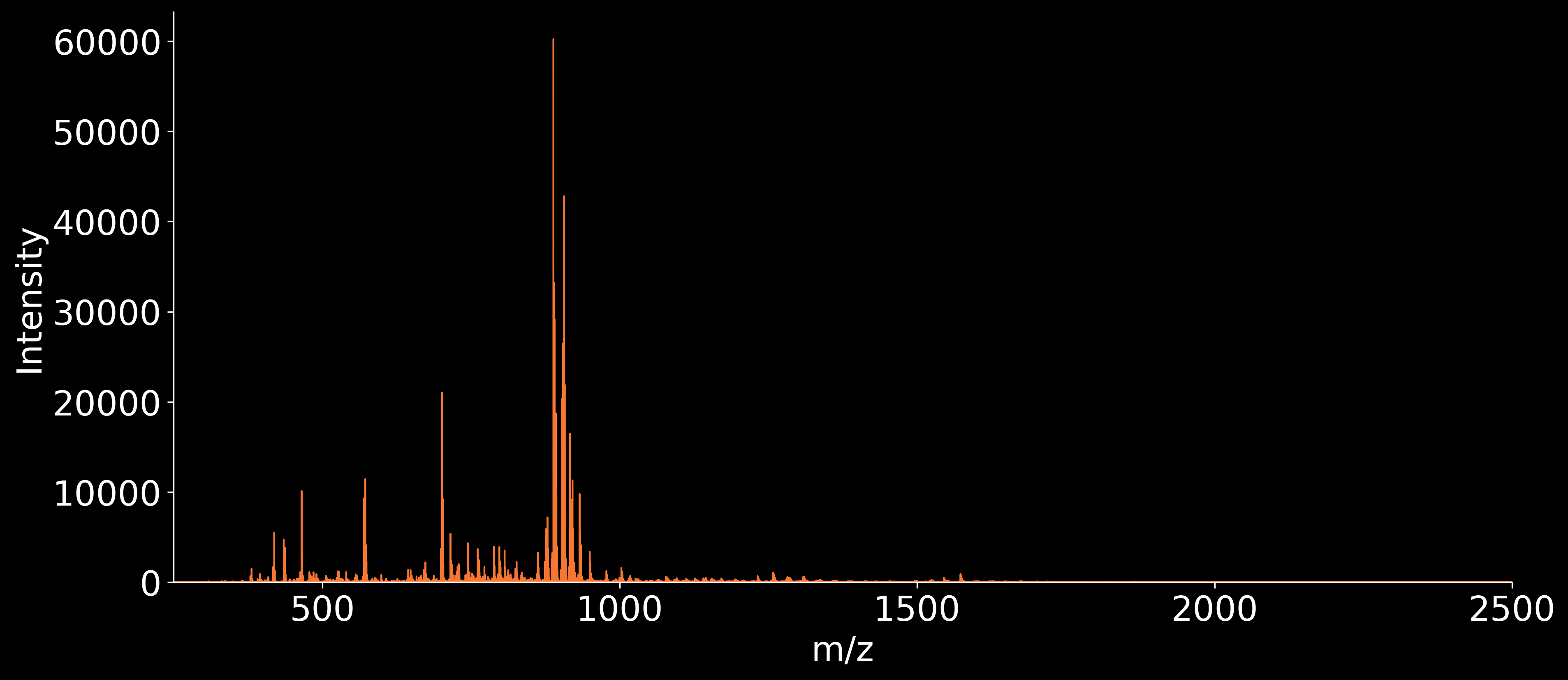

NAWM - Donor A anterior REPLICATE 1

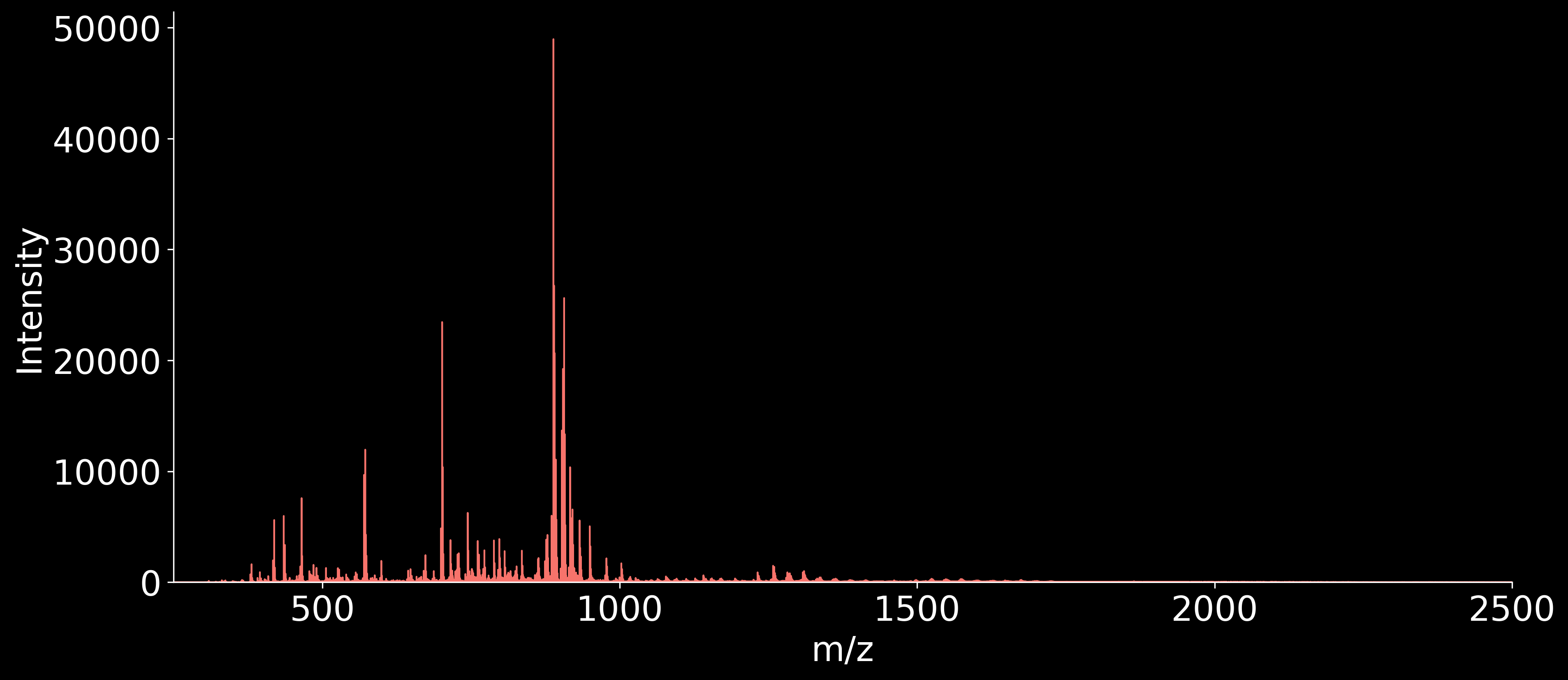

WMH - Donor A anterior REPLICATE 1

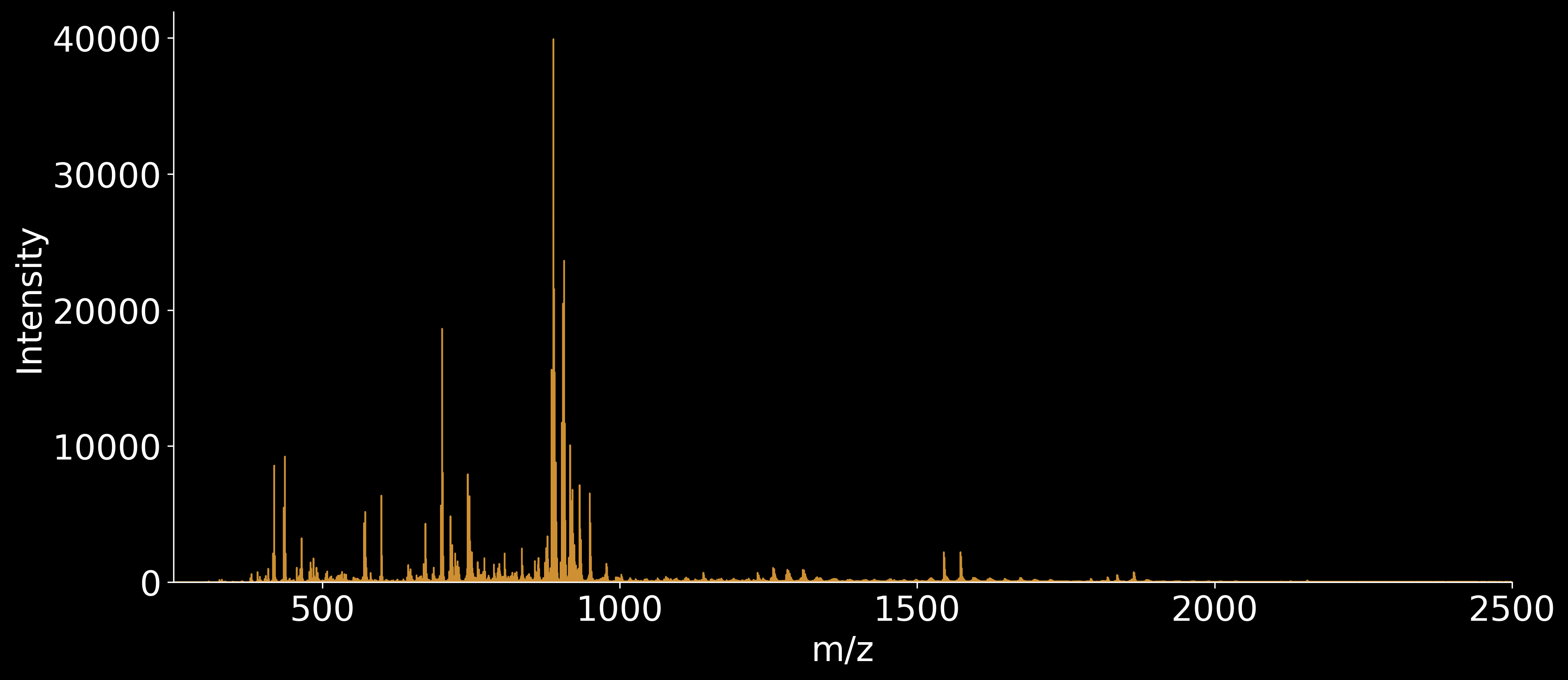

NAWM - Donor B anterior REPLICATE 1

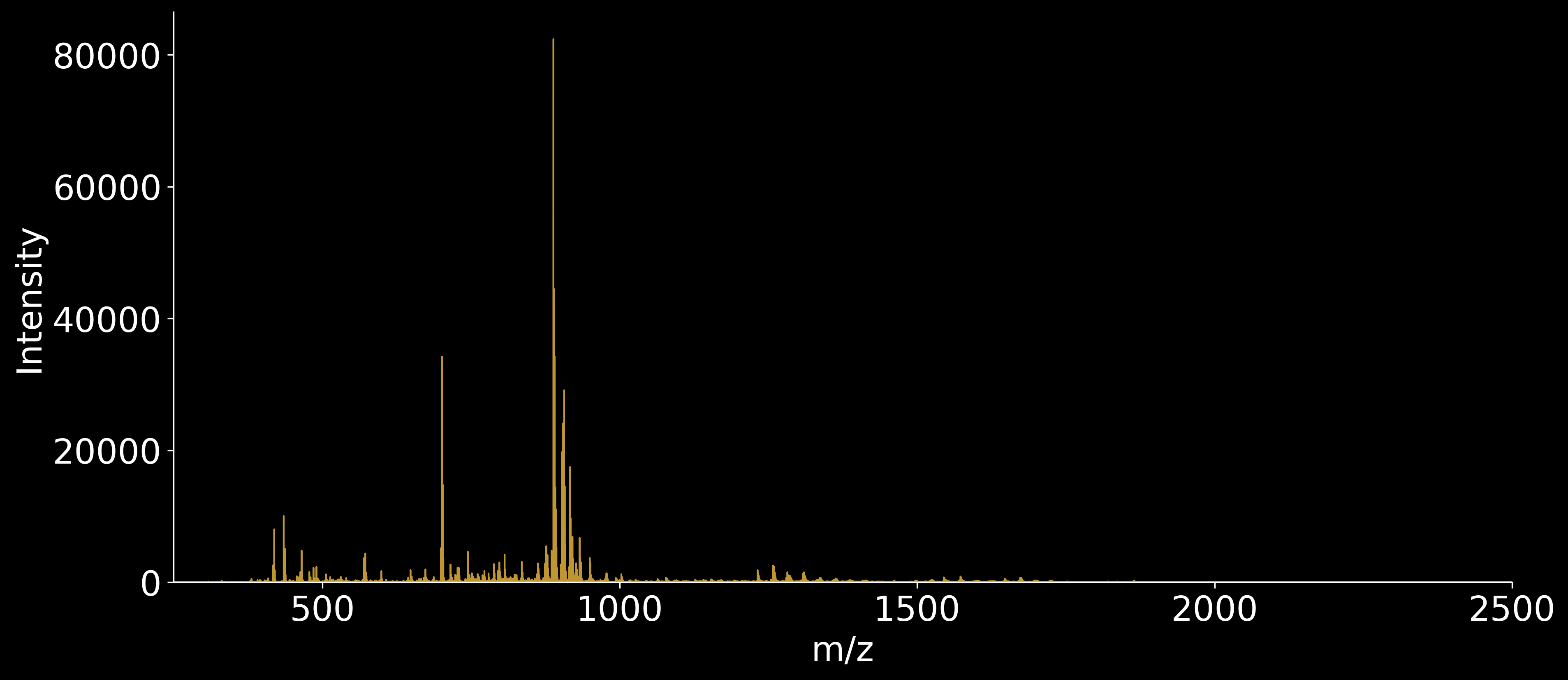

WMH - Donor B anterior REPLICATE 1

*(Figure S6 continued on next page)*

**Figure S6 (continued)**

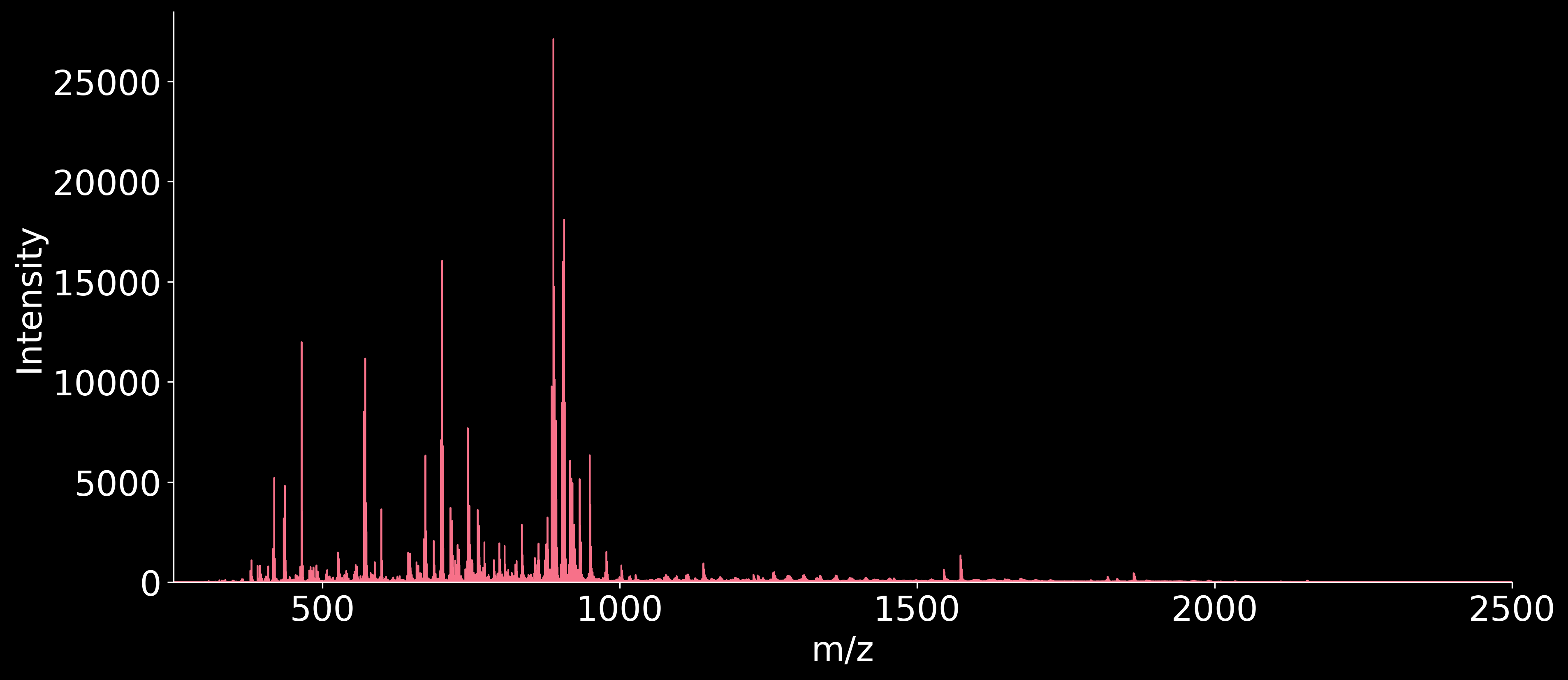

NAWM - Donor C anterior REPLICATE 1

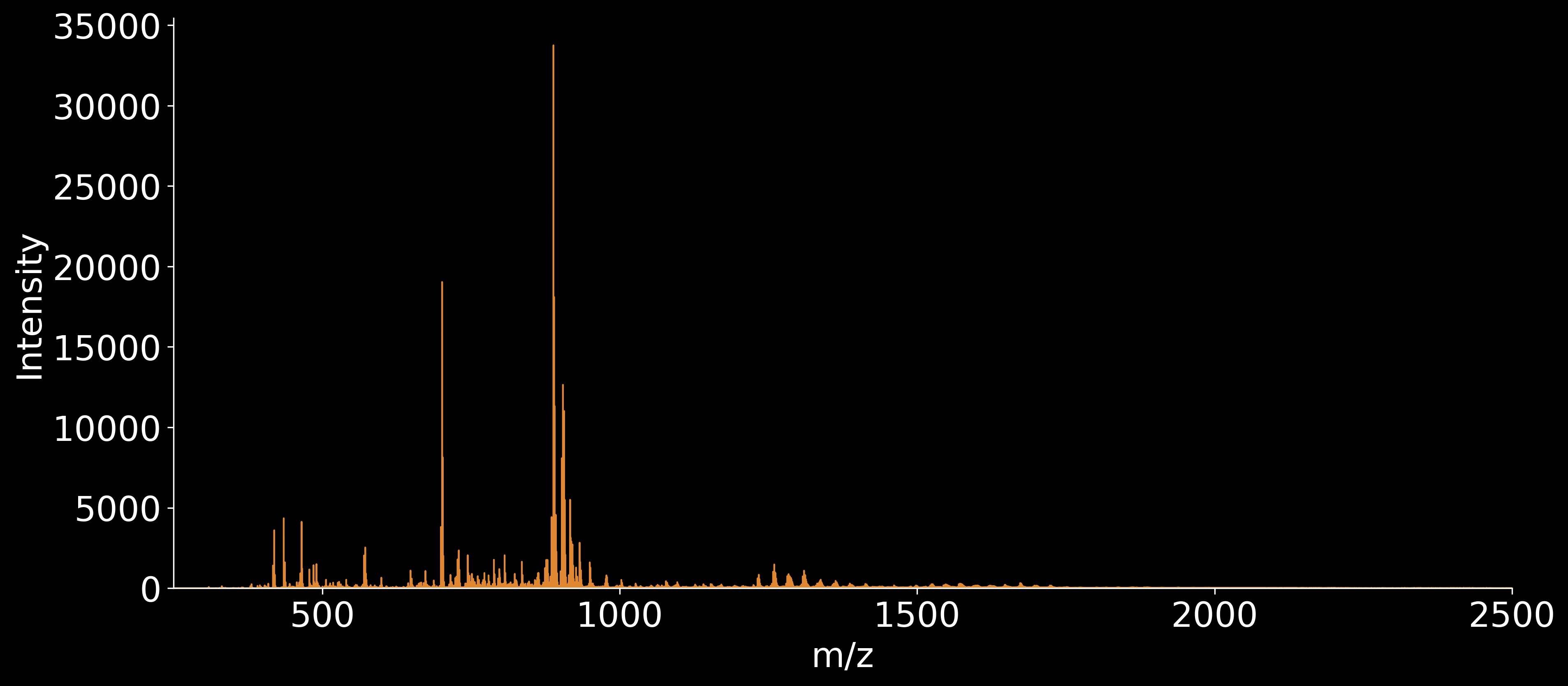

WMH - Donor C anterior REPLICATE 1

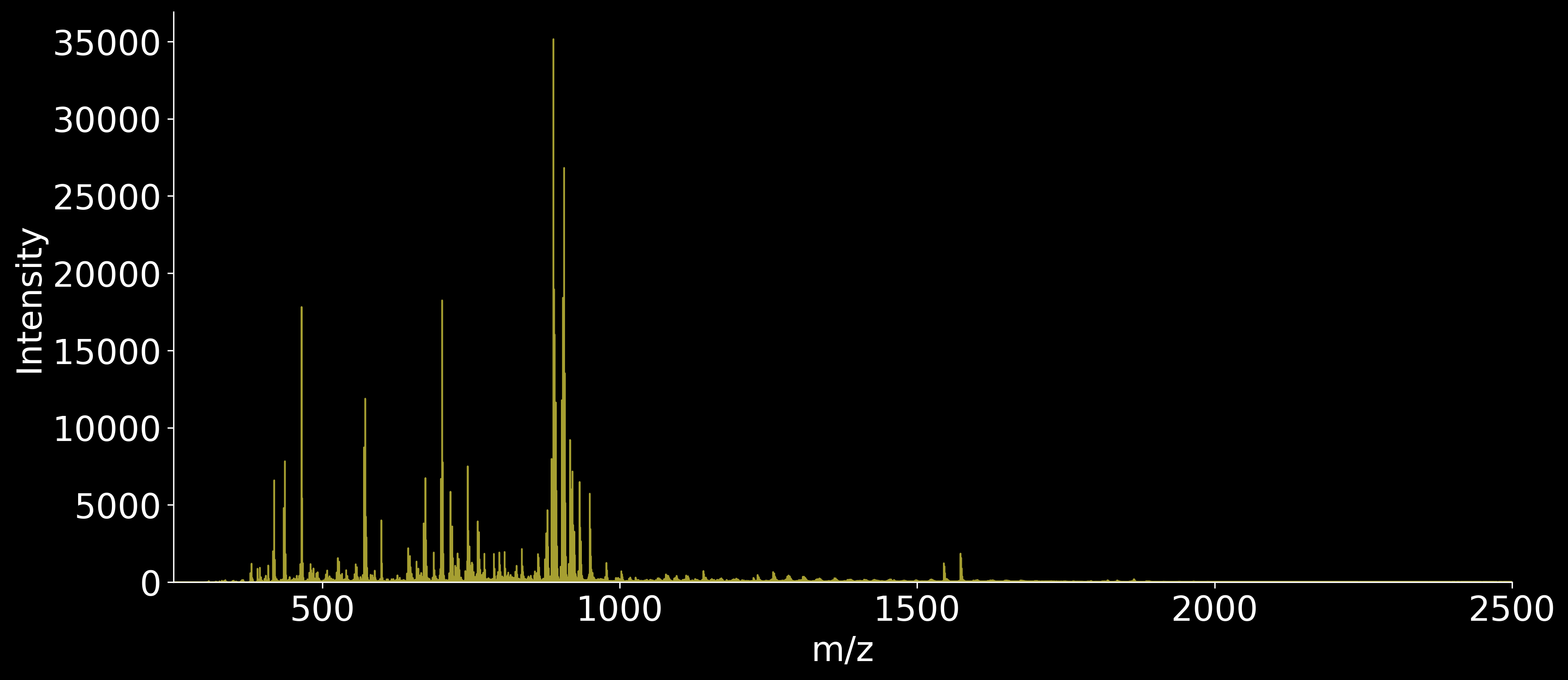

NAWM - Donor A anterior REPLICATE 2

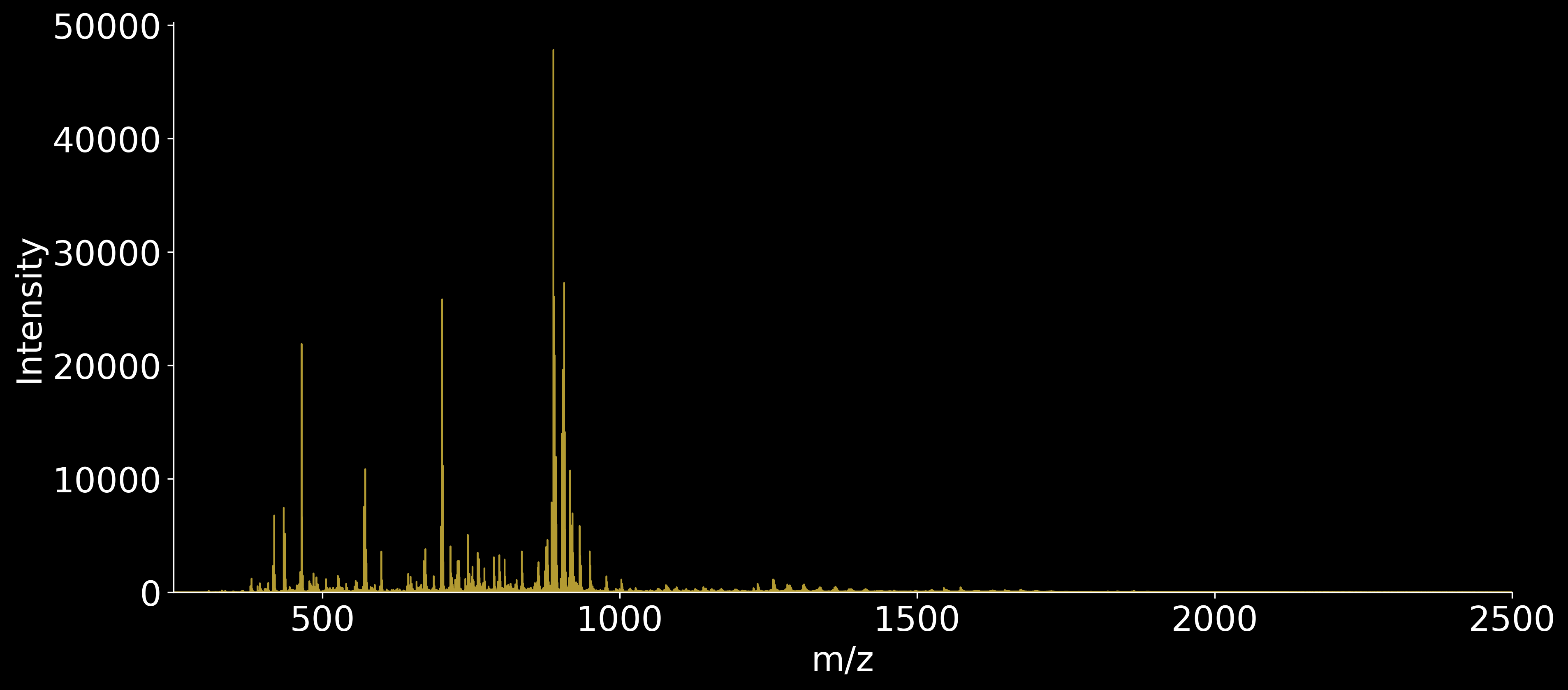

WMH - Donor A anterior REPLICATE 2

*(Figure S6 continued on next page)*

**Figure S6 (continued)**

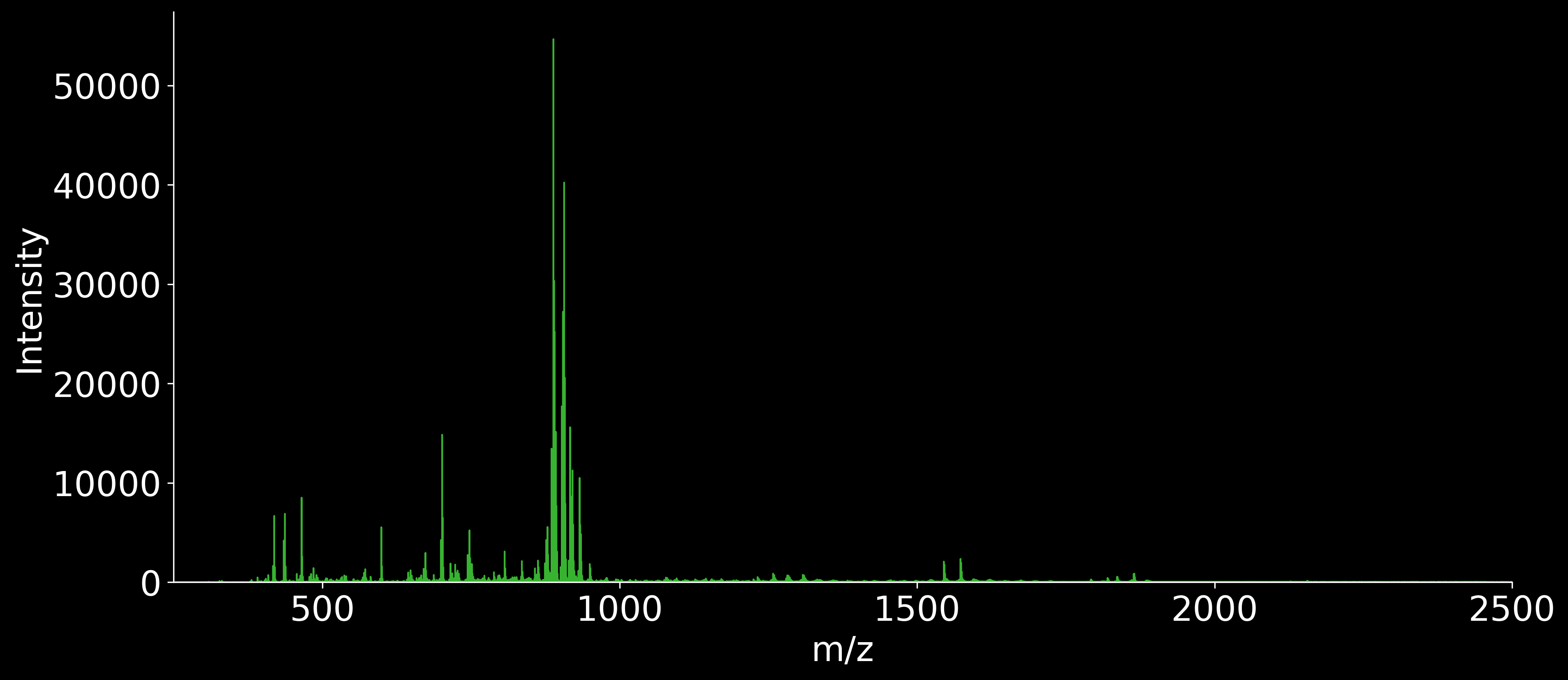

NAWM - Donor B anterior REPLICATE 2

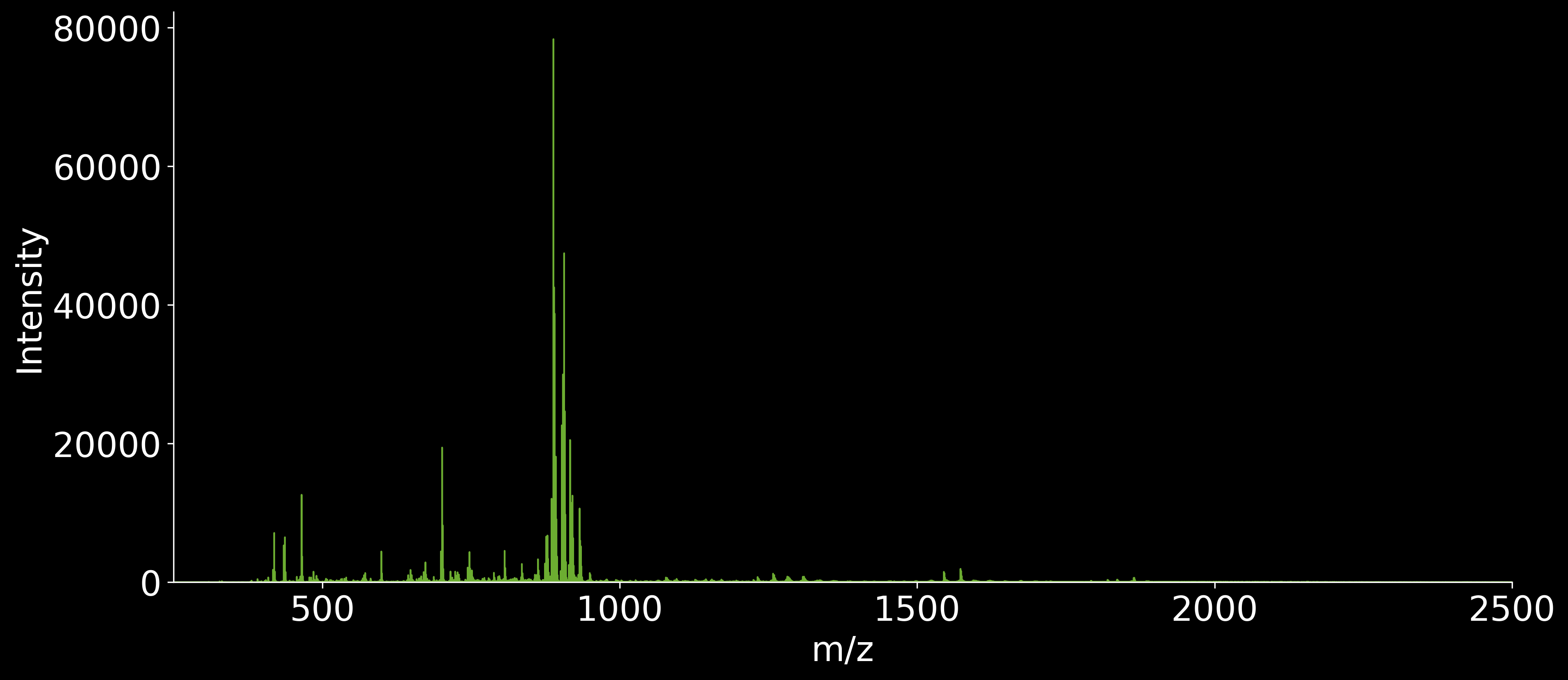

WMH - Donor B anterior REPLICATE 2

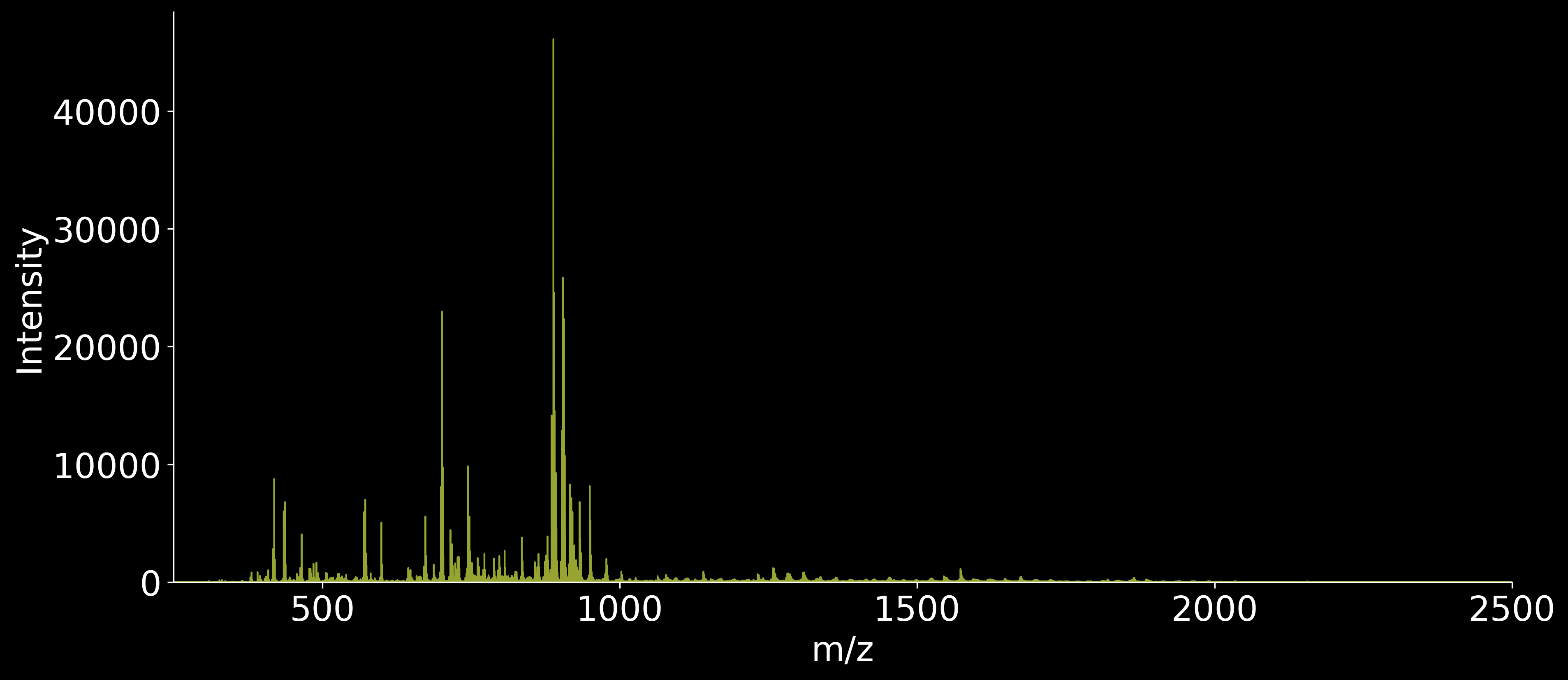

NAWM - Donor C anterior REPLICATE 2

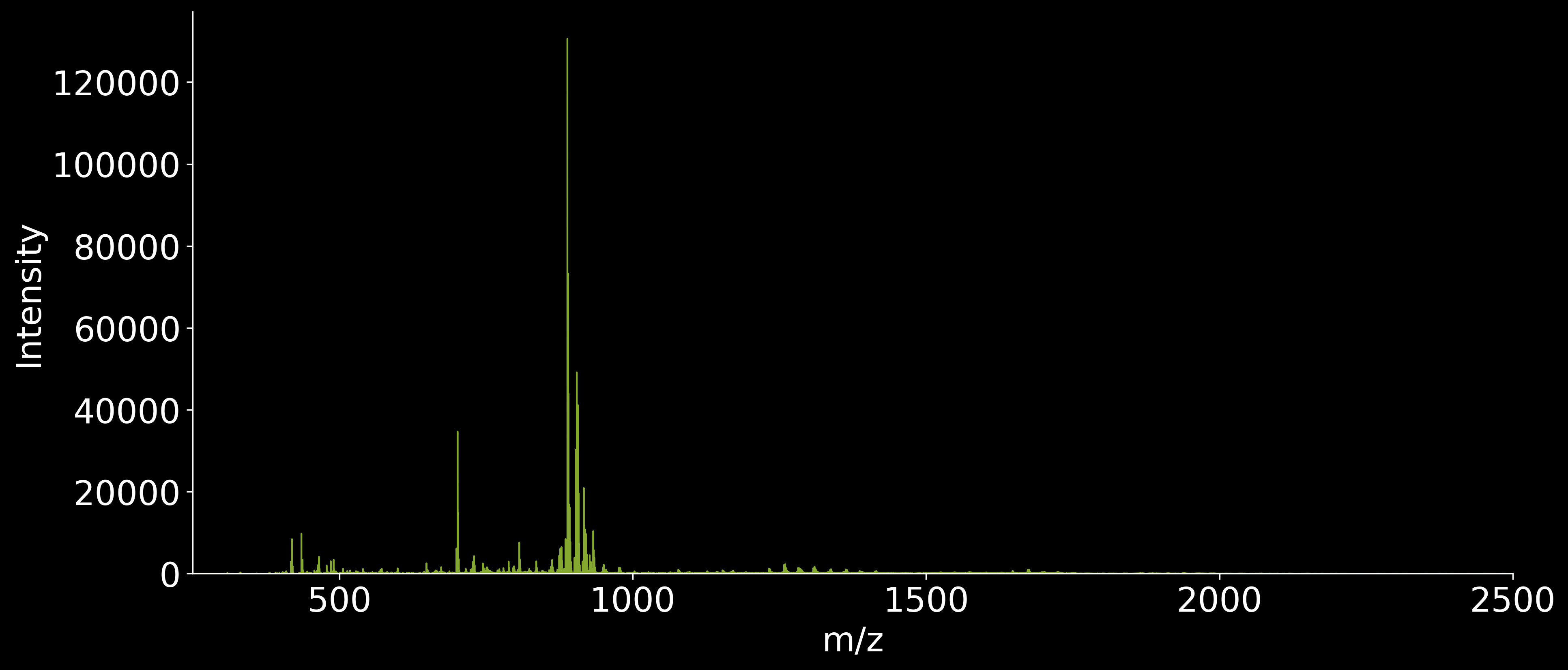

WMH - Donor C anterior REPLICATE 2

*(Figure S6 continued on next page)*

**Figure S6 (continued)**

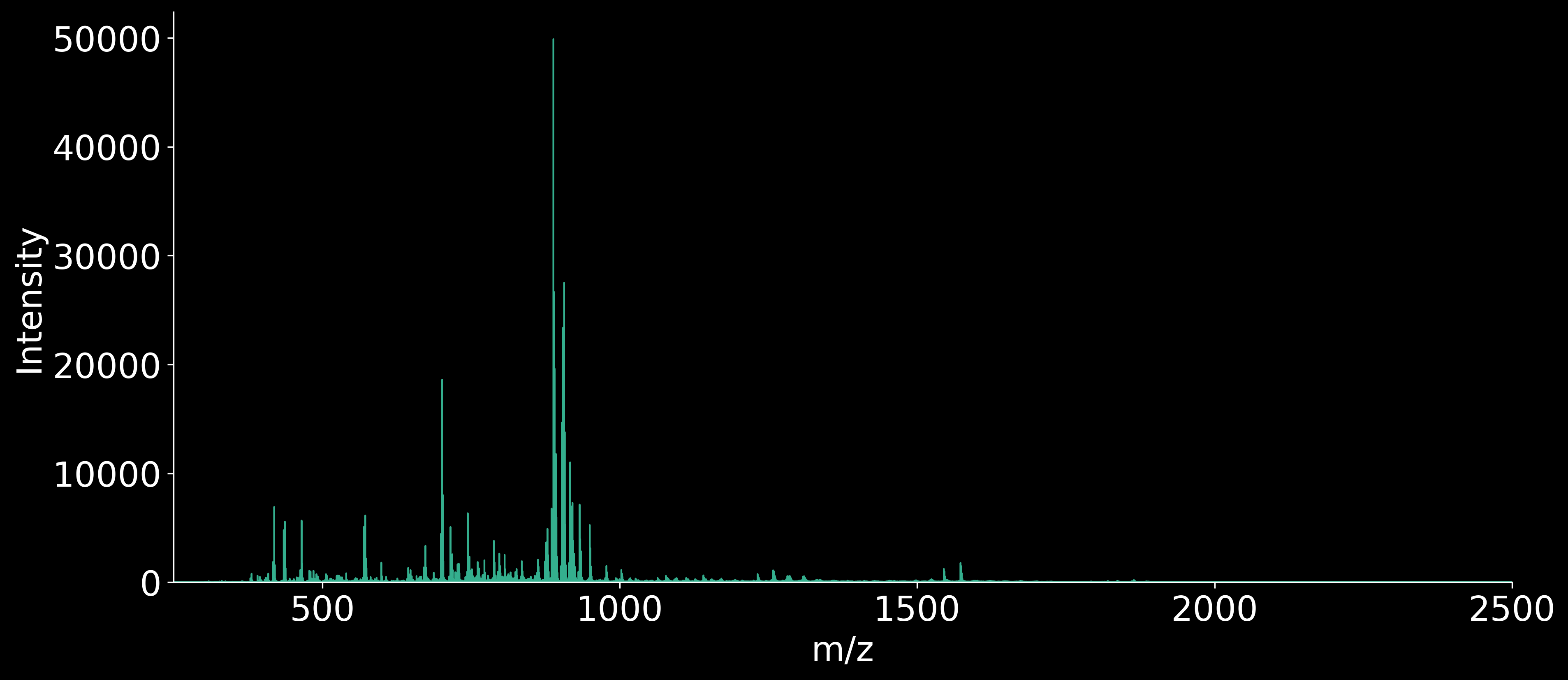

NAWM - Donor A anterior REPLICATE 3

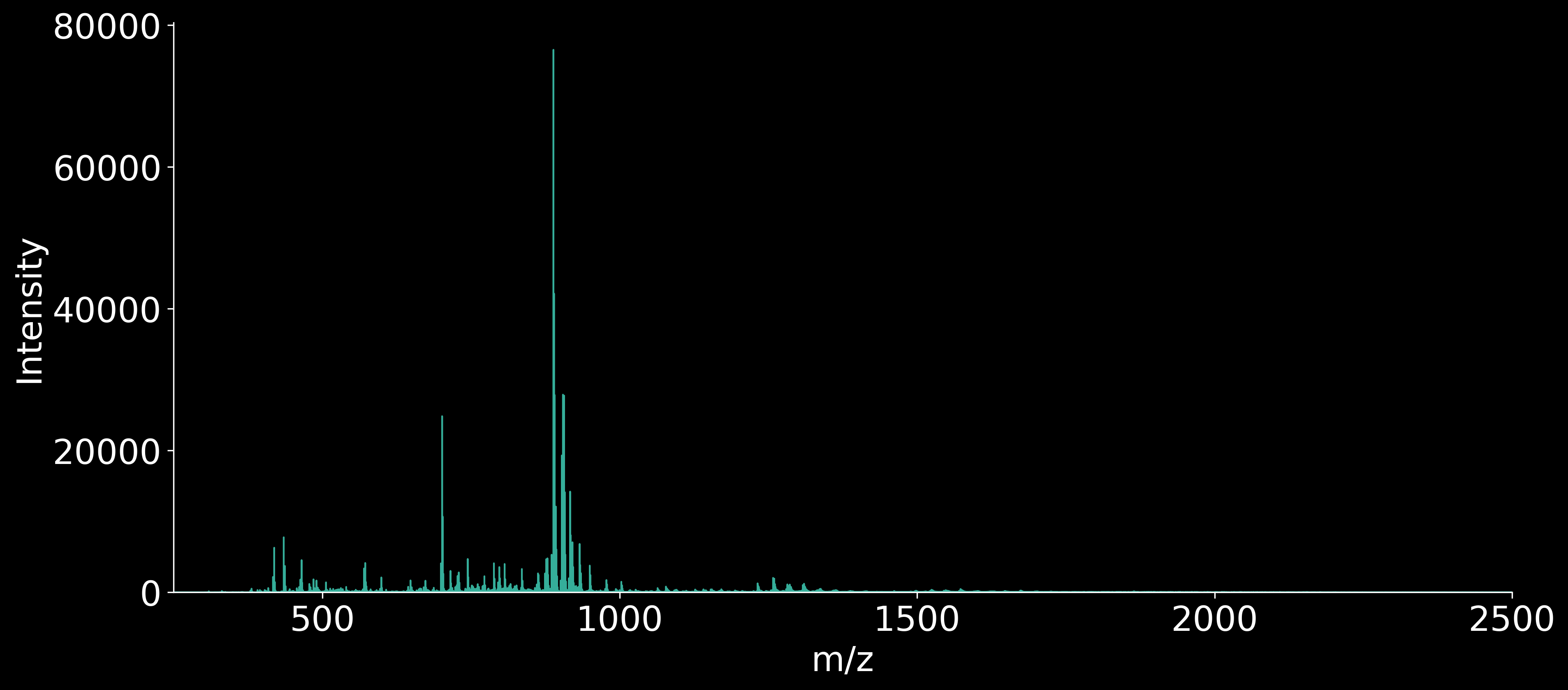

WMH - Donor A anterior REPLICATE 3

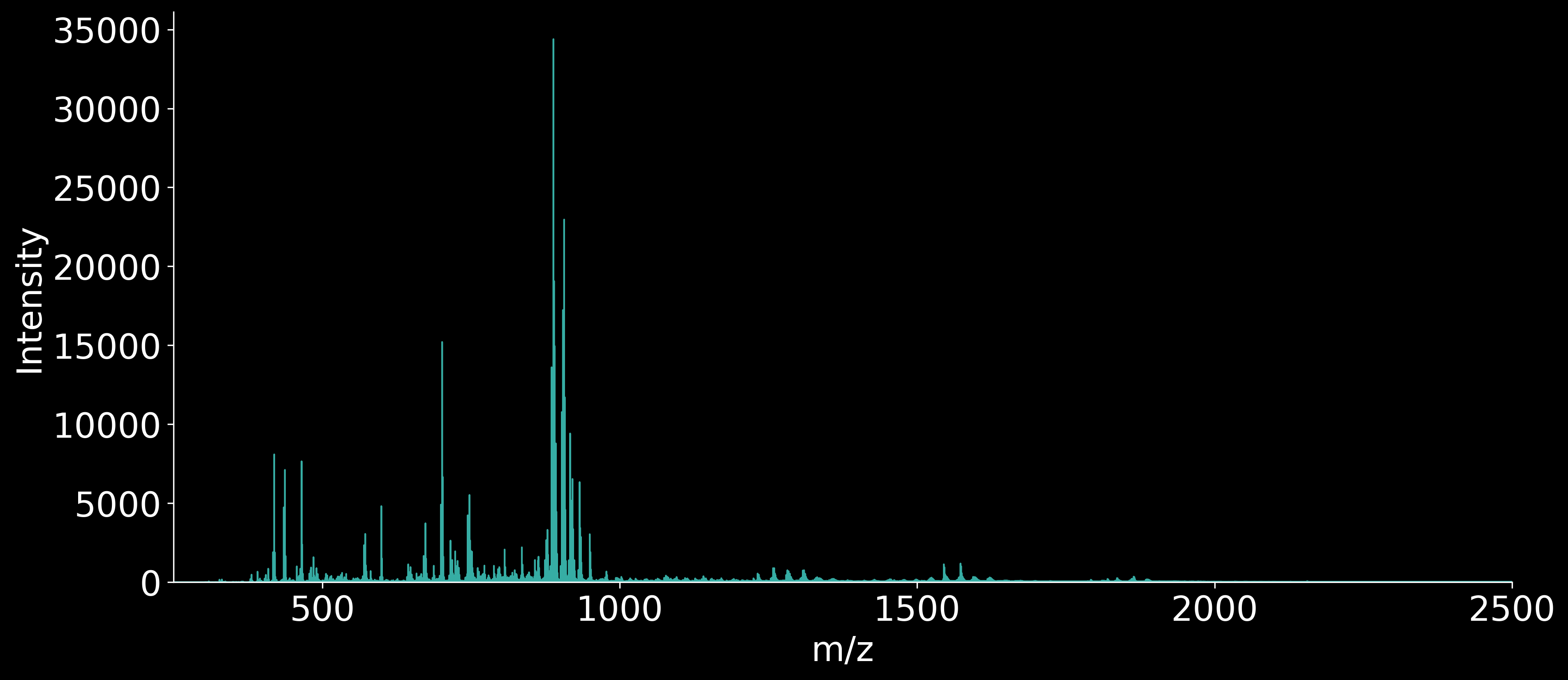

NAWM - Donor B anterior REPLICATE 3

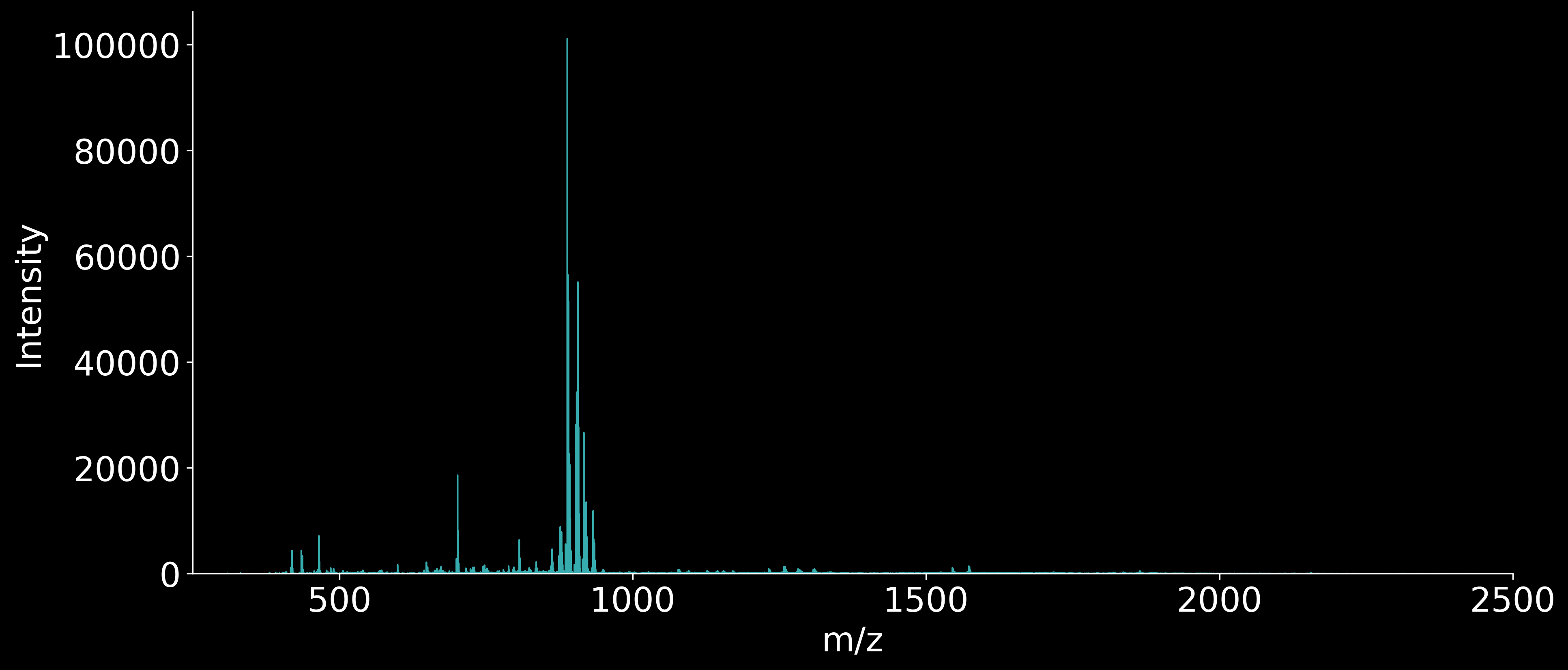

WMH - Donor B anterior REPLICATE 3

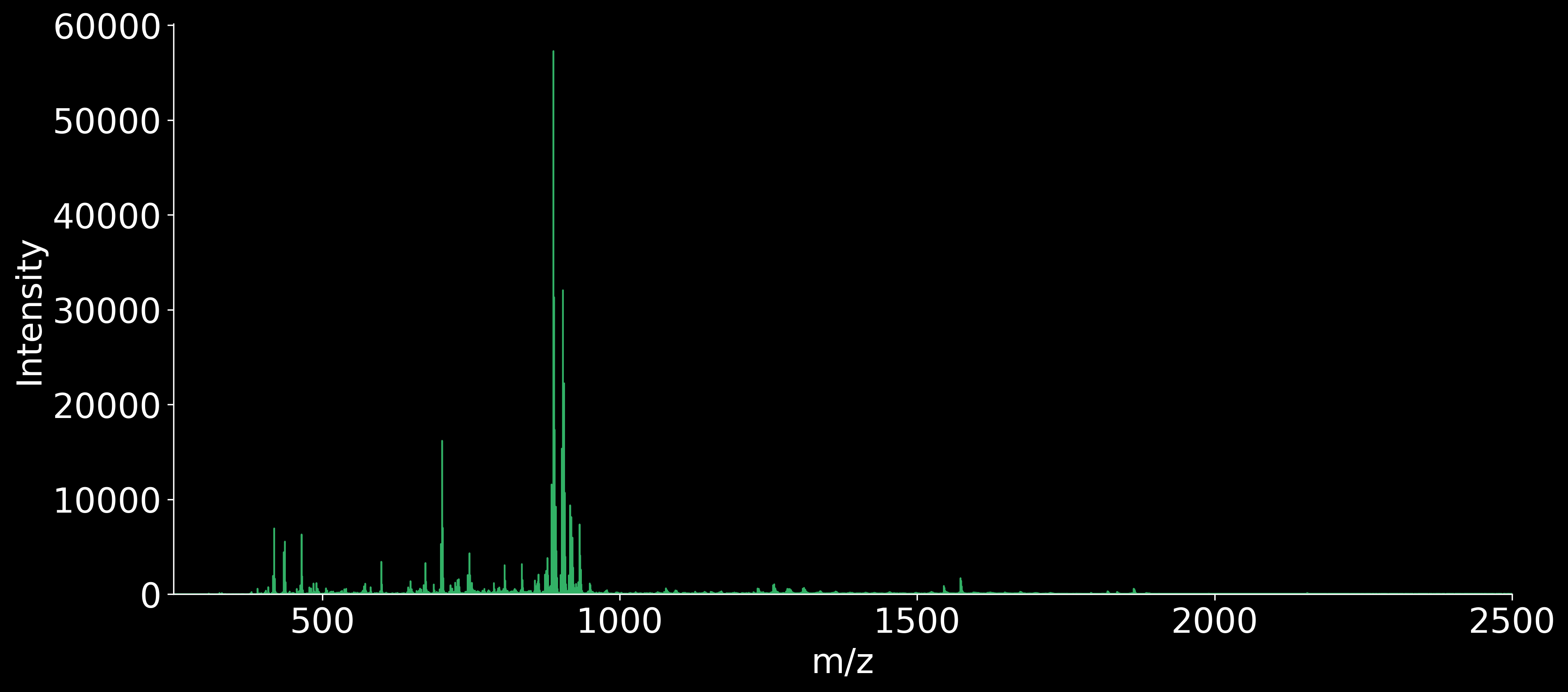

NAWM - Donor C anterior REPLICATE 3

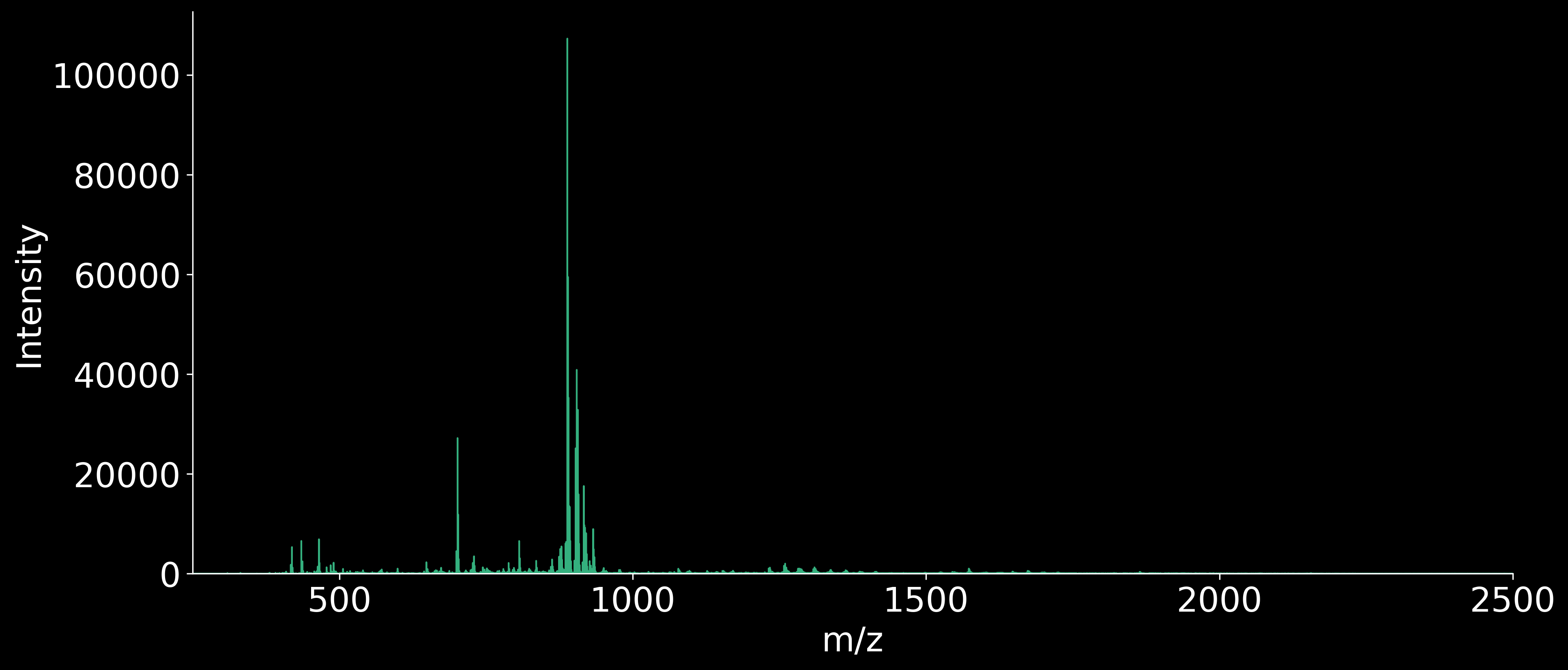

WMH - Donor C anterior REPLICATE 3

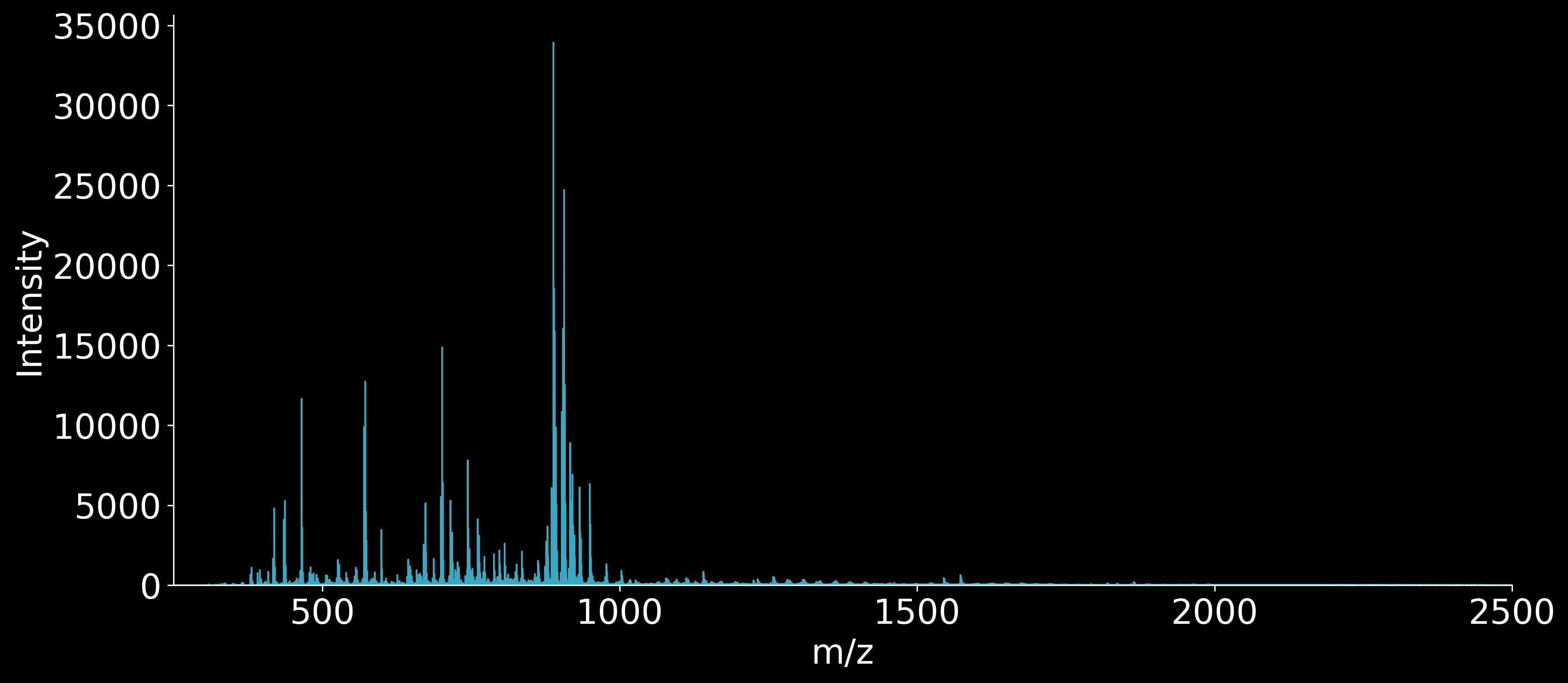

NAWM & WMH - Donor A posterior REPLICATE 1

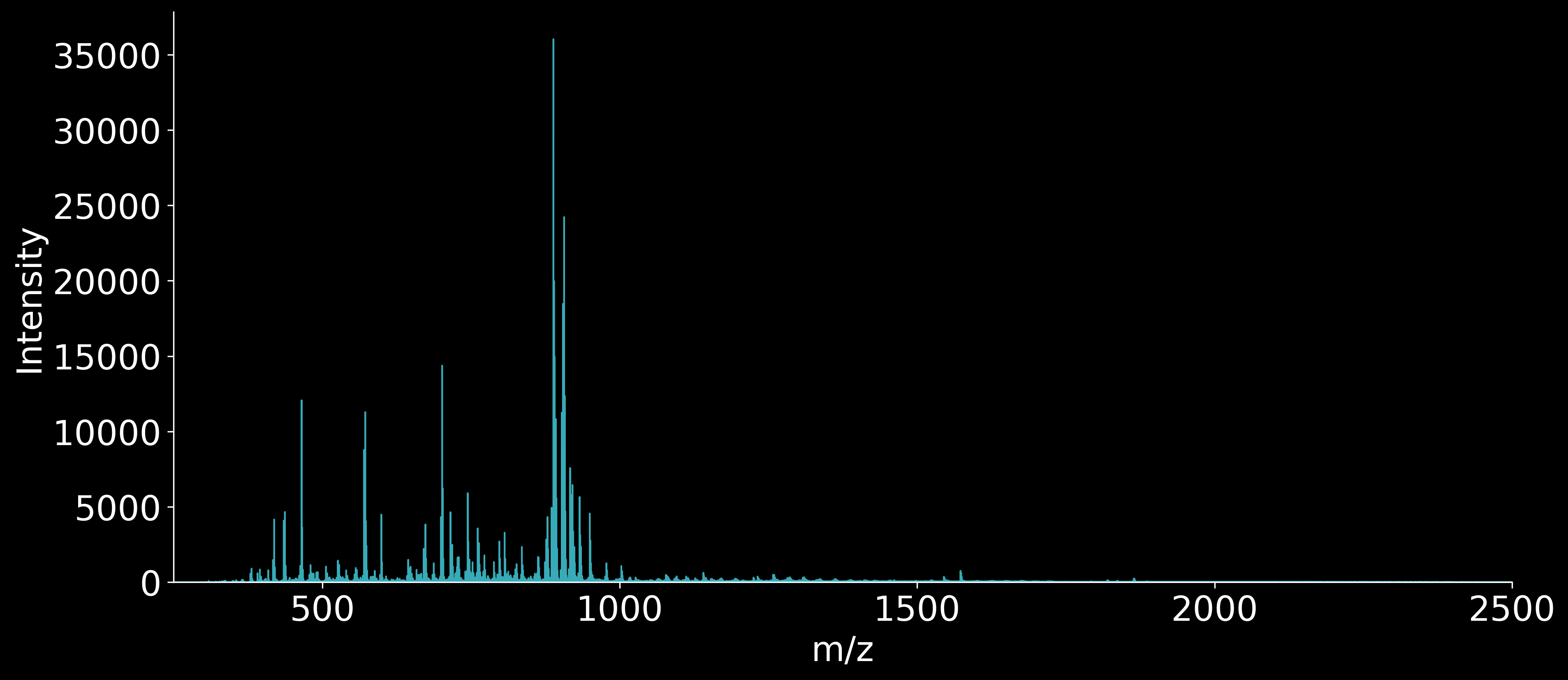

WMH - Donor B posterior REPLICATE 1

*(Figure S6 continued on next page)*

**Figure S6 (continued)**

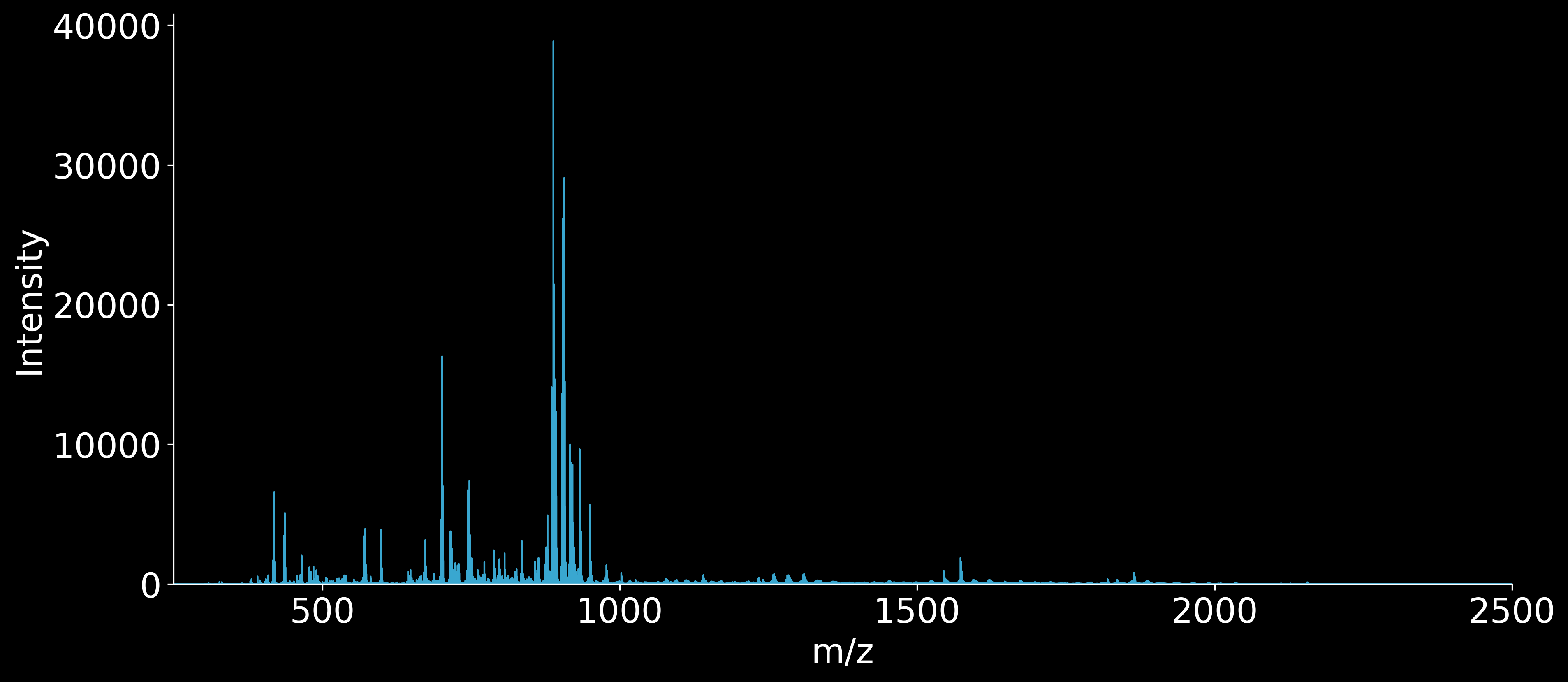

NAWM - Donor C posterior REPLICATE 1

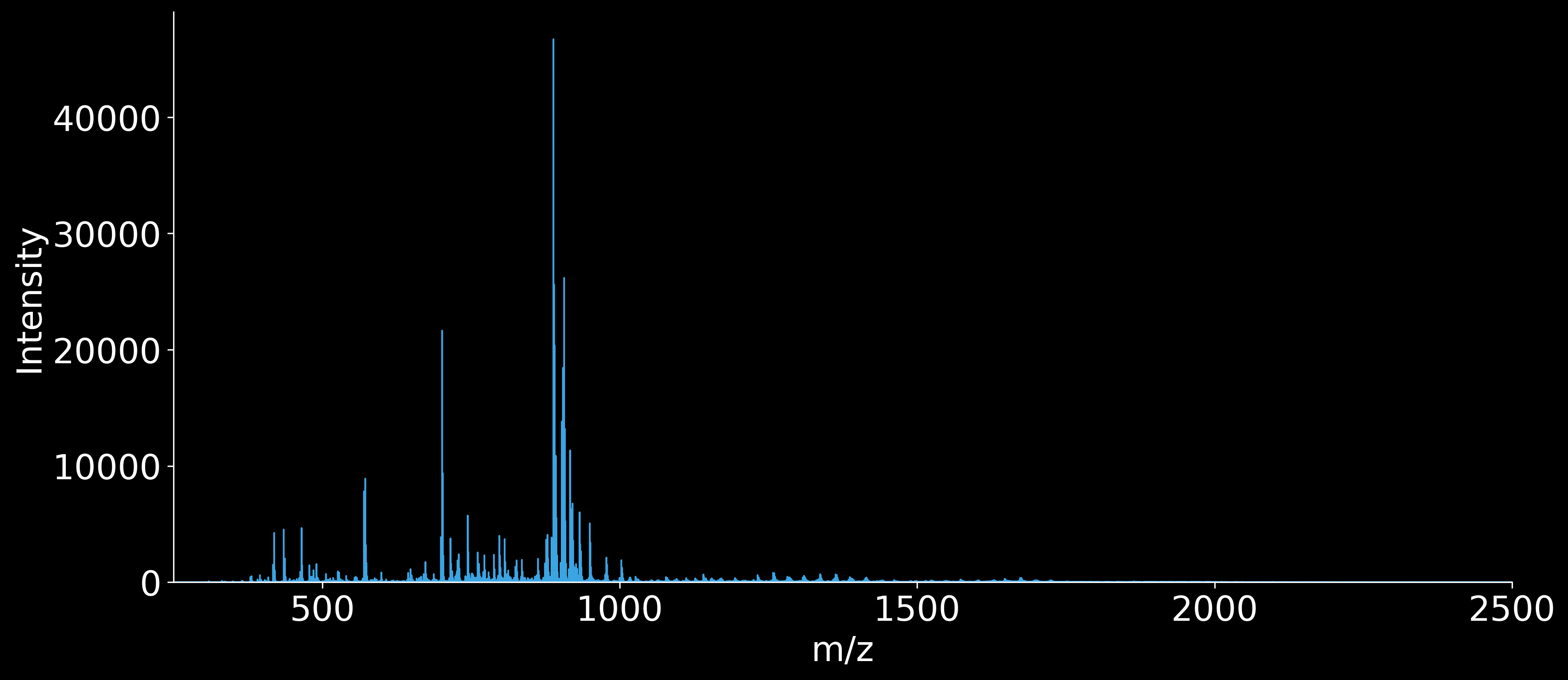

WMH - Donor C posterior REPLICATE 1

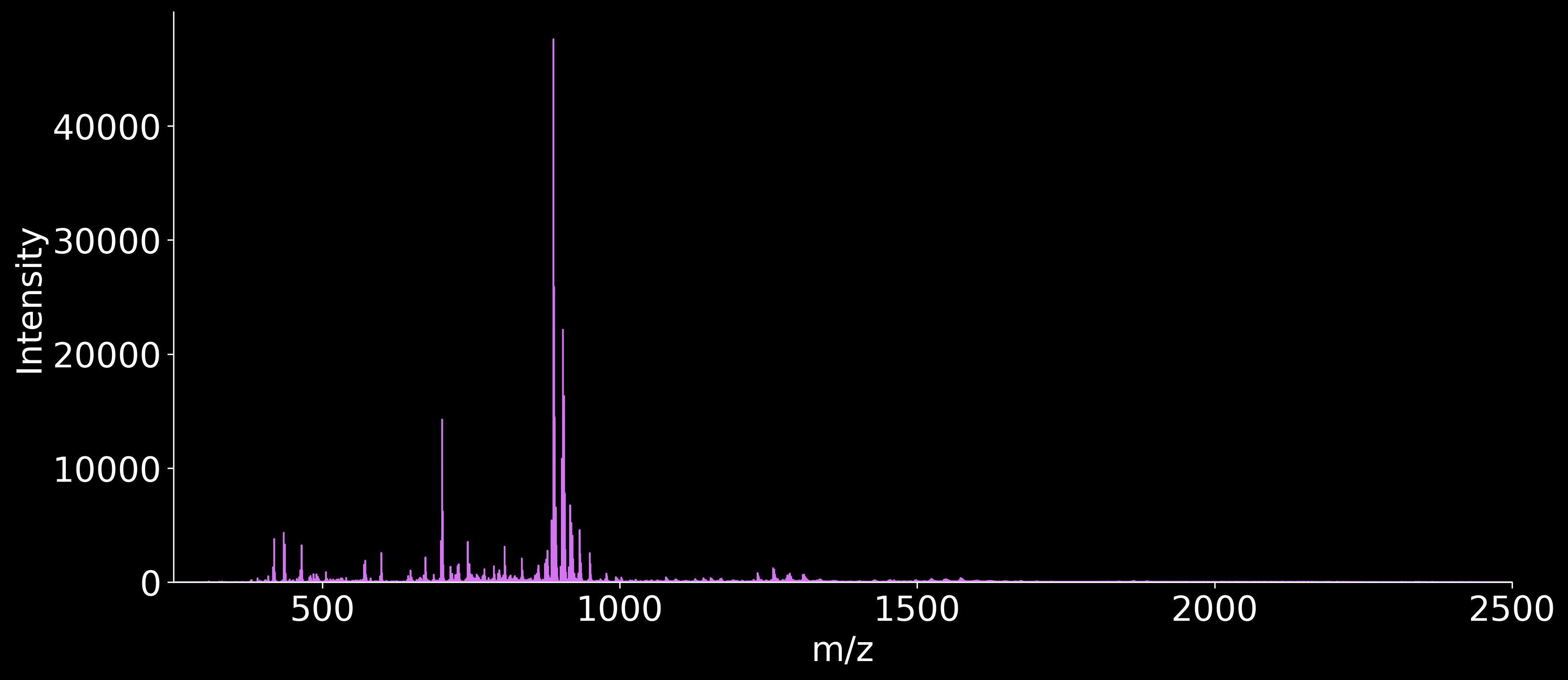

NAWM & WMH - Donor A posterior REPLICATE 2

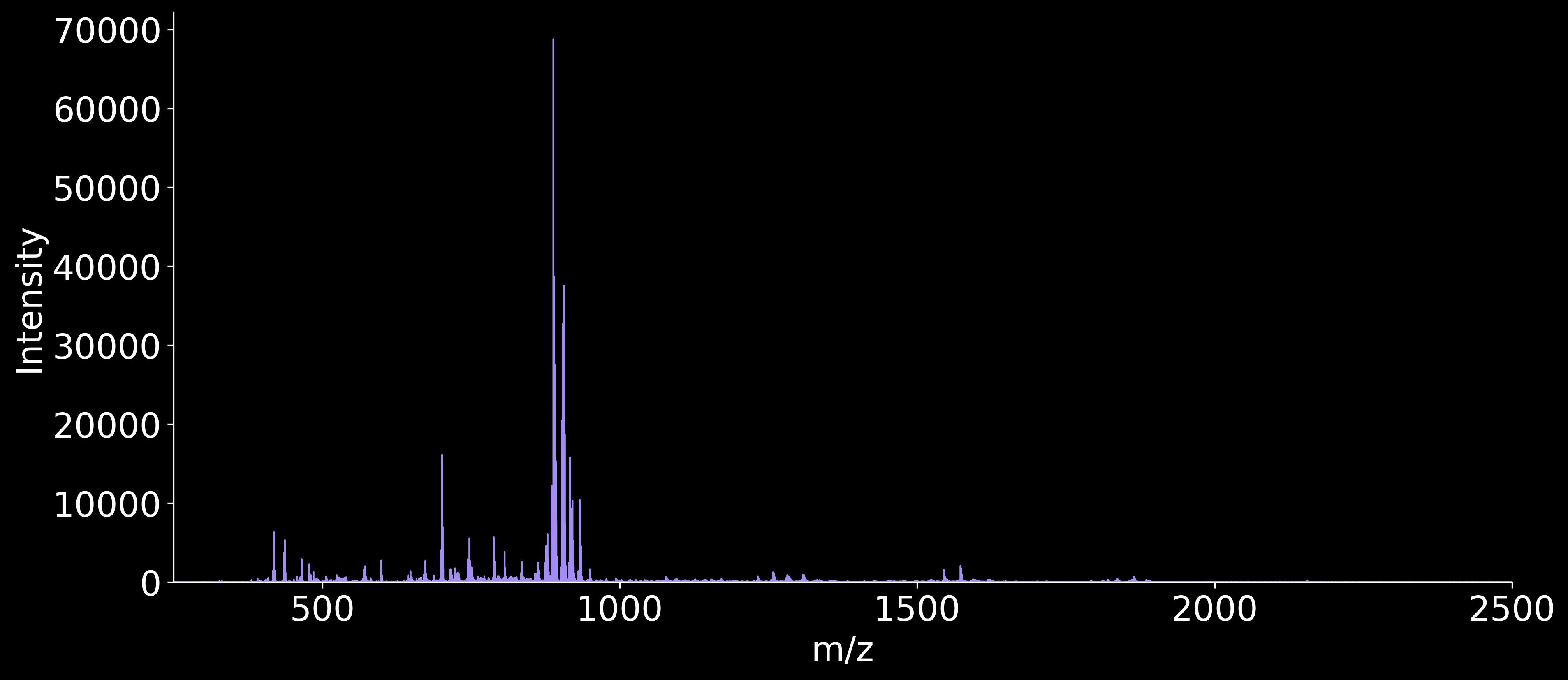

NAWM - Donor B posterior REPLICATE 2

*(Figure S6 continued on next page)*

**Figure S6 (continued)**

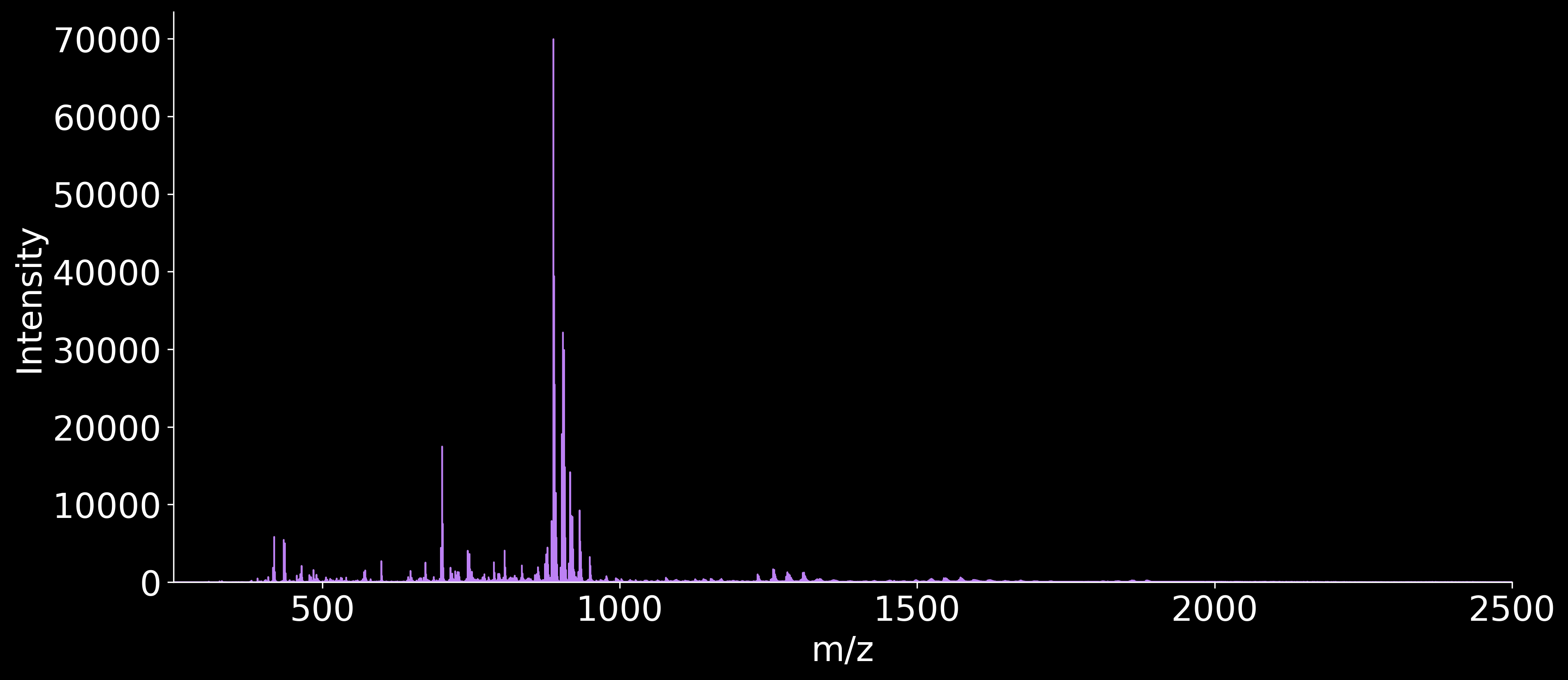

WMH - Donor B posterior REPLICATE 2

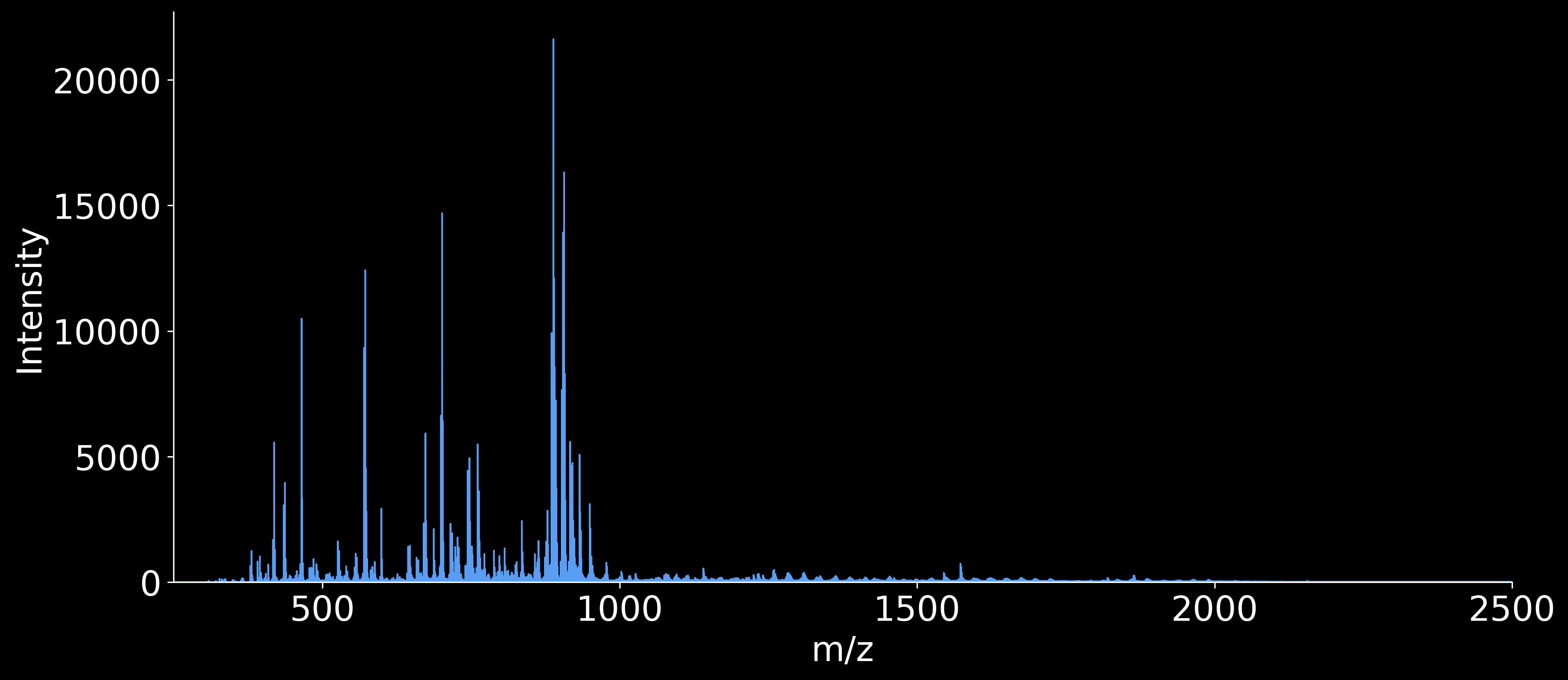

NAWM - Donor C posterior REPLICATE 2

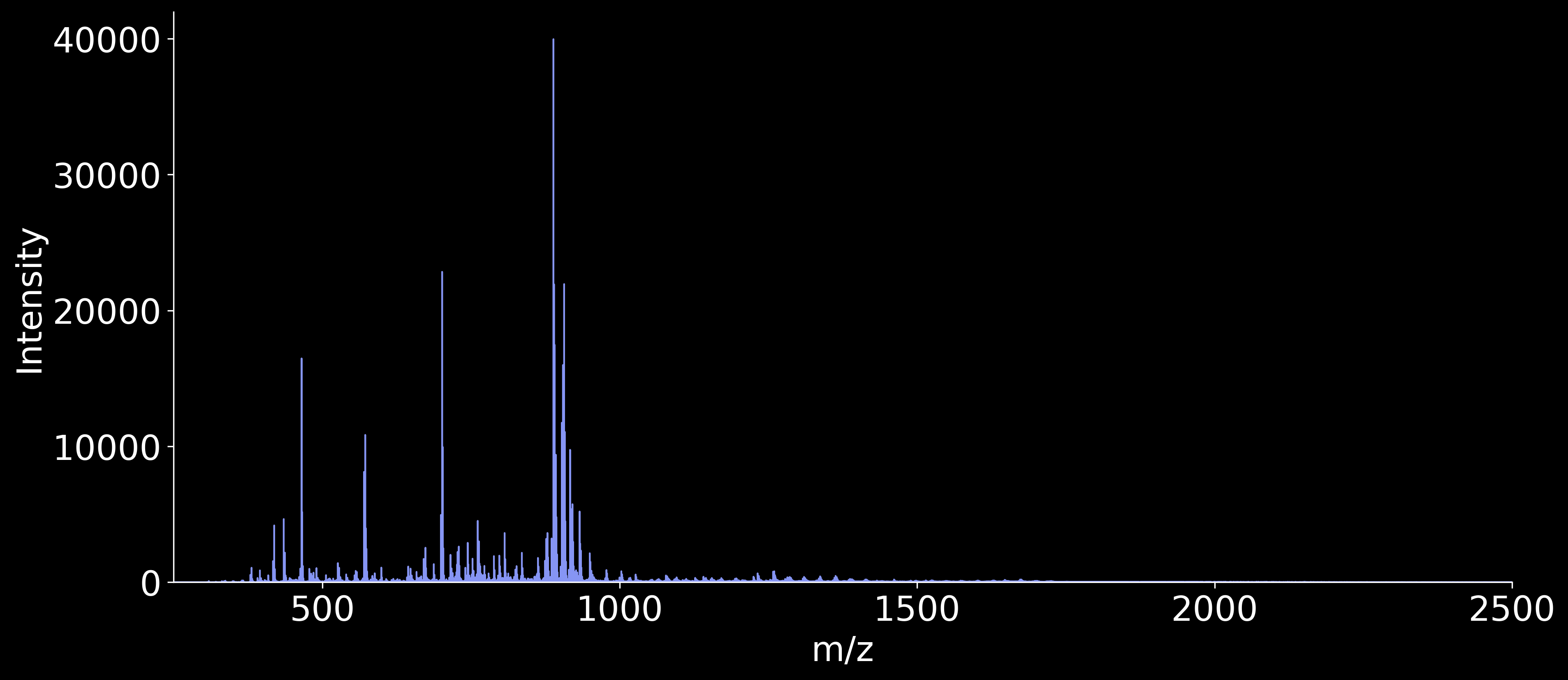

WMH - Donor C posterior REPLICATE 2

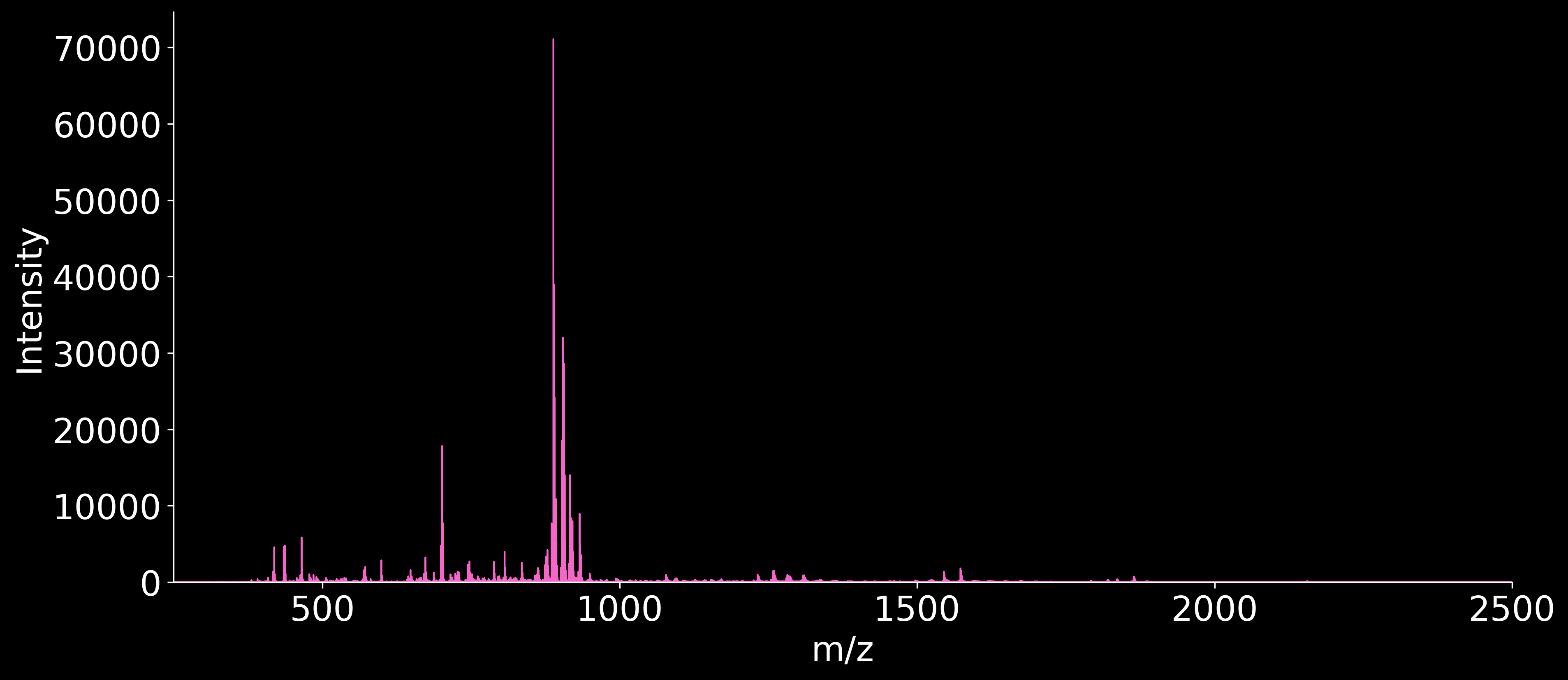

WMH - Donor B posterior REPLICATE 3

*(Figure S6 continued on next page)*

**Figure S6 (continued)**

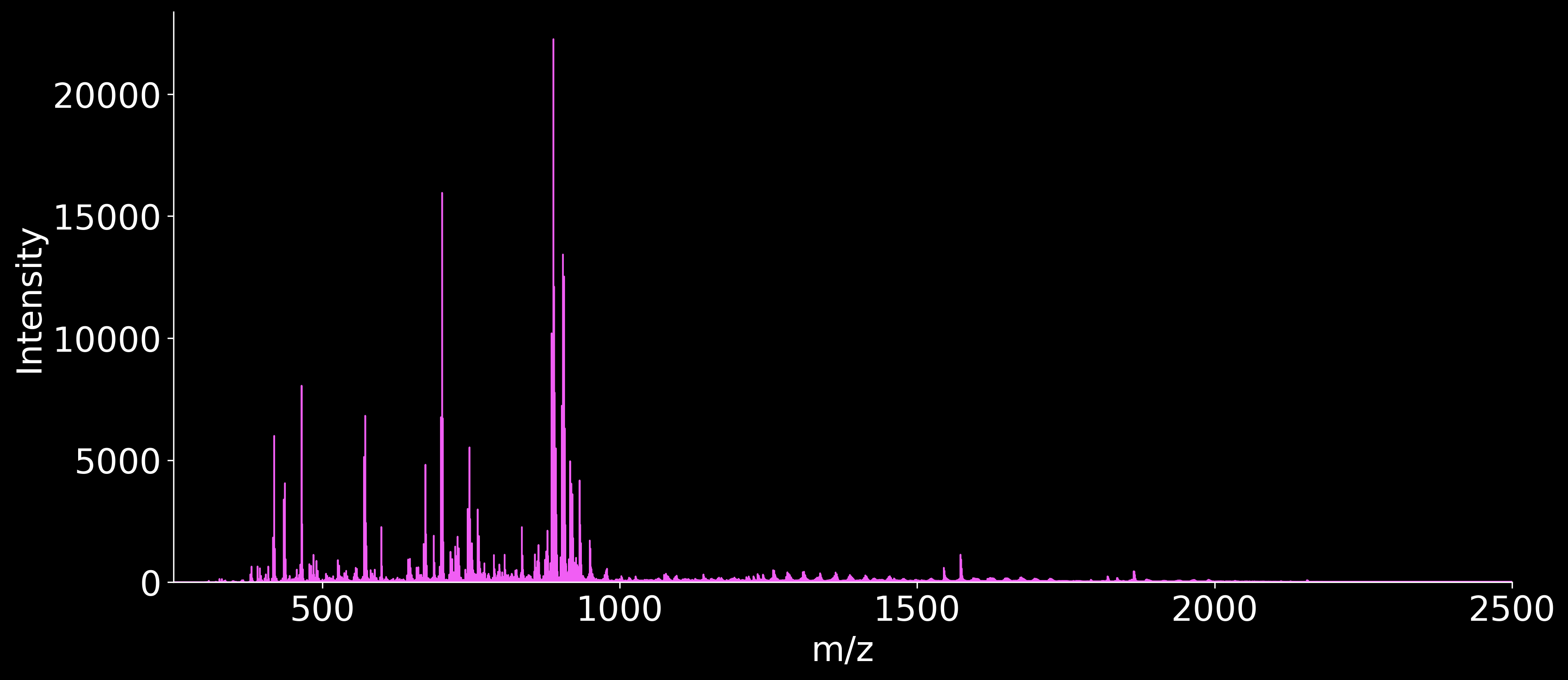

NAWM - Donor C posterior REPLICATE 3

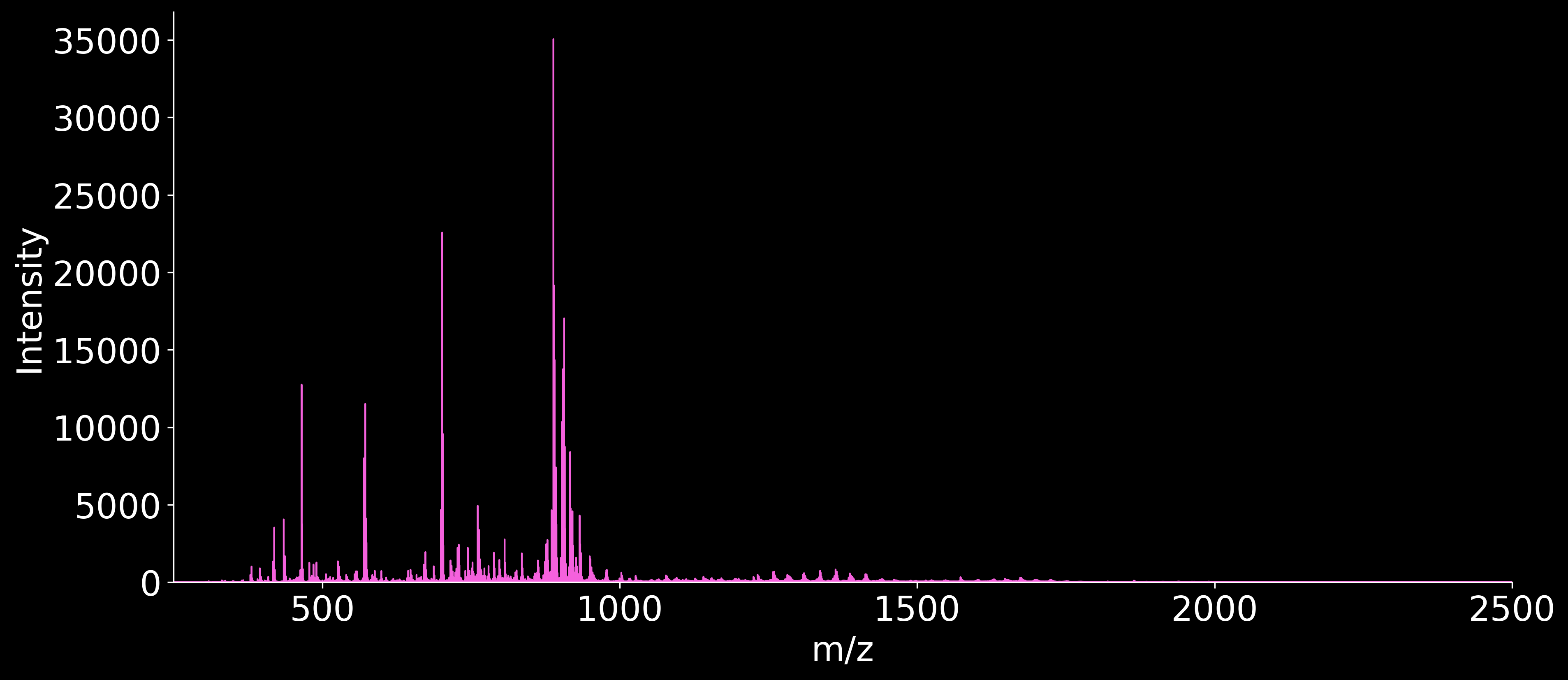

WMH - Donor C posterior REPLICATE 3

***Figure S6:*** *Average MALDI IMS Mass Spectra of each tissue sample, ordered by Donor and Replicate. The color of each spectrum is arbitrary.*

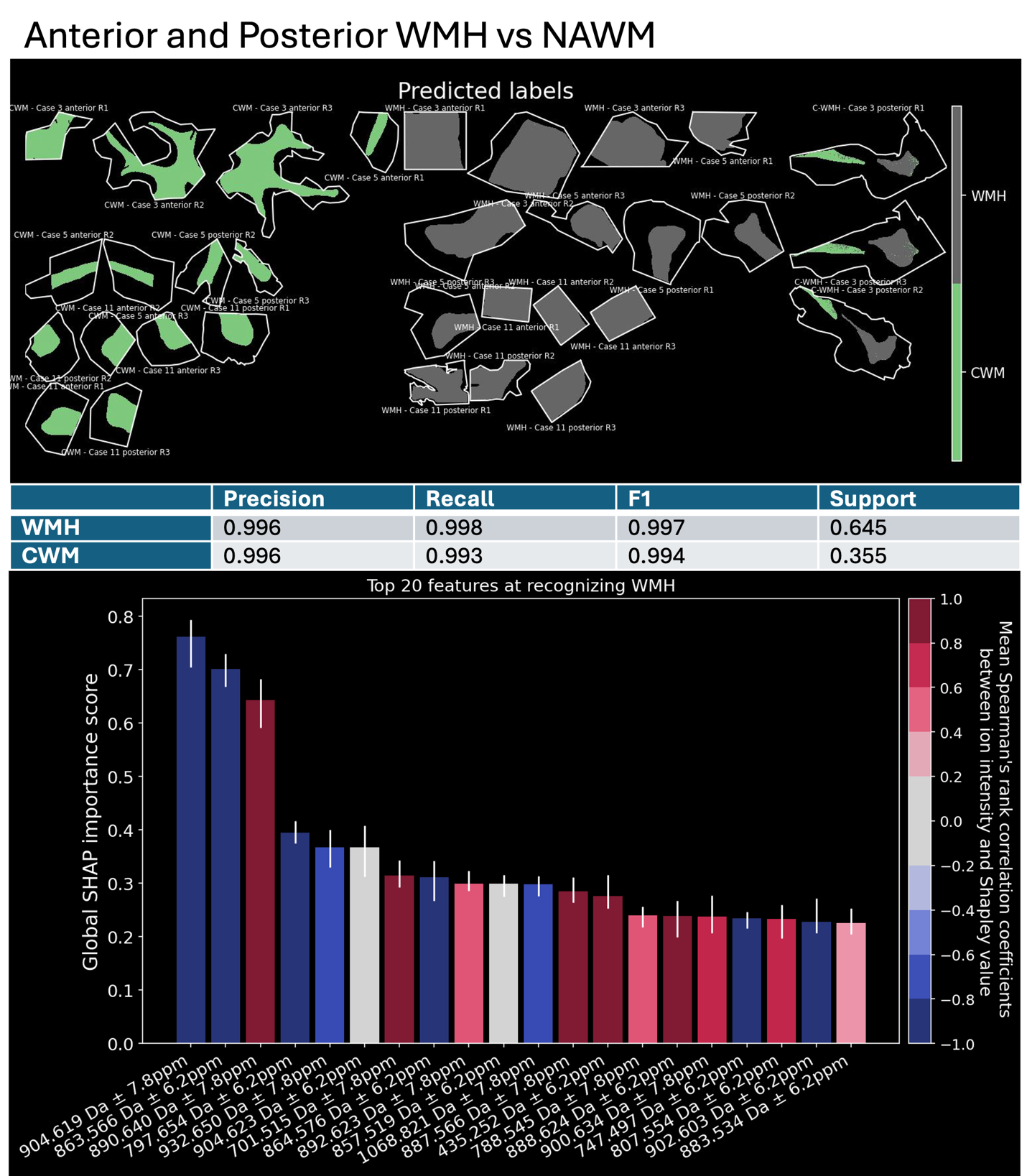

***Figure S7.*** *Predicted labels of XGBoost classification model and SHAP Precision, Recall, F1, and Support data, and SHAP bar plot of top 20 features in Anterior and Posterior WMH vs. NAWM. *Note:* ***Cases 3, 5, and 11 correspond to Donors A, B, and C, respectively****, matching the clinical profiles in Figure S1. WMH refers to white matter hyperintensity, CWM refers to control white matter, referenced in the main text of this paper as NAWM. Anterior / Posterior refers to the brain sample region, and “R#” refers to the replicate number (1-3).*

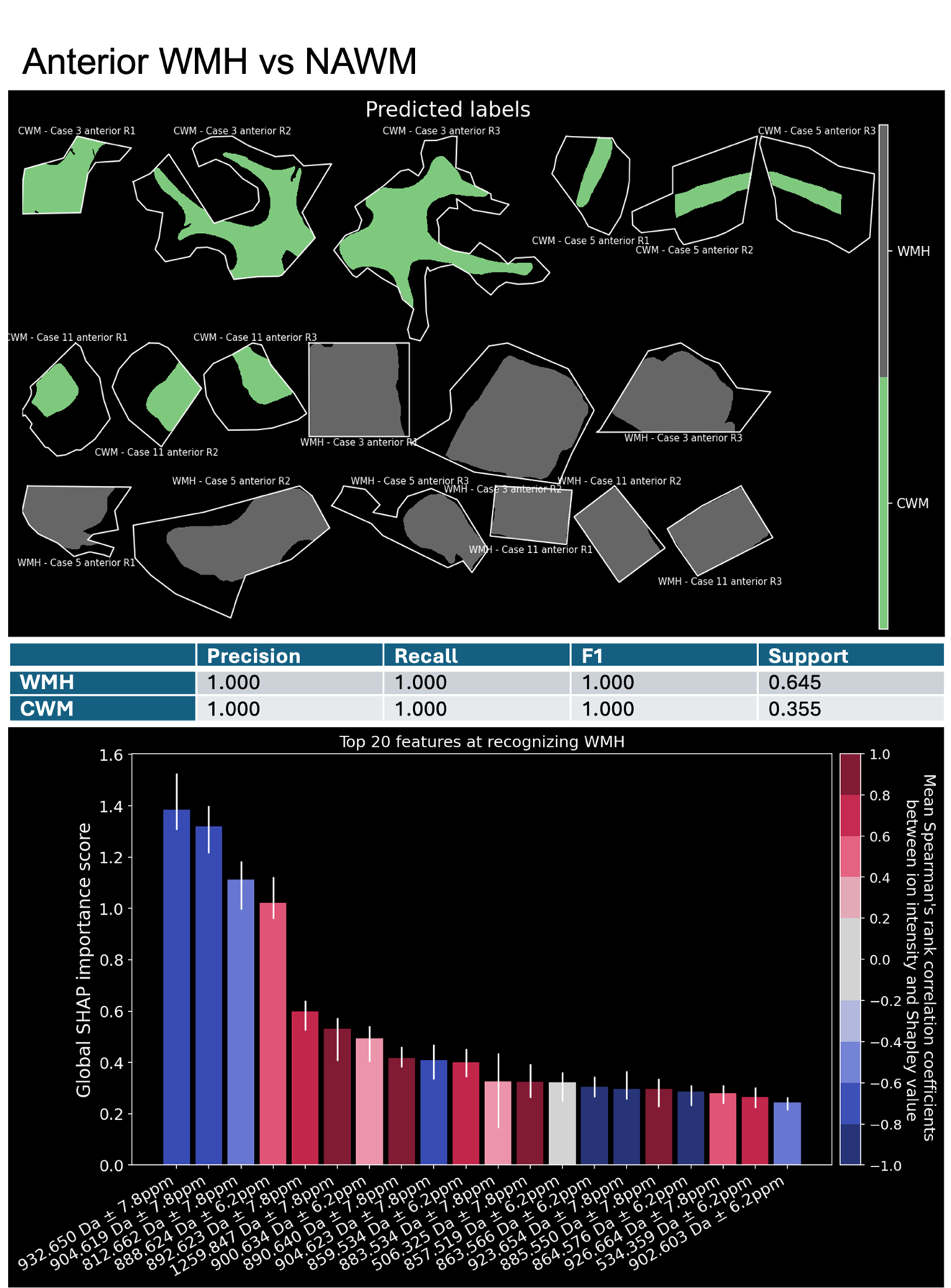

***Figure S8.*** *Predicted labels of XGBoost classification model and SHAP Precision, Recall, F1, and Support data, and SHAP bar plot of top 20 features in Anterior WMH vs. NAWM. *Note:* ***Cases 3, 5, and 11 correspond to Donors A, B, and C, respectively****, matching the clinical profiles in Figure S1. WMH refers to white matter hyperintensity, CWM refers to control white matter, referenced in the main text of this paper as NAWM. Anterior / Posterior refers to the brain sample region, and “R#” refers to the replicate number (1-3).*
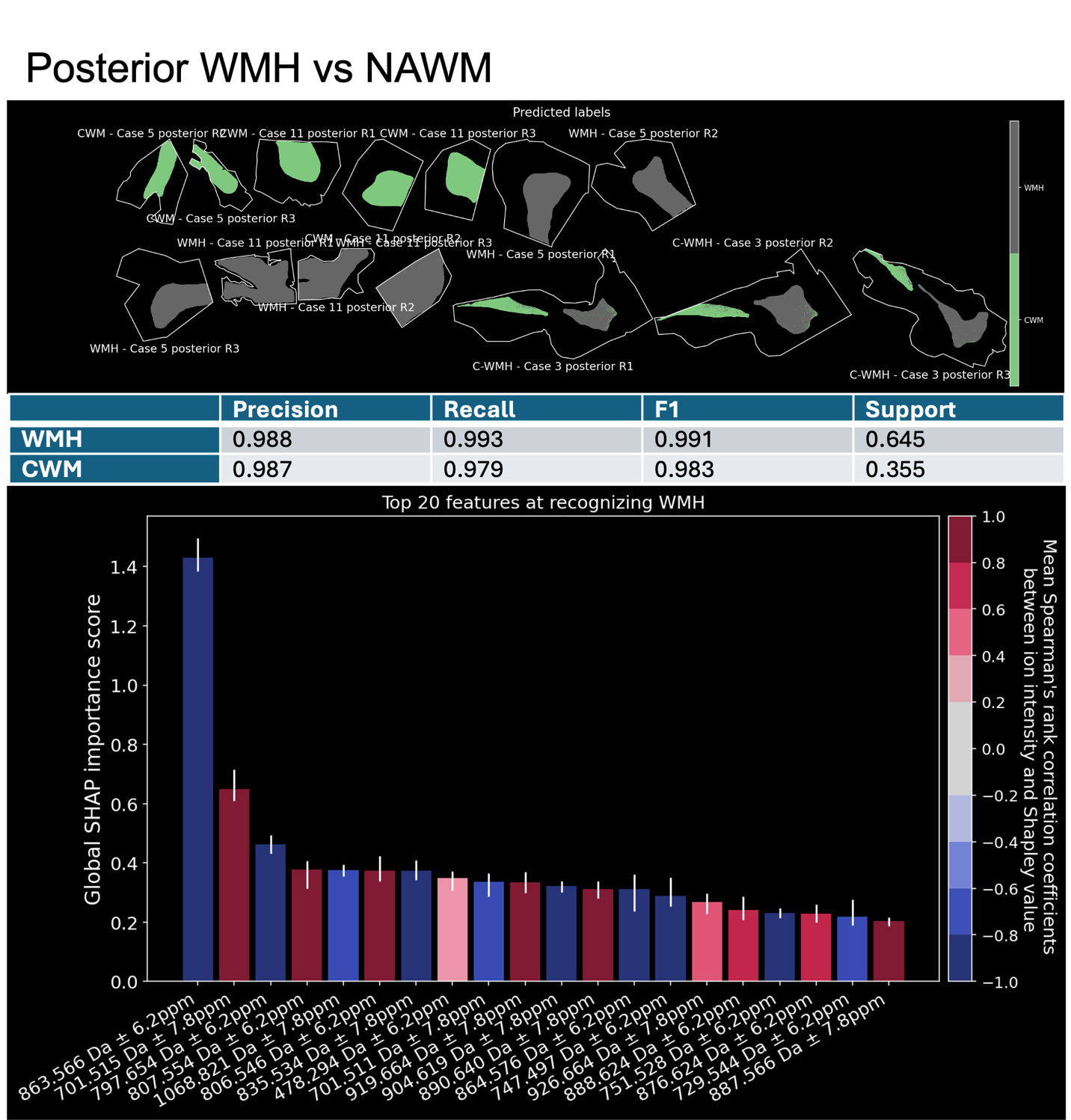

***Figure S9.*** *Predicted labels of XGBoost classification model and SHAP Precision, Recall, F1, and Support data, and SHAP bar plot of top 20 features in Posterior WMH vs. NAWM. *Note:* ***Cases 3, 5, and 11 correspond to Donors A, B, and C, respectively****, matching the clinical profiles in Figure S1. WMH refers to white matter hyperintensity, CWM refers to control white matter, referenced in the main text of this paper as NAWM. Anterior / Posterior refers to the brain sample region, and “R#” refers to the replicate number (1-3).*
