## Supplemental Doc 2 for "Elucidating Molecular Features of White Matter Hyperintensities in Alzheimer’s Disease through Multimodal Imaging and SHAP Analysis"

**SUPPLEMENT 2**

**LC-MS/MS Annotations of MALDI IMS-detected Lipids - Negative Mode**

| **abbrev** | **formula** | **chemical_class** | **has_ms/ms** |
| --- | --- | --- | --- |
| **FA 16:1** | C16H30O2 | FA | TRUE |
| **FA 16:0** | C16H32O2 | FA | TRUE |
| **FA 17:1** | C17H32O2 | FA | TRUE |
| **FA 17:0** | C17H34O2 | FA | TRUE |
| **FA 18:2** | C18H32O2 | FA | TRUE |
| **FA 18:1** | C18H34O2 | FA | TRUE |
| **FA 18:0** | C18H36O2 | FA | TRUE |
| **FA 19:2** | C19H34O2 | FA | TRUE |
| **FA 19:1** | C19H36O2 | FA | TRUE |
| **FA 19:0** | C19H38O2 | FA | TRUE |
| **NAGly 15:0** | C17H33NO3 | NAGly | TRUE |
| **FA 18:0;(2OH)** | C18H36O3 | OxFA | TRUE |
| **FA 20:4** | C20H32O2 | FA | TRUE |
| **FA 20:3** | C20H34O2 | FA | TRUE |
| **FA 20:2** | C20H36O2 | FA | TRUE |
| **FA 20:1** | C20H38O2 | FA | TRUE |
| **FA 20:0** | C20H40O2 | FA | TRUE |
| **FA 21:3** | C21H36O2 | FA | TRUE |
| **FA 21:2** | C21H38O2 | FA | TRUE |
| **AAHFA 4:0/15:2;O** | C19H32O4 | FAHFA | TRUE |
| **FA 21:1** | C21H40O2 | FA | TRUE |
| **Dodecylbenzenesulfonic acid** | C18H30O3S | Others | TRUE |
| **NAGly 17:0** | C19H37NO3 | NAGly | TRUE |
| **FA 22:6** | C22H32O2 | FA | TRUE |
| **FA 22:5** | C22H34O2 | FA | TRUE |
| **FA 22:4** | C22H36O2 | FA | TRUE |
| **FA 22:3** | C22H38O2 | FA | TRUE |
| **FA 22:2** | C22H40O2 | FA | TRUE |
| **FA 22:1** | C22H42O2 | FA | TRUE |
| **Norethisterone acetate** | C22H28O3 | Others | TRUE |
| **AAHFA 6:0/15:3;O** | C21H34O4 | FAHFA | TRUE |
| **AAHFA 6:0/15:2;O** | C21H36O4 | FAHFA | TRUE |
| **FA 23:1** | C23H44O2 | FA | TRUE |
| **FA 23:0** | C23H46O2 | FA | TRUE |
| **FA 24:6** | C24H36O2 | FA | FALSE |
| **FA 22:0;(2OH)** | C22H44O3 | OxFA | TRUE |
| **FA 24:5** | C24H38O2 | FA | TRUE |
| **FA 24:4** | C24H40O2 | FA | TRUE |
| **FA 24:3** | C24H42O2 | FA | TRUE |
| **FA 24:1** | C24H46O2 | FA | TRUE |
| **FA 24:0** | C24H48O2 | FA | TRUE |
| **FA 23:0;(2OH)** | C23H46O3 | OxFA | FALSE |
| **NAGlySer 13:1;O** | C18H32N2O6 | NAGlySer | TRUE |
| **AAHFA 5:0/18:5;O** | C23H34O4 | FAHFA | TRUE |
| **AAHFA 15:4/8:0;O** | C23H36O4 | FAHFA | TRUE |
| **AAHFA 15:3/8:0;O** | C23H38O4 | FAHFA | TRUE |
| **FA 25:1** | C25H48O2 | FA | TRUE |
| **FA 24:1;(2OH)** | C24H46O3 | OxFA | TRUE |
| **FA 25:0** | C25H50O2 | FA | TRUE |
| **LPG O-10:0** | C16H35O8P | EtherLPG | FALSE |
| **NAE 19:0** | C21H43NO2 | NAE | TRUE |
| **LPA 15:4** | C18H29O7P | LPA | TRUE |
| **FA 26:4** | C26H44O2 | FA | TRUE |
| **NATau 17:2;O** | C19H35NO5S | NATau | TRUE |
| **FA 26:3** | C26H46O2 | FA | TRUE |
| **NATau 17:1;O** | C19H37NO5S | NATau | FALSE |
| **NAGly 22:3** | C24H41NO3 | NAGly | TRUE |
| **NATau 17:0;O** | C19H39NO5S | NATau | FALSE |
| **Cer 15:3;2O/9:0** | C24H43NO3 | Cer_NS | TRUE |
| **FA 26:1** | C26H50O2 | FA | TRUE |
| **LPA 15:0** | C18H37O7P | LPA | TRUE |
| **FA 26:0** | C26H52O2 | FA | TRUE |
| **AAHFA 3:0/22:6;O** | C25H36O4 | FAHFA | TRUE |
| **Cer 12:1;3O/10:0;(2OH)** | C22H43NO5 | Cer_AP | TRUE |
| **AAHFA 7:0/18:5;O** | C25H38O4 | FAHFA | TRUE |
| **NAGlySer 15:0;O** | C20H38N2O6 | NAGlySer | TRUE |
| **FA 26:2;(2OH)** | C26H48O3 | OxFA | TRUE |
| **FA 27:1** | C27H52O2 | FA | TRUE |
| **FA 26:1;(2OH)** | C26H50O3 | OxFA | TRUE |
| **FA 27:0** | C27H54O2 | FA | TRUE |
| **PE O-8:0_4:0;1O** | C17H36NO8P | EtherOxPE | TRUE |
| **LPC O-8:0** | C16H36NO6P | EtherLPC | TRUE |
| **NATau 19:2;O** | C21H39NO5S | NATau | TRUE |
| **AAHFA 18:4/8:0;O** | C26H42O4 | FAHFA | TRUE |
| **NATau 19:1;O** | C21H41NO5S | NATau | FALSE |
| **FA 28:2** | C28H52O2 | FA | TRUE |
| **LPA 17:1** | C20H39O7P | LPA | TRUE |
| **FA 28:1** | C28H54O2 | FA | TRUE |
| **FA 28:0** | C28H56O2 | FA | TRUE |
| **Cer 12:0;2O/14:1** | C26H51NO3 | Cer_NDS | TRUE |
| **NAGlySer 17:1;O** | C22H40N2O6 | NAGlySer | TRUE |
| **AAHFA 5:0/22:6;O** | C27H40O4 | FAHFA | TRUE |
| **AAHFA 19:5/8:0;O** | C27H42O4 | FAHFA | TRUE |
| **AAHFA 19:4/8:0;O** | C27H44O4 | FAHFA | TRUE |
| **AAHFA 19:3/8:0;O** | C27H46O4 | FAHFA | TRUE |
| **LPA 18:1** | C21H41O7P | LPA | TRUE |
| **FA 29:1** | C29H56O2 | FA | TRUE |
| **LPA 18:0** | C21H43O7P | LPA | TRUE |
| **FA 29:0** | C29H58O2 | FA | TRUE |
| **LPA 19:5** | C22H35O7P | LPA | TRUE |
| **NATau 21:2;O** | C23H43NO5S | NATau | TRUE |
| **Cer 12:0;3O/10:0;(2OH)** | C22H45NO5 | Cer_AP | TRUE |
| **FA 30:1** | C30H58O2 | FA | TRUE |
| **LPE 16:1** | C21H42NO7P | LPE | TRUE |
| **FA 30:0** | C30H60O2 | FA | TRUE |
| **LPE 16:0** | C21H44NO7P | LPE | TRUE |
| **NAE 24:2** | C26H49NO2 | NAE | TRUE |
| **Cer 12:0;2O/16:1** | C28H55NO3 | Cer_NDS | TRUE |
| **LPE 17:1** | C22H44NO7P | LPE | TRUE |
| **NATau 24:4** | C26H45NO4S | NATau | TRUE |
| **NAPhe 21:3** | C30H45NO3 | NAPhe | TRUE |
| **SL 12:2;O/13:1** | C25H45NO5S | SL | TRUE |
| **LPG O-17:3** | C23H43O8P | EtherLPG | FALSE |
| **FA 32:1** | C32H62O2 | FA | TRUE |
| **LPE 18:1** | C23H46NO7P | LPE | TRUE |
| **LPE 18:0** | C23H48NO7P | LPE | TRUE |
| **PE O-8:0_10:0** | C23H48NO7P | EtherPE | TRUE |
| **Cer 12:0;2O/18:1** | C30H59NO3 | Cer_NDS | TRUE |
| **NATau 24:3;O** | C26H47NO5S | NATau | TRUE |
| **LPE 18:1(d7)** | C23H39D7NO7P | LPE | TRUE |
| **LPG O-18:3** | C24H45O8P | EtherLPG | TRUE |
| **NATau 26:5** | C28H47NO4S | NATau | TRUE |
| **CerP 12:0;2O/13:0** | C25H52NO6P | CerP | TRUE |
| **NAPhe 22:4;O** | C31H45NO4 | NAPhe | TRUE |
| **LPE O-20:0** | C25H54NO6P | EtherLPE | TRUE |
| **LPG 17:1** | C23H45O9P | LPG | TRUE |
| **SL 12:1;O/15:2** | C27H49NO5S | SL | TRUE |
| **NAGlySer 22:0;O** | C27H52N2O6 | NAGlySer | TRUE |
| **LPE 20:4** | C25H44NO7P | LPE | TRUE |
| **LPE 20:3** | C25H46NO7P | LPE | TRUE |
| **LPE 20:2** | C25H48NO7P | LPE | TRUE |
| **LPG O-19:3** | C25H47O8P | EtherLPG | TRUE |
| **LPC O-15:3** | C23H44NO6P | EtherLPC | TRUE |
| **LPE 20:1** | C25H50NO7P | LPE | TRUE |
| **NATau 26:5;O** | C28H47NO5S | NATau | TRUE |
| **LPE 20:0** | C25H52NO7P | LPE | TRUE |
| **PE O-8:0_12:0** | C25H52NO7P | EtherPE | TRUE |
| **Cer 12:0;2O/20:1** | C32H63NO3 | Cer_NDS | TRUE |
| **LPG 18:1** | C24H47O9P | LPG | TRUE |
| **NATau 28:6** | C30H49NO4S | NATau | TRUE |
| **LPE 21:1** | C26H52NO7P | LPE | TRUE |
| **NATau 28:5** | C30H51NO4S | NATau | TRUE |
| **LPG 19:2** | C25H47O9P | LPG | TRUE |
| **LPS 18:1** | C24H46NO9P | LPS | TRUE |
| **SL 12:1;O/17:4** | C29H49NO5S | SL | TRUE |
| **Cer 12:0;2O/21:1** | C33H65NO3 | Cer_NDS | TRUE |
| **LPE 22:6** | C27H44NO7P | LPE | TRUE |
| **LPS 18:0** | C24H48NO9P | LPS | TRUE |
| **SL 12:1;O/16:4;O** | C28H47NO6S | SL | TRUE |
| **NAGly 15:3;O(FA 15:4)** | C32H47NO5 | NAGly | TRUE |
| **LPE 22:5** | C27H46NO7P | LPE | TRUE |
| **LPC O-16:0** | C24H52NO6P | EtherLPC | TRUE |
| **FA 36:4** | C36H64O2 | FA | TRUE |
| **LPE 22:4** | C27H48NO7P | LPE | TRUE |
| **LPE 22:3** | C27H50NO7P | LPE | TRUE |
| **LPE 22:2** | C27H52NO7P | LPE | TRUE |
| **LPE 22:1** | C27H54NO7P | LPE | TRUE |
| **PE O-8:0_14:1** | C27H54NO7P | EtherPE | TRUE |
| **Cer 12:0;2O/22:2** | C34H65NO3 | Cer_NDS | TRUE |
| **SL 13:1;O/16:4;O** | C29H49NO6S | SL | TRUE |
| **Cer 12:0;2O/22:0** | C34H69NO3 | Cer_NDS | TRUE |
| **LPC 16:0** | C24H50NO7P | LPC | TRUE |
| **AAHFA 26:5/9:0;O** | C35H58O4 | FAHFA | TRUE |
| **LPC 17:4** | C25H44NO7P | LPC | TRUE |
| **PE O-8:0_15:2** | C28H54NO7P | EtherPE | TRUE |
| **LPC 17:3** | C25H46NO7P | LPC | TRUE |
| **CerP 12:2;2O/18:5** | C30H48NO6P | CerP | TRUE |
| **GM3 30:0;2O** | C53H98N2O21 | GM3 | TRUE |
| **LPE 23:1** | C28H56NO7P | LPE | TRUE |
| **NAPhe 26:5;O** | C35H51NO4 | NAPhe | TRUE |
| **SL 12:1;O/18:5;O** | C30H49NO6S | SL | TRUE |
| **Cer 12:0;2O/23:1** | C35H69NO3 | Cer_NDS | TRUE |
| **SL 12:1;O/18:4;O** | C30H51NO6S | SL | TRUE |
| **LPC O-18:1** | C26H54NO6P | EtherLPC | TRUE |
| **Cer 12:1;3O/21:1;(2OH)** | C33H63NO5 | Cer_AP | TRUE |
| **Cer 18:1;3O/15:1;(2OH)** | C33H63NO5 | Cer_AP | TRUE |
| **PE-Cer 27:5;2O** | C29H51N2O6P | PE_Cer | TRUE |
| **PE-Cer 12:2;2O/15:3** | C29H51N2O6P | PE_Cer | TRUE |
| **GM3 31:1;2O** | C54H98N2O21 | GM3 | TRUE |
| **LPE 24:5** | C29H50NO7P | LPE | TRUE |
| **LPC 17:0** | C25H52NO7P | LPC | TRUE |
| **LPC O-18:0** | C26H56NO6P | EtherLPC | TRUE |
| **Cer 12:1;3O/21:0;(2OH)** | C33H65NO5 | Cer_AP | TRUE |
| **FA 38:4** | C38H68O2 | FA | TRUE |
| **PE O-8:0_16:4** | C29H52NO7P | EtherPE | TRUE |
| **LPE 24:4** | C29H52NO7P | LPE | TRUE |
| **PA 8:0_18:3** | C29H51O8P | PA | TRUE |
| **LPE 24:3** | C29H54NO7P | LPE | TRUE |
| **PE O-8:0_16:3** | C29H54NO7P | EtherPE | TRUE |
| **LPC 18:4** | C26H46NO7P | LPC | FALSE |
| **LPE 24:2** | C29H56NO7P | LPE | TRUE |
| **LPA 27:1** | C30H59O7P | LPA | TRUE |
| **CerP 12:2;2O/19:5** | C31H50NO6P | CerP | TRUE |
| **LPE 24:1** | C29H58NO7P | LPE | TRUE |
| **PE O-8:0_16:1** | C29H58NO7P | EtherPE | TRUE |
| **CerP 12:1;2O/19:5** | C31H52NO6P | CerP | TRUE |
| **Cer 12:0;2O/24:1** | C36H71NO3 | Cer_NDS | TRUE |
| **LPI 16:3** | C25H43O12P | LPI | FALSE |
| **AAHFA 28:0/8:0;O** | C36H70O4 | FAHFA | TRUE |
| **LPC 18:1** | C26H52NO7P | LPC | TRUE |
| **SL 31:5;2O** | C31H53NO6S | SL | TRUE |
| **SL 13:1;O/18:4;O** | C31H53NO6S | SL | TRUE |
| **Cer 12:0;2O/24:0** | C36H73NO3 | Cer_NDS | TRUE |
| **LPS 22:6** | C28H44NO9P | LPS | TRUE |
| **PS 8:0_10:0;2O** | C24H46NO12P | OxPS | TRUE |
| **LPS 22:5** | C28H46NO9P | LPS | TRUE |
| **LPI 16:0** | C25H49O12P | LPI | TRUE |
| **LPA 28:2** | C31H59O7P | LPA | TRUE |
| **AAHFA 22:6/16:4;O** | C38H54O4 | FAHFA | TRUE |
| **LPC 19:4** | C27H48NO7P | LPC | TRUE |
| **SL 12:2;O/20:5;O** | C32H51NO6S | SL | TRUE |
| **LPC 19:3** | C27H50NO7P | LPC | TRUE |
| **NAPhe 28:5;O** | C37H55NO4 | NAPhe | TRUE |
| **Cer 12:1;3O/23:0;(2OH)** | C35H69NO5 | Cer_AP | TRUE |
| **PA 8:0_20:4** | C31H53O8P | PA | TRUE |
| **CL 12:0_12:0_12:0_15:0** | C60H116O17P2 | CL | TRUE |
| **Cer 12:0;3O/23:0;(2OH)** | C35H71NO5 | Cer_AP | TRUE |
| **PI 10:0_6:0** | C25H47O13P | PI | TRUE |
| **PA 8:0_20:3** | C31H55O8P | PA | TRUE |
| **SL 12:2;O/22:6** | C34H53NO5S | SL | FALSE |
| **PE O-8:0_18:3** | C31H58NO7P | EtherPE | TRUE |
| **SL 15:3;O/18:5;O** | C33H51NO6S | SL | TRUE |
| **LPC 20:4** | C28H50NO7P | LPC | TRUE |
| **CerP 33:7;2O** | C33H54NO6P | CerP | TRUE |
| **GM3 36:0;2O** | C59H110N2O21 | GM3 | TRUE |
| **GM3 37:5;2O** | C60H102N2O21 | GM3 | TRUE |
| **PE O-8:0_18:0** | C31H64NO7P | EtherPE | FALSE |
| **Cer 12:0;2O/26:1** | C38H75NO3 | Cer_NDS | TRUE |
| **SG 27:1;O;Hex** | C33H56O6 | SHex | TRUE |
| **GM3 37:3;2O** | C60H106N2O21 | GM3 | TRUE |
| **LPC 20:1** | C28H56NO7P | LPC | TRUE |
| **LPS 24:6** | C30H48NO9P | LPS | TRUE |
| **LPC 20:0** | C28H58NO7P | LPC | TRUE |
| **LPI 18:1** | C27H51O12P | LPI | TRUE |
| **Cer 12:0;3O/24:0;(2OH)** | C36H73NO5 | Cer_AP | TRUE |
| **LPI 18:0** | C27H53O12P | LPI | TRUE |
| **PI O-8:0_10:0** | C27H53O12P | EtherPI | TRUE |
| **PS O-8:0_16:4** | C30H52NO9P | EtherPS | FALSE |
| **SE 24:1;O4/13:1;1O** | C37H62O6 | DCAE | TRUE |
| **PS O-8:0_16:3** | C30H54NO9P | EtherPS | TRUE |
| **CerP 12:2;2O/22:6** | C34H54NO6P | CerP | TRUE |
| **SL 12:2;O/22:6;O** | C34H53NO6S | SL | TRUE |
| **PE-Cer 12:1;2O/18:0** | C32H65N2O6P | PE_Cer | TRUE |
| **SM 12:2;2O/12:0** | C29H57N2O6P | SM | TRUE |
| **Cer 12:0;2O/27:1** | C39H77NO3 | Cer_NDS | TRUE |
| **PC 16:2_3:0;1O** | C27H50NO9P | OxPC | TRUE |
| **AAHFA 21:2/19:4;O** | C40H66O4 | FAHFA | TRUE |
| **AAHFA 28:6/12:0;O** | C40H66O4 | FAHFA | TRUE |
| **PE 22:6_4:0;1O** | C31H50NO9P | OxPE | TRUE |
| **Cer 12:1;3O/25:0;(2OH)** | C37H73NO5 | Cer_AP | TRUE |
| **NAGlySer 8:0;O(FA 21:0)** | C34H64N2O7 | NAGlySer | TRUE |
| **AAHFA 28:5/12:0;O** | C40H68O4 | FAHFA | TRUE |
| **Cer 12:0;3O/25:0;(2OH)** | C37H75NO5 | Cer_AP | TRUE |
| **PS 20:5_3:0;1O** | C29H46NO11P | OxPS | FALSE |
| **LPC 22:4** | C30H54NO7P | LPC | TRUE |
| **PE O-8:0_20:2** | C33H64NO7P | EtherPE | TRUE |
| **Cer 13:1;3O/22:5;(2OH)** | C35H59NO5 | Cer_AP | TRUE |
| **Cer 12:0;2O/28:2** | C40H77NO3 | Cer_NDS | TRUE |
| **PA 15:3_16:4** | C34H53O8P | PA | TRUE |
| **GM3 41:5;2O** | C64H110N2O21 | GM3 | TRUE |
| **LPC 22:2** | C30H58NO7P | LPC | TRUE |
| **PE O-8:0_20:0** | C33H68NO7P | EtherPE | FALSE |
| **Cer 12:0;2O/28:1** | C40H79NO3 | Cer_NDS | TRUE |
| **CerP 13:1;2O/22:4** | C35H62NO6P | CerP | TRUE |
| **LPS 26:6** | C32H52NO9P | LPS | TRUE |
| **PE O-22:6_6:0;1O** | C33H56NO8P | EtherOxPE | TRUE |
| **AAHFA 28:5/13:0;O** | C41H70O4 | FAHFA | TRUE |
| **AAHFA 28:4/13:0;O** | C41H72O4 | FAHFA | TRUE |
| **AAHFA 28:3/13:0;O** | C41H74O4 | FAHFA | TRUE |
| **PC O-8:0_15:4** | C31H56NO7P | EtherPC | TRUE |
| **SL 13:1;O/24:6** | C37H61NO5S | SL | TRUE |
| **Cer 13:0;2O/28:3** | C41H77NO3 | Cer_NDS | TRUE |
| **Cer 13:0;2O/28:2** | C41H79NO3 | Cer_NDS | TRUE |
| **AAHFA 26:6/16:2;O** | C42H66O4 | FAHFA | TRUE |
| **PC 18:3_3:0;1O** | C29H52NO9P | OxPC | TRUE |
| **SL 12:1;O/24:5;O** | C36H61NO6S | SL | TRUE |
| **Cer 12:0;2O/29:1** | C41H81NO3 | Cer_NDS | TRUE |
| **ASG 27:1;O;Hex;FA 2:0** | C35H58O7 | AHexCS | TRUE |
| **PC 16:2_5:0;1O** | C29H54NO9P | OxPC | TRUE |
| **Cer 12:0;2O/29:0** | C41H83NO3 | Cer_NDS | FALSE |
| **AAHFA 28:6/14:0;O** | C42H70O4 | FAHFA | TRUE |
| **PI 14:0_6:0** | C29H55O13P | PI | TRUE |
| **PG O-8:0_20:5** | C34H59O9P | EtherPG | TRUE |
| **PS 20:5_5:0;1O** | C31H50NO11P | OxPS | TRUE |
| **PE O-8:0_22:3** | C35H66NO7P | EtherPE | TRUE |
| **LPC 24:4** | C32H58NO7P | LPC | TRUE |
| **Cer 13:1;3O/24:6;(2OH)** | C37H61NO5 | Cer_AP | TRUE |
| **PE O-8:0_22:2** | C35H68NO7P | EtherPE | TRUE |
| **SM 12:1;2O/15:2** | C32H61N2O6P | SM | TRUE |
| **CL 12:0_12:0_18:0_18:2** | C69H130O17P2 | CL | TRUE |
| **Cer 13:1;3O/24:5;(2OH)** | C37H63NO5 | Cer_AP | TRUE |
| **HexCer 16:0;2O/13:0;O** | C35H69NO9 | HexCer_HDS | TRUE |
| **PE 15:3_15:4** | C35H56NO8P | PE | TRUE |
| **SL 13:1;O/24:5;O** | C37H63NO6S | SL | TRUE |
| **Cer 13:1;3O/24:4;(2OH)** | C37H65NO5 | Cer_AP | TRUE |
| **Cer 12:0;2O/30:1** | C42H83NO3 | Cer_NDS | TRUE |
| **PE 6:0_24:6** | C35H58NO8P | PE | TRUE |
| **PC O-16:2_7:0;1O** | C31H60NO8P | EtherOxPC | FALSE |
| **CerP 13:1;2O/24:4** | C37H66NO6P | CerP | TRUE |
| **SHexCer 25:1;2O** | C31H59NO11S | SHexCer | TRUE |
| **SL 13:1;O/24:3;O** | C37H67NO6S | SL | TRUE |
| **SE 24:1;O4/17:3;1O** | C41H66O6 | DCAE | TRUE |
| **AAHFA 28:5/15:0;O** | C43H74O4 | FAHFA | TRUE |
| **CerP 13:1;2O/24:2** | C37H70NO6P | CerP | FALSE |
| **AAHFA 28:4/15:0;O** | C43H76O4 | FAHFA | TRUE |
| **PE-Cer 12:1;2O/22:1** | C36H71N2O6P | PE_Cer | TRUE |
| **SE 24:1;O4/17:1;O** | C41H70O6 | DCAE | TRUE |
| **Cer 13:0;2O/30:3** | C43H81NO3 | Cer_NDS | TRUE |
| **PE-Cer 12:1;2O/22:0** | C36H73N2O6P | PE_Cer | TRUE |
| **PE O-12:0_17:2;2O** | C34H66NO9P | EtherOxPE | FALSE |
| **CerP 12:1;2O/26:5** | C38H66NO6P | CerP | FALSE |
| **SL 12:1;O/26:5;O** | C38H65NO6S | SL | FALSE |
| **SL 13:1;O/26:4** | C39H69NO5S | SL | TRUE |
| **Cer 13:1;3O/28:2;(2OH)** | C41H77NO5 | Cer_AP | TRUE |
| **Cer 12:0;2O/31:1** | C43H85NO3 | Cer_NDS | TRUE |
| **PE 8:0_18:3;4O** | C31H56NO12P | OxPE | TRUE |
| **PE O-8:0_24:6** | C37H64NO7P | EtherPE | TRUE |
| **LPC 25:1** | C33H66NO7P | LPC | TRUE |
| **SL 12:1;O/26:3;O** | C38H69NO6S | SL | TRUE |
| **SL 13:1;O/26:2** | C39H73NO5S | SL | TRUE |
| **Cer 12:1;3O/29:0;(2OH)** | C41H81NO5 | Cer_AP | TRUE |
| **SE 24:1;O4/19:2** | C43H72O5 | DCAE | TRUE |
| **AAHFA 28:5/16:0;O** | C44H76O4 | FAHFA | TRUE |
| **PC 22:6_2:0;1O** | C32H52NO9P | OxPC | TRUE |
| **Cer 13:2;3O/26:6;(2OH)** | C39H63NO5 | Cer_AP | TRUE |
| **PE O-8:0_24:2** | C37H72NO7P | EtherPE | TRUE |
| **Cer 12:0;2O/32:3** | C44H83NO3 | Cer_NDS | TRUE |
| **Cer 13:1;3O/26:5;(2OH)** | C39H67NO5 | Cer_AP | TRUE |
| **Cer 12:0;2O/32:1** | C44H87NO3 | Cer_NDS | TRUE |
| **ASG 28:1;O;Hex;FA 4:0** | C38H64O7 | AHexCAS | TRUE |
| **LPC 26:1** | C34H68NO7P | LPC | TRUE |
| **HexCer 16:0;2O/12:0;O** | C34H67NO9 | HexCer_HDS | TRUE |
| **AAHFA 28:6/17:0;O** | C45H76O4 | FAHFA | TRUE |
| **LPC 26:0** | C34H70NO7P | LPC | TRUE |
| **AAHFA 24:6/22:6;O** | C46H66O4 | FAHFA | TRUE |
| **PC O-8:0_16:1;2O** | C32H64NO9P | EtherOxPC | TRUE |
| **FAHFA 22:4/24:6;O** | C46H70O4 | FAHFA | TRUE |
| **PE-Cer 36:2;2O** | C38H75N2O6P | PE_Cer | TRUE |
| **SE 24:1;O4/19:1;1O** | C43H74O6 | DCAE | TRUE |
| **PG 6:0_24:3** | C36H65O10P | PG | TRUE |
| **PA 8:0_27:1** | C38H73O8P | PA | TRUE |
| **LPC O-28:3** | C36H70NO6P | EtherLPC | FALSE |
| **Cer 13:0;2O/32:2** | C45H87NO3 | Cer_NDS | TRUE |
| **PE-Cer 12:0;2O/24:0** | C38H79N2O6P | PE_Cer | TRUE |
| **SL 13:1;O/28:4** | C41H73NO5S | SL | TRUE |
| **PC O-8:0_18:2;1O** | C34H66NO8P | EtherOxPC | TRUE |
| **PE 16:0_14:1;2O** | C35H68NO10P | OxPE | TRUE |
| **HexCer 16:1;3O/12:0;(2OH)** | C34H65NO10 | HexCer_AP | TRUE |
| **SL 12:0;O/28:5;O** | C40H71NO6S | SL | TRUE |
| **Cer 12:1;3O/31:1;(2OH)** | C43H83NO5 | Cer_AP | TRUE |
| **PI O-8:0_17:2** | C34H63O12P | EtherPI | FALSE |
| **CL 12:0_12:0_22:4_22:6** | C77H130O17P2 | CL | TRUE |
| **PE-Cer 12:1;2O/24:5;O** | C38H67N2O7P | PE_Cer | FALSE |
| **PC 10:0_15:1;1O** | C33H64NO9P | OxPC | TRUE |
| **GM3 52:8;2O** | C75H126N2O21 | GM3 | TRUE |
| **CL 12:0_12:0_22:3_22:6** | C77H132O17P2 | CL | TRUE |
| **PE-Cer 12:1;2O/24:4;O** | C38H69N2O7P | PE_Cer | FALSE |
| **AAHFA 28:5/18:0;O** | C46H80O4 | FAHFA | TRUE |
| **Cer 12:1;3O/28:1;(2OH)** | C40H77NO5 | Cer_AP | FALSE |
| **Cer 12:0;2O/34:5** | C46H83NO3 | Cer_NDS | TRUE |
| **PG O-8:0_24:5** | C38H67O9P | EtherPG | TRUE |
| **HexCer 16:0;2O/18:1** | C40H77NO8 | HexCer_NDS | TRUE |
| **HexCer 16:1;3O/16:1;(2OH)** | C38H71NO10 | HexCer_AP | TRUE |
| **Cer 12:0;3O/28:0;(2OH)** | C40H81NO5 | Cer_AP | TRUE |
| **SM 12:1;2O/19:2** | C36H69N2O6P | SM | TRUE |
| **NAGlySer 8:0;O(FA 28:4)** | C41H70N2O7 | NAGlySer | TRUE |
| **SE 24:1;O4/22:6** | C46H70O5 | DCAE | TRUE |
| **PE-Cer 12:1;2O/24:1;O** | C38H75N2O7P | PE_Cer | TRUE |
| **PE-Cer 12:1;2O/25:0** | C39H79N2O6P | PE_Cer | FALSE |
| **Cer 13:1;3O/28:5;(2OH)** | C41H71NO5 | Cer_AP | TRUE |
| **HexCer 16:0;2O/17:0;O** | C39H77NO9 | HexCer_HDS | TRUE |
| **SM 12:1;2O/19:1** | C36H71N2O6P | SM | TRUE |
| **SL 12:1;O/30:4** | C42H75NO5S | SL | FALSE |
| **LPC 28:1** | C36H72NO7P | LPC | TRUE |
| **Cer 13:1;2O/34:6** | C47H81NO3 | Cer_NS | TRUE |
| **AAHFA 28:6/19:0;O** | C47H80O4 | FAHFA | TRUE |
| **Cer 13:0;2O/34:6** | C47H83NO3 | Cer_NDS | TRUE |
| **PE 15:0_18:1(d7)** | C38H67D7NO8P | PE | TRUE |
| **AAHFA 28:5/19:0;O** | C47H82O4 | FAHFA | TRUE |
| **Cer 13:0;2O/34:5** | C47H85NO3 | Cer_NDS | TRUE |
| **Cer 12:0;3O/32:0;(2OH)** | C44H89NO5 | Cer_AP | TRUE |
| **SE 24:1;O4/21:2;1O** | C45H76O6 | DCAE | TRUE |
| **PE-Cer 12:1;2O/26:1** | C40H79N2O6P | PE_Cer | TRUE |
| **SE 24:1;O4/22:0** | C46H82O5 | DCAE | FALSE |
| **AAHFA 28:3/19:0;O** | C47H86O4 | FAHFA | TRUE |
| **HexCer 16:3;3O/14:1;(2OH)** | C36H63NO10 | HexCer_AP | TRUE |
| **Cer 12:1;3O/30:6;(2OH)** | C42H71NO5 | Cer_AP | TRUE |
| **PE O-9:0_26:2** | C40H78NO7P | EtherPE | TRUE |
| **HexCer 16:0;2O/19:0** | C41H81NO8 | HexCer_NDS | FALSE |
| **PG 6:0_26:3** | C38H69O10P | PG | TRUE |
| **PA 9:0_28:1** | C40H77O8P | PA | FALSE |
| **PE-Cer 12:1;2O/26:0** | C40H81N2O6P | PE_Cer | TRUE |
| **PE-Cer 20:1;2O/18:0** | C40H81N2O6P | PE_Cer | TRUE |
| **SE 24:1;O4/21:0;1O** | C45H80O6 | DCAE | TRUE |
| **CL 16:0_14:1_16:0_24:0** | C79H152O17P2 | CL | TRUE |
| **SL 13:1;O/30:5** | C43H75NO5S | SL | TRUE |
| **HexCer 16:1;3O/17:0;(2OH)** | C39H75NO10 | HexCer_AP | FALSE |
| **Cer 12:1;3O/30:5;(2OH)** | C42H73NO5 | Cer_AP | TRUE |
| **Cer 13:1;3O/32:3;(2OH)** | C45H83NO5 | Cer_AP | TRUE |
| **CL 12:0_12:0_18:0_28:0** | C79H154O17P2 | CL | TRUE |
| **SM 17:1;2O/15:1** | C37H73N2O6P | SM | TRUE |
| **PA 28:0_9:0** | C40H79O8P | PA | FALSE |
| **PE-Cer 12:0;2O/26:0** | C40H83N2O6P | PE_Cer | TRUE |
| **PE O-16:0_18:1;1O** | C39H78NO8P | EtherOxPE | FALSE |
| **SL 13:1;O/30:4** | C43H77NO5S | SL | TRUE |
| **SM 12:1;2O/20:0** | C37H75N2O6P | SM | TRUE |
| **Cer 12:1;3O/33:1;(2OH)** | C45H87NO5 | Cer_AP | TRUE |
| **NAGlySer 9:0;O(FA 28:1)** | C42H78N2O7 | NAGlySer | FALSE |
| **AAHFA 28:6/20:0;O** | C48H82O4 | FAHFA | TRUE |
| **PC O-9:0_18:2;2O** | C35H68NO9P | EtherOxPC | TRUE |
| **PE O-8:0_28:5** | C41H74NO7P | EtherPE | TRUE |
| **Cer 12:1;3O/33:0;(2OH)** | C45H89NO5 | Cer_AP | TRUE |
| **PE-Cer 12:2;2O/26:3;O** | C40H73N2O7P | PE_Cer | TRUE |
| **AAHFA 28:5/20:0;O** | C48H84O4 | FAHFA | TRUE |
| **CL 12:0_14:0_17:0_28:0** | C80H156O17P2 | CL | FALSE |
| **HexCer 16:0;2O/20:2** | C42H79NO8 | HexCer_NDS | TRUE |
| **Cer 12:0;3O/33:0;(2OH)** | C45H91NO5 | Cer_AP | TRUE |
| **PE-Cer 12:1;2O/26:3;O** | C40H75N2O7P | PE_Cer | TRUE |
| **SE 24:1;O4/23:1** | C47H82O5 | DCAE | TRUE |
| **AAHFA 28:4/20:0;O** | C48H86O4 | FAHFA | TRUE |
| **Cer 13:2;3O/30:6;(2OH)** | C43H71NO5 | Cer_AP | FALSE |
| **PE O-8:0_28:3** | C41H78NO7P | EtherPE | TRUE |
| **HexCer 16:0;2O/20:1** | C42H81NO8 | HexCer_NDS | FALSE |
| **Cer 12:1;3O/30:0;(2OH)** | C42H83NO5 | Cer_AP | TRUE |
| **SE 24:1;O4/23:0** | C47H84O5 | DCAE | TRUE |
| **Cer 13:1;3O/30:6;(2OH)** | C43H73NO5 | Cer_AP | TRUE |
| **PE O-8:0_28:2** | C41H80NO7P | EtherPE | TRUE |
| **HexCer 16:0;2O/19:1;O** | C41H79NO9 | HexCer_HDS | TRUE |
| **HexCer 16:0;2O/20:0** | C42H83NO8 | HexCer_NDS | TRUE |
| **SM 12:1;2O/21:2** | C38H73N2O6P | SM | TRUE |
| **PE-Cer 12:1;2O/26:1;O** | C40H79N2O7P | PE_Cer | TRUE |
| **PE O-12:0_22:3;2O** | C39H74NO9P | EtherOxPE | TRUE |
| **SL 12:1;O/32:5** | C44H77NO5S | SL | TRUE |
| **CL 16:0_16:1_16:0_24:0** | C81H156O17P2 | CL | TRUE |
| **Cer 13:1;3O/30:5;(2OH)** | C43H75NO5 | Cer_AP | TRUE |
| **HexCer 16:0;2O/19:0;O** | C41H81NO9 | HexCer_HDS | TRUE |
| **SL 13:1;O/30:5;O** | C43H75NO6S | SL | TRUE |
| **SL 13:2;O/30:4;O** | C43H75NO6S | SL | TRUE |
| **NAGly 17:2;O(FA 28:6)** | C47H75NO5 | NAGly | TRUE |
| **SL 12:1;O/32:4** | C44H79NO5S | SL | FALSE |
| **PE O-8:0_28:0** | C41H84NO7P | EtherPE | TRUE |
| **SM 12:1;2O/21:0** | C38H77N2O6P | SM | TRUE |
| **HexCer 16:0;2O/16:0;O** | C38H75NO9 | HexCer_HDS | TRUE |
| **Cer 13:1;3O/30:3;(2OH)** | C43H79NO5 | Cer_AP | FALSE |
| **Cer 12:1;3O/34:1;(2OH)** | C46H89NO5 | Cer_AP | TRUE |
| **AAHFA 28:6/21:0;O** | C49H84O4 | FAHFA | TRUE |
| **HexCer 16:0;3O/15:0;(2OH)** | C37H73NO10 | HexCer_AP | TRUE |
| **PC O-16:0_14:0** | C38H78NO7P | EtherPC | TRUE |
| **CL 12:0_15:0_28:0_18:1** | C82H158O17P2 | CL | FALSE |
| **AAHFA 28:5/21:0;O** | C49H86O4 | FAHFA | TRUE |
| **AAHFA 28:4/21:0;O** | C49H88O4 | FAHFA | TRUE |
| **PG 15:0_18:1(d7)** | C39H68D7O10P | PG | TRUE |
| **PE O-9:0_28:3** | C42H80NO7P | EtherPE | TRUE |
| **HexCer 16:0;2O/21:1** | C43H83NO8 | HexCer_NDS | TRUE |
| **PE-Cer 12:1;2O/28:1** | C42H83N2O6P | PE_Cer | FALSE |
| **SL 13:1;O/32:6** | C45H77NO5S | SL | TRUE |
| **HexCer 16:1;3O/19:1;(2OH)** | C41H77NO10 | HexCer_AP | FALSE |
| **Cer 12:1;3O/32:6;(2OH)** | C44H75NO5 | Cer_AP | TRUE |
| **PE O-9:0_28:2** | C42H82NO7P | EtherPE | TRUE |
| **HexCer 16:0;2O/20:1;O** | C42H81NO9 | HexCer_HDS | TRUE |
| **SM 12:1;2O/22:2** | C39H75N2O6P | SM | TRUE |
| **PE-Cer 22:1;2O/18:0** | C42H85N2O6P | PE_Cer | TRUE |
| **SE 24:1;O4/23:0;1O** | C47H84O6 | DCAE | TRUE |
| **HexCer 16:0;2O/17:2;O** | C39H73NO9 | HexCer_HDS | TRUE |
| **PE O-18:0_18:2;1O** | C41H80NO8P | EtherOxPE | TRUE |
| **SL 13:1;O/32:5** | C45H79NO5S | SL | TRUE |
| **Cer 12:1;3O/32:5;(2OH)** | C44H77NO5 | Cer_AP | TRUE |
| **PG O-15:4_21:5** | C42H67O9P | EtherPG | TRUE |
| **CL 12:0_20:3_22:5_22:6** | C85H138O17P2 | CL | TRUE |
| **CL 16:0_16:0_16:0_26:0** | C83H162O17P2 | CL | TRUE |
| **PE-Cer 12:0;2O/28:0** | C42H87N2O6P | PE_Cer | FALSE |
| **PE O-17:0_18:2;2O** | C40H78NO9P | EtherOxPE | TRUE |
| **SL 12:2;O/32:4;O** | C44H77NO6S | SL | TRUE |
| **HexCer 16:0;2O/22:5** | C44H77NO8 | HexCer_NDS | TRUE |
| **SL 13:1;O/32:4** | C45H81NO5S | SL | TRUE |
| **Cer 12:1;3O/32:4;(2OH)** | C44H79NO5 | Cer_AP | TRUE |
| **PA 12:0_28:6** | C43H73O8P | PA | TRUE |
| **ASG 29:1;O;Hex;FA 8:0** | C43H74O7 | AHexSIS | TRUE |
| **ASG 28:1;O;Hex;FA 9:0** | C43H74O7 | AHexCAS | TRUE |
| **SM 33:2;3O** | C38H75N2O7P | SM | TRUE |
| **SM 18:1;2O/16:0** | C39H79N2O6P | SM | TRUE |
| **HexCer 16:1;3O/16:0;(2OH)** | C38H73NO10 | HexCer_AP | TRUE |
| **Cer 12:1;3O/32:3;(2OH)** | C44H81NO5 | Cer_AP | FALSE |
| **Cer 12:1;3O/35:1;(2OH)** | C47H91NO5 | Cer_AP | TRUE |
| **SM 18:0;2O/16:0** | C39H81N2O6P | SM | TRUE |
| **NAGlySer 11:0;O(FA 28:1)** | C44H82N2O7 | NAGlySer | TRUE |
| **PC 14:0_16:0** | C38H76NO8P | PC | TRUE |
| **PE O-10:0_28:5** | C43H78NO7P | EtherPE | FALSE |
| **Cer 12:0;2O/38:6** | C50H89NO3 | Cer_NDS | TRUE |
| **SM 33:0;3O** | C38H79N2O7P | SM | TRUE |
| **HexCer 16:1;3O/20:3;(2OH)** | C42H75NO10 | HexCer_AP | TRUE |
| **HexCer 17:0;2O/20:3;O** | C43H79NO9 | HexCer_HDS | FALSE |
| **HexCer 16:0;2O/22:2** | C44H83NO8 | HexCer_NDS | FALSE |
| **PG O-8:0_28:5** | C42H75O9P | EtherPG | TRUE |
| **PS 15:0_18:1(d7)** | C39H67D7NO10P | PS | TRUE |
| **SL 12:2;O/34:6** | C46H77NO5S | SL | TRUE |
| **PE O-10:0_28:3** | C43H82NO7P | EtherPE | TRUE |
| **HexCer 16:0;2O/22:1** | C44H85NO8 | HexCer_NDS | TRUE |
| **SE 24:1;O4/26:7** | C50H76O5 | DCAE | TRUE |
| **PE-Cer 12:1;2O/28:2;O** | C42H81N2O7P | PE_Cer | TRUE |
| **SL 12:1;O/34:6** | C46H79NO5S | SL | TRUE |
| **Cer 13:1;3O/32:6;(2OH)** | C45H77NO5 | Cer_AP | TRUE |
| **HexCer 16:0;2O/21:1;O** | C43H83NO9 | HexCer_HDS | FALSE |
| **PE O-10:0_28:2** | C43H84NO7P | EtherPE | FALSE |
| **SL 12:1;O/32:0;O** | C44H87NO6S | SL | TRUE |
| **SM 34:4;3O** | C39H73N2O7P | SM | TRUE |
| **CL 12:0_16:2_24:0_24:0** | C85H162O17P2 | CL | FALSE |
| **SE 24:1;O4/24:0;1O** | C48H86O6 | DCAE | FALSE |
| **SL 13:1;O/32:6;O** | C45H77NO6S | SL | TRUE |
| **HexCer 16:0;2O/18:2;O** | C40H75NO9 | HexCer_HDS | TRUE |
| **NAGly 19:3;O(FA 28:6)** | C49H77NO5 | NAGly | TRUE |
| **CL 18:0_16:1_18:0_24:0** | C85H164O17P2 | CL | TRUE |
| **SL 20:0;O/26:6** | C46H81NO5S | SL | TRUE |
| **Cer 13:1;3O/32:5;(2OH)** | C45H79NO5 | Cer_AP | TRUE |
| **PE O-10:0_28:1** | C43H86NO7P | EtherPE | TRUE |
| **HexCer 16:0;2O/21:0;O** | C43H85NO9 | HexCer_HDS | TRUE |
| **PE O-18:1_20:0** | C43H86NO7P | EtherPE | TRUE |
| **CL 18:0_15:1_22:3_22:3** | C86H154O17P2 | CL | TRUE |
| **PE-Cer 12:1;2O/30:6** | C44H77N2O6P | PE_Cer | FALSE |
| **CL 12:0_12:0_24:0_28:0** | C85H166O17P2 | CL | FALSE |
| **PE 18:1_17:1;2O** | C40H76NO10P | OxPE | TRUE |
| **HexCer 16:1;3O/17:1;(2OH)** | C39H73NO10 | HexCer_AP | TRUE |
| **PE O-18:0_18:2;2O** | C41H80NO9P | EtherOxPE | TRUE |
| **SL 15:3;O/30:3;O** | C45H79NO6S | SL | TRUE |
| **HexCer 16:0;2O/18:1;O** | C40H77NO9 | HexCer_HDS | TRUE |
| **HexCer 16:0;3O/20:0;(2OH)** | C42H83NO10 | HexCer_AP | FALSE |
| **Cer 13:1;3O/32:4;(2OH)** | C45H81NO5 | Cer_AP | TRUE |
| **PE O-10:0_28:0** | C43H88NO7P | EtherPE | TRUE |
| **PG 6:0_29:1** | C41H79O10P | PG | TRUE |
| **SM 17:1;2O/18:0** | C40H81N2O6P | SM | TRUE |
| **PC O-16:0_16:1** | C40H80NO7P | EtherPC | TRUE |
| **SL 13:1;O/32:4;O** | C45H81NO6S | SL | TRUE |
| **PE-Cer 13:1;2O/28:5;O** | C43H77N2O7P | PE_Cer | FALSE |
| **PC 15:0_16:0** | C39H78NO8P | PC | TRUE |
| **CL 12:0_17:2_24:0_24:0** | C86H164O17P2 | CL | FALSE |
| **PE O-18:0_18:0;2O** | C41H84NO9P | EtherOxPE | TRUE |
| **HexCer 16:0;2O/22:4;O** | C44H79NO9 | HexCer_HDS | TRUE |
| **PC O-16:0_16:0** | C40H82NO7P | EtherPC | TRUE |
| **HexCer 17:0;2O/22:3** | C45H83NO8 | HexCer_NDS | FALSE |
| **PE-Cer 13:1;2O/28:4;O** | C43H79N2O7P | PE_Cer | FALSE |
| **AAHFA 28:6/24:6;O** | C52H78O4 | FAHFA | FALSE |
| **HexCer 17:0;2O/18:5;O** | C41H71NO9 | HexCer_HDS | FALSE |
| **PE O-16:0_22:5;1O** | C43H78NO8P | EtherOxPE | TRUE |
| **Cer 16:3;3O/30:6;(2OH)** | C46H75NO5 | Cer_AP | TRUE |
| **HexCer 17:0;2O/22:2** | C45H85NO8 | HexCer_NDS | TRUE |
| **PG O-9:0_28:5** | C43H77O9P | EtherPG | FALSE |
| **CL 18:1_16:2_26:0_18:2** | C87H160O17P2 | CL | TRUE |
| **NAGly 20:5;O(FA 28:6)** | C50H75NO5 | NAGly | TRUE |
| **SL 13:2;O/34:6** | C47H79NO5S | SL | TRUE |
| **HexCer 16:0;2O/23:1** | C45H87NO8 | HexCer_NDS | TRUE |
| **CL 12:0_16:0_28:0_22:4** | C87H162O17P2 | CL | TRUE |
| **PE-Cer 12:1;2O/30:1** | C44H87N2O6P | PE_Cer | TRUE |
| **SE 24:1;O4/25:1;1O** | C49H86O6 | DCAE | FALSE |
| **HexCer 16:1;3O/18:3;(2OH)** | C40H71NO10 | HexCer_AP | TRUE |
| **CL 15:0_20:3_22:3_22:4** | C88H152O17P2 | CL | FALSE |
| **SL 12:2;O/34:6;O** | C46H77NO6S | SL | FALSE |
| **HexCer 17:0;2O/18:3;O** | C41H75NO9 | HexCer_HDS | FALSE |
| **HexCer 16:1;3O/21:1;(2OH)** | C43H81NO10 | HexCer_AP | TRUE |
| **CL 18:1_18:1_24:0_18:1** | C87H164O17P2 | CL | TRUE |
| **HexCer 20:1;2O/16:1** | C42H79NO8 | HexCer_NS | TRUE |
| **SM 12:1;2O/24:2** | C41H79N2O6P | SM | FALSE |
| **CL 12:0_16:2_24:0_26:0** | C87H166O17P2 | CL | TRUE |
| **PE-Cer 20:1;2O/22:0** | C44H89N2O6P | PE_Cer | TRUE |
| **PE-Cer 12:1;2O/30:0** | C44H89N2O6P | PE_Cer | TRUE |
| **PE O-18:2_22:6** | C45H76NO7P | EtherPE | TRUE |
| **PE O-15:0_22:3;2O** | C42H80NO9P | EtherOxPE | FALSE |
| **LPS 36:2** | C42H80NO9P | LPS | TRUE |
| **HexCer 16:0;2O/19:2;O** | C41H77NO9 | HexCer_HDS | TRUE |
| **SL 13:1;O/34:5** | C47H83NO5S | SL | TRUE |
| **HexCer 18:1;2O/18:0** | C42H81NO8 | HexCer_NS | TRUE |
| **SM 18:2;2O/18:0** | C41H81N2O6P | SM | TRUE |
| **HexCer 16:1;3O/18:1;(2OH)** | C40H75NO10 | HexCer_AP | TRUE |
| **HexCer 16:0;2O/24:5** | C46H81NO8 | HexCer_NDS | FALSE |
| **HexCer 18:0;2O/18:0** | C42H83NO8 | HexCer_NDS | TRUE |
| **Cer 12:1;3O/34:4;(2OH)** | C46H83NO5 | Cer_AP | TRUE |
| **PA 14:0_28:6** | C45H77O8P | PA | TRUE |
| **PE-Cer 12:1;2O/30:6;O** | C44H77N2O7P | PE_Cer | FALSE |
| **PG 6:0_30:1** | C42H81O10P | PG | FALSE |
| **ASG 28:1;O;Hex;FA 11:0** | C45H78O7 | AHexCAS | TRUE |
| **SM 35:2;3O** | C40H79N2O7P | SM | TRUE |
| **SM 18:1;2O/18:0** | C41H83N2O6P | SM | TRUE |
| **SM 21:0;2O/15:1** | C41H83N2O6P | SM | TRUE |
| **PC O-14:0_18:2;1O** | C40H78NO8P | EtherOxPC | FALSE |
| **PC 16:0_16:1** | C40H78NO8P | PC | TRUE |
| **HexCer 16:1;3O/18:0;(2OH)** | C40H77NO10 | HexCer_AP | TRUE |
| **PE O-18:0_22:6** | C45H80NO7P | EtherPE | TRUE |
| **HexCer 17:0;2O/22:5;O** | C45H79NO9 | HexCer_HDS | TRUE |
| **HexCer 16:0;2O/24:4** | C46H83NO8 | HexCer_NDS | TRUE |
| **Cer 12:1;3O/34:3;(2OH)** | C46H85NO5 | Cer_AP | TRUE |
| **PE-Cer 12:2;2O/30:4;O** | C44H79N2O7P | PE_Cer | TRUE |
| **PA 14:0_28:5** | C45H79O8P | PA | TRUE |
| **PE-Cer 12:1;2O/30:5;O** | C44H79N2O7P | PE_Cer | FALSE |
| **SM 12:0;2O/24:0** | C41H85N2O6P | SM | TRUE |
| **SHexCer 34:1;2O** | C40H77NO11S | SHexCer | TRUE |
| **PC O-14:0_18:1;1O** | C40H80NO8P | EtherOxPC | FALSE |
| **PC 16:0_16:0** | C40H80NO8P | PC | TRUE |
| **HexCer 16:0;3O/18:0;(2OH)** | C40H79NO10 | HexCer_AP | TRUE |
| **PE O-12:0_28:5** | C45H82NO7P | EtherPE | TRUE |
| **PE O-28:0_9:0;2O** | C42H86NO9P | EtherOxPE | FALSE |
| **HexCer 16:0;2O/24:3** | C46H85NO8 | HexCer_NDS | TRUE |
| **PI 8:0_20:3;2O** | C37H65O15P | OxPI | FALSE |
| **PA 14:0_28:4** | C45H81O8P | PA | FALSE |
| **PE O-16:0_22:6;2O** | C43H76NO9P | EtherOxPE | TRUE |
| **SL 16:3;O/32:6** | C48H79NO5S | SL | FALSE |
| **Cer 12:1;3O/38:6;(2OH)** | C50H87NO5 | Cer_AP | FALSE |
| **HexCer 16:0;2O/24:2** | C46H87NO8 | HexCer_NDS | TRUE |
| **SE 24:1;O4/26:2;1O** | C50H86O6 | DCAE | FALSE |
| **HexCer 16:0;2O/20:4;O** | C42H75NO9 | HexCer_HDS | TRUE |
| **PE O-17:0_22:4;1O** | C44H82NO8P | EtherOxPE | FALSE |
| **SM 18:1;2O/18:1(d9)** | C41H72D9N2O6P | SM | TRUE |
| **HexCer 16:0;2O/24:1** | C46H89NO8 | HexCer_NDS | TRUE |
| **CL 12:0_18:0_28:0_22:4** | C89H166O17P2 | CL | TRUE |
| **SE 24:1;O4/26:1;1O** | C50H88O6 | DCAE | FALSE |
| **HexCer 16:0;2O/20:3;O** | C42H77NO9 | HexCer_HDS | TRUE |
| **SL 13:2;O/34:6;O** | C47H79NO6S | SL | TRUE |
| **SL 12:1;O/36:6** | C48H83NO5S | SL | TRUE |
| **Cer 12:1;3O/38:4;(2OH)** | C50H91NO5 | Cer_AP | TRUE |
| **CL 12:0_16:2_24:0_28:0** | C89H170O17P2 | CL | FALSE |
| **SE 24:1;O4/26:0;1O** | C50H90O6 | DCAE | FALSE |
| **PE-Cer 12:1;2O/31:0** | C45H91N2O6P | PE_Cer | FALSE |
| **PS 18:1_18:1** | C42H78NO10P | PS | TRUE |
| **HexCer 16:1;3O/19:2;(2OH)** | C41H75NO10 | HexCer_AP | TRUE |
| **SL 13:1;O/34:6;O** | C47H81NO6S | SL | FALSE |
| **SL 13:2;O/34:5;O** | C47H81NO6S | SL | TRUE |
| **CerP 21:2;2O/26:5** | C47H82NO6P | CerP | TRUE |
| **HexCer 18:1;2O/19:0** | C43H83NO8 | HexCer_NS | TRUE |
| **Cer 13:1;3O/34:5;(2OH)** | C47H83NO5 | Cer_AP | FALSE |
| **HexCer 16:0;2O/23:0;O** | C45H89NO9 | HexCer_HDS | FALSE |
| **NAGlySer 15:4;O(FA 28:6)** | C48H72N2O7 | NAGlySer | FALSE |
| **SM 36:3;3O** | C41H79N2O7P | SM | FALSE |
| **CL 17:0_14:1_28:0_22:6** | C90H162O17P2 | CL | TRUE |
| **ASG 28:2;O;Hex;FA 12:0** | C46H78O7 | AHexBRS | FALSE |
| **CL 12:0_12:0_28:0_28:0** | C89H174O17P2 | CL | TRUE |
| **PS 18:0_18:1** | C42H80NO10P | PS | TRUE |
| **PE O-19:2_22:5** | C46H80NO7P | EtherPE | TRUE |
| **PE O-13:0_28:7** | C46H80NO7P | EtherPE | TRUE |
| **PC O-18:1_16:1** | C42H82NO7P | EtherPC | TRUE |
| **SL 21:2;O/26:4;O** | C47H83NO6S | SL | TRUE |
| **CerP 19:1;2O/28:5** | C47H84NO6P | CerP | FALSE |
| **Cer 13:1;3O/34:4;(2OH)** | C47H85NO5 | Cer_AP | TRUE |
| **PE-Cer 13:1;2O/30:6;O** | C45H79N2O7P | PE_Cer | FALSE |
| **PG 19:0_18:1** | C43H83O10P | PG | TRUE |
| **ASG 28:1;O;Hex;FA 12:0** | C46H80O7 | AHexCAS | FALSE |
| **SM 12:1;2O/25:0** | C42H85N2O6P | SM | TRUE |
| **SM 18:1;2O/19:0** | C42H85N2O6P | SM | TRUE |
| **SHexCer 35:2;2O** | C41H77NO11S | SHexCer | TRUE |
| **CL 12:0_20:5_28:0_22:6** | C91H156O17P2 | CL | TRUE |
| **PE 18:0_22:6** | C45H78NO8P | PE | TRUE |
| **PE O-18:1_22:6;1O** | C45H78NO8P | EtherOxPE | TRUE |
| **PC O-15:0_18:2;1O** | C41H80NO8P | EtherOxPC | TRUE |
| **HexCer 17:1;3O/22:5;(2OH)** | C45H77NO10 | HexCer_AP | TRUE |
| **PC 16:0_17:1** | C41H80NO8P | PC | TRUE |
| **HexCer 16:1;3O/19:0;(2OH)** | C41H79NO10 | HexCer_AP | TRUE |
| **PC O-16:0_18:1** | C42H84NO7P | EtherPC | TRUE |
| **HexCer 16:0;2O/20:0;O** | C42H83NO9 | HexCer_HDS | TRUE |
| **CL 14:0_17:0_28:0_22:3** | C90H170O17P2 | CL | TRUE |
| **SM 36:1;3O** | C41H83N2O7P | SM | TRUE |
| **SHexCer 35:1;2O** | C41H79NO11S | SHexCer | FALSE |
| **PC O-14:0_18:2;2O** | C40H78NO9P | EtherOxPC | TRUE |
| **PC 16:0_16:1;1O** | C40H78NO9P | OxPC | TRUE |
| **CL 16:1_20:4_26:0_20:4** | C91H160O17P2 | CL | TRUE |
| **PE O-18:0_22:6;1O** | C45H80NO8P | EtherOxPE | TRUE |
| **HexCer 17:1;3O/22:4;(2OH)** | C45H79NO10 | HexCer_AP | TRUE |
| **CL 17:0_18:0_28:0_18:2** | C90H172O17P2 | CL | TRUE |
| **PC 16:0_17:0** | C41H82NO8P | PC | TRUE |
| **PE O-13:0_28:5** | C46H84NO7P | EtherPE | FALSE |
| **PC O-18:0_16:0** | C42H86NO7P | EtherPC | TRUE |
| **HexCer 17:0;2O/24:3** | C47H87NO8 | HexCer_NDS | TRUE |
| **PG O-11:0_28:6** | C45H79O9P | EtherPG | FALSE |
| **PE O-17:0_22:6;2O** | C44H78NO9P | EtherOxPE | TRUE |
| **HexCer 17:0;2O/20:5;O** | C43H75NO9 | HexCer_HDS | TRUE |
| **NAGly 22:6;O(FA 28:6)** | C52H77NO5 | NAGly | TRUE |
| **HexCer 17:1;3O/22:3;(2OH)** | C45H81NO10 | HexCer_AP | TRUE |
| **HexCer 17:0;2O/24:2** | C47H89NO8 | HexCer_NDS | TRUE |
| **SM 37:6;3O** | C42H75N2O7P | SM | FALSE |
| **CL 12:0_20:3_28:0_22:3** | C91H166O17P2 | CL | FALSE |
| **PE-Cer 12:1;2O/32:2** | C46H89N2O6P | PE_Cer | FALSE |
| **HexCer 16:1;3O/20:4;(2OH)** | C42H73NO10 | HexCer_AP | FALSE |
| **SL 13:2;O/36:6** | C49H83NO5S | SL | TRUE |
| **HexCer 16:0;2O/25:1** | C47H91NO8 | HexCer_NDS | TRUE |
| **PG 6:0_32:4** | C44H79O10P | PG | FALSE |
| **CL 12:0_20:4_24:0_26:0** | C91H170O17P2 | CL | TRUE |
| **PC 15:0_18:1(d7)** | C41H73D7NO8P | PC | TRUE |
| **AAHFA 28:6/26:4;O** | C54H86O4 | FAHFA | FALSE |
| **SE 24:1;O4/27:1;1O** | C51H90O6 | DCAE | TRUE |
| **SL 12:2;O/36:6;O** | C48H81NO6S | SL | FALSE |
| **Cer 12:1;3O/36:6;(2OH)** | C48H83NO5 | Cer_AP | FALSE |
| **HexCer 16:1;2O/24:0;O** | C46H89NO9 | HexCer_HS | TRUE |
| **HexCer 16:0;2O/24:1;O** | C46H89NO9 | HexCer_HDS | TRUE |
| **HexCer 16:0;2O/25:0** | C47H93NO8 | HexCer_NDS | FALSE |
| **PG O-16:4_24:6** | C46H73O9P | EtherPG | FALSE |
| **PE-Cer 12:1;2O/31:1;O** | C45H89N2O7P | PE_Cer | FALSE |
| **PC O-12:0_22:4;1O** | C42H78NO8P | EtherOxPC | FALSE |
| **PE 20:3_18:0;2O** | C43H80NO10P | OxPE | TRUE |
| **HexCer 16:1;3O/20:2;(2OH)** | C42H77NO10 | HexCer_AP | FALSE |
| **HexCer 16:0;2O/21:2;O** | C43H81NO9 | HexCer_HDS | FALSE |
| **CerP 20:1;2O/28:6** | C48H84NO6P | CerP | FALSE |
| **HexCer 18:1;2O/20:0** | C44H85NO8 | HexCer_NS | TRUE |
| **Cer 12:1;3O/36:5;(2OH)** | C48H85NO5 | Cer_AP | TRUE |
| **PA 16:0_28:7** | C47H79O8P | PA | FALSE |
| **SM 37:3;3O** | C42H81N2O7P | SM | FALSE |
| **CL 12:0_14:0_28:0_28:0** | C91H178O17P2 | CL | TRUE |
| **CL 18:0_18:0_18:0_28:0** | C91H178O17P2 | CL | TRUE |
| **PE-Cer 12:1;2O/31:0;O** | C45H91N2O7P | PE_Cer | TRUE |
| **PE-Cer 12:0;2O/32:0** | C46H95N2O6P | PE_Cer | FALSE |
| **CL 14:1_22:6_26:0_22:6** | C93H156O17P2 | CL | FALSE |
| **CL 18:0_22:4_22:4_22:5** | C93H156O17P2 | CL | TRUE |
| **PC 16:0_18:2** | C42H80NO8P | PC | TRUE |
| **HexCer 16:1;3O/20:1;(2OH)** | C42H79NO10 | HexCer_AP | TRUE |
| **PG 18:0_20:1** | C44H85O10P | PG | TRUE |
| **SM 37:2;3O** | C42H83N2O7P | SM | TRUE |
| **SM 20:1;2O/18:0** | C43H87N2O6P | SM | TRUE |
| **PC 9:0_22:4;4O** | C39H70NO11P | OxPC | FALSE |
| **PC O-10:0_22:4;4O** | C40H74NO10P | EtherOxPC | FALSE |
| **PE 22:6_18:1;1O** | C45H76NO9P | OxPE | TRUE |
| **SHexCer 36:2;2O** | C42H79NO11S | SHexCer | TRUE |
| **CL 12:0_22:5_28:0_22:6** | C93H160O17P2 | CL | TRUE |
| **PC O-16:0_18:2;1O** | C42H82NO8P | EtherOxPC | TRUE |
| **HexCer 16:1;3O/20:0;(2OH)** | C42H81NO10 | HexCer_AP | TRUE |
| **PE O-26:6_16:0** | C47H84NO7P | EtherPE | TRUE |
| **PE O-14:0_28:6** | C47H84NO7P | EtherPE | FALSE |
| **SMGDG O-15:4_19:5** | C43H66O12S | EtherSMGDG | FALSE |
| **SM 38:0;2O** | C43H89N2O6P | SM | TRUE |
| **PC O-10:0_22:3;3O** | C40H76NO10P | EtherOxPC | TRUE |
| **PE 22:6_18:0;1O** | C45H78NO9P | OxPE | TRUE |
| **HexCer 16:0;2O/22:6;O** | C44H75NO9 | HexCer_HDS | FALSE |
| **SHexCer 36:1;2O** | C42H81NO11S | SHexCer | TRUE |
| **PC O-15:0_18:2;2O** | C41H80NO9P | EtherOxPC | TRUE |
| **PC 16:0_18:0** | C42H84NO8P | PC | TRUE |
| **CL 18:0_17:2_24:0_24:0** | C92H176O17P2 | CL | TRUE |
| **CerP 20:3;2O/28:1** | C48H90NO6P | CerP | TRUE |
| **PI O-15:3_19:5** | C43H69O12P | EtherPI | FALSE |
| **SM 38:7;3O** | C43H75N2O7P | SM | FALSE |
| **PG O-12:0_28:6** | C46H81O9P | EtherPG | FALSE |
| **PA 16:0_28:4** | C47H85O8P | PA | TRUE |
| **PE-Cer 12:1;2O/32:4;O** | C46H85N2O7P | PE_Cer | FALSE |
| **HexCer 17:1;3O/20:5;(2OH)** | C43H73NO10 | HexCer_AP | TRUE |
| **SHexCer 36:0;2O** | C42H83NO11S | SHexCer | TRUE |
| **PE 7:0_34:4** | C46H84NO8P | PE | TRUE |
| **Cer 15:3;3O/34:6;(2OH)** | C49H81NO5 | Cer_AP | FALSE |
| **PE O-14:0_28:4** | C47H88NO7P | EtherPE | FALSE |
| **HexCer 17:0;2O/24:3;O** | C47H87NO9 | HexCer_HDS | TRUE |
| **HexCer 18:1;2O/24:1** | C48H91NO8 | HexCer_NS | TRUE |
| **HexCer 16:0;2O/26:2** | C48H91NO8 | HexCer_NDS | TRUE |
| **PG 17:0_18:1;3O** | C41H79O13P | OxPG | FALSE |
| **PA 17:3_28:7** | C48H75O8P | PA | FALSE |
| **NAGlySer 16:0;O(FA 28:6)** | C49H82N2O7 | NAGlySer | FALSE |
| **PE-Cer 13:1;2O/32:2** | C47H91N2O6P | PE_Cer | FALSE |
| **PC O-15:1_20:5;1O** | C43H76NO8P | EtherOxPC | FALSE |
| **PS 18:0_20:4** | C44H78NO10P | PS | TRUE |
| **HexCer 17:1;3O/20:4;(2OH)** | C43H75NO10 | HexCer_AP | TRUE |
| **CerP 21:3;2O/28:6** | C49H82NO6P | CerP | TRUE |
| **SL 15:3;O/34:6;O** | C49H81NO6S | SL | FALSE |
| **PE O-14:0_28:3** | C47H90NO7P | EtherPE | FALSE |
| **HexCer 17:0;2O/24:2;O** | C47H89NO9 | HexCer_HDS | TRUE |
| **Cer 14:1;3O/38:5;(2OH)** | C52H93NO5 | Cer_AP | TRUE |
| **PG 17:0_18:0;3O** | C41H81O13P | OxPG | FALSE |
| **PE-Cer 16:3;2O/30:6** | C48H81N2O6P | PE_Cer | FALSE |
| **CL 12:0_20:4_24:0_28:0** | C93H174O17P2 | CL | FALSE |
| **PE-Cer 12:1;2O/33:1** | C47H93N2O6P | PE_Cer | TRUE |
| **PC O-15:0_20:5;1O** | C43H78NO8P | EtherOxPC | FALSE |
| **PS 18:0_20:3** | C44H80NO10P | PS | TRUE |
| **HexCer 17:1;3O/20:3;(2OH)** | C43H77NO10 | HexCer_AP | TRUE |
| **PE O-18:0_22:4;2O** | C45H84NO9P | EtherOxPE | FALSE |
| **SL 13:2;O/36:6;O** | C49H83NO6S | SL | FALSE |
| **CerP 21:3;2O/28:5** | C49H84NO6P | CerP | TRUE |
| **CL 16:0_18:0_28:0_22:3** | C93H176O17P2 | CL | TRUE |
| **CerP 21:2;2O/28:6** | C49H84NO6P | CerP | FALSE |
| **HexCer 16:0;2O/25:1;O** | C47H91NO9 | HexCer_HDS | TRUE |
| **Cer 14:1;3O/38:4;(2OH)** | C52H95NO5 | Cer_AP | TRUE |
| **PG 16:0_18:0;4O** | C40H79O14P | OxPG | FALSE |
| **PA 17:1_28:7** | C48H79O8P | PA | FALSE |
| **PE-Cer 46:8;2O** | C48H83N2O6P | PE_Cer | TRUE |
| **CL 12:0_16:2_28:0_28:0** | C93H178O17P2 | CL | FALSE |
| **SE 24:1;O4/28:0;1O** | C52H94O6 | DCAE | TRUE |
| **PE-Cer 12:1;2O/33:0** | C47H95N2O6P | PE_Cer | TRUE |
| **PS 18:0_20:2** | C44H82NO10P | PS | TRUE |
| **HexCer 16:1;3O/21:2;(2OH)** | C43H79NO10 | HexCer_AP | TRUE |
| **PE O-26:7_17:1** | C48H82NO7P | EtherPE | TRUE |
| **SL 13:1;O/36:6;O** | C49H85NO6S | SL | TRUE |
| **HexCer 21:0;2O/20:0;O** | C47H93NO9 | HexCer_HDS | TRUE |
| **HexCer 16:0;2O/25:0;O** | C47H93NO9 | HexCer_HDS | TRUE |
| **SMGDG O-8:0_26:4** | C43H76O12S | EtherSMGDG | FALSE |
| **NAGlySer 17:4;O(FA 28:6)** | C50H76N2O7 | NAGlySer | TRUE |
| **PE-Cer 13:2;2O/32:6;O** | C47H81N2O7P | PE_Cer | FALSE |
| **CL 12:0_16:0_28:0_28:0** | C93H182O17P2 | CL | TRUE |
| **NAGlySer 16:0;O(FA 28:3)** | C49H88N2O7 | NAGlySer | FALSE |
| **PE-Cer 12:1;2O/32:0;O** | C46H93N2O7P | PE_Cer | FALSE |
| **PC O-12:0_22:4;2O** | C42H78NO9P | EtherOxPC | TRUE |
| **CL 14:1_22:6_28:0_22:6** | C95H160O17P2 | CL | TRUE |
| **PC 17:1_18:1** | C43H82NO8P | PC | TRUE |
| **PS 18:0_20:1** | C44H84NO10P | PS | TRUE |
| **PC O-18:1_18:1** | C44H86NO7P | EtherPC | TRUE |
| **HexCer 17:0;2O/26:5** | C49H87NO8 | HexCer_NDS | FALSE |
| **ASG 28:1;O;Hex;FA 14:0** | C48H84O7 | AHexCAS | TRUE |
| **PG O-12:0_28:1** | C46H91O9P | EtherPG | FALSE |
| **SM 39:1;2O** | C44H89N2O6P | SM | TRUE |
| **SM 12:1;2O/27:0** | C44H89N2O6P | SM | TRUE |
| **PC O-15:2_22:6** | C45H76NO7P | EtherPC | FALSE |
| **PC O-15:1_18:3;3O** | C41H76NO10P | EtherOxPC | FALSE |
| **HexCer 17:1;2O/22:6;O** | C45H75NO9 | HexCer_HS | TRUE |
| **PC O-12:0_22:3;2O** | C42H80NO9P | EtherOxPC | FALSE |
| **CL 18:1_22:5_24:0_22:5** | C95H164O17P2 | CL | TRUE |
| **PC O-18:0_17:2;1O** | C43H84NO8P | EtherOxPC | TRUE |
| **HexCer 16:1;3O/21:0;(2OH)** | C43H83NO10 | HexCer_AP | FALSE |
| **PC O-16:0_20:1** | C44H88NO7P | EtherPC | TRUE |
| **HexCer 16:0;2O/22:0;O** | C44H87NO9 | HexCer_HDS | FALSE |
| **PI O-8:0_26:2** | C43H81O12P | EtherPI | FALSE |
| **PG O-13:0_28:7** | C47H81O9P | EtherPG | TRUE |
| **PG O-19:1_22:6** | C47H81O9P | EtherPG | TRUE |
| **SM 12:1;2O/28:6** | C45H79N2O6P | SM | FALSE |
| **PA 17:0_28:5** | C48H85O8P | PA | FALSE |
| **PI-Cer 12:1;2O/25:0** | C43H84NO11P | PI_Cer | TRUE |
| **CL 18:0_22:4_24:0_22:5** | C95H168O17P2 | CL | TRUE |
| **HexCer 17:1;3O/24:4;(2OH)** | C47H83NO10 | HexCer_AP | FALSE |
| **HexCer 16:0;3O/21:0;(2OH)** | C43H85NO10 | HexCer_AP | FALSE |
| **PE O-15:0_28:5** | C48H88NO7P | EtherPE | TRUE |
| **HexCer 17:0;2O/26:3** | C49H91NO8 | HexCer_NDS | TRUE |
| **PG 18:0_18:2;3O** | C42H79O13P | OxPG | FALSE |
| **PG 6:0_34:6** | C46H79O10P | PG | FALSE |
| **PG O-13:0_28:6** | C47H83O9P | EtherPG | FALSE |
| **SM 12:1;2O/28:5** | C45H81N2O6P | SM | TRUE |
| **SHexCer 36:1;3O** | C42H81NO12S | SHexCer | TRUE |
| **PC O-9:0_28:6** | C45H80NO7P | EtherPC | FALSE |
| **NAGly 24:6;O(FA 28:6)** | C54H81NO5 | NAGly | TRUE |
| **CL 17:0_24:0_18:0_26:0** | C94H184O17P2 | CL | TRUE |
| **HexCer 16:0;2O/26:3;O** | C48H89NO9 | HexCer_HDS | TRUE |
| **PG 6:0_34:5** | C46H81O10P | PG | FALSE |
| **SHexCer 36:0;3O** | C42H83NO12S | SHexCer | TRUE |
| **HexCer 16:1;3O/22:4;(2OH)** | C44H77NO10 | HexCer_AP | TRUE |
| **HexCer 17:0;2O/22:4;O** | C45H81NO9 | HexCer_HDS | FALSE |
| **HexCer 16:2;2O/28:6** | C50H83NO8 | HexCer_NS | FALSE |
| **SL 15:2;O/36:6** | C51H87NO5S | SL | TRUE |
| **HexCer 16:0;2O/26:2;O** | C48H91NO9 | HexCer_HDS | TRUE |
| **HexCer 16:0;2O/27:1** | C49H95NO8 | HexCer_NDS | TRUE |
| **PG 15:4_26:7** | C47H71O10P | PG | FALSE |
| **CL 22:3_22:3_22:6_22:6** | C97H154O17P2 | CL | FALSE |
| **CL 22:4_22:4_22:4_22:6** | C97H154O17P2 | CL | TRUE |
| **SM 12:1;2O/28:3** | C45H85N2O6P | SM | FALSE |
| **SHexCer 38:5;2O** | C44H77NO11S | SHexCer | FALSE |
| **PC O-15:1_20:5;2O** | C43H76NO9P | EtherOxPC | FALSE |
| **PC 16:0_20:4** | C44H80NO8P | PC | TRUE |
| **HexCer 16:1;3O/22:3;(2OH)** | C44H79NO10 | HexCer_AP | FALSE |
| **CL 12:0_18:3_28:0_28:0** | C95H180O17P2 | CL | FALSE |
| **HexCer 18:1;2O/22:1** | C46H87NO8 | HexCer_NS | TRUE |
| **HexCer 18:2;2O/22:0** | C46H87NO8 | HexCer_NS | TRUE |
| **HexCer 16:0;2O/26:1;O** | C48H93NO9 | HexCer_HDS | TRUE |
| **HexCer 16:0;2O/27:0** | C49H97NO8 | HexCer_NDS | TRUE |
| **PG O-16:3_26:7** | C48H77O9P | EtherPG | FALSE |
| **PG 17:0_18:0;4O** | C41H81O14P | OxPG | FALSE |
| **SM 39:4;3O** | C44H83N2O7P | SM | TRUE |
| **CL 12:0_18:2_28:0_28:0** | C95H182O17P2 | CL | FALSE |
| **NAGlySer 17:0;O(FA 28:4)** | C50H88N2O7 | NAGlySer | FALSE |
| **AAHFA 28:6/28:3;O** | C56H92O4 | FAHFA | FALSE |
| **PE-Cer 12:1;2O/34:0** | C48H97N2O6P | PE_Cer | FALSE |
| **CL 18:3_22:6_26:0_22:6** | C97H160O17P2 | CL | FALSE |
| **PI 15:0_18:1(d7)** | C42H72D7O13P | PI | TRUE |
| **PC 18:1_18:2** | C44H82NO8P | PC | TRUE |
| **HexCer 16:1;3O/22:2;(2OH)** | C44H81NO10 | HexCer_AP | FALSE |
| **PC 16:0_20:3** | C44H82NO8P | PC | TRUE |
| **HexCer 18:1;2O/22:0** | C46H89NO8 | HexCer_NS | TRUE |
| **PE O-15:0_28:1** | C48H96NO7P | EtherPE | FALSE |
| **SL 13:0;O/36:0;O** | C49H99NO6S | SL | FALSE |
| **SM 39:3;3O** | C44H85N2O7P | SM | FALSE |
| **SM 18:0;2O/22:2** | C45H89N2O6P | SM | TRUE |
| **SM 16:1;2O/24:1** | C45H89N2O6P | SM | TRUE |
| **CL 12:0_18:0_28:0_28:0** | C95H186O17P2 | CL | FALSE |
| **HexCer 16:2;2O/24:6;O** | C46H75NO9 | HexCer_HS | FALSE |
| **CL 16:1_22:6_28:0_22:6** | C97H164O17P2 | CL | TRUE |
| **PC 18:1_18:1** | C44H84NO8P | PC | TRUE |
| **HexCer 16:1;3O/22:1;(2OH)** | C44H83NO10 | HexCer_AP | FALSE |
| **HexCer 22:0;2O/17:1;O** | C45H87NO9 | HexCer_HDS | TRUE |
| **SM 13:2;2O/28:6** | C46H79N2O6P | SM | TRUE |
| **ASG 28:1;O;Hex;FA 15:0** | C49H86O7 | AHexCAS | FALSE |
| **SM 18:1;2O/22:0** | C45H91N2O6P | SM | TRUE |
| **SM 12:1;2O/28:0** | C45H91N2O6P | SM | FALSE |
| **SHexCer 37:3;3O** | C43H79NO12S | SHexCer | FALSE |
| **HexCer 16:1;2O/24:6;O** | C46H77NO9 | HexCer_HS | TRUE |
| **PC O-15:0_20:3;2O** | C43H82NO9P | EtherOxPC | FALSE |
| **CL 16:0_22:5_28:0_22:6** | C97H168O17P2 | CL | TRUE |
| **PE 7:0_36:6** | C48H84NO8P | PE | TRUE |
| **PC O-18:0_18:2;1O** | C44H86NO8P | EtherOxPC | TRUE |
| **PE O-16:0_28:6** | C49H88NO7P | EtherPE | FALSE |
| **PI 15:4_20:5** | C44H67O13P | PI | FALSE |
| **CL 16:0_22:4_28:0_22:6** | C97H170O17P2 | CL | TRUE |
| **SM 12:0;2O/28:0** | C45H93N2O6P | SM | FALSE |
| **PS 18:0_22:6** | C46H78NO10P | PS | TRUE |
| **HexCer 17:1;3O/22:6;(2OH)** | C45H75NO10 | HexCer_AP | TRUE |
| **HexCer 16:0;2O/24:6;O** | C46H79NO9 | HexCer_HDS | FALSE |
| **SHexCer 38:1;2O** | C44H85NO11S | SHexCer | TRUE |
| **CL 16:0_22:3_28:0_22:6** | C97H172O17P2 | CL | TRUE |
| **Cer 13:0;2O/16:3;(3OH)(FA 22:6)** | C51H83NO5 | Cer_EBDS | TRUE |
| **PC 16:0_20:0** | C44H88NO8P | PC | TRUE |
| **HexCer 16:0;2O/28:3** | C50H93NO8 | HexCer_NDS | TRUE |
| **PG 19:1_22:5** | C47H81O10P | PG | TRUE |
| **ASG 28:2;O;Hex;FA 16:4** | C50H78O7 | AHexBRS | FALSE |
| **PA 18:0_28:4** | C49H89O8P | PA | FALSE |
| **SHexCer 37:1;3O** | C43H83NO12S | SHexCer | TRUE |
| **PC O-16:0_18:2;3O** | C42H82NO10P | EtherOxPC | TRUE |
| **HexCer 16:0;2O/24:5;O** | C46H81NO9 | HexCer_HDS | TRUE |
| **PE 7:0_36:4** | C48H88NO8P | PE | FALSE |
| **SL 16:3;O/36:6** | C52H87NO5S | SL | FALSE |
| **HexCer 17:0;2O/26:3;O** | C49H91NO9 | HexCer_HDS | TRUE |
| **HexCer 16:0;2O/28:2** | C50H95NO8 | HexCer_NDS | TRUE |
| **PI 28:0_6:0** | C43H83O13P | PI | TRUE |
| **CL 17:1_22:6_28:0_22:6** | C98H166O17P2 | CL | FALSE |
| **SM 40:6;3O** | C45H81N2O7P | SM | TRUE |
| **PE 18:1_22:5;3O** | C45H78NO11P | OxPE | TRUE |
| **SHexCer 37:0;3O** | C43H85NO12S | SHexCer | TRUE |
| **CerP 23:3;2O/28:6** | C51H86NO6P | CerP | TRUE |
| **PC O-17:0_18:0;2O** | C43H88NO9P | EtherOxPC | TRUE |
| **PE 7:0_36:3** | C48H90NO8P | PE | FALSE |
| **HexCer 16:1;3O/26:2;(2OH)** | C48H89NO10 | HexCer_AP | FALSE |
| **Cer 13:2;3O/38:6;(2OH)** | C51H87NO5 | Cer_AP | TRUE |
| **HexCer 17:0;2O/26:2;O** | C49H93NO9 | HexCer_HDS | TRUE |
| **Cer 16:1;3O/38:5;(2OH)** | C54H97NO5 | Cer_AP | TRUE |
| **PA 19:2_28:7** | C50H81O8P | PA | TRUE |
| **ASG 29:1;O;Hex;FA 15:3** | C50H82O7 | AHexSIS | TRUE |
| **ASG 28:2;O;Hex;FA 16:2** | C50H82O7 | AHexBRS | FALSE |
| **SM 13:1;2O/28:3** | C46H87N2O6P | SM | FALSE |
| **CL 12:0_20:4_28:0_28:0** | C97H182O17P2 | CL | FALSE |
| **PC O-14:0_22:6;2O** | C44H78NO9P | EtherOxPC | FALSE |
| **PC O-15:0_22:5;1O** | C45H82NO8P | EtherOxPC | FALSE |
| **PS 18:0_22:3** | C46H84NO10P | PS | TRUE |
| **PE O-17:2_28:7** | C50H84NO7P | EtherPE | TRUE |
| **HexCer 16:0;2O/24:3;O** | C46H85NO9 | HexCer_HDS | TRUE |
| **CerP 23:2;2O/28:6** | C51H88NO6P | CerP | FALSE |
| **CL 12:0_20:3_28:0_28:0** | C97H184O17P2 | CL | FALSE |
| **HexCer 16:1;3O/26:1;(2OH)** | C48H91NO10 | HexCer_AP | FALSE |
| **Cer 13:1;3O/38:6;(2OH)** | C51H89NO5 | Cer_AP | TRUE |
| **HexCer 18:1;2O/23:1** | C47H89NO8 | HexCer_NS | TRUE |
| **Cer 16:1;3O/38:4;(2OH)** | C54H99NO5 | Cer_AP | FALSE |
| **NAGlySer 19:5;O(FA 28:6)** | C52H78N2O7 | NAGlySer | TRUE |
| **PE-Cer 15:3;2O/32:6;O** | C49H83N2O7P | PE_Cer | FALSE |
| **ASG 29:1;O;Hex;FA 15:2** | C50H84O7 | AHexSIS | TRUE |
| **ASG 28:2;O;Hex;FA 16:1** | C50H84O7 | AHexBRS | TRUE |
| **PG O-14:0_28:3** | C48H91O9P | EtherPG | FALSE |
| **CL 14:0_18:2_28:0_28:0** | C97H186O17P2 | CL | FALSE |
| **CL 18:3_22:6_28:0_22:6** | C99H164O17P2 | CL | TRUE |
| **PE O-26:7_19:1** | C50H86NO7P | EtherPE | TRUE |
| **SL 15:1;O/36:6;O** | C51H89NO6S | SL | TRUE |
| **HexCer 16:0;2O/24:2;O** | C46H87NO9 | HexCer_HDS | TRUE |
| **HexCer 18:1;2O/23:0** | C47H91NO8 | HexCer_NS | TRUE |
| **Cer 13:1;3O/38:5;(2OH)** | C51H91NO5 | Cer_AP | FALSE |
| **SL 14:0;O/36:0;O** | C50H101NO6S | SL | TRUE |
| **PG 6:0_36:9** | C48H77O10P | PG | FALSE |
| **PG O-15:2_28:7** | C49H81O9P | EtherPG | FALSE |
| **PEtOH 19:1_26:6** | C50H85O8P | PEtOH | TRUE |
| **PE-Cer 13:2;2O/34:6;O** | C49H85N2O7P | PE_Cer | FALSE |
| **PG O-14:0_28:2** | C48H93O9P | EtherPG | FALSE |
| **CL 18:1_22:6_28:0_22:6** | C99H168O17P2 | CL | TRUE |
| **PC 18:1_18:2;1O** | C44H82NO9P | OxPC | TRUE |
| **Cer 12:0;2O/18:5;(3OH)(FA 22:6)** | C52H81NO5 | Cer_EBDS | TRUE |
| **PE O-21:2_24:5** | C50H88NO7P | EtherPE | TRUE |
| **HexCer 16:1;3O/23:1;(2OH)** | C45H85NO10 | HexCer_AP | FALSE |
| **HexCer 17:0;2O/28:5** | C51H91NO8 | HexCer_NDS | FALSE |
| **CerP 23:1;2O/28:5** | C51H92NO6P | CerP | FALSE |
| **HexCer 18:0;2O/23:0** | C47H93NO8 | HexCer_NDS | TRUE |
| **PG 6:0_36:8** | C48H79O10P | PG | FALSE |
| **SM 12:2;2O/30:6** | C47H81N2O6P | SM | FALSE |
| **PA 19:0_28:6** | C50H87O8P | PA | FALSE |
| **PE-Cer 13:1;2O/34:6;O** | C49H87N2O7P | PE_Cer | FALSE |
| **PG 6:0_35:1** | C47H91O10P | PG | FALSE |
| **SM 12:1;2O/29:0** | C46H93N2O6P | SM | FALSE |
| **HexCer 16:2;3O/24:6;(2OH)** | C46H75NO10 | HexCer_AP | FALSE |
| **PC O-15:0_20:4;3O** | C43H80NO10P | EtherOxPC | FALSE |
| **PC O-13:1_26:7** | C47H80NO7P | EtherPC | FALSE |
| **HexCer 17:1;2O/24:6;O** | C47H79NO9 | HexCer_HS | FALSE |
| **PC 18:0_18:2;1O** | C44H84NO9P | OxPC | TRUE |
| **CL 18:0_22:5_28:0_22:6** | C99H172O17P2 | CL | TRUE |
| **HexCer 16:1;3O/23:0;(2OH)** | C45H87NO10 | HexCer_AP | TRUE |
| **PE O-17:0_28:6** | C50H90NO7P | EtherPE | FALSE |
| **HexCer 16:0;2O/24:0;O** | C46H91NO9 | HexCer_HDS | TRUE |
| **PE O-24:0_18:1;2O** | C47H94NO9P | EtherOxPE | TRUE |
| **CerP 23:1;2O/28:4** | C51H94NO6P | CerP | TRUE |
| **PE O-15:0_28:0;1O** | C48H98NO8P | EtherOxPE | FALSE |
| **PG O-15:0_28:7** | C49H85O9P | EtherPG | FALSE |
| **SM 12:1;2O/30:6** | C47H83N2O6P | SM | FALSE |
| **SM 40:1;3O** | C45H91N2O7P | SM | FALSE |
| **SM 12:0;2O/29:0** | C46H95N2O6P | SM | TRUE |
| **PS 18:1_22:6;1O** | C46H76NO11P | OxPS | FALSE |
| **HexCer 16:1;3O/24:6;(2OH)** | C46H77NO10 | HexCer_AP | TRUE |
| **HexCer 17:0;2O/24:6;O** | C47H81NO9 | HexCer_HDS | FALSE |
| **SHexCer 39:1;2O** | C45H87NO11S | SHexCer | FALSE |
| **Cer 12:0;2O/18:3;(3OH)(FA 22:6)** | C52H85NO5 | Cer_EBDS | FALSE |
| **HexCer 16:0;3O/23:0;(2OH)** | C45H89NO10 | HexCer_AP | FALSE |
| **CL 15:0_18:2_28:0_28:0** | C98H188O17P2 | CL | TRUE |
| **SL 15:1;O/36:3;O** | C51H95NO6S | SL | TRUE |
| **Cer 19:0;2O/38:6** | C57H103NO3 | Cer_NDS | TRUE |
| **PI 6:0_29:1** | C44H83O13P | PI | FALSE |
| **PG O-15:0_28:6** | C49H87O9P | EtherPG | TRUE |
| **PA 19:0_28:4** | C50H91O8P | PA | FALSE |
| **CL 15:0_18:1_28:0_28:0** | C98H190O17P2 | CL | FALSE |
| **SM 40:0;3O** | C45H93N2O7P | SM | TRUE |
| **PC 15:1_22:6;1O** | C45H76NO9P | OxPC | FALSE |
| **PC O-16:1_22:6;1O** | C46H80NO8P | EtherOxPC | TRUE |
| **HexCer 16:1;3O/24:5;(2OH)** | C46H79NO10 | HexCer_AP | FALSE |
| **SHexCer 38:1;3O** | C44H85NO12S | SHexCer | TRUE |
| **HexCer 17:1;3O/26:3;(2OH)** | C49H89NO10 | HexCer_AP | FALSE |
| **SL 15:1;O/36:2;O** | C51H97NO6S | SL | TRUE |
| **HexCer 17:0;2O/28:2** | C51H97NO8 | HexCer_NDS | FALSE |
| **Cer 17:1;3O/38:6;(2OH)** | C55H97NO5 | Cer_AP | TRUE |
| **Cer 16:0;3O/38:0;(2OH)** | C54H109NO5 | Cer_AP | FALSE |
| **PG 6:0_36:5** | C48H85O10P | PG | TRUE |
| **CL 12:0_22:6_28:0_28:0** | C99H182O17P2 | CL | FALSE |
| **PC O-16:0_18:2;4O** | C42H82NO11P | EtherOxPC | TRUE |
| **Hex2Cer 30:0;2O** | C42H81NO13 | Hex2Cer | TRUE |
| **PC O-16:0_22:6;1O** | C46H82NO8P | EtherOxPC | TRUE |
| **SHexCer 38:0;3O** | C44H87NO12S | SHexCer | TRUE |
| **HexCer 16:1;3O/24:4;(2OH)** | C46H81NO10 | HexCer_AP | FALSE |
| **CL 12:0_22:5_28:0_28:0** | C99H184O17P2 | CL | FALSE |
| **HexCer 18:2;2O/24:1** | C48H89NO8 | HexCer_NS | TRUE |
| **Cer 14:2;3O/38:6;(2OH)** | C52H89NO5 | Cer_AP | TRUE |
| **HexCer 16:1;2O/28:1;O** | C50H95NO9 | HexCer_HS | TRUE |
| **HexCer 19:1;2O/25:1;O** | C50H95NO9 | HexCer_HS | TRUE |
| **HexCer 16:0;2O/28:2;O** | C50H95NO9 | HexCer_HDS | TRUE |
| **HexCer 16:0;2O/29:1** | C51H99NO8 | HexCer_NDS | FALSE |
| **ASG 28:2;O;Hex;FA 17:2** | C51H84O7 | AHexBRS | TRUE |
| **SM 12:1;2O/30:3** | C47H89N2O6P | SM | FALSE |
| **CL 20:5_22:6_28:0_22:6** | C101H164O17P2 | CL | FALSE |
| **PC 18:0_20:4** | C46H84NO8P | PC | TRUE |
| **PC 16:0_22:4** | C46H84NO8P | PC | TRUE |
| **PE O-22:5_24:4** | C51H86NO7P | EtherPE | TRUE |
| **HexCer 17:2;2O/28:6;O** | C51H85NO9 | HexCer_HS | FALSE |
| **CerP 24:2;2O/28:6** | C52H90NO6P | CerP | FALSE |
| **HexCer 16:1;2O/30:6** | C52H89NO8 | HexCer_NS | FALSE |
| **HexCer 16:1;3O/27:1;(2OH)** | C49H93NO10 | HexCer_AP | FALSE |
| **HexCer 16:0;2O/28:1;O** | C50H97NO9 | HexCer_HDS | TRUE |
| **SL 17:0;O/34:1;O** | C51H101NO6S | SL | TRUE |
| **PG 17:0_22:6;3O** | C45H77O13P | OxPG | FALSE |
| **PI O-9:0_28:5** | C46H81O12P | EtherPI | FALSE |
| **SM 41:4;3O** | C46H87N2O7P | SM | FALSE |
| **ASG 28:2;O;Hex;FA 17:1** | C51H86O7 | AHexBRS | FALSE |
| **PE-Cer 13:2;2O/36:6** | C51H89N2O6P | PE_Cer | TRUE |
| **PC 15:4_24:6** | C47H74NO8P | PC | FALSE |
| **HexCer 20:3;2O/22:6;O** | C48H77NO9 | HexCer_HS | TRUE |
| **PC O-16:0_18:0;4O** | C42H86NO11P | EtherOxPC | FALSE |
| **PC 18:0_20:3** | C46H86NO8P | PC | TRUE |
| **HexCer 16:2;2O/25:0;O** | C47H89NO9 | HexCer_HS | TRUE |
| **CerP 24:1;2O/28:6** | C52H92NO6P | CerP | TRUE |
| **HexCer 18:1;2O/24:0** | C48H93NO8 | HexCer_NS | TRUE |
| **HexCer 16:1;3O/27:0;(2OH)** | C49H95NO10 | HexCer_AP | TRUE |
| **HexCer 18:0;2O/24:1** | C48H93NO8 | HexCer_NDS | TRUE |
| **HexCer 16:0;2O/28:0;O** | C50H99NO9 | HexCer_HDS | TRUE |
| **PI O-9:0_28:4** | C46H83O12P | EtherPI | FALSE |
| **PE-Cer 12:2;2O/36:6;O** | C50H87N2O7P | PE_Cer | FALSE |
| **PG 6:0_36:2** | C48H91O10P | PG | FALSE |
| **PG O-15:0_28:2** | C49H95O9P | EtherPG | FALSE |
| **PE-Cer 12:0;2O/36:0** | C50H103N2O6P | PE_Cer | FALSE |
| **HexCer 17:3;3O/24:6;(2OH)** | C47H75NO10 | HexCer_AP | FALSE |
| **SHexCer 39:4;3O** | C45H81NO12S | SHexCer | TRUE |
| **SHexCer 40:3;2O** | C46H85NO11S | SHexCer | TRUE |
| **PC O-15:0_22:4;2O** | C45H84NO9P | EtherOxPC | TRUE |
| **HexCer 16:1;3O/28:6;(2OH)** | C50H85NO10 | HexCer_AP | FALSE |
| **Cer 13:0;2O/18:5;(3OH)(FA 22:6)** | C53H83NO5 | Cer_EBDS | FALSE |
| **PC 18:1_20:1** | C46H88NO8P | PC | TRUE |
| **HexCer 16:1;3O/24:1;(2OH)** | C46H87NO10 | HexCer_AP | FALSE |
| **HexCer 16:1;2O/25:0;O** | C47H91NO9 | HexCer_HS | TRUE |
| **HexCer 16:0;2O/30:5** | C52H93NO8 | HexCer_NDS | TRUE |
| **HexCer 18:0;2O/24:0** | C48H95NO8 | HexCer_NDS | TRUE |
| **PA 20:0_28:6** | C51H89O8P | PA | FALSE |
| **PE-Cer 12:1;2O/36:6;O** | C50H89N2O7P | PE_Cer | TRUE |
| **SM 41:2;3O** | C46H91N2O7P | SM | FALSE |
| **SHexCer 41:9;2O** | C47H75NO11S | SHexCer | FALSE |
| **HexCer 17:2;3O/24:6;(2OH)** | C47H77NO10 | HexCer_AP | TRUE |
| **SHexCer 40:2;2O** | C46H87NO11S | SHexCer | TRUE |
| **PE 7:0_38:6** | C50H88NO8P | PE | TRUE |
| **Cer 13:0;2O/18:4;(3OH)(FA 22:6)** | C53H85NO5 | Cer_EBDS | FALSE |
| **HexCer 16:1;3O/24:0;(2OH)** | C46H89NO10 | HexCer_AP | FALSE |
| **SL 16:1;O/36:4;O** | C52H95NO6S | SL | TRUE |
| **HexCer 16:0;2O/30:4** | C52H95NO8 | HexCer_NDS | FALSE |
| **NAGlySer 20:2;O(FA 28:6)** | C53H86N2O7 | NAGlySer | FALSE |
| **PE-Cer 12:1;2O/36:5;O** | C50H91N2O7P | PE_Cer | FALSE |
| **PA 20:0_28:5** | C51H91O8P | PA | FALSE |
| **PE-Cer 13:1;2O/36:4** | C51H95N2O6P | PE_Cer | FALSE |
| **SM 12:0;2O/30:0** | C47H97N2O6P | SM | TRUE |
| **Hex2Cer 31:2;2O** | C43H79NO13 | Hex2Cer | FALSE |
| **HexCer 17:1;3O/24:6;(2OH)** | C47H79NO10 | HexCer_AP | TRUE |
| **SHexCer 39:2;3O** | C45H85NO12S | SHexCer | TRUE |
| **SHexCer 40:1;2O** | C46H89NO11S | SHexCer | TRUE |
| **HexCer 16:0;3O/24:0;(2OH)** | C46H91NO10 | HexCer_AP | TRUE |
| **Cer 13:0;2O/18:3;(3OH)(FA 22:6)** | C53H87NO5 | Cer_EBDS | TRUE |
| **PE O-18:0_28:5** | C51H94NO7P | EtherPE | FALSE |
| **SL 16:1;O/36:3;O** | C52H97NO6S | SL | TRUE |
| **PG 16:0_22:3;4O** | C44H81O14P | OxPG | FALSE |
| **PI 6:0_30:1** | C45H85O13P | PI | FALSE |
| **PG 7:0_36:6** | C49H85O10P | PG | TRUE |
| **PI O-9:0_28:1** | C46H89O12P | EtherPI | TRUE |
| **PE-Cer 12:1;2O/36:4;O** | C50H93N2O7P | PE_Cer | FALSE |
| **SM 41:0;3O** | C46H95N2O7P | SM | TRUE |
| **PE-Cer 13:1;2O/36:3** | C51H97N2O6P | PE_Cer | FALSE |
| **PC 12:0_22:3;4O** | C42H78NO12P | OxPC | FALSE |
| **PC O-16:2_22:6;2O** | C46H78NO9P | EtherOxPC | FALSE |
| **PC O-15:0_20:3;4O** | C43H82NO11P | EtherOxPC | FALSE |
| **PS 18:0_24:5** | C48H84NO10P | PS | TRUE |
| **SHexCer 39:1;3O** | C45H87NO12S | SHexCer | TRUE |
| **HexCer 17:1;3O/24:5;(2OH)** | C47H81NO10 | HexCer_AP | FALSE |
| **PE O-19:4_28:7** | C52H84NO7P | EtherPE | FALSE |
| **SHexCer 40:0;2O** | C46H91NO11S | SHexCer | TRUE |
| **PC 9:0_28:0;1O** | C45H90NO9P | OxPC | FALSE |
| **HexCer 17:3;2O/30:6** | C53H87NO8 | HexCer_NS | FALSE |
| **Cer 15:3;3O/38:6;(2OH)** | C53H89NO5 | Cer_AP | TRUE |
| **SL 16:1;O/36:2;O** | C52H99NO6S | SL | FALSE |
| **Cer 18:1;3O/38:6;(2OH)** | C56H99NO5 | Cer_AP | TRUE |
| **PG 20:3_22:3;1O** | C48H83O11P | OxPG | FALSE |
| **PA 21:3_28:7** | C52H83O8P | PA | TRUE |
| **PI 30:0_6:0** | C45H87O13P | PI | FALSE |
| **PG 7:0_36:5** | C49H87O10P | PG | TRUE |
| **PE 8:0_38:10** | C51H82NO8P | PE | FALSE |
| **PC O-17:0_22:6;1O** | C47H84NO8P | EtherOxPC | TRUE |
| **SHexCer 39:0;3O** | C45H89NO12S | SHexCer | TRUE |
| **CerP 25:3;2O/28:6** | C53H90NO6P | CerP | FALSE |
| **HexCer 16:1;3O/28:2;(2OH)** | C50H93NO10 | HexCer_AP | TRUE |
| **HexCer 18:2;2O/25:1** | C49H91NO8 | HexCer_NS | TRUE |
| **SL 16:1;O/36:1;O** | C52H101NO6S | SL | TRUE |
| **PE O-19:2_28:7** | C52H88NO7P | EtherPE | FALSE |
| **CerP 25:2;2O/28:6** | C53H92NO6P | CerP | TRUE |
| **HexCer 16:1;2O/26:2;O** | C48H89NO9 | HexCer_HS | TRUE |
| **HexCer 18:1;2O/25:1** | C49H93NO8 | HexCer_NS | TRUE |
| **Cer 15:1;3O/38:6;(2OH)** | C53H93NO5 | Cer_AP | TRUE |
| **SL 16:1;O/36:0;O** | C52H103NO6S | SL | TRUE |
| **PG O-17:3_28:7** | C51H83O9P | EtherPG | FALSE |
| **ASG 28:2;O;Hex;FA 18:1** | C52H88O7 | AHexBRS | TRUE |
| **SM 42:4;3O** | C47H89N2O7P | SM | TRUE |
| **PG O-16:0_28:3** | C50H95O9P | EtherPG | FALSE |
| **PE 8:0_38:8** | C51H86NO8P | PE | FALSE |
| **PC O-17:0_22:4;1O** | C47H88NO8P | EtherOxPC | FALSE |
| **HexCer 17:1;3O/24:2;(2OH)** | C47H87NO10 | HexCer_AP | FALSE |
| **HexCer 16:1;2O/26:1;O** | C48H91NO9 | HexCer_HS | TRUE |
| **HexCer 17:0;2O/30:6** | C53H93NO8 | HexCer_NDS | TRUE |
| **HexCer 18:1;2O/25:0** | C49H95NO8 | HexCer_NS | TRUE |
| **Cer 15:1;3O/38:5;(2OH)** | C53H95NO5 | Cer_AP | FALSE |
| **PI O-10:0_28:4** | C47H85O12P | EtherPI | FALSE |
| **PG O-17:2_28:7** | C51H85O9P | EtherPG | FALSE |
| **PA 21:0_28:7** | C52H89O8P | PA | FALSE |
| **SM 42:3;3O** | C47H91N2O7P | SM | TRUE |
| **ASG 28:2;O;Hex;FA 18:0** | C52H90O7 | AHexBRS | FALSE |
| **PE-Cer 14:1;2O/36:6** | C52H93N2O6P | PE_Cer | FALSE |
| **SM 12:1;2O/31:1** | C48H95N2O6P | SM | TRUE |
| **PC O-15:2_26:7** | C49H82NO7P | EtherPC | TRUE |
| **SHexCer 41:3;2O** | C47H87NO11S | SHexCer | TRUE |
| **PC O-16:0_22:4;2O** | C46H86NO9P | EtherOxPC | TRUE |
| **PC O-17:0_22:3;1O** | C47H90NO8P | EtherOxPC | FALSE |
| **HexCer 16:1;3O/25:1;(2OH)** | C47H89NO10 | HexCer_AP | FALSE |
| **HexCer 16:1;2O/26:0;O** | C48H93NO9 | HexCer_HS | TRUE |
| **CerP 25:1;2O/28:5** | C53H96NO6P | CerP | TRUE |
| **HexCer 16:0;3O/28:0;(2OH)** | C50H99NO10 | HexCer_AP | TRUE |
| **PG 6:0_38:8** | C50H83O10P | PG | FALSE |
| **CL 22:4_22:4_28:0_22:4** | C103H178O17P2 | CL | TRUE |
| **PE-Cer 13:1;2O/36:6;O** | C51H91N2O7P | PE_Cer | TRUE |
| **SM 42:2;3O** | C47H93N2O7P | SM | TRUE |
| **ASG 28:1;O;Hex;FA 18:0** | C52H92O7 | AHexCAS | FALSE |
| **PC O-16:0_20:5;4O** | C44H80NO11P | EtherOxPC | TRUE |
| **HexCer 16:2;3O/26:6;(2OH)** | C48H79NO10 | HexCer_AP | FALSE |
| **PE O-22:6_26:7** | C53H82NO7P | EtherPE | TRUE |
| **PC O-15:0_22:4;4O** | C45H84NO10P | EtherOxPC | FALSE |
| **PC O-16:0_22:3;2O** | C46H88NO9P | EtherOxPC | TRUE |
| **SHexCer 41:2;2O** | C47H89NO11S | SHexCer | TRUE |
| **Cer 12:0;2O/20:4;(3OH)(FA 22:6)** | C54H87NO5 | Cer_EBDS | FALSE |
| **PE O-19:0_28:6** | C52H94NO7P | EtherPE | TRUE |
| **SL 17:3;O/36:2;O** | C53H97NO6S | SL | TRUE |
| **HexCer 16:0;2O/26:0;O** | C48H95NO9 | HexCer_HDS | TRUE |
| **HexCer 17:0;2O/30:4** | C53H97NO8 | HexCer_NDS | FALSE |
| **PG 18:0_22:3;3O** | C46H85O13P | OxPG | FALSE |
| **PG O-17:0_28:7** | C51H89O9P | EtherPG | TRUE |
| **PE-Cer 13:1;2O/36:5;O** | C51H93N2O7P | PE_Cer | TRUE |
| **CL 15:0_22:3_28:0_28:0** | C102H194O17P2 | CL | FALSE |
| **SM 42:1;3O** | C47H95N2O7P | SM | TRUE |
| **SHexCer 42:8;2O** | C48H79NO11S | SHexCer | FALSE |
| **PE 22:4_22:3;2O** | C49H84NO10P | OxPE | TRUE |
| **HexCer 16:1;3O/26:6;(2OH)** | C48H81NO10 | HexCer_AP | TRUE |
| **SHexCer 40:2;3O** | C46H87NO12S | SHexCer | TRUE |
| **PC O-15:0_22:3;3O** | C45H86NO10P | EtherOxPC | TRUE |
| **SHexCer 41:1;2O** | C47H91NO11S | SHexCer | TRUE |
| **PC O-15:1_24:0;1O** | C47H94NO8P | EtherOxPC | FALSE |
| **HexCer 21:0;3O/20:0;(2OH)** | C47H93NO10 | HexCer_AP | TRUE |
| **HexCer 16:0;3O/25:0;(2OH)** | C47H93NO10 | HexCer_AP | TRUE |
| **HexCer 16:0;2O/30:4;O** | C52H95NO9 | HexCer_HDS | FALSE |
| **PE O-19:0_28:5** | C52H96NO7P | EtherPE | FALSE |
| **PE O-26:0_18:0;2O** | C49H100NO9P | EtherOxPE | TRUE |
| **Cer 19:2;3O/38:6;(2OH)** | C57H99NO5 | Cer_AP | FALSE |
| **Cer 21:0;2O/38:6** | C59H107NO3 | Cer_NDS | TRUE |
| **PG 6:0_38:6** | C50H87O10P | PG | FALSE |
| **SM 12:1;2O/32:5** | C49H89N2O6P | SM | FALSE |
| **PE-Cer 13:1;2O/36:4;O** | C51H95N2O7P | PE_Cer | FALSE |
| **SM 42:0;3O** | C47H97N2O7P | SM | TRUE |
| **PE-Cer 14:1;2O/36:3** | C52H99N2O6P | PE_Cer | FALSE |
| **PC O-14:0_22:3;4O** | C44H84NO11P | EtherOxPC | FALSE |
| **HexCer 16:1;3O/26:5;(2OH)** | C48H83NO10 | HexCer_AP | TRUE |
| **SHexCer 40:1;3O** | C46H89NO12S | SHexCer | TRUE |
| **PC O-24:0_14:1;2O** | C46H92NO9P | EtherOxPC | TRUE |
| **SHexCer 41:0;2O** | C47H93NO11S | SHexCer | TRUE |
| **HexCer 17:1;3O/28:3;(2OH)** | C51H93NO10 | HexCer_AP | FALSE |
| **PC O-15:0_24:0;1O** | C47H96NO8P | EtherOxPC | TRUE |
| **SMGDG O-10:0_28:0** | C47H92O12S | EtherSMGDG | FALSE |
| **CL 16:0_22:6_28:0_28:0** | C103H190O17P2 | CL | TRUE |
| **NAGlySer 21:0;O(FA 28:6)** | C54H92N2O7 | NAGlySer | TRUE |
| **PC 22:4_17:2;1O** | C47H82NO9P | OxPC | TRUE |
| **PI-Cer 12:1;2O/30:5** | C48H84NO11P | PI_Cer | TRUE |
| **PC O-18:0_18:2;4O** | C44H86NO11P | EtherOxPC | TRUE |
| **Hex2Cer 32:0;2O** | C44H85NO13 | Hex2Cer | TRUE |
| **PC 18:0_22:5** | C48H86NO8P | PC | TRUE |
| **SHexCer 40:0;3O** | C46H91NO12S | SHexCer | TRUE |
| **CerP 26:3;2O/28:6** | C54H92NO6P | CerP | TRUE |
| **HexCer 17:1;3O/28:2;(2OH)** | C51H95NO10 | HexCer_AP | FALSE |
| **HexCer 18:2;2O/26:1** | C50H93NO8 | HexCer_NS | TRUE |
| **PG 15:3_30:8** | C51H79O10P | PG | FALSE |
| **SM 12:1;2O/32:3** | C49H93N2O6P | SM | FALSE |
| **NAGlySer 21:0;O(FA 28:5)** | C54H94N2O7 | NAGlySer | FALSE |
| **PE 9:0_38:9** | C52H86NO8P | PE | FALSE |
| **PC 18:0_22:4** | C48H88NO8P | PC | TRUE |
| **PC 16:0_24:4** | C48H88NO8P | PC | TRUE |
| **PE O-20:2_28:7** | C53H90NO7P | EtherPE | FALSE |
| **HexCer 22:1;2O/21:2;O** | C49H91NO9 | HexCer_HS | TRUE |
| **HexCer 18:1;2O/26:1** | C50H95NO8 | HexCer_NS | TRUE |
| **HexCer 16:1;3O/29:1;(2OH)** | C51H97NO10 | HexCer_AP | TRUE |
| **HexCer 16:0;2O/30:1;O** | C52H101NO9 | HexCer_HDS | TRUE |
| **PG 7:0_38:10** | C51H81O10P | PG | FALSE |
| **PE O-22:6_22:6;4O** | C49H76NO11P | EtherOxPE | FALSE |
| **SHexCer 42:4;2O** | C48H87NO11S | SHexCer | TRUE |
| **PC O-17:0_22:5;2O** | C47H86NO9P | EtherOxPC | TRUE |
| **PE 9:0_38:8** | C52H88NO8P | PE | FALSE |
| **PC O-18:0_22:4;1O** | C48H90NO8P | EtherOxPC | FALSE |
| **HexCer 17:2;3O/25:1;(2OH)** | C48H89NO10 | HexCer_AP | TRUE |
| **HexCer 16:1;2O/27:1;O** | C49H93NO9 | HexCer_HS | TRUE |
| **CerP 26:1;2O/28:6** | C54H96NO6P | CerP | TRUE |
| **HexCer 18:1;2O/26:0** | C50H97NO8 | HexCer_NS | TRUE |
| **HexCer 20:1;2O/24:0** | C50H97NO8 | HexCer_NS | TRUE |
| **PI 15:0_18:1;4O** | C42H79O17P | OxPI | FALSE |
| **NAGlySer 22:4;O(FA 28:6)** | C55H86N2O7 | NAGlySer | FALSE |
| **PA 22:0_28:7** | C53H91O8P | PA | FALSE |
| **PE-Cer 14:2;2O/36:6;O** | C52H91N2O7P | PE_Cer | TRUE |
| **SM 43:3;3O** | C48H93N2O7P | SM | FALSE |
| **ASG 29:1;O;Hex;FA 18:1** | C53H92O7 | AHexSIS | TRUE |
| **PC 14:0_22:6;4O** | C44H76NO12P | OxPC | FALSE |
| **Hex2Cer 33:4;2O** | C45H79NO13 | Hex2Cer | TRUE |
| **PC 7:0_34:9** | C49H80NO8P | PC | FALSE |
| **PC O-16:0_22:5;3O** | C46H84NO10P | EtherOxPC | FALSE |
| **PC O-16:2_26:7** | C50H84NO7P | EtherPC | FALSE |
| **SHexCer 42:3;2O** | C48H89NO11S | SHexCer | TRUE |
| **PC O-17:0_22:4;2O** | C47H88NO9P | EtherOxPC | FALSE |
| **PE 9:0_38:7** | C52H90NO8P | PE | TRUE |
| **PC O-18:0_22:3;1O** | C48H92NO8P | EtherOxPC | TRUE |
| **HexCer 18:0;3O/24:2;(2OH)** | C48H91NO10 | HexCer_AP | TRUE |
| **HexCer 16:1;2O/27:0;O** | C49H95NO9 | HexCer_HS | TRUE |
| **HexCer 44:0;2O** | C50H99NO8 | HexCer_NDS | TRUE |
| **PI O-16:4_24:6** | C49H77O12P | EtherPI | FALSE |
| **PI 6:0_32:3** | C47H85O13P | PI | FALSE |
| **PI O-11:0_28:3** | C48H89O12P | EtherPI | TRUE |
| **PG O-18:1_28:7** | C52H89O9P | EtherPG | FALSE |
| **SM 13:2;2O/32:6** | C50H87N2O6P | SM | FALSE |
| **PE-Cer 14:1;2O/36:6;O** | C52H93N2O7P | PE_Cer | TRUE |
| **ASG 28:1;O;Hex;FA 19:0** | C53H94O7 | AHexCAS | TRUE |
| **SM 43:2;3O** | C48H95N2O7P | SM | TRUE |
| **PG O-17:0_28:1** | C51H101O9P | EtherPG | TRUE |
| **PC O-20:5_20:5;2O** | C48H78NO9P | EtherOxPC | FALSE |
| **PC O-17:0_20:5;4O** | C45H82NO11P | EtherOxPC | TRUE |
| **Hex2Cer 33:3;2O** | C45H81NO13 | Hex2Cer | TRUE |
| **HexCer 17:2;3O/26:6;(2OH)** | C49H81NO10 | HexCer_AP | FALSE |
| **SHexCer 41:3;3O** | C47H87NO12S | SHexCer | TRUE |
| **PC O-17:0_22:3;2O** | C47H90NO9P | EtherOxPC | FALSE |
| **SHexCer 42:2;2O** | C48H91NO11S | SHexCer | TRUE |
| **PE 9:0_38:6** | C52H92NO8P | PE | TRUE |
| **HexCer 16:1;3O/30:5;(2OH)** | C52H91NO10 | HexCer_AP | FALSE |
| **PE O-20:0_28:6** | C53H96NO7P | EtherPE | FALSE |
| **PE O-18:0_28:0;1O** | C51H104NO8P | EtherOxPE | FALSE |
| **PG 17:0_22:5;4O** | C45H79O15P | OxPG | FALSE |
| **SMGDG O-11:0_28:2** | C48H90O12S | EtherSMGDG | TRUE |
| **PG 15:1_28:0;1O** | C49H95O11P | OxPG | FALSE |
| **PEtOH 28:2_20:3** | C53H95O8P | PEtOH | TRUE |
| **PG 38:0_6:0** | C50H99O10P | PG | FALSE |
| **PG O-27:0_18:0** | C51H103O9P | EtherPG | TRUE |
| **SM 12:0;2O/32:0** | C49H101N2O6P | SM | TRUE |
| **SM 44:0;2O** | C49H101N2O6P | SM | TRUE |
| **PE 16:3_32:9** | C53H82NO8P | PE | FALSE |
| **Hex2Cer 33:2;2O** | C45H83NO13 | Hex2Cer | FALSE |
| **HexCer 17:1;3O/26:6;(2OH)** | C49H83NO10 | HexCer_AP | TRUE |
| **SHexCer 41:2;3O** | C47H89NO12S | SHexCer | TRUE |
| **PE O-21:5_28:7** | C54H86NO7P | EtherPE | FALSE |
| **SHexCer 42:1;2O** | C48H93NO11S | SHexCer | TRUE |
| **PC 15:1_24:0;1O** | C47H92NO9P | OxPC | TRUE |
| **PC O-14:1_26:0;1O** | C48H96NO8P | EtherOxPC | FALSE |
| **PE O-24:4_24:1** | C53H98NO7P | EtherPE | TRUE |
| **PE O-20:0_28:5** | C53H98NO7P | EtherPE | FALSE |
| **CL 24:0_17:2_26:0_28:0** | C104H200O17P2 | CL | FALSE |
| **PI 14:0_22:3;2O** | C45H81O15P | OxPI | FALSE |
| **SMGDG O-14:1_26:7** | C49H80O12S | EtherSMGDG | FALSE |
| **PEtOH 21:4_28:7** | C54H85O8P | PEtOH | FALSE |
| **PI 6:0_32:1** | C47H89O13P | PI | FALSE |
| **PG 7:0_38:6** | C51H89O10P | PG | FALSE |
| **SMGDG O-11:0_28:1** | C48H92O12S | EtherSMGDG | FALSE |
| **PE-Cer 14:1;2O/36:4;O** | C52H97N2O7P | PE_Cer | FALSE |
| **PS O-18:5_28:7** | C52H80NO9P | EtherPS | TRUE |
| **PC O-18:2_22:6;2O** | C48H82NO9P | EtherOxPC | FALSE |
| **PC O-15:0_22:3;4O** | C45H86NO11P | EtherOxPC | TRUE |
| **HexCer 17:1;3O/26:5;(2OH)** | C49H85NO10 | HexCer_AP | TRUE |
| **SHexCer 41:1;3O** | C47H91NO12S | SHexCer | FALSE |
| **PE O-21:4_28:7** | C54H88NO7P | EtherPE | FALSE |
| **PC 24:0_14:1;2O** | C46H90NO10P | OxPC | FALSE |
| **HexCer 16:0;2O/28:5;O** | C50H89NO9 | HexCer_HDS | TRUE |
| **SHexCer 42:0;2O** | C48H95NO11S | SHexCer | TRUE |
| **PC O-12:0_28:0;1O** | C48H98NO8P | EtherOxPC | FALSE |
| **PG 20:3_22:5;3O** | C48H79O13P | OxPG | FALSE |
| **PC O-17:2_22:6;3O** | C47H80NO10P | EtherOxPC | TRUE |
| **HexCer 17:1;3O/26:4;(2OH)** | C49H87NO10 | HexCer_AP | FALSE |
| **SHexCer 41:0;3O** | C47H93NO12S | SHexCer | TRUE |
| **HexCer 16:0;2O/28:4;O** | C50H91NO9 | HexCer_HDS | FALSE |
| **CerP 27:3;2O/28:6** | C55H94NO6P | CerP | TRUE |
| **PG 16:3_30:8** | C52H81O10P | PG | FALSE |
| **PG O-19:4_28:7** | C53H85O9P | EtherPG | FALSE |
| **CL 18:0_22:4_28:0_28:0** | C105H198O17P2 | CL | FALSE |
| **SHexCer 42:6;3O** | C48H83NO12S | SHexCer | TRUE |
| **PC O-15:4_28:7** | C51H82NO7P | EtherPC | FALSE |
| **HexCer 17:3;3O/30:6;(2OH)** | C53H87NO10 | HexCer_AP | FALSE |
| **PE O-21:2_28:7** | C54H92NO7P | EtherPE | TRUE |
| **HexCer 16:1;2O/28:2;O** | C50H93NO9 | HexCer_HS | TRUE |
| **HexCer 16:0;2O/28:3;O** | C50H93NO9 | HexCer_HDS | TRUE |
| **PG O-19:3_28:7** | C53H87O9P | EtherPG | TRUE |
| **PE-Cer 16:2;2O/36:6** | C54H95N2O6P | PE_Cer | TRUE |
| **HexCer 17:2;3O/30:6;(2OH)** | C53H89NO10 | HexCer_AP | FALSE |
| **Cer 12:0;2O/22:6;(3OH)(FA 22:6)** | C56H87NO5 | Cer_EBDS | TRUE |
| **HexCer 16:1;2O/32:6;O** | C54H93NO9 | HexCer_HS | FALSE |
| **CerP 27:1;2O/28:6** | C55H98NO6P | CerP | TRUE |
| **HexCer 45:1;2O** | C51H99NO8 | HexCer_NS | TRUE |
| **PG 8:0_38:9** | C52H85O10P | PG | FALSE |
| **PA 23:0_28:7** | C54H93O8P | PA | TRUE |
| **ASG 28:2;O;Hex;FA 20:0** | C54H94O7 | AHexBRS | TRUE |
| **PE-Cer 16:1;2O/36:6** | C54H97N2O6P | PE_Cer | TRUE |
| **SM 23:2;2O/22:0** | C50H99N2O6P | SM | TRUE |
| **SM 12:1;2O/33:1** | C50H99N2O6P | SM | FALSE |
| **SM 12:2;2O/33:0** | C50H99N2O6P | SM | TRUE |
| **SHexCer 42:4;3O** | C48H87NO12S | SHexCer | TRUE |
| **PC O-20:4_22:6;1O** | C50H82NO8P | EtherOxPC | FALSE |
| **PC O-17:0_22:5;3O** | C47H86NO10P | EtherOxPC | TRUE |
| **SHexCer 43:3;2O** | C49H91NO11S | SHexCer | TRUE |
| **HexCer 17:1;3O/30:6;(2OH)** | C53H91NO10 | HexCer_AP | TRUE |
| **Cer 12:0;2O/22:5;(3OH)(FA 22:6)** | C56H89NO5 | Cer_EBDS | FALSE |
| **PE O-21:0_28:7** | C54H96NO7P | EtherPE | FALSE |
| **HexCer 17:2;3O/26:0;(2OH)** | C49H93NO10 | HexCer_AP | TRUE |
| **HexCer 16:1;2O/28:0;O** | C50H97NO9 | HexCer_HS | TRUE |
| **CerP 27:1;2O/28:5** | C55H100NO6P | CerP | TRUE |
| **HexCer 21:0;2O/23:1;O** | C50H97NO9 | HexCer_HDS | TRUE |
| **HexCer 17:0;2O/32:5** | C55H99NO8 | HexCer_NDS | TRUE |
| **PI 16:0_22:4;1O** | C47H83O14P | OxPI | TRUE |
| **PI 18:0_20:4;1O** | C47H83O14P | OxPI | TRUE |
| **SM 12:2;2O/34:6** | C51H89N2O6P | SM | TRUE |
| **PG O-19:1_28:7** | C53H91O9P | EtherPG | TRUE |
| **PE-Cer 15:1;2O/36:6;O** | C53H95N2O7P | PE_Cer | TRUE |
| **SM 44:2;3O** | C49H97N2O7P | SM | TRUE |
| **PE-Cer 16:1;2O/36:5** | C54H99N2O6P | PE_Cer | TRUE |
| **SM 12:1;2O/33:0** | C50H101N2O6P | SM | TRUE |
| **PS 22:3_22:5;1O** | C50H82NO11P | OxPS | FALSE |
| **PE 15:4_34:9** | C54H82NO8P | PE | FALSE |
| **Hex2Cer 34:3;2O** | C46H83NO13 | Hex2Cer | FALSE |
| **HexCer 16:2;3O/28:6;(2OH)** | C50H83NO10 | HexCer_AP | FALSE |
| **PE O-22:6_28:7** | C55H86NO7P | EtherPE | TRUE |
| **SHexCer 42:3;3O** | C48H89NO12S | SHexCer | TRUE |
| **SHexCer 43:2;2O** | C49H93NO11S | SHexCer | TRUE |
| **PE 10:0_38:6** | C53H94NO8P | PE | TRUE |
| **PC O-17:2_24:0;1O** | C49H96NO8P | EtherOxPC | FALSE |
| **PE O-21:0_28:6** | C54H98NO7P | EtherPE | FALSE |
| **PE O-28:0_18:1;2O** | C51H102NO9P | EtherOxPE | FALSE |
| **HexCer 22:0;2O/22:0;O** | C50H99NO9 | HexCer_HDS | TRUE |
| **PG 18:0_22:5;4O** | C46H81O15P | OxPG | FALSE |
| **SMGDG O-15:2_26:7** | C50H80O12S | EtherSMGDG | FALSE |
| **PG 8:0_38:7** | C52H89O10P | PG | FALSE |
| **SM 45:8;3O** | C50H87N2O7P | SM | FALSE |
| **PG O-19:0_28:7** | C53H93O9P | EtherPG | TRUE |
| **PE-Cer 15:1;2O/36:5;O** | C53H97N2O7P | PE_Cer | TRUE |
| **PG 38:0_7:0** | C51H101O10P | PG | FALSE |
| **PG O-18:0_28:0** | C52H105O9P | EtherPG | TRUE |
| **PC O-20:5_20:5;3O** | C48H78NO10P | EtherOxPC | FALSE |
| **PC O-16:1_22:3;4O** | C46H86NO11P | EtherOxPC | TRUE |
| **PE 15:3_34:9** | C54H84NO8P | PE | FALSE |
| **Hex2Cer 34:2;2O** | C46H85NO13 | Hex2Cer | FALSE |
| **SHexCer 42:2;3O** | C48H91NO12S | SHexCer | TRUE |
| **PE O-22:5_28:7** | C55H88NO7P | EtherPE | TRUE |
| **PC O-15:0_28:7** | C51H90NO7P | EtherPC | TRUE |
| **SHexCer 43:1;2O** | C49H95NO11S | SHexCer | TRUE |
| **PC O-26:0_15:1;1O** | C49H98NO8P | EtherOxPC | TRUE |
| **HexCer 16:0;3O/27:0;(2OH)** | C49H97NO10 | HexCer_AP | FALSE |
| **PE O-21:0_28:5** | C54H100NO7P | EtherPE | TRUE |
| **PI 6:0_34:8** | C49H79O13P | PI | FALSE |
| **PG 8:0_38:6** | C52H91O10P | PG | FALSE |
| **PG O-19:0_28:6** | C53H95O9P | EtherPG | FALSE |
| **SM 12:1;2O/34:5** | C51H93N2O6P | SM | FALSE |
| **PE-Cer 15:1;2O/36:4;O** | C53H99N2O7P | PE_Cer | FALSE |
| **PI-Cer 12:1;2O/32:6** | C50H86NO11P | PI_Cer | TRUE |
| **PS 22:3_22:3;1O** | C50H86NO11P | OxPS | FALSE |
| **PE 13:1_36:10** | C54H86NO8P | PE | TRUE |
| **Hex2Cer 34:1;2O** | C46H87NO13 | Hex2Cer | FALSE |
| **HexCer 16:1;3O/28:5;(2OH)** | C50H87NO10 | HexCer_AP | TRUE |
| **SHexCer 42:1;3O** | C48H93NO12S | SHexCer | TRUE |
| **PC 24:0_15:1;2O** | C47H92NO10P | OxPC | TRUE |
| **SHexCer 43:0;2O** | C49H97NO11S | SHexCer | FALSE |
| **HexCer 17:1;3O/30:3;(2OH)** | C53H97NO10 | HexCer_AP | FALSE |
| **PI 15:0_20:4;4O** | C44H77O17P | OxPI | FALSE |
| **PI 6:0_34:7** | C49H81O13P | PI | FALSE |
| **SMGDG O-13:0_28:7** | C50H84O12S | EtherSMGDG | TRUE |
| **PI 14:1_24:0;1O** | C47H89O14P | OxPI | FALSE |
| **PG 8:0_38:5** | C52H93O10P | PG | FALSE |
| **CL 20:3_22:3_28:0_28:0** | C107H198O17P2 | CL | TRUE |
| **PC O-18:5_26:7** | C52H82NO7P | EtherPC | FALSE |
| **PI-Cer 12:1;2O/32:5** | C50H88NO11P | PI_Cer | TRUE |
| **Hex2Cer 34:0;2O** | C46H89NO13 | Hex2Cer | TRUE |
| **HexCer 16:1;3O/28:4;(2OH)** | C50H89NO10 | HexCer_AP | FALSE |
| **SHexCer 42:0;3O** | C48H95NO12S | SHexCer | FALSE |
| **HexCer 17:0;2O/28:4;O** | C51H93NO9 | HexCer_HDS | TRUE |
| **PG 8:0_38:4** | C52H95O10P | PG | FALSE |
| **PE-Cer 17:3;2O/36:6** | C55H95N2O6P | PE_Cer | TRUE |
| **SHexCer 44:5;2O** | C50H89NO11S | SHexCer | TRUE |
| **PE 11:0_38:9** | C54H90NO8P | PE | FALSE |
| **PE O-22:2_28:7** | C55H94NO7P | EtherPE | TRUE |
| **PE O-20:3_28:0;1O** | C53H102NO8P | EtherOxPE | FALSE |
| **HexCer 16:1;3O/31:1;(2OH)** | C53H101NO10 | HexCer_AP | FALSE |
| **PE-Cer 16:3;2O/36:6;O** | C54H93N2O7P | PE_Cer | TRUE |
| **SM 45:4;3O** | C50H95N2O7P | SM | FALSE |
| **PE-Cer 17:2;2O/36:6** | C55H97N2O6P | PE_Cer | TRUE |
| **NAGlySer 23:0;O(FA 28:4)** | C56H100N2O7 | NAGlySer | TRUE |
| **PS 18:3_26:0;1O** | C50H92NO11P | OxPS | FALSE |
| **PE 11:0_38:8** | C54H92NO8P | PE | TRUE |
| **PC 18:1_24:2** | C50H94NO8P | PC | TRUE |
| **HexCer 17:1;2O/32:6;O** | C55H95NO9 | HexCer_HS | TRUE |
| **CL 28:0_14:1_28:0_28:0** | C107H208O17P2 | CL | TRUE |
| **CerP 28:1;2O/28:6** | C56H100NO6P | CerP | TRUE |
| **PG 20:3_22:3;4O** | C48H83O14P | OxPG | TRUE |
| **PI 6:0_34:4** | C49H87O13P | PI | TRUE |
| **PA 24:0_28:7** | C55H95O8P | PA | TRUE |
| **SM 45:3;3O** | C50H97N2O7P | SM | FALSE |
| **PG 23:1_23:1** | C52H99O10P | PG | TRUE |
| **PC 7:0_36:9** | C51H84NO8P | PC | FALSE |
| **SHexCer 44:3;2O** | C50H93NO11S | SHexCer | TRUE |
| **PI-Cer 12:1;2O/32:2** | C50H94NO11P | PI_Cer | TRUE |
| **PE O-22:0_28:7** | C55H98NO7P | EtherPE | TRUE |
| **HexCer 16:1;2O/29:0;O** | C51H99NO9 | HexCer_HS | TRUE |
| **PI 6:0_34:3** | C49H89O13P | PI | TRUE |
| **NAGlySer 24:3;O(FA 28:6)** | C57H92N2O7 | NAGlySer | FALSE |
| **PA 24:0_28:6** | C55H97O8P | PA | TRUE |
| **PG 8:0_38:1** | C52H101O10P | PG | TRUE |
| **SM 45:2;3O** | C50H99N2O7P | SM | TRUE |
| **PG 26:0_20:1** | C52H101O10P | PG | TRUE |
| **PG O-19:0_28:1** | C53H105O9P | EtherPG | TRUE |
| **PC O-17:2_22:3;4O** | C47H86NO11P | EtherOxPC | TRUE |
| **PC 7:0_36:8** | C51H86NO8P | PC | FALSE |
| **PC O-18:0_22:4;4O** | C48H90NO10P | EtherOxPC | FALSE |
| **SHexCer 43:3;3O** | C49H91NO12S | SHexCer | TRUE |
| **SHexCer 44:2;2O** | C50H95NO11S | SHexCer | TRUE |
| **PC 17:2_24:0;1O** | C49H94NO9P | OxPC | TRUE |
| **PI-Cer 17:2;2O/27:0** | C50H96NO11P | PI_Cer | TRUE |
| **HexCer 16:1;3O/32:5;(2OH)** | C54H95NO10 | HexCer_AP | FALSE |
| **PC 16:0_26:1** | C50H98NO8P | PC | TRUE |
| **HexCer 17:0;2O/32:5;O** | C55H99NO9 | HexCer_HDS | TRUE |
| **PI 20:5_20:5;1O** | C49H75O14P | OxPI | TRUE |
| **PI 6:0_34:2** | C49H91O13P | PI | FALSE |
| **PA 24:0_28:5** | C55H99O8P | PA | TRUE |
| **PG 38:0_8:0** | C52H103O10P | PG | TRUE |
| **PS 22:3_22:5;2O** | C50H82NO12P | OxPS | TRUE |
| **SHexCer 45:8;2O** | C51H85NO11S | SHexCer | FALSE |
| **PC O-17:1_22:3;4O** | C47H88NO11P | EtherOxPC | FALSE |
| **Hex2Cer 35:2;2O** | C47H87NO13 | Hex2Cer | TRUE |
| **HexCer 17:1;3O/28:6;(2OH)** | C51H87NO10 | HexCer_AP | FALSE |
| **SHexCer 43:2;3O** | C49H93NO12S | SHexCer | TRUE |
| **SHexCer 44:1;2O** | C50H97NO11S | SHexCer | TRUE |
| **Cer 18:1;3O/38:2;(2OH)** | C56H107NO5 | Cer_AP | FALSE |
| **Cer 24:0;2O/38:6** | C62H113NO3 | Cer_NDS | TRUE |
| **PI 6:0_34:1** | C49H93O13P | PI | FALSE |
| **PI O-13:0_28:1** | C50H97O12P | EtherPI | FALSE |
| **SMGDG O-13:0_28:1** | C50H96O12S | EtherSMGDG | FALSE |
| **PS 15:3_32:9** | C53H80NO10P | PS | TRUE |
| **PS 22:3_22:4;2O** | C50H84NO12P | OxPS | TRUE |
| **PC O-20:3_22:5;2O** | C50H86NO9P | EtherOxPC | FALSE |
| **PC O-17:0_22:3;4O** | C47H90NO11P | EtherOxPC | TRUE |
| **SHexCer 43:1;3O** | C49H95NO12S | SHexCer | TRUE |
| **PC O-24:0_16:2;3O** | C48H94NO10P | EtherOxPC | TRUE |
| **HexCer 16:0;2O/30:5;O** | C52H93NO9 | HexCer_HDS | TRUE |
| **SHexCer 44:0;2O** | C50H99NO11S | SHexCer | TRUE |
| **PG 22:3_22:5;3O** | C50H83O13P | OxPG | FALSE |
| **PG 24:0_18:2;4O** | C48H91O14P | OxPG | FALSE |
| **PMeOH 24:3_28:7** | C56H91O8P | PMeOH | TRUE |
| **CL 22:3_22:3_28:0_28:0** | C109H202O17P2 | CL | TRUE |
| **PS 22:3_22:3;2O** | C50H86NO12P | OxPS | TRUE |
| **PC O-19:5_26:7** | C53H84NO7P | EtherPC | FALSE |
| **SHexCer 45:6;2O** | C51H89NO11S | SHexCer | FALSE |
| **PE 12:0_38:10** | C55H90NO8P | PE | FALSE |
| **PE 20:4_30:6** | C55H90NO8P | PE | TRUE |
| **SHexCer 43:0;3O** | C49H97NO12S | SHexCer | TRUE |
| **PC O-24:0_16:1;3O** | C48H96NO10P | EtherOxPC | FALSE |
| **PC O-24:0_17:0;2O** | C49H100NO9P | EtherOxPC | TRUE |
| **PG 24:0_18:1;4O** | C48H93O14P | OxPG | TRUE |
| **PMeOH 24:2_28:7** | C56H93O8P | PMeOH | TRUE |
| **PG 9:0_38:4** | C53H97O10P | PG | FALSE |
| **SM 46:5;3O** | C51H95N2O7P | SM | FALSE |
| **PE-Cer 18:3;2O/36:6** | C56H97N2O6P | PE_Cer | FALSE |
| **PC O-17:4_28:7** | C53H86NO7P | EtherPC | FALSE |
| **PC O-20:3_22:3;2O** | C50H90NO9P | EtherOxPC | TRUE |
| **PE 12:0_38:9** | C55H92NO8P | PE | TRUE |
| **PC O-24:0_16:0;3O** | C48H98NO10P | EtherOxPC | TRUE |
| **HexCer 16:0;2O/30:3;O** | C52H97NO9 | HexCer_HDS | FALSE |
| **PG 20:3_22:5;4O** | C48H79O15P | OxPG | TRUE |
| **PG 24:0_18:0;4O** | C48H95O14P | OxPG | TRUE |
| **PG O-28:5_21:5** | C55H91O9P | EtherPG | TRUE |
| **PA 25:1_28:7** | C56H95O8P | PA | TRUE |
| **PE-Cer 18:2;2O/36:6** | C56H99N2O6P | PE_Cer | TRUE |
| **PC O-18:1_22:6;4O** | C48H84NO11P | EtherOxPC | FALSE |
| **PE 12:0_38:8** | C55H94NO8P | PE | TRUE |
| **PE O-23:1_28:7** | C56H98NO7P | EtherPE | TRUE |
| **HexCer 16:0;2O/30:2;O** | C52H99NO9 | HexCer_HDS | TRUE |
| **PI O-26:7_17:4** | C52H81O12P | EtherPI | TRUE |
| **PG O-21:2_28:7** | C55H93O9P | EtherPG | FALSE |
| **PEtOH 23:0_28:7** | C56H97O8P | PEtOH | TRUE |
| **SM 46:3;3O** | C51H99N2O7P | SM | TRUE |
| **PG O-20:0_28:2** | C54H105O9P | EtherPG | FALSE |
| **PE 15:4_36:10** | C56H84NO8P | PE | FALSE |
| **SHexCer 45:3;2O** | C51H95NO11S | SHexCer | FALSE |
| **PC 18:1_25:1** | C51H98NO8P | PC | TRUE |
| **PE O-23:0_28:7** | C56H100NO7P | EtherPE | FALSE |
| **SL 21:1;O/36:5;O** | C57H103NO6S | SL | FALSE |
| **Cer 25:2;2O/38:6** | C63H111NO3 | Cer_NS | TRUE |
| **SM 12:2;2O/36:6** | C53H93N2O6P | SM | FALSE |
| **PE-Cer 17:1;2O/36:6;O** | C55H99N2O7P | PE_Cer | FALSE |
| **PG 9:0_38:1** | C53H103O10P | PG | TRUE |
| **PC O-18:2_22:3;4O** | C48H88NO11P | EtherOxPC | FALSE |
| **Hex2Cer 36:3;2O** | C48H87NO13 | Hex2Cer | FALSE |
| **SHexCer 44:3;3O** | C50H93NO12S | SHexCer | TRUE |
| **SHexCer 45:2;2O** | C51H97NO11S | SHexCer | TRUE |
| **PC O-24:0_18:3;2O** | C50H96NO9P | EtherOxPC | FALSE |
| **PE O-23:0_28:6** | C56H102NO7P | EtherPE | TRUE |
| **Cer 25:1;2O/38:6** | C63H113NO3 | Cer_NS | TRUE |
| **Cer 22:1;3O/38:1;(2OH)** | C60H117NO5 | Cer_AP | TRUE |
| **PG 22:5_22:6;4O** | C50H77O14P | OxPG | TRUE |
| **PI 42:9** | C51H81O13P | PI | TRUE |
| **SMGDG O-15:2_28:7** | C52H84O12S | EtherSMGDG | FALSE |
| **PE-Cer 17:1;2O/36:5;O** | C55H101N2O7P | PE_Cer | TRUE |
| **SHexCer 46:8;2O** | C52H87NO11S | SHexCer | FALSE |
| **PI-Cer 12:2;2O/34:6** | C52H88NO11P | PI_Cer | FALSE |
| **PC O-18:1_22:3;4O** | C48H90NO11P | EtherOxPC | TRUE |
| **Hex2Cer 36:2;2O** | C48H89NO13 | Hex2Cer | FALSE |
| **PC O-22:3_22:5;1O** | C52H90NO8P | EtherOxPC | TRUE |
| **SHexCer 44:2;3O** | C50H95NO12S | SHexCer | TRUE |
| **PC O-24:0_18:2;2O** | C50H98NO9P | EtherOxPC | FALSE |
| **PC O-15:1_28:0;1O** | C51H102NO8P | EtherOxPC | FALSE |
| **Cer 25:0;2O/38:6** | C63H115NO3 | Cer_NDS | TRUE |
| **PI 16:0_22:4;4O** | C47H83O16P | OxPI | TRUE |
| **PI O-15:1_28:7** | C52H87O12P | EtherPI | FALSE |
| **PG O-21:0_28:6** | C55H99O9P | EtherPG | FALSE |
| **SHexCer 46:7;2O** | C52H89NO11S | SHexCer | TRUE |
| **PC 24:0_16:2;3O** | C48H92NO11P | OxPC | TRUE |
| **Hex2Cer 36:1;2O** | C48H91NO13 | Hex2Cer | TRUE |
| **HexCer 22:0;3O/24:6;(2OH)** | C52H91NO10 | HexCer_AP | TRUE |
| **SHexCer 44:1;3O** | C50H97NO12S | SHexCer | TRUE |
| **PC 24:0_17:1;2O** | C49H96NO10P | OxPC | TRUE |
| **SHexCer 46:6;2O** | C52H91NO11S | SHexCer | TRUE |
| **PC O-18:5_28:7** | C54H86NO7P | EtherPC | FALSE |
| **PE 13:0_38:10** | C56H92NO8P | PE | TRUE |
| **PC 22:1_22:4** | C52H94NO8P | PC | TRUE |
| **HexCer 16:1;3O/30:4;(2OH)** | C52H93NO10 | HexCer_AP | TRUE |
| **SHexCer 44:0;3O** | C50H99NO12S | SHexCer | TRUE |
| **PE O-24:3_28:7** | C57H96NO7P | EtherPE | FALSE |
| **PC 24:0_17:0;2O** | C49H98NO10P | OxPC | TRUE |
| **SMGDG O-15:0_28:6** | C52H90O12S | EtherSMGDG | TRUE |
| **PA 26:2_28:7** | C57H95O8P | PA | TRUE |
| **PC O-20:3_22:4;4O** | C50H88NO10P | EtherOxPC | FALSE |
| **PC O-18:4_28:7** | C54H88NO7P | EtherPC | TRUE |
| **PE 13:0_38:9** | C56H94NO8P | PE | TRUE |
| **PE O-24:2_28:7** | C57H98NO7P | EtherPE | TRUE |
| **HexCer 16:1;3O/30:3;(2OH)** | C52H95NO10 | HexCer_AP | TRUE |
| **PA 26:1_28:7** | C57H97O8P | PA | TRUE |
| **PE-Cer 19:2;2O/36:6** | C57H101N2O6P | PE_Cer | FALSE |
| **PG O-21:0_28:3** | C55H105O9P | EtherPG | TRUE |
| **PC 7:0_38:10** | C53H86NO8P | PC | FALSE |
| **PE 13:0_38:8** | C56H96NO8P | PE | TRUE |
| **PE O-24:1_28:7** | C57H100NO7P | EtherPE | FALSE |
| **PI 6:0_36:4** | C51H91O13P | PI | FALSE |
| **SM 15:3;2O/34:6** | C54H93N2O6P | SM | TRUE |
| **PA 26:0_28:7** | C57H99O8P | PA | TRUE |
| **ASG 28:2;O;Hex;FA 23:0** | C57H100O7 | AHexBRS | TRUE |
| **PC 7:0_38:9** | C53H88NO8P | PC | FALSE |
| **SHexCer 46:3;2O** | C52H97NO11S | SHexCer | TRUE |
| **PE 13:0_38:7** | C56H98NO8P | PE | FALSE |
| **PC 18:1_26:1** | C52H100NO8P | PC | TRUE |
| **HexCer 17:0;2O/34:6;O** | C57H101NO9 | HexCer_HDS | TRUE |
| **PG O-22:1_28:7** | C56H97O9P | EtherPG | TRUE |
| **SM 13:2;2O/36:6** | C54H95N2O6P | SM | FALSE |
| **PG 10:0_38:1** | C54H105O10P | PG | FALSE |
| **PS 24:0_20:3;3O** | C50H92NO13P | OxPS | TRUE |
| **PC 7:0_38:8** | C53H90NO8P | PC | TRUE |
| **PC O-18:1_28:7** | C54H94NO7P | EtherPC | FALSE |
| **PC 17:2_26:0;1O** | C51H98NO9P | OxPC | FALSE |
| **PE 13:0_38:6** | C56H100NO8P | PE | TRUE |
| **PG 24:0_20:4;4O** | C50H91O14P | OxPG | TRUE |
| **PG O-22:0_28:7** | C56H99O9P | EtherPG | FALSE |
| **PG 10:0_38:0** | C54H107O10P | PG | FALSE |
| **SHexCer 46:9;3O** | C52H85NO12S | SHexCer | TRUE |
| **SHexCer 47:8;2O** | C53H89NO11S | SHexCer | TRUE |
| **PE 16:2_36:10** | C57H90NO8P | PE | TRUE |
| **SHexCer 45:2;3O** | C51H97NO12S | SHexCer | TRUE |
| **Cer 23:1;3O/38:0;(2OH)** | C61H121NO5 | Cer_AP | TRUE |
| **PG 24:0_20:3;4O** | C50H93O14P | OxPG | TRUE |
| **PS 15:3_34:9** | C55H84NO10P | PS | TRUE |
| **PS 26:0_18:1;3O** | C50H96NO13P | OxPS | TRUE |
| **SHexCer 45:1;3O** | C51H99NO12S | SHexCer | TRUE |
| **PC O-24:0_18:2;3O** | C50H98NO10P | EtherOxPC | FALSE |
| **PG 26:0_18:2;4O** | C50H95O14P | OxPG | TRUE |
| **Hex2Cer 38:7;2O** | C50H83NO13 | Hex2Cer | TRUE |
| **PC O-19:5_28:7** | C55H88NO7P | EtherPC | TRUE |
| **PI-Cer 13:1;2O/34:5** | C53H94NO11P | PI_Cer | TRUE |
| **PS 26:0_18:0;3O** | C50H98NO13P | OxPS | FALSE |
| **PC O-24:0_18:1;3O** | C50H100NO10P | EtherOxPC | TRUE |
| **PG 22:4_22:5;4O** | C50H81O15P | OxPG | TRUE |
| **PG 26:0_18:1;4O** | C50H97O14P | OxPG | TRUE |
| **PC 16:3_30:8** | C54H86NO8P | PC | TRUE |
| **SHexCer 46:6;3O** | C52H91NO12S | SHexCer | FALSE |
| **PC O-19:4_28:7** | C55H90NO7P | EtherPC | TRUE |
| **SHexCer 47:5;2O** | C53H95NO11S | SHexCer | TRUE |
| **PE 14:0_38:9** | C57H96NO8P | PE | TRUE |
| **SMGDG O-19:5_26:7** | C54H82O12S | EtherSMGDG | TRUE |
| **PA 27:1_28:7** | C58H99O8P | PA | TRUE |
| **PC O-20:4_22:3;4O** | C50H88NO11P | EtherOxPC | TRUE |
| **PC 8:0_38:10** | C54H88NO8P | PC | FALSE |
| **SHexCer 46:5;3O** | C52H93NO12S | SHexCer | TRUE |
| **SHexCer 47:4;2O** | C53H97NO11S | SHexCer | TRUE |
| **PE 14:0_38:8** | C57H98NO8P | PE | TRUE |
| **PC 7:0_38:3** | C53H100NO8P | PC | FALSE |
| **PE O-25:1_28:7** | C58H102NO7P | EtherPE | FALSE |
| **PI O-21:5_24:6** | C54H85O12P | EtherPI | TRUE |
| **PA 27:0_28:7** | C58H101O8P | PA | TRUE |
| **PE 15:4_38:10** | C58H88NO8P | PE | TRUE |
| **PC 8:0_38:9** | C54H90NO8P | PC | FALSE |
| **PC O-19:2_28:7** | C55H94NO7P | EtherPC | TRUE |
| **PE 14:0_38:7** | C57H100NO8P | PE | TRUE |
| **Cer 27:2;2O/38:6** | C65H115NO3 | Cer_NS | TRUE |
| **Cer 24:1;3O/38:2;(2OH)** | C62H119NO5 | Cer_AP | TRUE |
| **PG 12:0_38:8** | C56H95O10P | PG | TRUE |
| **SM 12:2;2O/38:6** | C55H97N2O6P | SM | TRUE |
| **NAGlySer 26:0;O(FA 28:2)** | C59H110N2O7 | NAGlySer | FALSE |
| **PE 15:3_38:10** | C58H90NO8P | PE | TRUE |
| **PS 28:0_18:2;2O** | C52H98NO12P | OxPS | TRUE |
| **PE 14:0_38:6** | C57H102NO8P | PE | FALSE |
| **HexCer 17:1;3O/34:5;(2OH)** | C57H101NO10 | HexCer_AP | FALSE |
| **Cer 27:1;2O/38:6** | C65H117NO3 | Cer_NS | TRUE |
| **SMGDG O-16:0_28:2** | C53H100O12S | EtherSMGDG | FALSE |
| **PG O-23:0_28:7** | C57H101O9P | EtherPG | TRUE |
| **PE-Cer 19:1;2O/36:5;O** | C57H105N2O7P | PE_Cer | FALSE |
| **SHexCer 48:8;2O** | C54H91NO11S | SHexCer | TRUE |
| **PC 24:0_18:3;3O** | C50H94NO11P | OxPC | TRUE |
| **PC 22:3_24:4** | C54H94NO8P | PC | TRUE |
| **SHexCer 46:2;3O** | C52H99NO12S | SHexCer | TRUE |
| **PE 14:0_38:5** | C57H104NO8P | PE | TRUE |
| **SM 12:1;2O/38:5** | C55H101N2O6P | SM | TRUE |
| **SHexCer 47:8;3O** | C53H89NO12S | SHexCer | FALSE |
| **SHexCer 48:7;2O** | C54H93NO11S | SHexCer | TRUE |
| **PC O-24:0_18:3;4O** | C50H96NO11P | EtherOxPC | TRUE |
| **Hex2Cer 38:1;2O** | C50H95NO13 | Hex2Cer | TRUE |
| **PC 26:0_17:1;2O** | C51H100NO10P | OxPC | FALSE |
| **CL 28:0_22:6_28:0_28:0** | C115H214O17P2 | CL | TRUE |
| **Hex2Cer 39:7;2O** | C51H85NO13 | Hex2Cer | FALSE |
| **PC O-20:5_28:7** | C56H90NO7P | EtherPC | TRUE |
| **SHexCer 48:6;2O** | C54H95NO11S | SHexCer | TRUE |
| **PE 15:0_38:10** | C58H96NO8P | PE | TRUE |
| **PC 24:1_22:4** | C54H98NO8P | PC | TRUE |
| **PE O-26:3_28:7** | C59H100NO7P | EtherPE | TRUE |
| **PC O-20:4_22:5;4O** | C50H84NO12P | EtherOxPC | FALSE |
| **PE 15:0_38:9** | C58H98NO8P | PE | TRUE |
| **PC O-20:5_26:0;1O** | C54H100NO8P | EtherOxPC | TRUE |
| **HexCer 16:1;3O/32:3;(2OH)** | C54H99NO10 | HexCer_AP | FALSE |
| **PE-Cer 20:3;2O/36:6;O** | C58H101N2O7P | PE_Cer | TRUE |
| **SHexCer 47:5;3O** | C53H95NO12S | SHexCer | TRUE |
| **PE 15:0_38:8** | C58H100NO8P | PE | TRUE |
| **SMGDG O-22:5_24:6** | C55H86O12S | EtherSMGDG | TRUE |
| **PG 24:0_22:6;4O** | C52H91O14P | OxPG | TRUE |
| **PC 20:3_22:3;4O** | C50H88NO12P | OxPC | TRUE |
| **PS 24:0_22:4;4O** | C52H94NO13P | OxPS | TRUE |
| **PC 9:0_38:9** | C55H92NO8P | PC | FALSE |
| **HexCer 17:3;3O/32:6;(2OH)** | C55H91NO10 | HexCer_AP | TRUE |
| **PC O-20:2_28:7** | C56H96NO7P | EtherPC | TRUE |
| **HexCer 16:1;3O/36:6;(2OH)** | C58H101NO10 | HexCer_AP | TRUE |
| **Cer 28:2;2O/38:6** | C66H117NO3 | Cer_NS | TRUE |
| **SMGDG O-18:3_28:7** | C55H88O12S | EtherSMGDG | FALSE |
| **PI O-18:3_28:7** | C55H89O12P | EtherPI | TRUE |
| **PI 6:0_38:3** | C53H97O13P | PI | TRUE |
| **SMGDG O-17:0_28:3** | C54H100O12S | EtherSMGDG | TRUE |
| **SHexCer 49:9;2O** | C55H91NO11S | SHexCer | TRUE |
| **PE 16:3_38:10** | C59H92NO8P | PE | TRUE |
| **HexCer 16:1;3O/36:5;(2OH)** | C58H103NO10 | HexCer_AP | TRUE |
| **HexCer 16:1;3O/32:0;(2OH)** | C54H105NO10 | HexCer_AP | TRUE |
| **Cer 28:1;2O/38:6** | C66H119NO3 | Cer_NS | TRUE |
| **Cer 25:1;3O/38:1;(2OH)** | C63H123NO5 | Cer_AP | TRUE |
| **SMGDG O-18:2_28:7** | C55H90O12S | EtherSMGDG | TRUE |
| **PG 26:0_20:4;4O** | C52H95O14P | OxPG | TRUE |
| **PG O-23:0_28:0** | C57H115O9P | EtherPG | TRUE |
| **SHexCer 49:8;2O** | C55H93NO11S | SHexCer | TRUE |
| **PE 16:2_38:10** | C59H94NO8P | PE | TRUE |
| **HexCer 16:1;3O/36:4;(2OH)** | C58H105NO10 | HexCer_AP | FALSE |
| **NAGlySer 27:0;O(FA 28:0)** | C60H116N2O7 | NAGlySer | TRUE |
| **Hex2Cer 40:8;2O** | C52H85NO13 | Hex2Cer | FALSE |
| **PS O-24:5_28:7** | C58H92NO9P | EtherPS | TRUE |
| **SHexCer 49:7;2O** | C55H95NO11S | SHexCer | TRUE |
| **PE 16:1_38:10** | C59H96NO8P | PE | TRUE |
| **PI 7:0_38:7** | C54H91O13P | PI | TRUE |
| **PI O-18:0_28:7** | C55H95O12P | EtherPI | FALSE |
| **Hex2Cer 40:7;2O** | C52H87NO13 | Hex2Cer | TRUE |
| **PC 9:0_38:5** | C55H100NO8P | PC | FALSE |
| **PI 18:0_22:3;4O** | C49H89O17P | OxPI | TRUE |
| **PI 22:3_22:4;1O** | C53H89O14P | OxPI | TRUE |
| **PC 16:2_32:9** | C56H90NO8P | PC | FALSE |
| **PE 16:0_38:9** | C59H100NO8P | PE | TRUE |
| **HexCer 17:1;3O/32:3;(2OH)** | C55H101NO10 | HexCer_AP | FALSE |
| **HexCer 16:2;2O/38:6;O** | C60H103NO9 | HexCer_HS | TRUE |
| **Cer 23:1;3O/38:6;(2OH)** | C61H109NO5 | Cer_AP | TRUE |
| **PI O-21:5_26:7** | C56H87O12P | EtherPI | TRUE |
| **SMGDG O-19:5_28:7** | C56H86O12S | EtherSMGDG | TRUE |
| **PC O-22:4_22:3;4O** | C52H92NO11P | EtherOxPC | TRUE |
| **SHexCer 48:5;3O** | C54H97NO12S | SHexCer | FALSE |
| **HexCer 17:2;3O/36:6;(2OH)** | C59H101NO10 | HexCer_AP | TRUE |
| **PI 20:5_24:0;1O** | C53H93O14P | OxPI | TRUE |
| **PE 17:4_38:10** | C60H92NO8P | PE | TRUE |
| **SHexCer 48:4;3O** | C54H99NO12S | SHexCer | TRUE |
| **PE 16:0_38:7** | C59H104NO8P | PE | FALSE |
| **HexCer 17:1;3O/36:6;(2OH)** | C59H103NO10 | HexCer_AP | TRUE |
| **Cer 29:2;2O/38:6** | C67H119NO3 | Cer_NS | TRUE |
| **SMGDG O-19:3_28:7** | C56H90O12S | EtherSMGDG | TRUE |
| **PI 20:4_24:0;1O** | C53H95O14P | OxPI | TRUE |
| **SM 51:9;3O** | C56H97N2O7P | SM | TRUE |
| **PS 26:0_20:4;4O** | C52H94NO14P | OxPS | TRUE |
| **PC O-24:0_20:5;4O** | C52H96NO11P | EtherOxPC | FALSE |
| **PC O-21:1_28:7** | C57H100NO7P | EtherPC | TRUE |
| **PE 16:0_38:6** | C59H106NO8P | PE | TRUE |
| **HexCer 16:0;2O/38:5;O** | C60H109NO9 | HexCer_HDS | TRUE |
| **Cer 29:1;2O/38:6** | C67H121NO3 | Cer_NS | TRUE |
| **PG O-25:0_28:7** | C59H105O9P | EtherPG | TRUE |
| **SHexCer 50:8;2O** | C56H95NO11S | SHexCer | TRUE |
| **Hex2Cer 40:2;2O** | C52H97NO13 | Hex2Cer | TRUE |
| **PI 8:0_38:8** | C55H91O13P | PI | TRUE |
| **PG O-25:0_28:6** | C59H107O9P | EtherPG | TRUE |
| **PC O-22:6_28:7** | C58H92NO7P | EtherPC | TRUE |
| **PE 17:1_38:10** | C60H98NO8P | PE | TRUE |
| **HexCer 20:0;3O/30:6;(2OH)** | C56H99NO10 | HexCer_AP | TRUE |
| **PI 24:0_18:3;3O** | C51H93O16P | OxPI | TRUE |
| **PE-Cer 21:1;2O/36:3;O** | C59H113N2O7P | PE_Cer | FALSE |
| **Hex2Cer 41:7;2O** | C53H89NO13 | Hex2Cer | TRUE |
| **PC O-22:5_28:7** | C58H94NO7P | EtherPC | TRUE |
| **PE 17:0_38:10** | C60H100NO8P | PE | TRUE |
| **HexCer 16:1;3O/34:4;(2OH)** | C56H101NO10 | HexCer_AP | FALSE |
| **Cer 24:2;3O/38:6;(2OH)** | C62H109NO5 | Cer_AP | TRUE |
| **PI 20:4_22:6;4O** | C51H79O17P | OxPI | TRUE |
| **PG 15:1_38:10** | C59H95O10P | PG | TRUE |
| **PI O-19:0_28:6** | C56H99O12P | EtherPI | TRUE |
| **PC 13:1_36:10** | C57H92NO8P | PC | TRUE |
| **SHexCer 49:6;3O** | C55H97NO12S | SHexCer | FALSE |
| **PE 17:0_38:9** | C60H102NO8P | PE | TRUE |
| **HexCer 16:1;3O/37:1;(2OH)** | C59H113NO10 | HexCer_AP | TRUE |
| **Cer 29:0;2O/38:3** | C67H129NO3 | Cer_NDS | TRUE |
| **PG 15:0_38:10** | C59H97O10P | PG | TRUE |
| **HexCer 16:1;3O/34:2;(2OH)** | C56H105NO10 | HexCer_AP | TRUE |
| **PE 16:0_38:1** | C59H116NO8P | PE | TRUE |
| **HexCer 18:0;2O/38:6** | C62H111NO8 | HexCer_NDS | TRUE |
| **PG 26:0_22:6;4O** | C54H95O14P | OxPG | TRUE |
| **PE-Cer 22:2;2O/36:6;O** | C60H107N2O7P | PE_Cer | TRUE |
| **PC O-24:0_22:5;3O** | C54H100NO10P | EtherOxPC | TRUE |
| **PE 17:0_38:7** | C60H106NO8P | PE | TRUE |
| **HexCer 16:1;3O/34:1;(2OH)** | C56H107NO10 | HexCer_AP | TRUE |
| **PG 18:5_36:10** | C60H89O10P | PG | TRUE |
| **PG 28:0_20:5;4O** | C54H97O14P | OxPG | TRUE |
| **PE-Cer 22:1;2O/36:6;O** | C60H109N2O7P | PE_Cer | TRUE |
| **PS 24:0_22:5;4O** | C52H92NO15P | OxPS | FALSE |
| **SHexCer 51:9;2O** | C57H95NO11S | SHexCer | TRUE |
| **HexCer 16:1;3O/38:5;(2OH)** | C60H107NO10 | HexCer_AP | TRUE |
| **SMGDG O-20:2_28:7** | C57H94O12S | EtherSMGDG | TRUE |
| **PI O-20:2_28:7** | C57H95O12P | EtherPI | TRUE |
| **PC 24:0_20:4;4O** | C52H96NO12P | OxPC | TRUE |
| **HexCer 17:1;3O/34:6;(2OH)** | C57H99NO10 | HexCer_AP | TRUE |
| **PI 24:0_20:3;2O** | C53H97O15P | OxPI | TRUE |
| **PG 26:0_22:3;4O** | C54H101O14P | OxPG | TRUE |
| **Hex2Cer 41:1;2O** | C53H101NO13 | Hex2Cer | TRUE |
| **SHexCer 50:0;2O** | C56H111NO11S | SHexCer | FALSE |
| **PC 16:3_34:9** | C58H92NO8P | PC | TRUE |
| **PS O-26:4_28:7** | C60H98NO9P | EtherPS | TRUE |
| **PC 26:0_18:2;4O** | C52H100NO12P | OxPC | TRUE |
| **Hex2Cer 41:0;2O** | C53H103NO13 | Hex2Cer | TRUE |
| **Cer 28:1;3O/38:5;(2OH)** | C66H121NO5 | Cer_AP | TRUE |
| **PI 26:0_18:1;2O** | C53H101O15P | OxPI | TRUE |
| **PC 14:1_36:10** | C58H94NO8P | PC | TRUE |
| **Cer 25:1;3O/38:6;(2OH)** | C63H113NO5 | Cer_AP | TRUE |
| **PI 26:0_18:0;2O** | C53H103O15P | OxPI | TRUE |
| **PS 28:0_22:4;2O** | C56H102NO12P | OxPS | TRUE |
| **HexCer 16:0;2O/36:2;O** | C58H111NO9 | HexCer_HDS | TRUE |
| **HexCer 19:0;2O/38:6** | C63H113NO8 | HexCer_NDS | TRUE |
| **Cer 25:1;3O/38:5;(2OH)** | C63H115NO5 | Cer_AP | TRUE |
| **SMGDG O-21:4_28:7** | C58H92O12S | EtherSMGDG | TRUE |
| **PS 28:0_22:3;2O** | C56H104NO12P | OxPS | TRUE |
| **PS O-26:1_28:7** | C60H104NO9P | EtherPS | FALSE |
| **HexCer 16:0;2O/36:1;O** | C58H113NO9 | HexCer_HDS | TRUE |
| **PG 19:5_36:10** | C61H91O10P | PG | TRUE |
| **PI 20:4_26:0;1O** | C55H99O14P | OxPI | TRUE |
| **PI O-20:0_28:3** | C57H107O12P | EtherPI | TRUE |
| **ASG 29:1;O;Hex;FA 28:7** | C63H100O7 | AHexSIS | TRUE |
| **SM 52:2;3O** | C57H113N2O7P | SM | TRUE |
| **PS 28:0_20:4;4O** | C54H98NO14P | OxPS | TRUE |
| **HexCer 16:2;3O/36:6;(2OH)** | C58H99NO10 | HexCer_AP | TRUE |
| **Cer 28:1;3O/38:1;(2OH)** | C66H129NO5 | Cer_AP | TRUE |
| **PG O-28:7_28:7** | C62H97O9P | EtherPG | TRUE |
| **HexCer 22:1;3O/30:6;(2OH)** | C58H101NO10 | HexCer_AP | TRUE |
| **PI 24:0_20:4;3O** | C53H95O16P | OxPI | TRUE |
| **PC O-24:6_28:7** | C60H96NO7P | EtherPC | TRUE |
| **Hex2Cer 42:1;2O** | C54H103NO13 | Hex2Cer | TRUE |
| **PI 22:5_22:6;4O** | C53H81O17P | OxPI | TRUE |
| **PC 15:2_36:10** | C59H94NO8P | PC | TRUE |
| **SHexCer 52:6;2O** | C58H103NO11S | SHexCer | TRUE |
| **PC O-22:6_28:0;1O** | C58H106NO8P | EtherOxPC | TRUE |
| **HexCer 17:1;3O/38:2;(2OH)** | C61H115NO10 | HexCer_AP | TRUE |
| **SMGDG O-22:6_28:7** | C59H90O12S | EtherSMGDG | TRUE |
| **PG 17:1_38:10** | C61H99O10P | PG | TRUE |
| **Hex2Cer 43:6;2O** | C55H95NO13 | Hex2Cer | TRUE |
| **PC 13:1_38:10** | C59H96NO8P | PC | TRUE |
| **HexCer 16:1;3O/36:3;(2OH)** | C58H107NO10 | HexCer_AP | TRUE |
| **SMGDG O-22:5_28:7** | C59H92O12S | EtherSMGDG | TRUE |
| **Hex3Cer 31:2;2O** | C49H89NO18 | Hex3Cer | TRUE |
| **PE 19:0_38:8** | C62H108NO8P | PE | TRUE |
| **PC O-22:4_28:0;1O** | C58H110NO8P | EtherOxPC | TRUE |
| **HexCer 16:1;3O/36:2;(2OH)** | C58H109NO10 | HexCer_AP | TRUE |
| **PI 24:0_22:6;2O** | C55H95O15P | OxPI | TRUE |
| **PG 16:0_38:2** | C60H115O10P | PG | FALSE |
| **ASG 29:1;O;Hex;FA 28:1** | C63H112O7 | AHexSIS | TRUE |
| **Hex3Cer 31:1;2O** | C49H91NO18 | Hex3Cer | TRUE |
| **PE 19:0_38:7** | C62H110NO8P | PE | TRUE |
| **HexCer 16:1;3O/36:1;(2OH)** | C58H111NO10 | HexCer_AP | TRUE |
| **PC O-23:0_28:2** | C59H116NO7P | EtherPC | FALSE |
| **PI 24:0_22:5;2O** | C55H97O15P | OxPI | TRUE |
| **PI O-22:3_28:7** | C59H97O12P | EtherPI | TRUE |
| **PG 16:0_38:1** | C60H117O10P | PG | TRUE |
| **SM 53:2;3O** | C58H115N2O7P | SM | TRUE |
| **HexCer 16:1;3O/36:0;(2OH)** | C58H113NO10 | HexCer_AP | TRUE |
| **PI 22:3_22:3;4O** | C53H91O17P | OxPI | TRUE |
| **SM 17:1;2O/38:6** | C60H109N2O6P | SM | FALSE |
| **Hex2Cer 43:2;2O** | C55H103NO13 | Hex2Cer | TRUE |
| **PE 19:0_38:5** | C62H114NO8P | PE | FALSE |
| **Hex2Cer 43:1;2O** | C55H105NO13 | Hex2Cer | TRUE |
| **PC 16:2_36:10** | C60H96NO8P | PC | TRUE |
| **PC 22:6_28:0;1O** | C58H104NO9P | OxPC | TRUE |
| **HexCer 16:0;2O/38:4;O** | C60H111NO9 | HexCer_HDS | FALSE |
| **PG O-28:0_28:4** | C62H117O9P | EtherPG | TRUE |
| **PC 14:1_38:10** | C60H98NO8P | PC | TRUE |
| **HexCer 21:1;2O/38:6** | C65H115NO8 | HexCer_NS | TRUE |
| **Cer 27:1;3O/38:6;(2OH)** | C65H117NO5 | Cer_AP | TRUE |
| **PG O-28:0_28:3** | C62H119O9P | EtherPG | TRUE |
| **PS O-28:2_28:7** | C62H106NO9P | EtherPS | TRUE |
| **HexCer 16:0;2O/38:2;O** | C60H115NO9 | HexCer_HDS | TRUE |
| **HexCer 21:0;2O/38:6** | C65H117NO8 | HexCer_NDS | TRUE |
| **SM 18:3;2O/38:6** | C61H107N2O6P | SM | TRUE |
| **SM 54:3;3O** | C59H115N2O7P | SM | TRUE |
| **PE 20:0_38:7** | C63H112NO8P | PE | TRUE |
| **PI 20:4_28:0;1O** | C57H103O14P | OxPI | TRUE |
| **SM 54:2;3O** | C59H117N2O7P | SM | TRUE |
| **SHexCer 53:2;2O** | C59H113NO11S | SHexCer | TRUE |
| **Hex2Cer 44:2;2O** | C56H105NO13 | Hex2Cer | TRUE |
| **PI-Cer 16:1;2O/36:1;O** | C58H112NO12P | PI_Cer | TRUE |
| **Hex2Cer 44:1;2O** | C56H107NO13 | Hex2Cer | TRUE |
| **PC 14:0_38:6** | C60H108NO8P | PC | TRUE |
| **PI-Cer 18:1;2O/36:5** | C60H108NO11P | PI_Cer | TRUE |
| **PE 21:0_38:10** | C64H108NO8P | PE | TRUE |
| **HexCer 16:1;3O/38:4;(2OH)** | C60H109NO10 | HexCer_AP | TRUE |
| **SHexCer 52:0;3O** | C58H115NO12S | SHexCer | TRUE |
| **SMGDG O-24:6_28:7** | C61H94O12S | EtherSMGDG | TRUE |
| **PG 18:0_38:4** | C62H115O10P | PG | TRUE |
| **PE 22:6_38:10** | C65H98NO8P | PE | TRUE |
| **PE 21:0_38:9** | C64H110NO8P | PE | TRUE |
| **HexCer 16:1;3O/38:3;(2OH)** | C60H111NO10 | HexCer_AP | TRUE |
| **PE 22:5_38:10** | C65H100NO8P | PE | TRUE |
| **SHexCer 54:4;2O** | C60H111NO11S | SHexCer | TRUE |
| **HexCer 16:1;3O/38:2;(2OH)** | C60H113NO10 | HexCer_AP | TRUE |
| **SM 19:3;2O/38:6** | C62H109N2O6P | SM | TRUE |
| **PG 18:0_38:2** | C62H119O10P | PG | TRUE |
| **PC 26:0_22:6;4O** | C56H100NO12P | OxPC | TRUE |
| **PC O-26:2_28:7** | C62H108NO7P | EtherPC | FALSE |
| **PI-Cer 18:1;2O/36:2** | C60H114NO11P | PI_Cer | TRUE |
| **SHexCer 54:3;2O** | C60H113NO11S | SHexCer | TRUE |
| **PC 14:0_38:2** | C60H116NO8P | PC | TRUE |
| **HexCer 16:1;3O/38:1;(2OH)** | C60H115NO10 | HexCer_AP | TRUE |
| **HexCer 22:0;2O/38:5** | C66H121NO8 | HexCer_NDS | TRUE |
| **PI O-24:3_28:7** | C61H101O12P | EtherPI | TRUE |
| **SM 55:2;3O** | C60H119N2O7P | SM | TRUE |
| **PE 21:0_38:6** | C64H116NO8P | PE | TRUE |
| **Cer 30:0;2O/38:2** | C68H133NO3 | Cer_NDS | TRUE |
| **PC O-28:0_22:3;3O** | C58H112NO10P | EtherOxPC | TRUE |
| **HexCer 18:0;2O/38:6;O** | C62H111NO9 | HexCer_HDS | TRUE |
| **PE 21:0_38:5** | C64H118NO8P | PE | TRUE |
| **PI O-23:0_28:1** | C60H117O12P | EtherPI | FALSE |
| **SHexCer 54:0;2O** | C60H119NO11S | SHexCer | TRUE |
| **PI O-23:0_28:0** | C60H119O12P | EtherPI | TRUE |
| **HexCer 18:0;2O/38:4;O** | C62H115NO9 | HexCer_HDS | TRUE |
| **Cer 29:2;3O/38:6;(2OH)** | C67H119NO5 | Cer_AP | TRUE |
| **PI 24:0_22:3;4O** | C55H101O17P | OxPI | TRUE |
| **SM 56:5;3O** | C61H115N2O7P | SM | TRUE |
| **PI-Cer 19:1;2O/36:4** | C61H112NO11P | PI_Cer | TRUE |
| **PC 15:0_38:4** | C61H114NO8P | PC | TRUE |
| **Cer 29:1;3O/38:6;(2OH)** | C67H121NO5 | Cer_AP | TRUE |
| **PI-Cer 19:1;2O/36:3** | C61H114NO11P | PI_Cer | TRUE |
| **SHexCer 55:4;2O** | C61H113NO11S | SHexCer | TRUE |
| **PE 22:0_38:8** | C65H114NO8P | PE | TRUE |
| **Cer 29:1;3O/38:5;(2OH)** | C67H123NO5 | Cer_AP | TRUE |
| **PI-Cer 18:1;2O/36:3;O** | C60H112NO12P | PI_Cer | TRUE |
| **HexCer 19:2;2O/38:6;O** | C63H109NO9 | HexCer_HS | TRUE |
| **PE 22:0_38:7** | C65H116NO8P | PE | TRUE |
| **HexCer 18:0;2O/38:1;O** | C62H121NO9 | HexCer_HDS | TRUE |
| **PI 22:4_28:0;1O** | C59H107O14P | OxPI | TRUE |
| **SM 20:2;2O/38:6** | C63H113N2O6P | SM | TRUE |
| **HexCer 18:2;3O/38:6;(2OH)** | C62H107NO10 | HexCer_AP | TRUE |
| **PI-Cer 18:1;2O/36:2;O** | C60H114NO12P | PI_Cer | TRUE |
| **SHexCer 54:3;3O** | C60H113NO12S | SHexCer | TRUE |
| **PE 22:0_38:6** | C65H118NO8P | PE | TRUE |
| **SHexCer 54:2;3O** | C60H115NO12S | SHexCer | TRUE |
| **PG 20:0_38:6** | C64H115O10P | PG | FALSE |
| **SHexCer 56:7;2O** | C62H109NO11S | SHexCer | TRUE |
| **PI-Cer 20:1;2O/36:6** | C62H110NO11P | PI_Cer | TRUE |
| **HexCer 18:1;3O/38:5;(2OH)** | C62H111NO10 | HexCer_AP | TRUE |
| **SHexCer 54:1;3O** | C60H117NO12S | SHexCer | TRUE |
| **PI-Cer 18:1;2O/36:0;O** | C60H118NO12P | PI_Cer | TRUE |
| **SHexCer 56:6;2O** | C62H111NO11S | SHexCer | TRUE |
| **PI-Cer 20:1;2O/36:5** | C62H112NO11P | PI_Cer | TRUE |
| **PE 23:0_38:10** | C66H112NO8P | PE | TRUE |
| **PC 16:0_38:5** | C62H114NO8P | PC | TRUE |
| **SHexCer 54:0;3O** | C60H119NO12S | SHexCer | TRUE |
| **HexCer 18:1;3O/38:4;(2OH)** | C62H113NO10 | HexCer_AP | TRUE |
| **Cer 30:2;3O/38:6;(2OH)** | C68H121NO5 | Cer_AP | TRUE |
| **SHexCer 56:5;2O** | C62H113NO11S | SHexCer | TRUE |
| **PE 23:0_38:9** | C66H114NO8P | PE | TRUE |
| **PC 16:0_38:4** | C62H116NO8P | PC | TRUE |
| **PE 30:1_30:1** | C65H126NO8P | PE | TRUE |
| **Cer 30:1;3O/38:6;(2OH)** | C68H123NO5 | Cer_AP | TRUE |
| **PG 20:0_38:3** | C64H121O10P | PG | TRUE |
| **SHexCer 56:4;2O** | C62H115NO11S | SHexCer | TRUE |
| **PI-Cer 20:1;2O/36:3** | C62H116NO11P | PI_Cer | TRUE |
| **HexCer 18:1;3O/38:2;(2OH)** | C62H117NO10 | HexCer_AP | TRUE |
| **HexCer 19:0;2O/38:2;O** | C63H121NO9 | HexCer_HDS | TRUE |
| **SMGDG O-25:0_28:4** | C62H114O12S | EtherSMGDG | FALSE |
| **PG 20:0_38:2** | C64H123O10P | PG | TRUE |
| **SM 20:1;2O/38:1** | C63H125N2O6P | SM | TRUE |
| **PC O-28:2_28:7** | C64H112NO7P | EtherPC | FALSE |
| **SHexCer 56:3;2O** | C62H117NO11S | SHexCer | TRUE |
| **PI-Cer 20:1;2O/36:2** | C62H118NO11P | PI_Cer | TRUE |
| **PE 23:0_38:7** | C66H118NO8P | PE | TRUE |
| **HexCer 18:1;3O/38:1;(2OH)** | C62H119NO10 | HexCer_AP | TRUE |
| **HexCer 19:0;2O/38:1;O** | C63H123NO9 | HexCer_HDS | TRUE |
| **SM 21:2;2O/38:6** | C64H115N2O6P | SM | FALSE |
| **PG 20:0_38:1** | C64H125O10P | PG | FALSE |
| **Hex2Cer 47:3;2O** | C59H109NO13 | Hex2Cer | TRUE |
| **SHexCer 55:3;3O** | C61H115NO12S | SHexCer | TRUE |
| **PI-Cer 19:1;2O/36:2;O** | C61H116NO12P | PI_Cer | TRUE |
| **PC O-28:1_28:7** | C64H114NO7P | EtherPC | TRUE |
| **PE 23:0_38:6** | C66H120NO8P | PE | TRUE |
| **SM 58:8;3O** | C63H113N2O7P | SM | FALSE |
| **SMGDG O-25:0_28:2** | C62H118O12S | EtherSMGDG | TRUE |
| **PI O-25:0_28:2** | C62H119O12P | EtherPI | TRUE |
| **PC O-28:0_28:7** | C64H116NO7P | EtherPC | TRUE |
| **SHexCer 56:1;2O** | C62H121NO11S | SHexCer | TRUE |
| **SMGDG O-25:0_28:1** | C62H120O12S | EtherSMGDG | TRUE |
| **Hex2Cer 47:1;2O** | C59H113NO13 | Hex2Cer | TRUE |
| **PC O-26:0_28:0;1O** | C62H126NO8P | EtherOxPC | TRUE |
| **PG 22:2_38:10** | C66H107O10P | PG | TRUE |
| **HexCer 20:0;2O/38:4;O** | C64H119NO9 | HexCer_HDS | TRUE |
| **PE 23:0_38:3** | C66H126NO8P | PE | TRUE |
| **HexCer 22:1;3O/38:2;(2OH)** | C66H125NO10 | HexCer_AP | TRUE |
| **Cer 19:0;2O/28:1;(3OH)(FA 22:6)** | C69H123NO5 | Cer_EBDS | TRUE |
| **PI 26:0_22:3;4O** | C57H105O17P | OxPI | TRUE |
| **PI-Cer 20:1;2O/36:5;O** | C62H112NO12P | PI_Cer | TRUE |
| **HexCer 20:0;2O/38:3;O** | C64H121NO9 | HexCer_HDS | TRUE |
| **PE 23:0_38:2** | C66H128NO8P | PE | TRUE |
| **Cer 13:0;2O/28:6;(3OH)(FA 28:0)** | C69H125NO5 | Cer_EBDS | TRUE |
| **SHexCer 56:5;3O** | C62H113NO12S | SHexCer | TRUE |
| **HexCer 21:3;2O/38:6;O** | C65H111NO9 | HexCer_HS | TRUE |
| **HexCer 19:1;3O/38:2;(2OH)** | C63H119NO10 | HexCer_AP | TRUE |
| **HexCer 20:0;2O/38:2;O** | C64H123NO9 | HexCer_HDS | TRUE |
| **PE 23:0_38:1** | C66H130NO8P | PE | TRUE |
| **Cer 13:0;2O/28:5;(3OH)(FA 28:0)** | C69H127NO5 | Cer_EBDS | TRUE |
| **SM 22:3;2O/38:6** | C65H115N2O6P | SM | TRUE |
| **SM 21:1;2O/38:1** | C64H127N2O6P | SM | TRUE |
| **PS 22:5_38:10** | C66H100NO10P | PS | TRUE |
| **SHexCer 56:4;3O** | C62H115NO12S | SHexCer | TRUE |
| **PI-Cer 20:1;2O/36:3;O** | C62H116NO12P | PI_Cer | TRUE |
| **PE 24:0_38:7** | C67H120NO8P | PE | TRUE |
| **HexCer 19:1;3O/38:1;(2OH)** | C63H121NO10 | HexCer_AP | TRUE |
| **HexCer 20:0;2O/38:1;O** | C64H125NO9 | HexCer_HDS | TRUE |
| **PG 21:0_38:1** | C65H127O10P | PG | FALSE |
| **PC 18:0_38:8** | C64H112NO8P | PC | FALSE |
| **SHexCer 56:3;3O** | C62H117NO12S | SHexCer | TRUE |
| **PG 22:0_38:7** | C66H117O10P | PG | TRUE |
| **SM 59:8;3O** | C64H115N2O7P | SM | FALSE |
| **PI-Cer 22:2;2O/36:6** | C64H112NO11P | PI_Cer | TRUE |
| **SHexCer 56:2;3O** | C62H119NO12S | SHexCer | TRUE |
| **HexCer 21:0;2O/38:6;O** | C65H117NO9 | HexCer_HDS | TRUE |
| **PE 24:0_38:5** | C67H124NO8P | PE | TRUE |
| **PG 22:0_38:6** | C66H119O10P | PG | FALSE |
| **HexCer 20:1;3O/38:5;(2OH)** | C64H115NO10 | HexCer_AP | TRUE |
| **PI-Cer 20:1;2O/36:0;O** | C62H122NO12P | PI_Cer | TRUE |
| **HexCer 22:0;2O/38:4** | C66H123NO8 | HexCer_NDS | TRUE |
| **Cer 13:0;2O/28:1;(3OH)(FA 28:0)** | C69H135NO5 | Cer_EBDS | TRUE |
| **PI 26:0_22:5;4O** | C57H101O18P | OxPI | TRUE |
| **SMGDG O-27:0_28:7** | C64H112O12S | EtherSMGDG | TRUE |
| **PI-Cer 22:1;2O/36:5** | C64H116NO11P | PI_Cer | TRUE |
| **PE 25:0_38:10** | C68H116NO8P | PE | TRUE |
| **PC 18:0_38:5** | C64H118NO8P | PC | TRUE |
| **SHexCer 56:0;3O** | C62H123NO12S | SHexCer | TRUE |
| **PI-Cer 20:0;2O/36:0;O** | C62H124NO12P | PI_Cer | FALSE |
| **Cer 20:0;2O/28:1;(3OH)(FA 22:6)** | C70H125NO5 | Cer_EBDS | TRUE |
| **PI 24:0_28:0;1O** | C61H119O14P | OxPI | TRUE |
| **PE 25:0_38:9** | C68H118NO8P | PE | TRUE |
| **HexCer 20:1;3O/38:3;(2OH)** | C64H119NO10 | HexCer_AP | TRUE |
| **Cer 14:0;2O/28:6;(3OH)(FA 28:0)** | C70H127NO5 | Cer_EBDS | TRUE |
| **PG 22:0_38:3** | C66H125O10P | PG | FALSE |
| **PI-Cer 22:1;2O/36:3** | C64H120NO11P | PI_Cer | TRUE |
| **HexCer 20:1;3O/38:2;(2OH)** | C64H121NO10 | HexCer_AP | TRUE |
| **PE 24:0_38:1** | C67H132NO8P | PE | TRUE |
| **PI O-27:0_28:4** | C64H119O12P | EtherPI | TRUE |
| **SM 23:3;2O/38:6** | C66H117N2O6P | SM | FALSE |
| **SM 22:1;2O/38:1** | C65H129N2O6P | SM | TRUE |
| **Hex2Cer 49:4;2O** | C61H111NO13 | Hex2Cer | TRUE |
| **PI-Cer 21:1;2O/36:3;O** | C63H118NO12P | PI_Cer | TRUE |
| **HexCer 20:1;3O/38:1;(2OH)** | C64H123NO10 | HexCer_AP | TRUE |
| **HexCer 21:0;2O/38:1;O** | C65H127NO9 | HexCer_HDS | TRUE |
| **SM 60:9;3O** | C65H115N2O7P | SM | TRUE |
| **PG 22:0_38:1** | C66H129O10P | PG | TRUE |
| **Hex2Cer 49:3;2O** | C61H113NO13 | Hex2Cer | TRUE |
| **PC 18:0_38:1** | C64H126NO8P | PC | TRUE |
| **PE 25:0_38:6** | C68H124NO8P | PE | TRUE |
| **PI 16:0_38:2** | C63H119O13P | PI | FALSE |
| **PI O-27:0_28:2** | C64H123O12P | EtherPI | TRUE |
| **SM 23:1;2O/38:6** | C66H121N2O6P | SM | TRUE |
| **PI-Cer 22:3;2O/36:6;O** | C64H110NO12P | PI_Cer | TRUE |
| **Hex2Cer 49:2;2O** | C61H115NO13 | Hex2Cer | TRUE |
| **PI-Cer 21:1;2O/36:1;O** | C63H122NO12P | PI_Cer | TRUE |
| **HexCer 21:1;3O/38:6;(2OH)** | C65H115NO10 | HexCer_AP | TRUE |
| **SMGDG O-28:1_28:7** | C65H112O12S | EtherSMGDG | TRUE |
| **PI 16:0_38:1** | C63H121O13P | PI | TRUE |
| **Hex2Cer 50:8;2O** | C62H105NO13 | Hex2Cer | TRUE |
| **PI-Cer 22:2;2O/36:6;O** | C64H112NO12P | PI_Cer | TRUE |
| **PE 26:1_38:10** | C69H116NO8P | PE | TRUE |
| **Hex2Cer 49:1;2O** | C61H117NO13 | Hex2Cer | TRUE |
| **PC O-28:0_28:0;1O** | C64H130NO8P | EtherOxPC | FALSE |
| **PG 24:2_38:10** | C68H111O10P | PG | TRUE |
| **PI-Cer 22:1;2O/36:6;O** | C64H114NO12P | PI_Cer | TRUE |
| **Hex2Cer 49:0;2O** | C61H119NO13 | Hex2Cer | TRUE |
| **HexCer 22:0;2O/38:4;O** | C66H123NO9 | HexCer_HDS | TRUE |
| **PE 25:0_38:3** | C68H130NO8P | PE | TRUE |
| **SMGDG O-28:0_28:6** | C65H116O12S | EtherSMGDG | FALSE |
| **SM 23:1;2O/38:3** | C66H127N2O6P | SM | TRUE |
| **PI-Cer 22:1;2O/36:5;O** | C64H116NO12P | PI_Cer | TRUE |
| **PE 26:0_38:9** | C69H120NO8P | PE | TRUE |
| **HexCer 22:0;2O/38:3;O** | C66H125NO9 | HexCer_HDS | TRUE |
| **PE 25:0_38:2** | C68H132NO8P | PE | TRUE |
| **PI-Cer 22:1;2O/36:4;O** | C64H118NO12P | PI_Cer | TRUE |
| **HexCer 21:1;3O/38:2;(2OH)** | C65H123NO10 | HexCer_AP | TRUE |
| **HexCer 22:0;2O/38:2;O** | C66H127NO9 | HexCer_HDS | TRUE |
| **SMGDG O-28:0_28:4** | C65H120O12S | EtherSMGDG | TRUE |
| **SM 24:3;2O/38:6** | C67H119N2O6P | SM | TRUE |
| **Hex2Cer 50:4;2O** | C62H113NO13 | Hex2Cer | TRUE |
| **PC 28:1_30:8** | C66H114NO8P | PC | TRUE |
| **PI-Cer 22:1;2O/36:3;O** | C64H120NO12P | PI_Cer | TRUE |
| **HexCer 21:1;3O/38:1;(2OH)** | C65H125NO10 | HexCer_AP | TRUE |
| **HexCer 22:0;2O/38:1;O** | C66H129NO9 | HexCer_HDS | TRUE |
| **PI 17:0_38:3** | C64H119O13P | PI | TRUE |
| **SM 61:9;3O** | C66H117N2O7P | SM | TRUE |
| **SM 24:2;2O/38:6** | C67H121N2O6P | SM | TRUE |
| **PG 23:0_38:1** | C67H131O10P | PG | TRUE |
| **PI-Cer 22:1;2O/36:2;O** | C64H122NO12P | PI_Cer | TRUE |
| **SMGDG O-28:0_28:2** | C65H124O12S | EtherSMGDG | TRUE |
| **SM 24:1;2O/38:6** | C67H123N2O6P | SM | TRUE |
| **PI-Cer 22:1;2O/36:1;O** | C64H124NO12P | PI_Cer | TRUE |
| **PE 26:0_38:5** | C69H128NO8P | PE | TRUE |
| **PE 27:1_38:10** | C70H118NO8P | PE | TRUE |
| **Hex2Cer 50:1;2O** | C62H119NO13 | Hex2Cer | TRUE |
| **PC 20:0_38:6** | C66H120NO8P | PC | TRUE |
| **PI 17:0_38:0** | C64H125O13P | PI | FALSE |
| **PE 27:0_38:10** | C70H120NO8P | PE | FALSE |
| **PC 34:2_24:3** | C66H122NO8P | PC | TRUE |
| **PI-Cer 22:0;2O/36:0;O** | C64H128NO12P | PI_Cer | TRUE |
| **SM 24:1;2O/38:3** | C67H129N2O6P | SM | FALSE |
| **HexCer 22:1;3O/38:3;(2OH)** | C66H123NO10 | HexCer_AP | TRUE |
| **PE 27:0_38:8** | C70H124NO8P | PE | TRUE |
| **PG 25:0_38:9** | C69H119O10P | PG | TRUE |
| **PC 21:0_38:9** | C67H116NO8P | PC | TRUE |
| **PE 27:0_38:7** | C70H126NO8P | PE | TRUE |
| **PC 20:0_38:2** | C66H128NO8P | PC | TRUE |
| **HexCer 22:1;3O/38:1;(2OH)** | C66H127NO10 | HexCer_AP | TRUE |
| **PG 25:0_38:8** | C69H121O10P | PG | TRUE |
| **SM 25:2;2O/38:6** | C68H123N2O6P | SM | TRUE |
| **PG 24:0_38:1** | C68H133O10P | PG | TRUE |
| **PC 21:0_38:8** | C67H118NO8P | PC | FALSE |
| **PC 20:0_38:1** | C66H130NO8P | PC | TRUE |
| **PE 27:0_38:6** | C70H128NO8P | PE | TRUE |
| **PE 28:2_38:10** | C71H118NO8P | PE | TRUE |
| **PG 26:3_38:10** | C70H113O10P | PG | TRUE |
| **PG 25:0_38:6** | C69H125O10P | PG | TRUE |
| **PC 22:3_38:10** | C68H110NO8P | PC | TRUE |
| **PE 28:1_38:10** | C71H120NO8P | PE | TRUE |
| **Hex2Cer 51:1;2O** | C63H121NO13 | Hex2Cer | TRUE |
| **Cer 16:0;2O/28:1;(3OH)(FA 28:0)** | C72H141NO5 | Cer_EBDS | TRUE |
| **PG 26:2_38:10** | C70H115O10P | PG | FALSE |
| **PC 21:0_38:4** | C67H126NO8P | PC | TRUE |
| **PG 26:0_38:10** | C70H119O10P | PG | TRUE |
| **PG 25:0_38:3** | C69H131O10P | PG | FALSE |
| **PE 28:0_38:8** | C71H126NO8P | PE | TRUE |
| **Hex2Cer 52:4;2O** | C64H117NO13 | Hex2Cer | TRUE |
| **PE 28:0_38:7** | C71H128NO8P | PE | TRUE |
| **PI 19:0_38:3** | C66H123O13P | PI | TRUE |
| **PG 26:0_38:8** | C70H123O10P | PG | FALSE |
| **SM 26:2;2O/38:6** | C69H125N2O6P | SM | TRUE |
| **Hex2Cer 52:3;2O** | C64H119NO13 | Hex2Cer | TRUE |
| **SM 26:1;2O/38:6** | C69H127N2O6P | SM | TRUE |
| **Hex2Cer 52:2;2O** | C64H121NO13 | Hex2Cer | TRUE |
| **PE 28:0_38:5** | C71H132NO8P | PE | TRUE |
| **PI 19:0_38:1** | C66H127O13P | PI | TRUE |
| **PE 29:1_38:10** | C72H122NO8P | PE | FALSE |
| **Hex2Cer 52:1;2O** | C64H123NO13 | Hex2Cer | TRUE |
| **PC 60:6** | C68H124NO8P | PC | TRUE |
| **Cer 17:0;2O/28:1;(3OH)(FA 28:0)** | C73H143NO5 | Cer_EBDS | TRUE |
| **PI 21:4_38:10** | C68H105O13P | PI | TRUE |
| **PI 19:0_38:0** | C66H129O13P | PI | TRUE |
| **PE 29:0_38:10** | C72H124NO8P | PE | TRUE |
| **PC 38:1_22:4** | C68H126NO8P | PC | TRUE |
| **PG 30:8_36:10** | C72H107O10P | PG | TRUE |
| **Hex3Cer 41:3;2O** | C59H107NO18 | Hex3Cer | TRUE |
| **PE 29:0_38:9** | C72H126NO8P | PE | TRUE |
| **PC 22:0_38:4** | C68H128NO8P | PC | TRUE |
| **PG 28:7_38:10** | C72H109O10P | PG | TRUE |
| **PE 29:0_38:8** | C72H128NO8P | PE | TRUE |
| **PI 21:1_38:10** | C68H111O13P | PI | TRUE |
| **PG 27:0_38:9** | C71H123O10P | PG | TRUE |
| **PG 26:0_38:2** | C70H135O10P | PG | TRUE |
| **PC 23:0_38:9** | C69H120NO8P | PC | TRUE |
| **PE 29:0_38:7** | C72H130NO8P | PE | TRUE |
| **PC 22:0_38:2** | C68H132NO8P | PC | TRUE |
| **Hex2Cer 53:3;2O** | C65H121NO13 | Hex2Cer | TRUE |
| **PC 22:0_38:1** | C68H134NO8P | PC | TRUE |
| **Hex2Cer 53:2;2O** | C65H123NO13 | Hex2Cer | TRUE |
| **PC 24:3_38:10** | C70H114NO8P | PC | TRUE |
| **PE 30:1_38:10** | C73H124NO8P | PE | TRUE |
| **Pentaerythritol tetrakis(3,5-di-tert-butyl-4-hydroxyhydrocinnamate)** | C73H108O12 | Others | TRUE |
| **GM3 36:2;2O** | C59H106N2O21 | GM3 | TRUE |
| **GM3 36:1;2O** | C59H108N2O21 | GM3 | TRUE |
| **PC 24:0_38:9** | C70H122NO8P | PC | TRUE |
| **Hex2Cer 54:3;2O** | C66H123NO13 | Hex2Cer | TRUE |
| **Hex2Cer 54:2;2O** | C66H125NO13 | Hex2Cer | TRUE |
| **PE 31:1_38:10** | C74H126NO8P | PE | TRUE |
| **Hex2Cer 54:1;2O** | C66H127NO13 | Hex2Cer | TRUE |
| **PG 32:9_36:10** | C74H109O10P | PG | TRUE |
| **Hex2Cer 54:0;2O** | C66H129NO13 | Hex2Cer | TRUE |
| **PG 30:8_38:10** | C74H111O10P | PG | TRUE |
| **PC 24:0_38:4** | C70H132NO8P | PC | TRUE |
| **PG 30:7_38:10** | C74H113O10P | PG | TRUE |
| **PG 29:0_38:10** | C73H125O10P | PG | TRUE |
| **PE 31:0_38:8** | C74H132NO8P | PE | TRUE |
| **SM 29:3;2O/38:6** | C72H129N2O6P | SM | TRUE |
| **PE 31:0_38:7** | C74H134NO8P | PE | TRUE |
| **Hex2Cer 55:3;2O** | C67H125NO13 | Hex2Cer | TRUE |
| **PE 32:2_38:10** | C75H126NO8P | PE | TRUE |
| **PE 31:0_38:3** | C74H142NO8P | PE | TRUE |
| **PC 26:0_38:10** | C72H124NO8P | PC | TRUE |
| **GM3 38:1;2O** | C61H112N2O21 | GM3 | TRUE |
| **Hex2Cer 56:4;2O** | C68H125NO13 | Hex2Cer | TRUE |
| **PI 24:0_38:9** | C71H121O13P | PI | TRUE |
| **PS 30:3_38:10** | C74H120NO10P | PS | TRUE |
| **SM 30:1;2O/38:1** | C73H145N2O6P | SM | FALSE |
| **PE 34:4_38:10** | C77H126NO8P | PE | TRUE |
| **PE 32:0_38:0** | C75H150NO8P | PE | TRUE |
| **PE 34:3_38:10** | C77H128NO8P | PE | TRUE |
| **PE 34:2_38:10** | C77H130NO8P | PE | TRUE |
| **PI 26:2_38:10** | C73H119O13P | PI | TRUE |
| **Hex2Cer 58:5;2O** | C70H127NO13 | Hex2Cer | TRUE |
| **PI 26:1_38:10** | C73H121O13P | PI | FALSE |
| **PG 31:0_38:2** | C75H145O10P | PG | TRUE |
| **Hex3Cer 46:0;2O** | C64H123NO18 | Hex3Cer | TRUE |
| **SM 68:1;3O** | C73H147N2O7P | SM | TRUE |
| **PE 35:1_38:10** | C78H134NO8P | PE | TRUE |
| **PG 34:7_38:10** | C78H121O10P | PG | TRUE |
| **PE 36:5_38:10** | C79H128NO8P | PE | TRUE |
| **PI 30:8_36:10** | C75H111O13P | PI | TRUE |
| **PE 36:3_38:10** | C79H132NO8P | PE | TRUE |
| **PI 28:4_38:10** | C75H119O13P | PI | TRUE |
| **Hex3Cer 48:4;2O** | C66H119NO18 | Hex3Cer | TRUE |
| **PS 34:8_38:10** | C78H118NO10P | PS | TRUE |
| **PI 28:3_38:10** | C75H121O13P | PI | TRUE |
| **GM3 42:2;2O** | C65H118N2O21 | GM3 | TRUE |
| **PS 34:6_38:10** | C78H122NO10P | PS | TRUE |
| **PG 33:0_38:2** | C77H149O10P | PG | TRUE |
| **PI 27:0_38:3** | C74H139O13P | PI | TRUE |
| **PG 34:0_38:8** | C78H139O10P | PG | TRUE |
| **PE 38:7_38:10** | C81H128NO8P | PE | TRUE |
| **PG 36:8_38:10** | C80H123O10P | PG | TRUE |
| **PI 28:0_38:6** | C75H135O13P | PI | TRUE |
| **PE 38:5_38:10** | C81H132NO8P | PE | TRUE |
| **Hex3Cer 50:7;2O** | C68H117NO18 | Hex3Cer | TRUE |
| **PI 30:6_38:10** | C77H119O13P | PI | TRUE |
| **PI 29:0_38:9** | C76H131O13P | PI | TRUE |
| **PS 36:10_38:10** | C80H118NO10P | PS | TRUE |
| **PI 30:5_38:10** | C77H121O13P | PI | TRUE |
| **PS 36:8_38:10** | C80H122NO10P | PS | TRUE |
| **PS 36:7_38:10** | C80H124NO10P | PS | TRUE |
| **PI 30:2_38:10** | C77H127O13P | PI | TRUE |
| **PI 32:8_38:10** | C79H119O13P | PI | TRUE |
| **GM3 45:1;2O** | C68H126N2O21 | GM3 | TRUE |
| **CL 12:0_14:1_18:0_17:1** | C70H132O17P2 | CL | TRUE |
| **Hex3Cer 52:7;2O** | C70H121NO18 | Hex3Cer | TRUE |
| **PC 34:7_38:8** | C80H130NO8P | PC | TRUE |
| **PI 32:6_38:10** | C79H123O13P | PI | TRUE |
| **PS 38:10_38:10** | C82H122NO10P | PS | TRUE |
| **PI 32:5_38:10** | C79H125O13P | PI | TRUE |
| **PI 32:4_38:10** | C79H127O13P | PI | TRUE |
| **PS 38:7_38:10** | C82H128NO10P | PS | TRUE |
| **PI 32:2_38:10** | C79H131O13P | PI | TRUE |
| **PI 31:0_38:5** | C78H143O13P | PI | TRUE |
| **PI 32:1_38:10** | C79H133O13P | PI | TRUE |
| **PI 32:0_38:10** | C79H135O13P | PI | TRUE |
| **CL 12:0_12:0_17:0_22:5** | C72H130O17P2 | CL | TRUE |
| **PI 36:10_36:10** | C81H119O13P | PI | TRUE |
| **Hex3Cer 54:9;2O** | C72H121NO18 | Hex3Cer | TRUE |
| **PC 36:7_38:10** | C82H130NO8P | PC | TRUE |
| **PI 34:8_38:10** | C81H123O13P | PI | TRUE |
| **Hex3Cer 54:8;2O** | C72H123NO18 | Hex3Cer | TRUE |
| **Hex3Cer 54:7;2O** | C72H125NO18 | Hex3Cer | TRUE |
| **PI 34:6_38:10** | C81H127O13P | PI | TRUE |
| **Hex3Cer 54:6;2O** | C72H127NO18 | Hex3Cer | TRUE |
| **PI 34:4_38:10** | C81H131O13P | PI | TRUE |
| **Hex3Cer 54:4;2O** | C72H131NO18 | Hex3Cer | TRUE |
| **CL 12:0_12:0_24:0_16:2** | C73H138O17P2 | CL | TRUE |
| **PC 37:1_38:10** | C83H144NO8P | PC | TRUE |
| **PC 37:0_38:10** | C83H146NO8P | PC | TRUE |
| **PI 36:8_38:10** | C83H127O13P | PI | TRUE |
| **PI 34:0_38:3** | C81H153O13P | PI | TRUE |
| **PC 37:0_38:8** | C83H150NO8P | PC | TRUE |
| **PI 36:5_38:10** | C83H133O13P | PI | TRUE |
| **CL 12:0_12:0_24:0_18:3** | C75H140O17P2 | CL | TRUE |
| **CL 12:0_12:0_26:0_16:2** | C75H142O17P2 | CL | TRUE |
| **CL 15:1_17:1_15:1_20:5** | C76H132O17P2 | CL | TRUE |
| **PI 36:0_38:4** | C83H155O13P | PI | TRUE |
| **PI 36:0_38:3** | C83H157O13P | PI | TRUE |
| **PI 37:0_38:9** | C84H147O13P | PI | TRUE |
| **PI 38:4_38:10** | C85H139O13P | PI | TRUE |
| **CL 12:0_12:0_26:0_18:3** | C77H144O17P2 | CL | TRUE |
| **CL 12:0_12:0_28:0_16:2** | C77H146O17P2 | CL | TRUE |
| **PI 38:0_38:5** | C85H157O13P | PI | TRUE |
| **PI 38:0_38:4** | C85H159O13P | PI | TRUE |
| **CL 12:0_14:0_22:3_22:4** | C79H140O17P2 | CL | TRUE |
| **CL 12:0_12:0_24:0_22:6** | C79H142O17P2 | CL | TRUE |
| **CL 12:0_12:0_26:0_20:4** | C79H146O17P2 | CL | TRUE |
| **CL 12:0_12:0_28:0_18:2** | C79H150O17P2 | CL | TRUE |
| **CL 17:1_16:2_17:1_22:6** | C81H138O17P2 | CL | TRUE |
| **CL 12:0_16:0_22:3_22:6** | C81H140O17P2 | CL | TRUE |
| **CL 12:0_16:0_22:3_22:5** | C81H142O17P2 | CL | TRUE |
| **CL 12:0_14:1_24:0_22:6** | C81H144O17P2 | CL | TRUE |
| **CL 12:0_12:0_26:0_22:6** | C81H146O17P2 | CL | FALSE |
| **CL 12:0_12:0_28:0_20:5** | C81H148O17P2 | CL | TRUE |
| **CL 12:0_12:0_28:0_20:4** | C81H150O17P2 | CL | TRUE |
| **CL 12:0_12:0_28:0_20:3** | C81H152O17P2 | CL | TRUE |
| **CL 12:0_18:0_22:4_22:6** | C83H142O17P2 | CL | TRUE |
| **CL 12:0_18:0_22:3_22:6** | C83H144O17P2 | CL | TRUE |
| **CL 12:0_16:2_24:0_22:6** | C83H146O17P2 | CL | TRUE |
| **CL 12:0_14:1_26:0_22:6** | C83H148O17P2 | CL | TRUE |
| **CL 12:0_15:0_18:0_28:0** | C82H160O17P2 | CL | FALSE |
| **CL 12:0_12:0_28:0_22:6** | C83H150O17P2 | CL | TRUE |
| **CL 12:0_20:3_22:3_22:6** | C85H142O17P2 | CL | TRUE |
| **CL 12:0_15:0_28:0_20:5** | C84H154O17P2 | CL | TRUE |
| **CL 12:0_20:3_22:3_22:5** | C85H144O17P2 | CL | TRUE |
| **CL 15:0_18:1_24:0_18:3** | C84H156O17P2 | CL | TRUE |
| **CL 12:0_20:3_22:3_22:4** | C85H146O17P2 | CL | TRUE |
| **CL 12:0_15:0_28:0_20:3** | C84H158O17P2 | CL | TRUE |
| **CL 12:0_18:3_24:0_22:6** | C85H148O17P2 | CL | TRUE |
| **CL 12:0_16:2_26:0_22:6** | C85H150O17P2 | CL | TRUE |
| **CL 12:0_15:1_24:0_24:0** | C84H162O17P2 | CL | TRUE |
| **CL 17:0_18:1_17:0_24:0** | C85H164O17P2 | CL | TRUE |
| **CL 16:0_17:2_24:0_20:5** | C86H154O17P2 | CL | TRUE |
| **CL 12:0_24:0_16:0_24:0** | C85H166O17P2 | CL | TRUE |
| **CL 12:0_15:0_28:0_22:6** | C86H156O17P2 | CL | TRUE |
| **CL 12:0_15:0_28:0_22:4** | C86H160O17P2 | CL | TRUE |
| **CL 17:1_18:2_18:0_24:0** | C86H162O17P2 | CL | TRUE |
| **CL 12:0_24:0_17:0_24:0** | C86H168O17P2 | CL | TRUE |
| **CL 18:0_14:1_26:0_20:4** | C87H160O17P2 | CL | TRUE |
| **CL 12:0_16:0_28:0_22:3** | C87H164O17P2 | CL | TRUE |
| **CL 14:1_17:2_26:0_22:6** | C88H154O17P2 | CL | TRUE |
| **CL 16:0_16:0_28:0_18:2** | C87H166O17P2 | CL | TRUE |
| **CL 12:0_17:2_28:0_22:6** | C88H156O17P2 | CL | TRUE |
| **CL 12:0_17:1_28:0_22:6** | C88H158O17P2 | CL | TRUE |
| **CL 17:0_22:3_18:1_22:3** | C88H158O17P2 | CL | TRUE |
| **CL 17:0_22:3_18:0_22:3** | C88H160O17P2 | CL | TRUE |
| **CL 12:0_17:0_28:0_22:5** | C88H162O17P2 | CL | TRUE |
| **CL 18:2_18:3_22:3_22:3** | C89H152O17P2 | CL | TRUE |
| **CL 12:0_17:0_28:0_22:4** | C88H164O17P2 | CL | TRUE |
| **CL 18:1_18:2_24:0_20:4** | C89H160O17P2 | CL | TRUE |
| **CL 15:0_24:0_16:0_24:0** | C88H172O17P2 | CL | TRUE |
| **CL 14:0_24:0_17:0_24:0** | C88H172O17P2 | CL | TRUE |
| **CL 18:1_16:2_28:0_18:3** | C89H162O17P2 | CL | TRUE |
| **CL 16:0_24:0_18:0_22:5** | C89H164O17P2 | CL | TRUE |
| **CL 20:5_20:5_24:0_17:1** | C90H154O17P2 | CL | TRUE |
| **CL 17:2_20:5_24:0_20:3** | C90H156O17P2 | CL | TRUE |
| **CL 15:0_22:4_22:3_22:3** | C90H156O17P2 | CL | TRUE |
| **CL 12:0_18:0_28:0_22:3** | C89H168O17P2 | CL | FALSE |
| **CL 14:1_17:2_28:0_22:6** | C90H158O17P2 | CL | TRUE |
| **CL 18:3_20:5_26:0_17:1** | C90H158O17P2 | CL | TRUE |
| **CL 14:0_17:2_28:0_22:6** | C90H160O17P2 | CL | TRUE |
| **CL 12:0_14:1_26:0_28:0** | C89H172O17P2 | CL | FALSE |
| **CL 14:0_17:1_28:0_22:6** | C90H162O17P2 | CL | TRUE |
| **CL 15:0_22:6_28:0_16:1** | C90H162O17P2 | CL | TRUE |
| **CL 18:0_17:1_24:0_22:5** | C90H164O17P2 | CL | TRUE |
| **CL 14:0_17:0_28:0_22:6** | C90H164O17P2 | CL | TRUE |
| **CL 15:0_18:2_28:0_20:4** | C90H164O17P2 | CL | TRUE |
| **CL 18:2_18:2_24:0_22:6** | C91H158O17P2 | CL | TRUE |
| **CL 15:0_17:2_28:0_22:6** | C91H162O17P2 | CL | TRUE |
| **CL 17:0_22:3_22:5_22:6** | C92H152O17P2 | CL | TRUE |
| **CL 17:2_22:4_22:3_22:4** | C92H154O17P2 | CL | TRUE |
| **CL 16:0_24:0_20:3_22:3** | C91H166O17P2 | CL | TRUE |
| **CL 18:0_18:1_26:0_20:4** | C91H168O17P2 | CL | TRUE |
| **CL 12:0_18:3_24:0_28:0** | C91H172O17P2 | CL | TRUE |
| **CL 15:0_20:4_28:0_20:4** | C92H164O17P2 | CL | TRUE |
| **CL 15:0_18:2_28:0_22:6** | C92H164O17P2 | CL | TRUE |
| **CL 15:0_18:1_28:0_22:6** | C92H166O17P2 | CL | FALSE |
| **CL 17:0_24:0_17:0_24:0** | C91H178O17P2 | CL | TRUE |
| **CL 17:0_18:1_28:0_20:5** | C92H168O17P2 | CL | TRUE |
| **CL 17:0_18:2_28:0_20:3** | C92H170O17P2 | CL | TRUE |
| **CL 16:0_17:0_28:0_22:4** | C92H172O17P2 | CL | TRUE |
| **CL 15:0_18:0_28:0_22:4** | C92H172O17P2 | CL | TRUE |
| **CL 12:0_22:4_28:0_22:6** | C93H162O17P2 | CL | FALSE |
| **CL 17:0_18:0_28:0_20:3** | C92H174O17P2 | CL | TRUE |
| **CL 18:0_20:4_24:0_22:5** | C93H164O17P2 | CL | TRUE |
| **CL 12:0_15:0_28:0_28:0** | C92H180O17P2 | CL | FALSE |
| **CL 15:1_22:6_26:0_22:6** | C94H158O17P2 | CL | TRUE |
| **CL 17:0_22:6_24:0_22:6** | C94H160O17P2 | CL | TRUE |
| **CL 15:0_20:5_28:0_22:6** | C94H162O17P2 | CL | TRUE |
| **CL 18:3_22:6_28:0_17:2** | C94H162O17P2 | CL | TRUE |
| **CL 15:0_20:3_28:0_22:6** | C94H166O17P2 | CL | TRUE |
| **CL 15:0_20:3_28:0_22:5** | C94H168O17P2 | CL | FALSE |
| **CL 15:0_26:0_26:0_17:1** | C93H180O17P2 | CL | TRUE |
| **CL 17:0_18:1_28:0_22:6** | C94H170O17P2 | CL | TRUE |
| **CL 18:0_22:4_28:0_17:2** | C94H172O17P2 | CL | TRUE |
| **CL 14:0_22:6_28:0_22:6** | C95H162O17P2 | CL | TRUE |
| **CL 20:4_20:4_26:0_20:4** | C95H162O17P2 | CL | TRUE |
| **CL 18:0_22:4_28:0_17:1** | C94H174O17P2 | CL | TRUE |
| **CL 17:0_18:1_28:0_22:4** | C94H174O17P2 | CL | TRUE |
| **CL 17:0_18:1_28:0_22:3** | C94H176O17P2 | CL | TRUE |
| **CL 12:0_17:0_28:0_28:0** | C94H184O17P2 | CL | TRUE |
| **CL 15:1_22:6_28:0_22:6** | C96H162O17P2 | CL | TRUE |
| **CL 12:0_22:6_24:0_28:0** | C95H174O17P2 | CL | FALSE |
| **CL 12:0_24:0_28:0_22:4** | C95H178O17P2 | CL | TRUE |
| **CL 26:0_16:2_26:0_18:1** | C95H180O17P2 | CL | TRUE |
| **CL 15:1_22:4_28:0_22:4** | C96H170O17P2 | CL | TRUE |
| **CL 17:0_26:0_26:0_17:2** | C95H182O17P2 | CL | TRUE |
| **CL 17:0_20:3_28:0_22:5** | C96H172O17P2 | CL | TRUE |
| **CL 15:1_22:6_24:0_26:0** | C96H174O17P2 | CL | TRUE |
| **CL 15:0_22:3_28:0_22:3** | C96H176O17P2 | CL | TRUE |
| **CL 18:0_22:6_26:0_22:6** | C97H166O17P2 | CL | TRUE |
| **CL 15:0_20:5_24:0_28:0** | C96H178O17P2 | CL | TRUE |
| **CL 15:0_20:4_24:0_28:0** | C96H180O17P2 | CL | TRUE |
| **CL 14:1_17:2_28:0_28:0** | C96H182O17P2 | CL | TRUE |
| **CL 28:0_15:1_28:0_16:2** | C96H182O17P2 | CL | TRUE |
| **CL 20:3_22:3_24:0_22:3** | C97H172O17P2 | CL | TRUE |
| **CL 14:0_17:1_28:0_28:0** | C96H186O17P2 | CL | TRUE |
| **CL 22:5_22:5_28:0_17:2** | C98H168O17P2 | CL | TRUE |
| **CL 17:0_22:4_28:0_22:6** | C98H172O17P2 | CL | TRUE |
| **CL 12:0_26:0_28:0_22:3** | C97H184O17P2 | CL | TRUE |
| **CL 17:0_22:3_28:0_22:5** | C98H176O17P2 | CL | TRUE |
| **CL 14:0_18:1_28:0_28:0** | C97H188O17P2 | CL | FALSE |
| **CL 22:3_22:3_28:0_17:1** | C98H178O17P2 | CL | TRUE |
| **CL 12:0_24:0_24:0_28:0** | C97H190O17P2 | CL | TRUE |
| **CL 17:0_22:3_28:0_22:3** | C98H180O17P2 | CL | TRUE |
| **CL 26:0_17:1_26:0_20:5** | C98H180O17P2 | CL | TRUE |
| **CL 18:1_22:5_28:0_22:5** | C99H172O17P2 | CL | TRUE |
| **CL 15:0_24:0_28:0_22:4** | C98H184O17P2 | CL | TRUE |
| **CL 18:0_22:4_28:0_22:6** | C99H174O17P2 | CL | FALSE |
| **CL 18:0_22:3_28:0_22:5** | C99H178O17P2 | CL | TRUE |
| **CL 16:0_28:0_28:0_17:1** | C98H190O17P2 | CL | TRUE |
| **CL 18:0_22:3_28:0_22:3** | C99H182O17P2 | CL | TRUE |
| **CL 24:0_16:2_28:0_22:3** | C99H184O17P2 | CL | TRUE |
| **CL 15:0_22:6_26:0_28:0** | C100H184O17P2 | CL | FALSE |
| **CL 20:3_22:3_28:0_22:6** | C101H174O17P2 | CL | TRUE |
| **CL 15:0_20:5_28:0_28:0** | C100H186O17P2 | CL | TRUE |
| **CL 20:3_22:4_28:0_22:4** | C101H176O17P2 | CL | TRUE |
| **CL 28:0_17:2_28:0_18:2** | C100H188O17P2 | CL | TRUE |
| **CL 28:0_17:1_28:0_18:3** | C100H188O17P2 | CL | TRUE |
| **CL 17:0_24:0_28:0_22:3** | C100H190O17P2 | CL | TRUE |
| **CL 14:1_22:6_28:0_28:0** | C101H184O17P2 | CL | TRUE |
| **CL 24:0_18:3_28:0_22:3** | C101H186O17P2 | CL | TRUE |
| **CL 14:0_22:5_28:0_28:0** | C101H188O17P2 | CL | TRUE |
| **CL 16:0_28:0_28:0_20:3** | C101H192O17P2 | CL | TRUE |
| **CL 14:0_22:3_28:0_28:0** | C101H192O17P2 | CL | TRUE |
| **CL 12:0_24:0_28:0_28:0** | C101H198O17P2 | CL | FALSE |
| **CL 17:0_28:0_28:0_20:5** | C102H190O17P2 | CL | TRUE |
| **CL 24:0_22:6_28:0_20:4** | C103H182O17P2 | CL | TRUE |
| **CL 16:2_22:6_28:0_28:0** | C103H186O17P2 | CL | TRUE |
| **CL 24:0_15:1_26:0_28:0** | C102H198O17P2 | CL | FALSE |
| **CL 16:1_22:6_28:0_28:0** | C103H188O17P2 | CL | TRUE |
| **CL 28:0_16:2_28:0_22:4** | C103H190O17P2 | CL | TRUE |
| **CL 24:0_22:3_24:0_24:0** | C103H196O17P2 | CL | TRUE |
| **CL 24:0_16:2_26:0_28:0** | C103H198O17P2 | CL | FALSE |
| **CL 24:0_14:1_28:0_28:0** | C103H200O17P2 | CL | FALSE |
| **CL 12:0_26:0_28:0_28:0** | C103H202O17P2 | CL | TRUE |
| **CL 24:0_22:6_28:0_22:5** | C105H184O17P2 | CL | TRUE |
| **CL 17:0_22:4_28:0_28:0** | C104H196O17P2 | CL | FALSE |
| **CL 20:4_22:6_26:0_28:0** | C105H186O17P2 | CL | TRUE |
| **CL 24:0_22:5_28:0_22:4** | C105H188O17P2 | CL | TRUE |
| **CL 18:1_22:6_28:0_28:0** | C105H192O17P2 | CL | FALSE |
| **CL 18:0_22:3_28:0_28:0** | C105H200O17P2 | CL | TRUE |
| **CL 24:0_16:2_28:0_28:0** | C105H202O17P2 | CL | TRUE |
| **CL 20:5_22:6_28:0_28:0** | C107H188O17P2 | CL | TRUE |
| **CL 20:4_22:6_28:0_28:0** | C107H190O17P2 | CL | TRUE |
| **CL 20:3_22:6_28:0_28:0** | C107H192O17P2 | CL | TRUE |
| **CL 20:3_22:4_28:0_28:0** | C107H196O17P2 | CL | TRUE |
| **CL 24:0_18:3_28:0_28:0** | C107H204O17P2 | CL | TRUE |
| **CL 22:6_22:6_28:0_28:0** | C109H190O17P2 | CL | TRUE |
| **CL 22:5_22:6_28:0_28:0** | C109H192O17P2 | CL | TRUE |
| **CL 22:4_22:6_28:0_28:0** | C109H194O17P2 | CL | TRUE |
| **CL 22:3_22:6_28:0_28:0** | C109H196O17P2 | CL | TRUE |
| **CL 28:0_17:2_28:0_28:0** | C110H212O17P2 | CL | TRUE |
| **Cer 34:1;2O** | C34H67NO3 | Cer_NS | TRUE |
| **Cer 18:1;2O/16:0** | C34H67NO3 | Cer_NS | TRUE |
| **Cer 36:2;2O** | C36H69NO3 | Cer_NS | TRUE |
| **Cer 18:2;2O/18:0** | C36H69NO3 | Cer_NS | TRUE |
| **Cer 36:1;2O** | C36H71NO3 | Cer_NS | TRUE |
| **Cer 18:1;2O/18:0** | C36H71NO3 | Cer_NS | TRUE |
| **Cer 34:2;2O** | C34H65NO3 | Cer_NS | TRUE |
| **Cer 18:2;2O/16:0** | C34H65NO3 | Cer_NS | TRUE |
| **Cer 36:0;2O** | C36H73NO3 | Cer_NDS | TRUE |
| **Cer 18:0;2O/18:0** | C36H73NO3 | Cer_NDS | TRUE |
| **Cer 40:1;2O** | C40H79NO3 | Cer_NS | TRUE |
| **Cer 20:1;2O/20:0** | C40H79NO3 | Cer_NS | TRUE |
| **Cer 38:1;4O** | C38H75NO5 | Cer_AP | TRUE |
| **Cer 21:0;3O/17:1;(2OH)** | C38H75NO5 | Cer_AP | TRUE |
| **Cer 38:2;2O** | C38H73NO3 | Cer_NS | TRUE |
| **Cer 18:2;2O/20:0** | C38H73NO3 | Cer_NS | TRUE |
| **Cer 38:1;2O** | C38H75NO3 | Cer_NS | TRUE |
| **Cer 18:1;2O/20:0** | C38H75NO3 | Cer_NS | TRUE |
| **Cer 38:0;2O** | C38H77NO3 | Cer_NDS | TRUE |
| **Cer 18:0;2O/20:0** | C38H77NO3 | Cer_NDS | TRUE |
| **Cer 42:2;2O** | C42H81NO3 | Cer_NS | TRUE |
| **Cer 18:1;2O/24:1** | C42H81NO3 | Cer_NS | TRUE |
| **Cer 40:2;2O** | C40H77NO3 | Cer_NS | TRUE |
| **Cer 18:1;2O/22:1** | C40H77NO3 | Cer_NS | TRUE |
| **Cer 18:1;2O/22:0** | C40H79NO3 | Cer_NS | TRUE |
| **Cer 41:3;2O** | C41H77NO3 | Cer_NS | TRUE |
| **Cer 18:2;2O/23:1** | C41H77NO3 | Cer_NS | TRUE |
| **Cer 41:2;2O** | C41H79NO3 | Cer_NS | TRUE |
| **Cer 18:1;2O/23:1** | C41H79NO3 | Cer_NS | TRUE |
| **Cer 41:1;2O** | C41H81NO3 | Cer_NS | TRUE |
| **Cer 18:1;2O/23:0** | C41H81NO3 | Cer_NS | TRUE |
| **Cer 41:0;2O** | C41H83NO3 | Cer_NDS | TRUE |
| **Cer 18:0;2O/23:0** | C41H83NO3 | Cer_NDS | TRUE |
| **PE 32:2** | C37H70NO8P | PE | TRUE |
| **PE 16:1_16:1** | C37H70NO8P | PE | TRUE |
| **PMeOH 34:1** | C38H73O8P | PMeOH | TRUE |
| **PMeOH 16:0_18:1** | C38H73O8P | PMeOH | TRUE |
| **PE-Cer 36:1;2O** | C38H77N2O6P | PE_Cer | TRUE |
| **PE-Cer 14:0;2O/22:1** | C38H77N2O6P | PE_Cer | TRUE |
| **PE 32:1** | C37H72NO8P | PE | TRUE |
| **PE 16:0_16:1** | C37H72NO8P | PE | TRUE |
| **Cer 42:3;2O** | C42H79NO3 | Cer_NS | TRUE |
| **Cer 18:2;2O/24:1** | C42H79NO3 | Cer_NS | TRUE |
| **Cer 18:1;2O/24:0** | C42H83NO3 | Cer_NS | TRUE |
| **Cer 42:1;2O** | C42H83NO3 | Cer_NDS | TRUE |
| **Cer 18:0;2O/24:1** | C42H83NO3 | Cer_NDS | TRUE |
| **PA 36:2** | C39H73O8P | PA | TRUE |
| **PA 18:1_18:1** | C39H73O8P | PA | TRUE |
| **PE O-34:2** | C39H76NO7P | EtherPE | TRUE |
| **PE O-16:1_18:1** | C39H76NO7P | EtherPE | TRUE |
| **PA 36:1** | C39H75O8P | PA | TRUE |
| **PA 18:0_18:1** | C39H75O8P | PA | TRUE |
| **PE O-34:1** | C39H78NO7P | EtherPE | TRUE |
| **PE O-16:0_18:1** | C39H78NO7P | EtherPE | TRUE |
| **PA 36:0** | C39H77O8P | PA | TRUE |
| **PA 18:0_18:0** | C39H77O8P | PA | TRUE |
| **Cer 43:3;2O** | C43H81NO3 | Cer_NS | TRUE |
| **Cer 18:2;2O/25:1** | C43H81NO3 | Cer_NS | TRUE |
| **Cer 43:2;2O** | C43H83NO3 | Cer_NS | TRUE |
| **Cer 18:1;2O/25:1** | C43H83NO3 | Cer_NS | TRUE |
| **Cer 18:1;2O/25:0** | C43H85NO3 | Cer_NS | TRUE |
| **Cer 43:1;2O** | C43H85NO3 | Cer_NDS | TRUE |
| **Cer 18:0;2O/25:1** | C43H85NO3 | Cer_NDS | TRUE |
| **PE 34:2** | C39H74NO8P | PE | TRUE |
| **PE 16:1_18:1** | C39H74NO8P | PE | TRUE |
| **PE 34:1** | C39H76NO8P | PE | TRUE |
| **PE 16:0_18:1** | C39H76NO8P | PE | TRUE |
| **PE 34:0** | C39H78NO8P | PE | TRUE |
| **PE 16:0_18:0** | C39H78NO8P | PE | TRUE |
| **Cer 44:3;2O** | C44H83NO3 | Cer_NS | TRUE |
| **Cer 18:1;2O/26:2** | C44H83NO3 | Cer_NS | TRUE |
| **Cer 44:2;2O** | C44H85NO3 | Cer_NS | TRUE |
| **Cer 18:1;2O/26:1** | C44H85NO3 | Cer_NS | TRUE |
| **PG 32:0** | C38H75O10P | PG | TRUE |
| **PG 16:0_16:0** | C38H75O10P | PG | TRUE |
| **PC 28:0** | C36H72NO8P | PC | TRUE |
| **PC 12:0_16:0** | C36H72NO8P | PC | TRUE |
| **PEtOH 36:1** | C41H79O8P | PEtOH | TRUE |
| **PEtOH 18:0_18:1** | C41H79O8P | PEtOH | TRUE |
| **PE O-36:1** | C41H82NO7P | EtherPE | TRUE |
| **PE O-16:0_20:1** | C41H82NO7P | EtherPE | TRUE |
| **PE 36:4** | C41H74NO8P | PE | TRUE |
| **PE 16:0_20:4** | C41H74NO8P | PE | TRUE |
| **PE 36:3** | C41H76NO8P | PE | TRUE |
| **PE 18:1_18:2** | C41H76NO8P | PE | TRUE |
| **PE 36:2** | C41H78NO8P | PE | TRUE |
| **PE 18:1_18:1** | C41H78NO8P | PE | TRUE |
| **PE 36:1** | C41H80NO8P | PE | TRUE |
| **PE 18:0_18:1** | C41H80NO8P | PE | TRUE |
| **PG 34:2** | C40H75O10P | PG | TRUE |
| **PG 16:1_18:1** | C40H75O10P | PG | TRUE |
| **PE O-38:7** | C43H74NO7P | EtherPE | TRUE |
| **PE O-16:1_22:6** | C43H74NO7P | EtherPE | TRUE |
| **PA 40:6** | C43H73O8P | PA | TRUE |
| **PA 18:0_22:6** | C43H73O8P | PA | TRUE |
| **PG 34:1** | C40H77O10P | PG | TRUE |
| **PG 16:0_18:1** | C40H77O10P | PG | TRUE |
| **PE O-16:1_22:4** | C43H78NO7P | EtherPE | TRUE |
| **PE O-38:5** | C43H78NO7P | EtherPE | TRUE |
| **PE O-18:1_20:4** | C43H78NO7P | EtherPE | TRUE |
| **PA 40:4** | C43H77O8P | PA | TRUE |
| **PA 18:0_22:4** | C43H77O8P | PA | TRUE |
| **PE O-16:0_22:4** | C43H80NO7P | EtherPE | TRUE |
| **PE O-38:4** | C43H80NO7P | EtherPE | TRUE |
| **PE O-18:0_20:4** | C43H80NO7P | EtherPE | TRUE |
| **PE 36:2;O** | C41H78NO9P | OxPE | TRUE |
| **PE 18:1_18:1;O** | C41H78NO9P | OxPE | TRUE |
| **PE O-38:1** | C43H86NO7P | EtherPE | TRUE |
| **PE O-18:0_20:1** | C43H86NO7P | EtherPE | TRUE |
| **SL 45:6;2O** | C45H79NO6S | SL | TRUE |
| **SL 15:0;O/30:6;O** | C45H79NO6S | SL | TRUE |
| **PE 36:1;O** | C41H80NO9P | OxPE | TRUE |
| **PE 18:1_18:0;O** | C41H80NO9P | OxPE | TRUE |
| **PE 38:6** | C43H74NO8P | PE | TRUE |
| **PE 16:0_22:6** | C43H74NO8P | PE | TRUE |
| **PC O-32:1** | C40H80NO7P | EtherPC | TRUE |
| **PE 38:5** | C43H76NO8P | PE | TRUE |
| **PE 16:0_22:5** | C43H76NO8P | PE | TRUE |
| **PE 18:0_20:4** | C43H78NO8P | PE | TRUE |
| **PE 38:4** | C43H78NO8P | PE | TRUE |
| **PE 16:0_22:4** | C43H78NO8P | PE | TRUE |
| **PE 38:3** | C43H80NO8P | PE | TRUE |
| **PE 18:0_20:3** | C43H80NO8P | PE | TRUE |
| **PG 36:4** | C42H75O10P | PG | TRUE |
| **PG 16:0_20:4** | C42H75O10P | PG | TRUE |
| **PE 38:2** | C43H82NO8P | PE | TRUE |
| **PE 18:1_20:1** | C43H82NO8P | PE | TRUE |
| **PG 36:3** | C42H77O10P | PG | TRUE |
| **PG 18:1_18:2** | C42H77O10P | PG | TRUE |
| **PG O-37:3** | C43H81O9P | EtherPG | TRUE |
| **PG O-19:2_18:1** | C43H81O9P | EtherPG | TRUE |
| **PE 38:1** | C43H84NO8P | PE | TRUE |
| **PE 18:0_20:1** | C43H84NO8P | PE | TRUE |
| **PG 16:0_20:2** | C42H79O10P | PG | TRUE |
| **PG 36:2** | C42H79O10P | PG | TRUE |
| **PG 18:1_18:1** | C42H79O10P | PG | TRUE |
| **PE O-40:7** | C45H78NO7P | EtherPE | TRUE |
| **PE O-18:1_22:6** | C45H78NO7P | EtherPE | TRUE |
| **PG 36:1** | C42H81O10P | PG | TRUE |
| **PG 18:0_18:1** | C42H81O10P | PG | TRUE |
| **PE O-40:6** | C45H80NO7P | EtherPE | TRUE |
| **PE O-18:2_22:4** | C45H80NO7P | EtherPE | TRUE |
| **PE O-40:5** | C45H82NO7P | EtherPE | TRUE |
| **PE O-18:1_22:4** | C45H82NO7P | EtherPE | TRUE |
| **PE O-40:4** | C45H84NO7P | EtherPE | TRUE |
| **PE O-18:0_22:4** | C45H84NO7P | EtherPE | TRUE |
| **PE 38:4;O** | C43H78NO9P | OxPE | TRUE |
| **PE 18:1_20:3;O** | C43H78NO9P | OxPE | TRUE |
| **PE-Cer 44:6;2O** | C46H83N2O6P | PE_Cer | TRUE |
| **PE-Cer 20:1;2O/24:5** | C46H83N2O6P | PE_Cer | TRUE |
| **PE 40:5** | C45H80NO8P | PE | TRUE |
| **PE 18:0_22:5** | C45H80NO8P | PE | TRUE |
| **PE 40:4** | C45H82NO8P | PE | TRUE |
| **PE 18:0_22:4** | C45H82NO8P | PE | TRUE |
| **PE 40:3** | C45H84NO8P | PE | TRUE |
| **PE 18:0_22:3** | C45H84NO8P | PE | TRUE |
| **PG 38:4** | C44H79O10P | PG | TRUE |
| **PG 18:0_20:4** | C44H79O10P | PG | TRUE |
| **PE 38:4;2O** | C43H78NO10P | OxPE | TRUE |
| **PE 20:4_18:0;2O** | C43H78NO10P | OxPE | TRUE |
| **PG 38:2** | C44H83O10P | PG | TRUE |
| **PG 18:1_20:1** | C44H83O10P | PG | TRUE |
| **PC O-35:1** | C43H86NO7P | EtherPC | TRUE |
| **PC O-17:0_18:1** | C43H86NO7P | EtherPC | TRUE |
| **PG 38:0** | C44H87O10P | PG | TRUE |
| **PG 19:0_19:0** | C44H87O10P | PG | TRUE |
| **PC 33:1;O** | C41H80NO9P | OxPC | TRUE |
| **PC 18:1_15:0;O** | C41H80NO9P | OxPC | TRUE |
| **SMGDG O-33:1** | C42H80O12S | EtherSMGDG | TRUE |
| **SMGDG O-17:0_16:1** | C42H80O12S | EtherSMGDG | TRUE |
| **PE 40:5;O** | C45H80NO9P | OxPE | TRUE |
| **PE 18:1_22:4;O** | C45H80NO9P | OxPE | TRUE |
| **PE O-42:4** | C47H88NO7P | EtherPE | TRUE |
| **PE O-20:0_22:4** | C47H88NO7P | EtherPE | TRUE |
| **PI 32:0** | C41H79O13P | PI | TRUE |
| **PI 16:0_16:0** | C41H79O13P | PI | TRUE |
| **PG O-40:5** | C46H83O9P | EtherPG | TRUE |
| **PG O-18:1_22:4** | C46H83O9P | EtherPG | TRUE |
| **PE 18:1_22:3;O** | C45H82NO9P | OxPE | TRUE |
| **PE 40:4;O** | C45H82NO9P | OxPE | TRUE |
| **PE 22:3_18:1;O** | C45H82NO9P | OxPE | TRUE |
| **PC 16:1_18:1;O** | C42H80NO9P | OxPC | TRUE |
| **PC 34:2;O** | C42H80NO9P | OxPC | TRUE |
| **PC 16:0_18:2;O** | C42H80NO9P | OxPC | TRUE |
| **PC 17:0_17:1;O** | C42H82NO9P | OxPC | TRUE |
| **PC 34:1;O** | C42H82NO9P | OxPC | TRUE |
| **PC 16:0_18:1;O** | C42H82NO9P | OxPC | TRUE |
| **PC 36:4** | C44H80NO8P | PC | TRUE |
| **PC 16:1_20:3** | C44H80NO8P | PC | TRUE |
| **PI 34:2** | C43H79O13P | PI | TRUE |
| **PI 16:0_18:2** | C43H79O13P | PI | TRUE |
| **PI 34:1** | C43H81O13P | PI | TRUE |
| **PI 16:0_18:1** | C43H81O13P | PI | TRUE |
| **PC O-38:6** | C46H82NO7P | EtherPC | TRUE |
| **PC O-16:2_22:4** | C46H82NO7P | EtherPC | TRUE |
| **PI 34:0** | C43H83O13P | PI | TRUE |
| **PI 16:0_18:0** | C43H83O13P | PI | TRUE |
| **PC 36:4;O** | C44H80NO9P | OxPC | TRUE |
| **PC 16:0_20:4;O** | C44H80NO9P | OxPC | TRUE |
| **HexCer 40:2;3O** | C46H87NO9 | HexCer_HS | TRUE |
| **HexCer 16:2;2O/24:0;O** | C46H87NO9 | HexCer_HS | TRUE |
| **SMGDG O-36:4** | C45H80O12S | EtherSMGDG | TRUE |
| **SMGDG O-16:0_20:4** | C45H80O12S | EtherSMGDG | TRUE |
| **SM 41:2;2O** | C46H91N2O6P | SM | TRUE |
| **SM 18:1;2O/23:1** | C46H91N2O6P | SM | TRUE |
| **PC O-38:2** | C46H90NO7P | EtherPC | TRUE |
| **PC O-18:1_20:1** | C46H90NO7P | EtherPC | TRUE |
| **SM 41:1;2O** | C46H93N2O6P | SM | TRUE |
| **SM 18:1;2O/23:0** | C46H93N2O6P | SM | TRUE |
| **PC 36:2;O** | C44H84NO9P | OxPC | TRUE |
| **PC 18:1_18:1;O** | C44H84NO9P | OxPC | TRUE |
| **PC 37:1** | C45H88NO8P | PC | TRUE |
| **PC 18:0_19:1** | C45H88NO8P | PC | TRUE |
| **PI 35:2** | C44H81O13P | PI | TRUE |
| **PI 17:1_18:1** | C44H81O13P | PI | TRUE |
| **SM 41:0;2O** | C46H95N2O6P | SM | TRUE |
| **SM 18:0;2O/23:0** | C46H95N2O6P | SM | TRUE |
| **PC 36:1;O** | C44H86NO9P | OxPC | TRUE |
| **PC 18:1_18:0;O** | C44H86NO9P | OxPC | TRUE |
| **PI 35:1** | C44H83O13P | PI | TRUE |
| **PI 18:0_17:1** | C44H83O13P | PI | TRUE |
| **PI 34:1;O** | C43H81O14P | OxPI | TRUE |
| **PI 16:0_18:1;O** | C43H81O14P | OxPI | TRUE |
| **SM 18:2;2O/24:1** | C47H91N2O6P | SM | TRUE |
| **SM 42:3;2O** | C47H91N2O6P | SM | TRUE |
| **SM 14:1;2O/28:2** | C47H91N2O6P | SM | TRUE |
| **HexCer 41:2;3O** | C47H89NO9 | HexCer_HS | TRUE |
| **HexCer 16:1;2O/25:1;O** | C47H89NO9 | HexCer_HS | TRUE |
| **PI 36:4** | C45H79O13P | PI | TRUE |
| **PI 16:0_20:4** | C45H79O13P | PI | TRUE |
| **SM 42:2;2O** | C47H93N2O6P | SM | TRUE |
| **SM 18:1;2O/24:1** | C47H93N2O6P | SM | TRUE |
| **PI 16:0_20:3** | C45H81O13P | PI | TRUE |
| **PI 36:3** | C45H81O13P | PI | TRUE |
| **PI 18:1_18:2** | C45H81O13P | PI | TRUE |
| **SM 18:1;2O/24:0** | C47H95N2O6P | SM | TRUE |
| **SM 42:1;2O** | C47H95N2O6P | SM | TRUE |
| **SM 18:0;2O/24:1** | C47H95N2O6P | SM | TRUE |
| **PC 38:1** | C46H90NO8P | PC | TRUE |
| **PC 18:0_20:1** | C46H90NO8P | PC | TRUE |
| **HexCer 41:0;3O** | C47H93NO9 | HexCer_HDS | TRUE |
| **PI 18:1_18:1** | C45H83O13P | PI | TRUE |
| **PI 36:2** | C45H83O13P | PI | TRUE |
| **PI 18:0_18:2** | C45H83O13P | PI | TRUE |
| **SM 42:0;2O** | C47H97N2O6P | SM | TRUE |
| **SM 18:0;2O/24:0** | C47H97N2O6P | SM | TRUE |
| **PI 36:1** | C45H85O13P | PI | TRUE |
| **PI 18:0_18:1** | C45H85O13P | PI | TRUE |
| **PC O-40:4** | C48H90NO7P | EtherPC | TRUE |
| **PC O-18:3_22:1** | C48H90NO7P | EtherPC | TRUE |
| **PI O-38:5** | C47H83O12P | EtherPI | TRUE |
| **PI O-18:1_20:4** | C47H83O12P | EtherPI | TRUE |
| **SM 43:3;2O** | C48H93N2O6P | SM | TRUE |
| **SM 23:2;2O/20:1** | C48H93N2O6P | SM | TRUE |
| **PC 38:4;O** | C46H84NO9P | OxPC | TRUE |
| **PC 18:0_20:4;O** | C46H84NO9P | OxPC | TRUE |
| **PI 37:4** | C46H81O13P | PI | TRUE |
| **PI 17:0_20:4** | C46H81O13P | PI | TRUE |
| **SM 43:2;2O** | C48H95N2O6P | SM | TRUE |
| **SM 18:1;2O/25:1** | C48H95N2O6P | SM | TRUE |
| **SM 43:1;2O** | C48H97N2O6P | SM | TRUE |
| **SM 18:1;2O/25:0** | C48H97N2O6P | SM | TRUE |
| **SM 43:0;2O** | C48H99N2O6P | SM | TRUE |
| **SM 18:0;2O/25:0** | C48H99N2O6P | SM | TRUE |
| **PC 40:6** | C48H84NO8P | PC | TRUE |
| **PC 18:0_22:6** | C48H84NO8P | PC | TRUE |
| **PC 40:5** | C48H86NO8P | PC | TRUE |
| **PC 20:1_20:4** | C48H86NO8P | PC | TRUE |
| **PI 38:6** | C47H79O13P | PI | TRUE |
| **PI 16:0_22:6** | C47H79O13P | PI | TRUE |
| **HexCer 43:3;3O** | C49H91NO9 | HexCer_HS | TRUE |
| **HexCer 21:1;2O/22:2;O** | C49H91NO9 | HexCer_HS | TRUE |
| **PI 38:5** | C47H81O13P | PI | TRUE |
| **PI 18:1_20:4** | C47H81O13P | PI | TRUE |
| **SM 44:3;2O** | C49H95N2O6P | SM | TRUE |
| **SM 18:1;2O/26:2** | C49H95N2O6P | SM | TRUE |
| **PI 18:1_20:3** | C47H83O13P | PI | TRUE |
| **PI 38:4** | C47H83O13P | PI | TRUE |
| **PI 18:0_20:4** | C47H83O13P | PI | TRUE |
| **SM 44:2;2O** | C49H97N2O6P | SM | TRUE |
| **SM 18:1;2O/26:1** | C49H97N2O6P | SM | TRUE |
| **PI 38:3** | C47H85O13P | PI | TRUE |
| **PI 18:0_20:3** | C47H85O13P | PI | TRUE |
| **SM 44:1;2O** | C49H99N2O6P | SM | TRUE |
| **SM 18:1;2O/26:0** | C49H99N2O6P | SM | TRUE |
| **HexCer 44:3;3O** | C50H93NO9 | HexCer_HS | TRUE |
| **PI O-40:5** | C49H87O12P | EtherPI | TRUE |
| **PI O-18:1_22:4** | C49H87O12P | EtherPI | TRUE |
| **PI 18:0_20:5;O** | C47H81O14P | OxPI | TRUE |
| **PI 38:5;O** | C47H81O14P | OxPI | TRUE |
| **PI 18:1_20:4;O** | C47H81O14P | OxPI | TRUE |
| **PI 38:4;O** | C47H83O14P | OxPI | TRUE |
| **PI 18:0_20:4;O** | C47H83O14P | OxPI | TRUE |
| **PC O-42:1** | C50H100NO7P | EtherPC | TRUE |
| **PC O-16:0_26:1** | C50H100NO7P | EtherPC | TRUE |
| **PI 40:6** | C49H83O13P | PI | TRUE |
| **PI 18:0_22:6** | C49H83O13P | PI | TRUE |
| **PI 40:5** | C49H85O13P | PI | TRUE |
| **PI 18:0_22:5** | C49H85O13P | PI | TRUE |
| **PI 40:4** | C49H87O13P | PI | TRUE |
| **PI 18:0_22:4** | C49H87O13P | PI | TRUE |
| **SM 46:2;2O** | C51H101N2O6P | SM | TRUE |
| **SM 20:1;2O/26:1** | C51H101N2O6P | SM | TRUE |
| **PC 42:2** | C50H96NO8P | PC | TRUE |
| **PC 18:1_24:1** | C50H96NO8P | PC | TRUE |
| **PI 40:3** | C49H89O13P | PI | TRUE |
| **PI 18:0_22:3** | C49H89O13P | PI | TRUE |
| **PC 42:1** | C50H98NO8P | PC | TRUE |
| **PC 24:0_18:1** | C50H98NO8P | PC | TRUE |
| **PI 38:4;2O** | C47H83O15P | OxPI | TRUE |
| **PI 18:0_20:4;2O** | C47H83O15P | OxPI | TRUE |
| **PC 42:0** | C50H100NO8P | PC | TRUE |
| **PC 16:0_26:0** | C50H100NO8P | PC | TRUE |
| **HexCer 46:2;3O** | C52H99NO9 | HexCer_HDS | TRUE |
| **HexCer 46:1;3O** | C52H101NO9 | HexCer_HS | TRUE |
| **HexCer 16:1;2O/30:0;O** | C52H101NO9 | HexCer_HS | TRUE |
| **PC 16:0_27:1** | C51H100NO8P | PC | TRUE |
| **PC 43:1** | C51H100NO8P | PC | TRUE |
| **PC 25:0_18:1** | C51H100NO8P | PC | TRUE |
| **PC 43:0** | C51H102NO8P | PC | TRUE |
| **PC 16:0_27:0** | C51H102NO8P | PC | TRUE |
| **PC 44:6** | C52H92NO8P | PC | TRUE |
| **PC 22:3_22:3** | C52H92NO8P | PC | TRUE |
| **PC O-46:12** | C54H86NO7P | EtherPC | TRUE |
| **PC O-24:6_22:6** | C54H86NO7P | EtherPC | TRUE |
| **PC 18:0_26:1** | C52H102NO8P | PC | TRUE |
| **PC 44:1** | C52H102NO8P | PC | TRUE |
| **PC 26:0_18:1** | C52H102NO8P | PC | TRUE |
| **PC 44:0** | C52H104NO8P | PC | TRUE |
| **PC 18:0_26:0** | C52H104NO8P | PC | TRUE |
| **PC 45:2** | C53H102NO8P | PC | TRUE |
| **PC 18:1_27:1** | C53H102NO8P | PC | TRUE |
| **PC O-45:2;1O** | C53H104NO8P | EtherOxPC | TRUE |
| **PC O-28:0_17:2;1O** | C53H104NO8P | EtherOxPC | TRUE |
| **PC 46:5** | C54H98NO8P | PC | TRUE |
| **PC 26:1_20:4** | C54H98NO8P | PC | TRUE |
| **PC 46:3** | C54H102NO8P | PC | TRUE |
| **PC 18:1_28:2** | C54H102NO8P | PC | TRUE |
| **PC 46:2** | C54H104NO8P | PC | TRUE |
| **PC 18:1_28:1** | C54H104NO8P | PC | TRUE |
| **PC 46:1** | C54H106NO8P | PC | TRUE |
| **PC 28:0_18:1** | C54H106NO8P | PC | TRUE |
| **PC 47:2** | C55H106NO8P | PC | TRUE |
| **PC 18:1_29:1** | C55H106NO8P | PC | TRUE |
| **PC 48:5** | C56H102NO8P | PC | TRUE |
| **PC 28:1_20:4** | C56H102NO8P | PC | TRUE |
| **PC 48:3** | C56H106NO8P | PC | TRUE |
| **PC 18:1_30:2** | C56H106NO8P | PC | TRUE |
| **PC 48:2** | C56H108NO8P | PC | TRUE |
| **PC 18:1_30:1** | C56H108NO8P | PC | TRUE |
| **PC 52:4** | C60H112NO8P | PC | TRUE |
| **PC 16:0_36:4** | C60H112NO8P | PC | TRUE |
| **PC 54:5** | C62H114NO8P | PC | TRUE |
| **PC 18:1_36:4** | C62H114NO8P | PC | TRUE |
| **PC 56:5** | C64H118NO8P | PC | TRUE |
| **PC 18:1_38:4** | C64H118NO8P | PC | TRUE |
| **PC 57:3** | C65H124NO8P | PC | TRUE |
| **PC 37:0_20:3** | C65H124NO8P | PC | TRUE |
| **PG 62:1** | C68H133O10P | PG | TRUE |
| **PG 37:0_25:1** | C68H133O10P | PG | TRUE |
| **CL 58:2** | C67H126O17P2 | CL | TRUE |
| **CL 29:0_29:2** | C67H126O17P2 | CL | TRUE |
| **CL 58:1** | C67H128O17P2 | CL | TRUE |
| **CL 26:0_32:1** | C67H128O17P2 | CL | TRUE |
| **CL 60:2** | C69H130O17P2 | CL | TRUE |
| **CL 24:0_36:2** | C69H130O17P2 | CL | TRUE |
| **CL 60:1** | C69H132O17P2 | CL | TRUE |
| **CL 24:0_36:1** | C69H132O17P2 | CL | TRUE |
| **CL 64:4** | C73H134O17P2 | CL | TRUE |
| **CL 24:0_40:4** | C73H134O17P2 | CL | TRUE |
